# Evolutionary innovation through fusion of sequences from across the tree of life

**DOI:** 10.1101/2025.08.30.672725

**Authors:** Rishabh R. Kapoor, Evelyn E. Schwager, Supanat Phuangphong, Emily L. Rivard, Chandrashekar Kuyyamudi, Suhrid Ghosh, Isobel Ronai, Cassandra G. Extavour

## Abstract

Novel genes arise through multiple mechanisms, including gene duplication, gene fusion and horizontal gene transfer (HGT). While HGT has increasingly been documented in animals, the post-transfer evolutionary fate of horizontally-acquired genes is less well understood. We hypothesized that fusion with endogenous sequences in animal genomes might generate what we call “HGT-chimeras”: genes with regions of non-metazoan and metazoan descent in the same open reading frame. To test this hypothesis, we developed a molecular phylogenetics pipeline that enables the identification of HGT-chimeras. We applied our pipeline to 319 high-quality annotated arthropod genomes and uncovered a high-confidence set of 274 HGT-chimeras corresponding to 104 independent origination events across diverse arthropods. HGT-chimeras contain intervals acquired from across the tree of life, and many likely originated via a gene duplication-based mechanism. To assess whether HGT-chimeras might be functionally important, we performed RT-PCR and Sanger sequencing of tissues from 20 arthropod species predicted to harbor HGT-chimeras in their genome. We found evidence for the expression of contiguous chimeric mRNAs for 36 of 41 tested HGT-chimeras across 18 of 20 different tested species. We also found evidence that HGT chimeras evolve under purifying selection and have acquired potentially functional domain architectures, consistent with the hypothesis that these genes are in active use and may participate in diverse biological processes. These results illuminate an underappreciated combinatorial mechanism underlying the origin of novel genes across the largest animal phylum, and suggest that interdomain sequence fusion can play important roles in animal biology and evolution.

**Significance:** Evolution forges novelty through the repurposing of available parts. Can recently acquired parts, previously foreign to an organism, be similarly repurposed? Applying a rigorous methodology to 319 genomes from arthropods, the largest phylum of animals, we uncover 104 novel genes that arose from the fusion of animal genes with fragments acquired via horizontal gene transfer from bacteria, viruses, fungi, and plants. RNA-Seq and RT-PCR across multiple species show that many of these novel genes are expressed as mRNAs. Many show signatures of evolutionary conservation and coherent domain architectures, suggesting that these chimeric genes may play important roles in diverse biological processes. These results reveal an understudied path to evolutionary innovation via “bricolage” of genes from across the tree of life.

## Introduction

Novel genes contribute to adaptation and essential biological functions across the tree of life (1, 2), and have emerged through mechanisms such as duplication (1–3), fission (1, 2), fusion (4–8), and *de novo* birth (9, 10). Among these mechanisms, horizontal gene transfer (HGT) allows organisms to acquire new genes in a non-Mendelian fashion from other species (3, 11–15) and is widely appreciated as a fundamental force of evolution in prokaryotes. (12) Beyond an initial burst of HGT during early stages of eukaryotic symbiogenesis, HGT was long assumed to be vanishingly rare and therefore unimportant in eukaryotic evolution (3, 11, 16). Arthropods, the phylum of animals that constitute at least 80% of all described animal species (17), are emerging as an exception to this expectation. Horizontally acquired genes from bacteria (13–15, 18–23), viruses (14, 24–26), fungi (14, 27, 28) and even plants (14, 29) have been identified in diverse arthropods. In many cases, these genes contribute adaptive functions ranging from ecological defense (23, 29) to metabolism (28, 30). A recent systematic screen for HGT across 218 insect genomes uncovered 741 independent HGT events, placing the rate of HGT on par with that of other rare novel gene formation events within this phylum (14).

Whether horizontally acquired genes retain similar gene structures and functions following transfer, or, like other kinds of novel genes (7, 31–34), rapidly diverge from their original forms, has received less attention than detection of the transfer phenomenon. Here we focus on gene fusion as a possible mechanism of post-transfer divergence. Fusion of endogenous gene sequences is a well-documented source of novelty in protein structure and function (4–8). We hypothesized that, as a result of the fusion of horizontally acquired sequences with pre-existing host genome sequences, animal genomes would contain “HGT-chimeras”: genes with at least one region horizontally transferred from a non-metazoan source in the same open reading frame as a region of ancient metazoan ancestry. To our knowledge, no large-scale systematic screen for HGT-chimeras has been conducted in multicellular eukaryotes, but a few examples have been identified idiosyncratically across diverse taxa (21, 35–38). For instance, we previously attributed the origin of *oskar*, a gene required for essential developmental processes in diverse insect species (39–42), to the formation of a HGT-chimera between a metazoan LOTUS domain (43) and a bacterial GDSL hydrolase domain (44, 45) in the last common ancestor of extant insects (46). In another example, a screen of 20 microbial genomes identified 37 putative interkingdom HGT-chimeras in 11 unicellular species, but did not examine multicellular eukaryotes (47). Here, we present the results of a systematic bioinformatic screen for HGT-chimeras across 319 arthropod genomes. We find evidence that HGT-chimeric genes are not only widespread across the tree of arthropods, but also display transcriptional and molecular evolutionary characteristics that suggest they play active roles in organismal biology and evolution.

## Results and Discussion

### Development of an HGT-chimera detection pipeline

We designed a computational pipeline to search for HGT-chimeras in arthropod genomes (SI Figure 1), which we describe in brief here and detail in SI Methods. The pipeline proceeds in four broad phases (SI Figure 1). In the first phase, we assembled a search set of 7,702,369 protein inputs from 319 RefSeq scaffold or chromosome-level genomes (SI Table 1) across eight arthropod classes (SI Figure 4A), excluding proteins from scaffolds <100 kb to reduce the influence of contamination. We clustered the initial search set to 610,348 proteins using MMSeqs2 to promote computational efficiency (48). In the second phase, we used the results of DIAMOND-BLASTp (49) against the NR database (50) to perform an initial screen for putative chimeric proteins. Non-arthropod BLASTp results were first used to partition query proteins into intervals of potentially distinct evolutionary history using a customized implementation of a previously described algorithm (51) (Figure 1A, SI Figure 2A-C, SI File 1). We hypothesized that intervals with a preponderance of hits in non-metazoan taxa and/or lower E-values for non-metazoan than for metazoan hits were of putative HGT ancestry (right-hand interval of Figure 1A), and that those whose lowest E-value hits were in other metazoan taxa or had a preponderance of metazoan hits were of ancient metazoan ancestry (left-hand interval of Figure 1A). We then subjected proteins containing at least one interval of putative HGT ancestry and at least one interval of metazoan ancestry to a second round of BLASTp, using each of the two types of intervals as separate queries to further examine potential HGT and metazoan ancestry annotations from the first round of BLASTp (SI Figure 2D-E). In the third phase, we removed potential confounders of HGT inference (ankyrin repeats and metazoan transposable elements, see SI Methods) and binned HGT-chimeras into groups that we call “similarity clusters” based on their predicted protein domain architecture, using a custom approach that required all sequences in the same cluster to have the same chimeric architecture as determined by BLASTp and hmmsearch (HMMER suite (52). Except where otherwise indicated, downstream analyses (Figures 2-4) were performed with a single representative sequence per similarity cluster. In the fourth and final phase of the pipeline, we assessed the support for HGT and metazoan ancestries of the distinct intervals of each cluster representative using maximum likelihood phylogenetic trees (IQ-TREE (53)) (Figure 1B, SI Figure 3). To improve the sensitivity of homology detection, we constructed these trees from the results of hmmsearch (52) using profile hidden Markov models (HMMs) built from arthropod BLASTp hits as queries, or from BLASTp hits directly when <2 unique, high-confidence (see SI Methods) arthropod BLASTp hits were recovered (E-value for both <1e-2). We considered proteins with at least one interval of metazoan origin, at least one interval of non-metazoan origin, and support for tree topologies with >70% support at the relevant node (bootstrap only considered for HGT intervals, see SI Figure 3), to be candidate HGT-chimeras: genes formed by horizontal sequence transfer followed by in-frame fusion.

**Figure 1.**
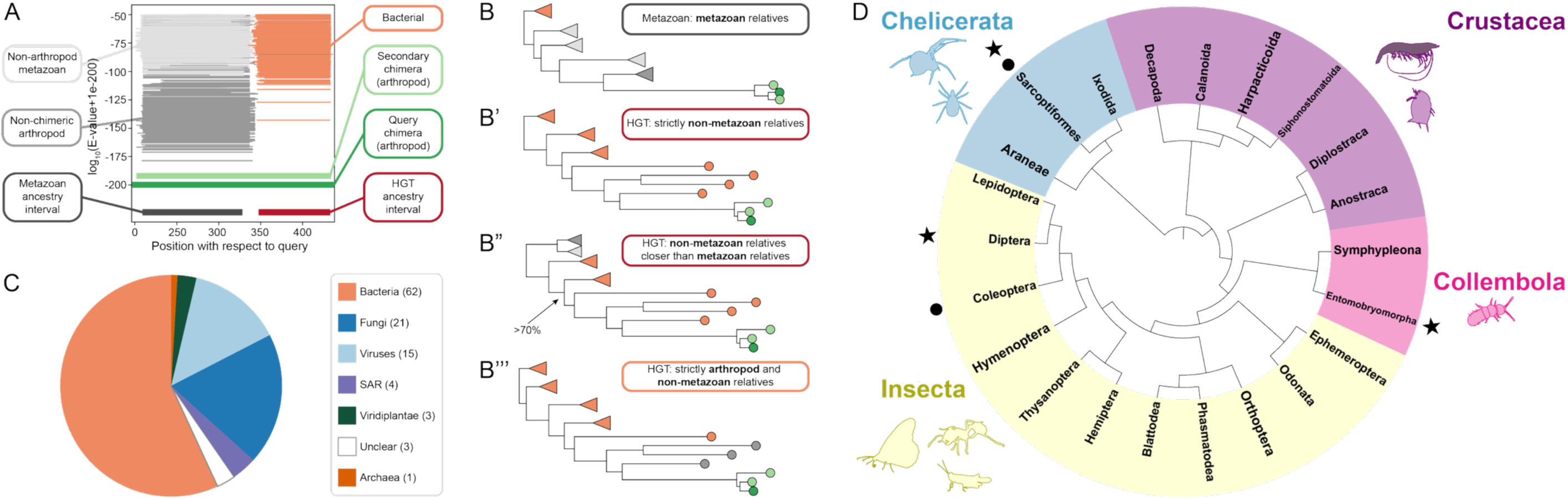
HGT-chimera detection across Arthropoda. **(A)** The taxonomic distribution of BLASTp hits across a query sequence are used to hypothesize chimeric ancestry. In this hypothetical example, the interval 1-325 amino acids is hypothesized to be of metazoan ancestry (darkest grey) because the closest non-arthropod hits are from non-arthropod metazoan taxa. The interval from 350-450 amino acids is hypothesized to be of HGT ancestry (red) because the closest hits are of non-metazoan origin. In addition to the original query sequence (dark green), secondary chimeric sequences of the same domain architecture (light green) are retrieved from other arthropods. Plots for all HGT-chimeras are available in SI File 1. **(B)** HGT and metazoan ancestries are further tested for each interval using maximum likelihood phylogenetics on hmmsearch or BLASTp hits as described in SI Methods. Circles and triangles represent single sequences and collapsed multi-sequence clades, respectively. **B** shows a hypothetical interval of metazoan ancestry due to the absence of close non-metazoan relatives of the chimeric sequences. **B’** shows the simplest case consistent with HGT from a non-metazoan donor, in which all detected phylogenetic relatives outside of other orthologous arthropod chimeras are from non-metazoan taxa. In **B’’**, non-chimeric metazoan relatives are detected, but an HGT origin is supported by nesting of the chimeric sequences within a clade of non-metazoan sequences with >70% bootstrap support. In **B’’’**, the closest relatives are non-chimeric arthropod sequences followed by non-metazoan sequences, supporting HGT from non-metazoans to arthropods prior to the chimera-forming fusion event. **(C)** Plausible HGT donor taxa at the domain/superkingdom level inferred from the taxonomic distribution of sister and cousin clades on maximum likelihood phylogenies for 109 HGT intervals from 104 HGT-chimera clusters (see SI Methods). SAR refers to the eukaryotic supergroup consisting of stramenopiles, alveolates, and rhizarians. **(D)** Arthropod orders with species containing at least one HGT-chimera. Colored ranges indicate taxonomic class (Insecta, Collembola) or subphylum (Chelicerata, Crustacea). Data include sequences from both the primary search set of 319 RefSeq genomes and 197 additional GenBank genomes (see SI Methods). Symbols outside the phylogram indicate the presence of HGT-chimeras in clusters 18 (circles) and 3 (stars), which are notable for their sparse phylogenetic distribution (SI Figure 6-7).

**Figure 2.**
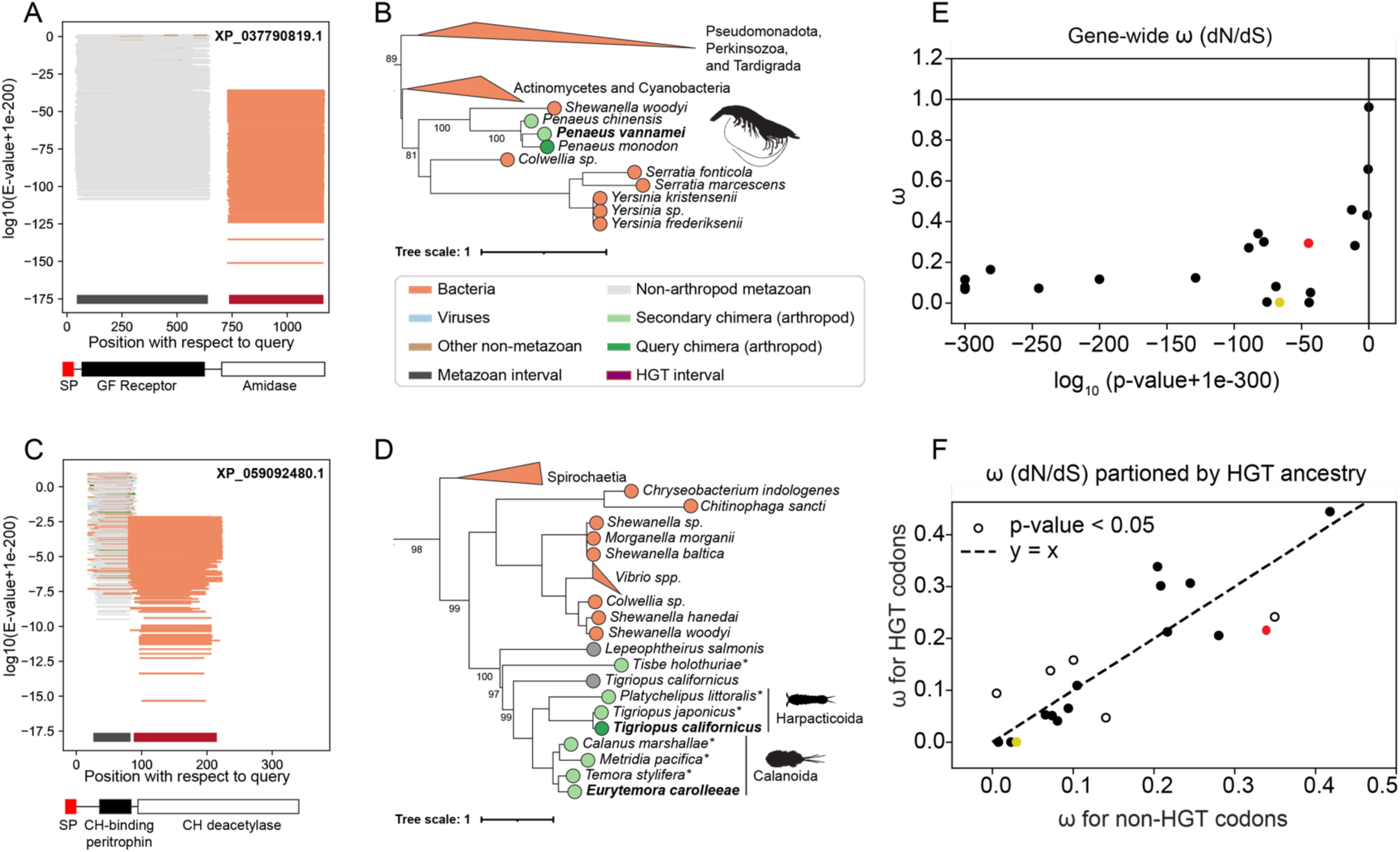
Crustacean chimeras of varying age have evolved under purifying selection. **(A)** BLASTp plot as in Figure 1A, but excluding arthropod hits, for the query chimera XP_037790819.1 from the shrimp *Penaeus monodon* (cluster 12). Metazoan ancestry is inferred for the interval (45, 640) and HGT ancestry for the interval (736, 1165). Domain annotations obtained via NCBI CDD search and InterProScan (SI Table 17) are shown across the bottom of the plot, with an N-terminal signal peptide predicted by SignalP-6.0 (cleavage site between positions 25-26). See SI Figure 11 for detailed annotation. **(B)** Maximum likelihood tree for the HGT interval (736, 1165) from XP_037790819.1, including intervals from secondary chimers found in the two other congeneric shrimp *P. vannamei* (XP_069990332.1) and *P. chinensis* (XP_047489075.1). **(C)** BLASTp plot excluding arthropod hits for the query chimera XP_059092480.1 (cluster 14) from the copepod *Tigriopus californicus*. Metazoan ancestry is inferred for the interval (26, 83) and HGT ancestry for the interval (88, 215). Domain annotations are shown across the bottom of the plot, with an N-terminal signal peptide cleaved between positions 29-30. See SI Figure 12 for detailed annotation. **(D)** Maximum likelihood tree for the HGT interval (88, 215), including secondary chimera sequences from the genome of the copepod *Eurytemora carolleeae* (XP_023343432.1) and additional sequences obtained from transcriptome shotgun assembly data from seven additional copepod species (indicated with an asterisk). The phylogenetic split among HGT-chimeras from the copepod orders Harpacticoida and Calanoida is indicated with vertical lines. In B and D, collapsed multi-sequence clades are represented as triangles. Triangle color represents the taxonomic origin of <u>the majority of</u> contained sequences, and the lengths of the two edges connecting each triangle to its parent node represents the summed branch length to the longest and shortest branch in the clade. Bolded text in B and D indicate sequences verified by RT-PCR, scale bars indicate one substitution per amino acid site, and numerical values at internal nodes indicate ultrafast bootstrap support values for select nodes relevant for HGT inference. Only a portion of both trees is shown here; the full trees for both metazoan and HGT trees with accessions are available on iTOL. **(E)** Gene-wide ω (dN/dS) values fitted for 21 of the 23 HGT-chimera similarity clusters that were found in the genomes of at least two different species (excluding clusters 3 and 18). The x-axis represents the log-transformed Benjamini-Hochberg corrected p-values from a likelihood ratio test comparing the likelihood of the free ω ratio values (y-axis) to a null model with ω fixed at 1. **(F)** Each of 23 HGT-derived intervals from the same 21 HGT-chimeras as in (E) were permitted to take a different ω value from the rest of the protein using fixed-site models. Benjami-Hochberg corrected p-values (indicated by dot fill) were obtained by comparing the likelihood of a model in which codons in the HGT interval differed from the remaining codons in their ω values; open dots indicate statistically significant deviation of dN/dS values for an HGT-derived interval from the rest of the coding region. Three intervals with y-axis values >0.50 and p_adj >0.05 have been ommited from this plot. Red and yellow circles in (E) and (F) highlight clusters 12 and 14, respectively.

### HGT-chimeras are detected across Arthropoda

Excluding possible within-genome paralogs (see SI Table 6 for copy numbers), we found a final set of 274 HGT-chimera genes across these query genomes, corresponding to 104 independent origination events inferred by similarity clustering (SI Table 2). We detected HGT-chimeras across the arthropod phylogeny (Figure 1D), including members of Chelicerata and Pancrustacea. A majority (193/319, 60.5%) of examined RefSeq genomes had at least one HGT-chimera (SI Figure 4B, SI Table 3). We note that *Folsomia candida*, the species with the greatest predicted number of HGT-chimeras, was previously reported to harbor large numbers of non-chimeric horizontally acquired genes (20, 27, 30).

To identify the broadest range of arthropod species containing each HGT-chimera (SI Table 4), we additionally searched for representatives of HGT-chimera similarity clusters in an expanded set of arthropod genomes, including an additional 197 genome annotations from GenBank (SI Table 1). We found that 81/104 similarity clusters were restricted to a single species (SI Figure 4C, SI Figure 5A), a result that may reflect a relatively recent evolutionary origin or the sparse genomic sampling of many arthropod taxa (SI text 1). For similarity clusters found in more than one species, we inferred minimum evolutionary gene ages by phylostratigraphy (54) (SI Figure 5B and SI Table 5). The youngest HGT-chimeras (clusters 20 and 21) appeared in the congeneric microcrustaceans *Daphnia pulex* and *D. pulicaria*, but were absent in the three other *Daphnia* genomes available at the time of writing. This suggests an origin for these HGT-chimeras as recently as 82,000 years ago in the *D. pulex*-*D. pulicaria* common ancestor (55) We detected a member of the similarity cluster corresponding to the gene *oskar* (cluster 1) in 181 insects spanning the subclass Pterygota, consistent with our prior finding that *oskar* originated via chimeric HGT in or prior to the last common ancestor of winged insects (>400 million years ago [MYA] (46)). Collectively, these findings suggest that HGT-chimera formation has occurred throughout arthropod evolution.

Unexpectedly, we found two clusters with members from multiple distantly related classes of arthropods (Figure 1D). Cluster 18 was found exclusively in the firefly *Photinus pyralis* (order: Coleoptera, class: Insecta) and in the mite *Oppia nitens* (order: Sarcoptiformes, class: Arachnida), a finding consistent with convergent evolution of analogous HGT-chimera architectures via independent HGT and gene fusion events (SI Figure 6). In the case of cluster 3, molecular phylogenetic analyses suggested that its sparse phylogenetic distribution could have arisen from the transfer of these HGT-chimeras between springtails, fungus gnats, and mites following the formation of this gene (SI Text 2, SI Figure 7). We note that inter-metazoan HGT, although thought to be rare (see for example 56–58), has nevertheless been reported by other studies (30, 58–60), including among these specific groups of arthropods (30, 61). Thus, HGT appears not only to have contributed to the origination of this HGT-chimera, but also to have contributed to its spread across species following its formation.

### HGT-chimeras are derived from diverse donor lineages

We used HGT interval trees (Figure 1B) to infer donor taxa and found that the majority of HGT intervals plausibly originated in bacterial donors (Figure 1C, SI Table 7). Among 62 sequences of inferred bacterial ancestry, close phylogenetic relatives of 37 of them are from bacterial genera of known arthropod symbionts (SI Table 8), including the endosymbionts *Wolbachia*, *Rickettsia* and *Hamiltonella*. This finding is consistent with prior reports of HGT between pathogenic and endosymbiotic bacteria and arthropod genomes (14, 18, 22, 46).

In addition to symbiotic bacterial donors, we observed 25 putative transfers from donors with plausible ecological associations with the recipient arthropods, including 15 cases of HGT from arthropod-infecting viruses, one from fungi to the fungus-consuming gnat *Bradysia coprophila*, and three from soil-dwelling bacteria to the soil-dwelling mite *Oppia nitens* (SI Table 8). In all, these data are consistent with the hypothesis that close ecological associations with food, pathogens and symbionts may provide opportunities for HGT (12, 18, 30, 37, 59).

### HGT-chimeras are transcribed

Nucleotide composition, codon usage and publicly available RNA-seq data suggested that most HGT-chimeras we predict from genome assemblies and annotations exist in their respective genomes and are expressed as functional gene products (SI Figure 8A-C, SI Table 11-12). For 41 chimeras from 20 species for which we could obtain tissue samples, we used RT-PCR and Sanger sequencing to test for the existence of continuous mRNA transcripts containing both predicted HGT and metazoan intervals in the predicted organisms (SI Figure 8D, SI Table 13, SI File 2). We detected transcripts of ∼88% (36/41) of examined HGT-chimeras, corresponding to 24 distinct similarity clusters across 18 species. This provides strong evidence that the observed HGT-chimeric transcripts are genuine rather than artifacts of genomic contamination, bioinformatic mis-assembly, or mis-annotation.

We additionally leveraged a variety of *in silico* tools to predict the domain architecture, structure, and function of HGT-chimeras (see Methods and SI Table 17). In the following sections, we use four RT-PCR-validated examples to illustrate the diverse origins, evolutionary trajectories, and predicted functions of these genes.

### A shrimp chimera illustrates that HGT-chimeras can rapidly come under the influence of purifying selection

HGT-chimera cluster 12 (representative sequence XP_037790819.1) is found in three species of the genus of marine shrimp *Penaeus* (order Decapoda). HGT intervals of these chimeras are phylogenetically nested within a clade of bacterial sequences with high statistical support (Figure 2B; tree topology tests in SI Figure 10), supporting a bacterial origin. The closest bacterial relative of the chimera clade is from the marine genus *Shewanella*, members of which are *Penaeus* pathogens (62), suggesting plausible ecological contact. The HGT interval is a predicted amidase domain with intact catalytic residues (SI Figure 11), and the metazoan-derived interval is annotated as a putative growth factor receptor. This unusual domain architecture (Figure 2A, SI Figure 11A) juxtaposes a metazoan cell signaling domain with a bacterial hydrolase in a likely soluble and extracellular protein (DeepLoc (63) probabilities >0.76 and >0.71, respectively; SignalP (64) likelihood > 0.99).

The three shrimp-derived sequences are monophyletic in both the HGT and metazoan interval trees (Figure 2B), supporting a single origin of this chimera as recently as 83.1 MYA (SI Figure 5B) (65). We can further place an upper bound on this chimera’s age due to its absence in other examined decapod genomes, including *P. japonicus*, which diverged from its chimera-bearing congeneric relatives 114 MYA. Despite its recent origin, this chimera has come under the influence of purifying selection, with gene-wide dN/dS = 0.29 (likelihood ratio test [LRT] p-value 1.06E-45) and no evidence of relaxed constraint in HGT-derived vs non-HGT codons as assessed by fixed-site (partition) models.

Similar patterns hold across the broader set of 23 HGT-chimeras found in more than one species: all have gene-wide dN/dS < 1, with neutrality rejected in 21 cases (Figure 2E, SI Table 14). Only three clusters showed significant relaxation of constraint in HGT versus non-HGT codons after FDR correction (Figure 2F, SI Table 15). This paucity of relaxed constraint supports the interpretation that many HGT-chimeras are functionally expressed and that horizontally acquired regions contribute to their activity.

### A copepod chimera illustrates deep conservation and functional similarity among chimera intervals

We found representatives of cluster 14 (representative sequence XP_059092480.1) in the genomes of two species of aquatic microcrustaceans, *Eurytemora coralleeae* (order Calanoida) and *Tigriopus californicus* (order Harpacticoida), implying an origin at least as ancient as the last common ancestor of extant copepods ∼446 MYA (Figure 2D). Further supporting our inference of deep functional conservation, we observed a strong signature of purifying selection on this gene (dN/dS=0.00374, LRT p-value = 3.18E-67) and detected representatives of this cluster in transcriptomes of 6/7 additional copepod species across two orders.

The HGT intervals of this chimera are phylogenetically adjacent to sequences from aquatic bacterial genera (Figure 2D), including *Shewanella* (as in cluster 12 above) and *Vibrio*, the latter of which is a well-documented copepod symbiont (66, 67). We note that the HGT-derived region is a predicted chitin deacetylase with intact catalytic residues (Figure 2A, SI Figure 12A-D), and this enzyme has been implicated in the association of *Vibrio* with their chitinous copepod hosts (68). Strikingly, the metazoan-derived region is annotated as a peritrophin (SI Figure 12E), a class of extracellular chitin-binding proteins that have been implicated in copepod transcriptional responses to *Vibrio* (66). Future work should examine whether this chimera, like its metazoan and bacterial relatives, functions at the host-microbe interface (SI Text 3).

The shared chitin-interacting functionality in the HGT and metazoan intervals of this chimera illustrates a broader pattern. Over one third (38/104) of HGT-chimeras displayed putative functional similarity in their HGT- and metazoan intervals, including eight cases in which both HGT and metazoan intervals were predicted to bind to or chemically modify carbohydrates (SI Table 18). These observations, coupled with the repeated observation of fusions of functionally related genes in other studies (4, 47, 69), could mean that HGT-chimeras formed from functionally similar components may be more likely to encode beneficial functions, contributing to their fixation and long-term preservation under purifying selection (4).

### A mosquito chimera illustrates that preexisting non-chimeric HGT genes may contribute to HGT-chimera formation

HGT-chimera cluster 9 (representative sequence XP_021699539.1) is found in three closely related species of mosquitoes (order Diptera), suggesting a minimum age of 92 MYA (65). It consists of a metazoan C2H2 zinc finger region and three tandem HGT-intervals, all annotated by CENSOR (70) as fragments of the auxiliary (non-transposase) protein of fungal Harbinger transposons (Figure 3A). Tree topologies confirm a close relationship of the latter with fungal sequences, but display a complex interleaving of fungal, plant, and oomycete sequences---consistent with documented cross-taxon transposon mobility (37, 71, 72) ---that precludes exact donor inference (Figure 3B). Notably, the closest non-chimeric relatives of the HGT intervals are from multiple dipterans (analogous to Figure 1B’’’), including one that harbors the HGT-chimera itself (*Aedes albopictus)*. The observations suggest a multistep model for the evolution of this HGT-chimera (Figure 3C): 1. a horizontal transfer event in a dipteran ancestor resulted in a non-chimeric HGT gene, 2. the standalone HGT gene duplicated, 3. at least one resulting duplicate fused with a metazoan C2H2 zinc finger to form the HGT-chimera (possibly coincident with the duplication event, see SI Figure 9C-D). We found analogous evidence across the full set of HGT chimeras, suggesting that non-chimeric HGT genes often precede HGT-chimera formation, and that gene duplication may contribute to HGT-chimera origination (SI text 4, SI Figure 9).

**Figure 3.**
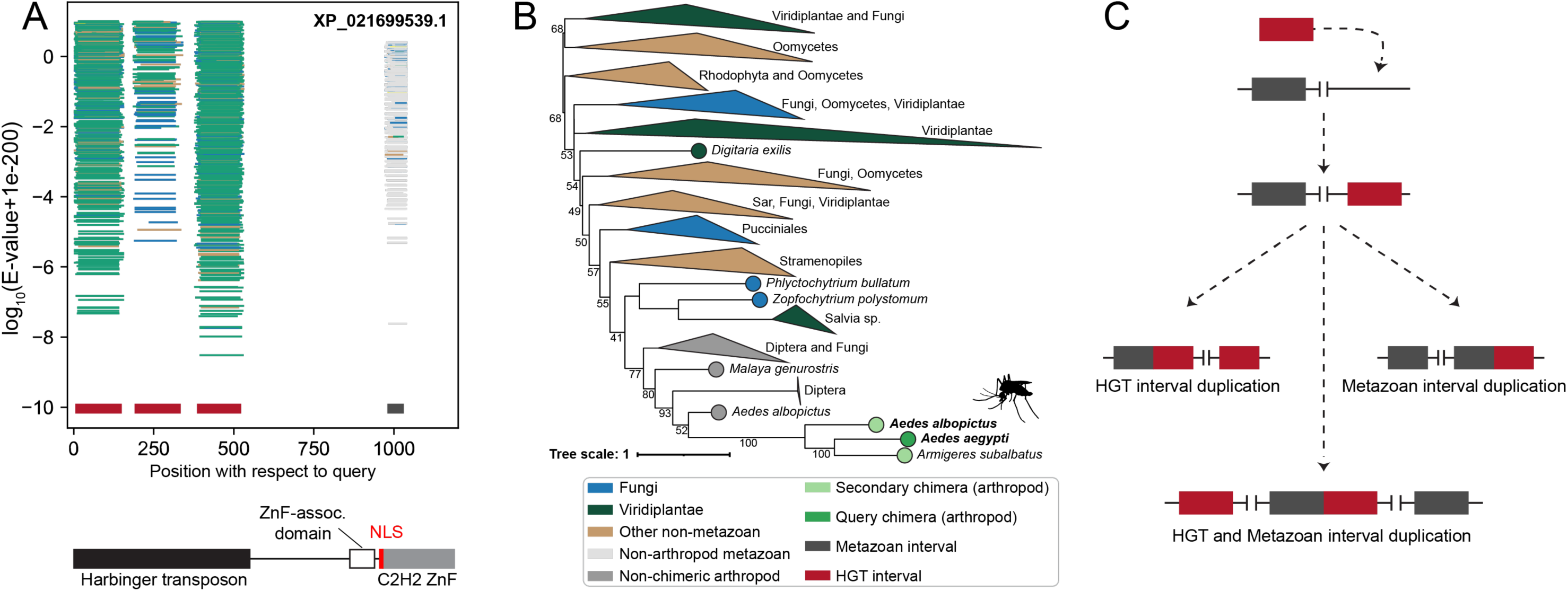
An HGT-chimera from mosquitoes illustrates that non-chimeric HGT can precede HGT-chimera formation. **(A)** BLASTp plot excluding arthropod hits for the query chimera XP_021699539.1 from the mosquito *Aedes aegypti* (cluster 9). Metazoan ancestry is inferred for the interval (980-1031) and HGT ancestry for the intervals (5-150), (190-334), (385-523). Domain annotations obtained via NCBI CDD search and InterProScan (SI Table 17) are shown across the bottom of the plot, along with a nuclear localization sequence from residues 975-985 as predicted by cNLS Mapper (96). See SI Figure 13 for detailed annotation. **(B)** Full maximum likelihood tree for the HGT interval (190,394), including secondary chimeras from the mosquitoes *Armigeres subalbatus* (XP_062544500.1) and *Aedes albopictus* (XP_029735553.1). Bolded text indicates that the chimera was successfully verified via RT-PCR. Collapsed clades are represented as in Figure 2. **(C)** Schematic representation of a model of HGT-chimera formation in which HGT precedes a duplication of HGT and/or metazoan genes. This model posits the existence of non-chimeric relatives to the HGT or metazoan intervals within the same genome or in the genomes of closely related arthropods. Note that the expected outcome of fusion via tandem duplication (bottom example) is a reversal of the order of metazoan and HGT segments on either side of the resulting chimera (97). See SI Figure 9 and SI Text 4 for supporting evidence.

Both auxiliary Harbinger proteins (73) and C2H2 zinc fingers (74) are DNA binding, and we predict intact DNA-binding residues in both the HGT- and metazoan-derived intervals of this HGT-chimera (SI Figure 13), along with an intact nuclear localization sequence (DeepLoc nuclear probability > 0.90). Coupled with gene-wide purifying selection (dN/dS=.27, p-value = 2.54E-90), these findings suggest that HGT-derived transposon fragments may have been co-opted into a transcriptional regulatory role, as has been found for other cases of transcription factor – transposon fusions (75, 76). We document 22 HGT chimeras in which both HGT and metazoan intervals are predicted to function in nucleic acid-associated processes (SI Table 18), further illustrating the recurrent pattern of functional similarity raised by the chitin-interacting copepod chimera above.

### A damselfly chimera illustrates the potential contribution of metazoan gene duplication and neofunctionalization to HGT-chimera evolution

Cluster 41 (XP_046403459.1) is restricted to a single species, the damselfly *Ischnura elegans* (order Odonata). It encodes a predicted transmembrane protein (DeepLoc probability > 0.99), containing a fragment of a metazoan anoctamin transmembrane transporter fused to a fragment of a bacterial ABC transporter likely derived from bacterial endosymbionts (Figure 4A-B). Using MembraneFold (77), we predicted two transmembrane helices of metazoan origin followed by a C-terminal HGT-derived transmembrane helix (Figure 4C). This chimera therefore illustrates the assembly of a novel transmembrane protein via fusion of parts of transmembrane proteins from different domains of life.

**Figure 4.**
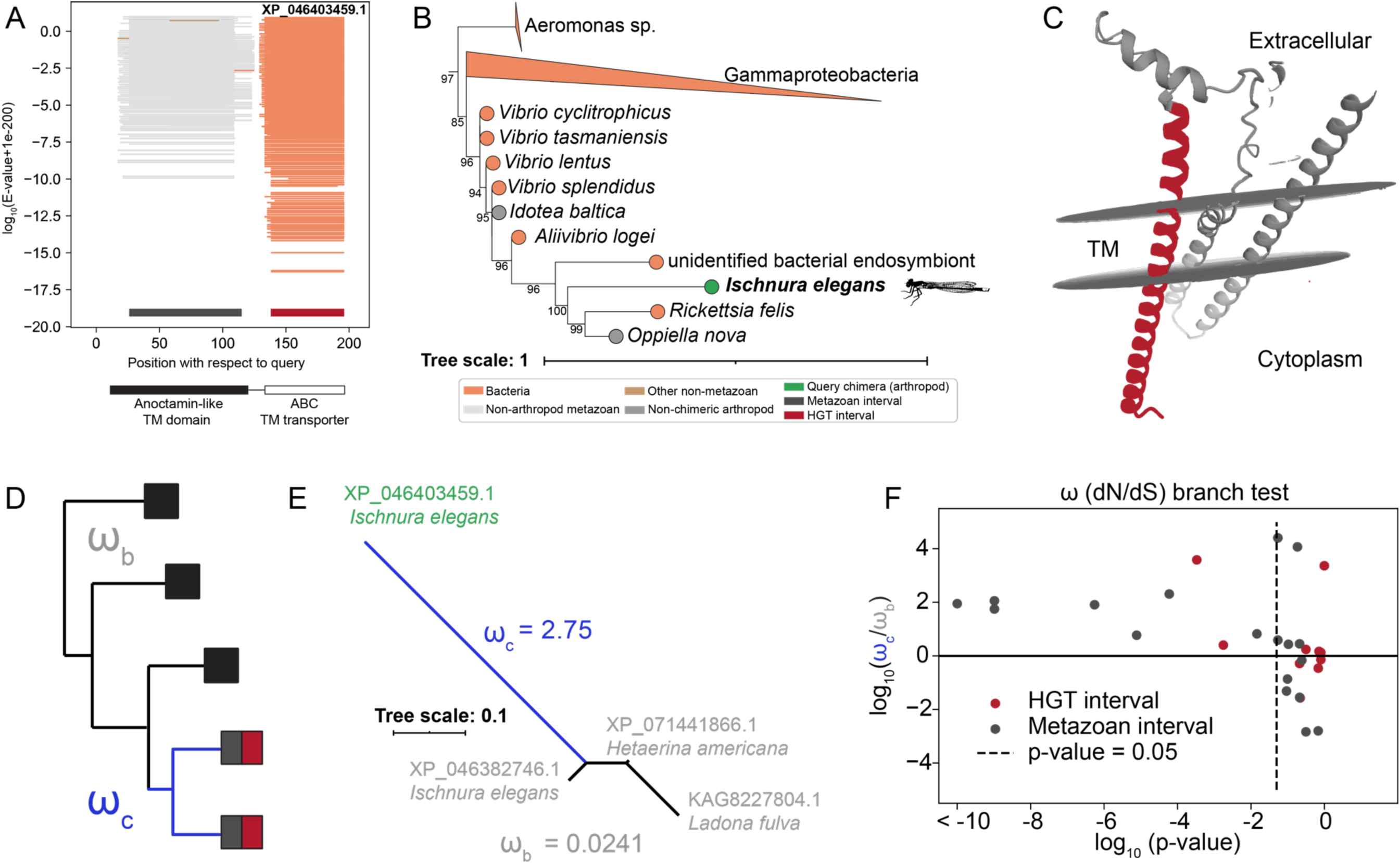
A transmembrane HGT-chimera from a damselfly illustrates post-fusion neofunctionalization. **(A)** BLASTp plot excluding arthropod hits for the query chimera XP_046403459.1 from the damselfly *Ischnura elegans* (cluster 41). Metazoan ancestry is inferred for the interval (26-115) and HGT ancestry for the interval (138-196). Domain annotations obtained via NCBI CDD search and InterProScan are shown across the top of the plot. **(B)** Maximum likelihood tree for the HGT interval (138-196), pruned to show sequences closely related to the query chimera (full tree available on the iTOL webserver at https://itol.embl.de/shared/rkapoor). Internal node values indicate ultrafast bootstrap support. Per GenBank, the “unidentified bacterial endosymbiont” was isolated from the midge *Chironomus riparius*, and *Rickettsia felis* from the louse *Liposcelis bostrychophila*. **(C)** Transmembrane protein structure and topology for XP_046403459.1 predicted with MembraneFold version 0.0.71 (77) with OmegaFold (98) and DeepTMHMM (99). The HGT-derived interval is colored in dark red. **(D)** Schematic representation of branch test performed to compare ω values of an interval before (gray) and after (blue) an HGT-chimera-forming gene fusion event. Separate gene trees were built for HGT and metazoan intervals using phylogenetic relatives within the same taxonomic order as HGT-chimeras. To best capture the early dynamics of HGT-chimera evolution, we restricted our analysis to 28 intervals from young (species or genus-restricted) HGT-chimeras in which relatives were detected both within the same genome as the HGT-chimera and in the genomes of other arthropods in the same taxonomic order (SI Methods). **(E)** Maximum likelihood tree constructed from the metazoan interval of the damselfly chimera XP_046403459.1 and its non-chimeric phylogenetic relatives in the same taxonomic order (Odonata), including a non-chimeric relative gene in the same genome as the chimera (XP_046382746.1). ω (dN/dS) values from the branch test explained in (D) are superimposed for chimeric and non-chimeric branches. **(F)** The distribution of the fold-change in ω between chimeric and non-chimeric branches of gene trees is plotted on the y-axis, for 28 HGT-chimera intervals. Benjamini-Hochberg corrected p-values (x-axis) were obtained using a likelihood ratio test of a two-branch model to a model with a single ω value.

The metazoan interval tree further shows that the closest relative of this chimera is a non-chimeric anoctamin gene in the *I. elegans* genome (Figure 4D). Thus, paralleling our hypothesis that duplication of preexisting non-chimeric HGT genes contributes to HGT-chimera origination, we posit that duplication of a metazoan anoctamin gene facilitated the fusion of one duplicate with an HGT gene (Figure 3C). We observed a significant elongation of the branch of this tree leading to the HGT-chimera relative to the remaining damselfly branches, suggesting that the fusion event was followed by accelerated sequence evolution. Consistently, we fit a two-ratio dN/dS model (78–80) to this metazoan interval tree and found that the non-chimeric metazoan sequences have evolved under strong purifying selection (dN/dS=.0241), while the HGT-chimera diverged under the influence of strong positive selection (dN/dS = 2.75, LRT p-value = 1.1E-10).

More broadly, across species- or genus-specific HGT-chimera intervals with non-chimeric phylogenetic relatives in the same taxonomic order, 64.3% (18/28) have higher dN/dS on the chimeric branches. A higher dN/dS on chimeric vs. non-chimeric branches is statistically supported by an LRT in 32.1% (9/28) of such intervals (Figure 4E-F, SI Table 16). This suggests that accelerated protein sequence evolution may be a common fate of incipient HGT-chimeras, consistent with the functional divergence (neofunctionalization) observed in other novel genes that have evolved through gene fusion and/or duplication (4, 7, 8, 31, 34, 81).

Post-fusion functional divergence has been experimentally shown for the HGT-derived domain of the HGT-chimeric gene *oskar*. The closest bacterial relatives of *oskar*’s HGT-derived domain, called the OSK domain (44), are lipid hydrolases (40, 45, 46), but in insects this domain lacks key hydrolytic residues and instead binds mRNAs (44, 82). Thus, HGT-chimeras reflect evolutionary innovation not by only the juxtaposition of pre-existing sequences from different domains of life, but also through post-fusion sequence divergence.

## Conclusions

Novel protein structure and function can emerge from several elementary events, including amino acid substitutions, gene duplication, gene fusion and HGT (3). In this study, we have shown that multiple of these elementary events can operate together in the origins of a previously understudied class of genes that we call HGT-chimeras. In at least 104 independent events across the history of the arthropods, HGT-chimeras evolved via the fusion of endogenous genes of ancient metazoan ancestry with genes acquired via horizontal transfer from non-metazoan sources. Sequence duplication within the host genome played a plausible role in the formation of many HGT-chimeras (Figure 3B-C, Figure 4D, SI Figure 9), and we additionally detected a signature of accelerated amino acid substitution following HGT-chimera birth (figure 4D). Collectively, these findings reveal a rich post-transfer history of horizontally acquired sequences in arthropods.

We propose that although chimeric genes of this nature have been reported rarely (21, 35–38, 46, 47) despite significant prior work on HGT (12–16, 18, 19, 22, 24–29), we were able to uncover these HGT-chimeras by better demarcating intervals of distinct history (Figure 1A, SI Figure 2). HGT-chimeras are unlikely to be detected by traditional HGT detection methods, which assume a common evolutionary history throughout the length of the open reading frame. In almost half (41/104) of HGT-chimeras, a naive interpretation of gene-wide BLASTp results via the same criteria used to infer interval ancestry would lead to a false inference of gene-wide metazoan ancestry (see SI Table 19). For the remaining genes, constructing a single gene tree for genic intervals with distinct histories would be methodologically problematic and, even if it resulted in a sensible topology, would obscure the true chimeric history of the gene. We expect that the application of methods like the one we describe here, that allow for intragenic phylogenetic discordance, could lead to the discovery of more HGT-chimeras, especially in lineages with frequent HGT such as grasses and parasitic plants (83, 84), fungi (85, 86) and prokaryotes (3, 12).

The biological significance of HGT has been questioned by some researchers, as many transferred segments show evidence of pseudogenization (15, 22, 87), and significant barriers to the transfer and functional integration of genes from divergent lineages have been posited (11, 16, 88–90). Motivated by the functionally coherent fusions discussed in the insect examples above Figures 2-4), we speculate that fusion with pre-existing gene sequences might facilitate the integration of novel horizontally transferred sequences into pre-existing functional networks. Many HGT-chimeras show signatures consistent with biological function, including multiple lines of evidence for active transcription (SI Figure 8, SI Tables 12-13) and gene-wide purifying selection (Figure 2E-F, SI Tables 14-15). Experimental studies in diverse model systems have demonstrated important functions for many young and species-specific genes (6, 10, 33, 91–93). Thus, we consider it plausible that even the young HGT-chimeras detected here may play important biological roles. Our *in silico* functional predictions (SI Tables 17-18) provide insights into the potential range of biological processes impacted by HGT-chimeras, warranting experimental validation in future studies.

To conclude, our novel computational approach has revealed that fusion of horizontally acquired sequence with endogenous ones has been widespread throughout the history of arthropods, illustrating a previously underappreciated versatility in evolutionary innovation via genomic “bricolage” (94).

## Methods

Software used in the “Development of an HGT-chimera detection pipeline” section included HMMER v3.3.2 (52), MUSCLE v5.1 (99), trimAl (100), and iTOL v5 (95). dN/dS ratios were computed with PAL2NAL v14.1 (101) and PAML v4.10.0 (68–70). All BLASTp searches were performed with DIAMOND v2.0.15 (49). RT-PCR and Sanger sequencing were performed on arthropod samples from donations or lab cultures (SI Table 13). Full methods are provided in SI.

## Supporting information

ST Tables 1 through 21

## Acknowledgements

We are grateful to Gregg Thomas, Sean Eddy, Andrew W. Murray, Gonzalo Giribet and the Extavour lab for critical feedback and discussion. Computational resources were provided and maintained by Harvard Faculty of Arts and Sciences Research Computing. Arthropod specimens for RT-PCR were generously donated by the researchers listed in SI Table 13. A special thank you to Avinash Tope and Andrew Ray (Kentucky State University) for allowing us to use their lab space and materials for *Cherax quadricarinatus* and *Penaeus vannamei* sample preparation, to John Stivers for assistance with the dissection of these specimens, to Yu Shirai (Extavour lab) for *Tribolium castaneum* RT-PCR, and to Tarun Kumar (Extavour lab) for *Bombyx mori* dissection and extraction of *Tyrophagus putrescentiae* cDNA.

## Data availability

All data required to reproduce associated figures is included in Supporting Information tables. Maximum likelihood trees for HGT and metazoan intervals of the final set of HGT-chimera similarity clusters are freely accessible in the public project “Arthropod HGT-chimera interval trees 8/27/2025” on the iTOL webserver at https://itol.embl.de/shared/rkapoor. Associated BLAST results, hmmsearch results, multiple sequence alignments, and Newick trees have been deposited at the Dryad digital repository for this study https://doi.org/10.5061/dryad.t1g1jwtdz. Scripts used to implement the HGT-chimera detection pipeline and downstream analysis are available on the github repository https://github.com/rishabhrajkapoor/Arthropod-HGT-chimeras-2025 with commit ID 6a974df. Sanger-verified cDNA sequences have been deposited at GenBank, with accession numbers available in SI Table 13.

## Author contributions

Project was conceptualized by CGE and RK. RK performed bioinformatic analyses. SP, EES and IR prepared cDNA libraries. RT-PCRs were performed by EES, ELR, CK, SG, SP, and IR. EES inspected and manually corrected Sanger sequencing results. RK, EES, ELR, and CGE wrote the manuscript and created associated figures. All authors reviewed the manuscript.

