## Supplementary material for "Evolutionary innovation through fusion of sequences from across the tree of life": SI Files 1 and 2: HGTc_SI_PDF_1_v3.pdf

Drosophila innubila GCF_004354385.1;XP_034487048.1 (cluster 1)

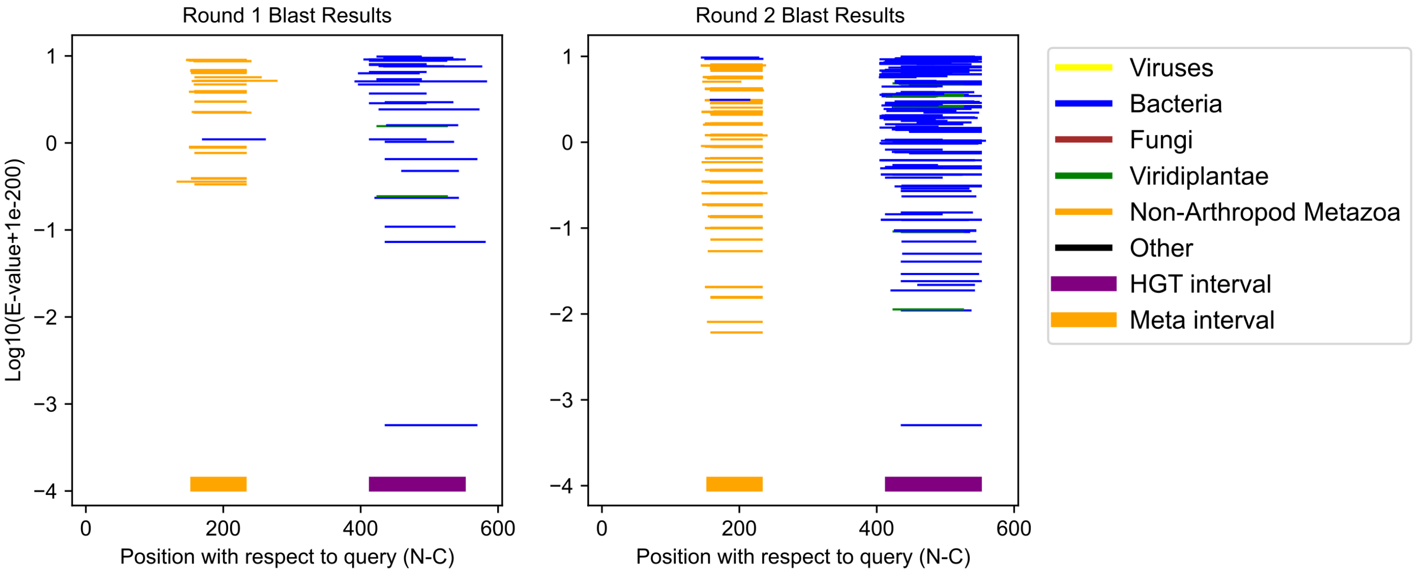

Homarus americanus GCF_018991925.1;XP_042220148.1 (cluster 2)

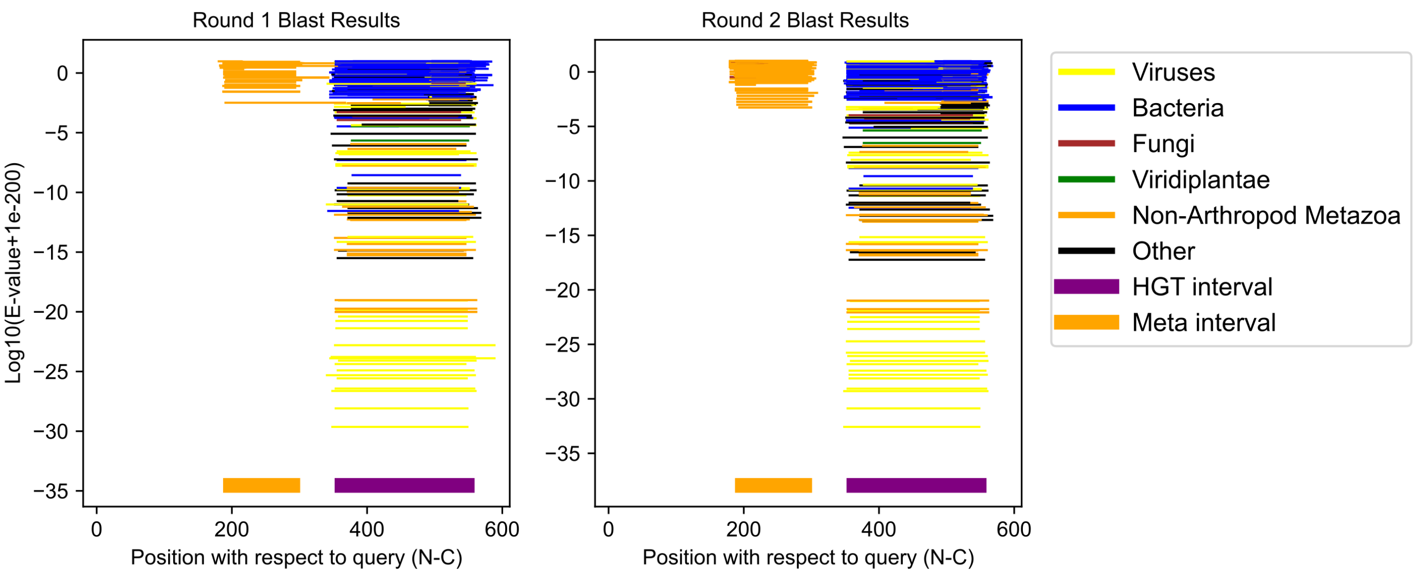

Folsomia candida GCF_002217175.1;XP_035715507.1 (cluster 3)

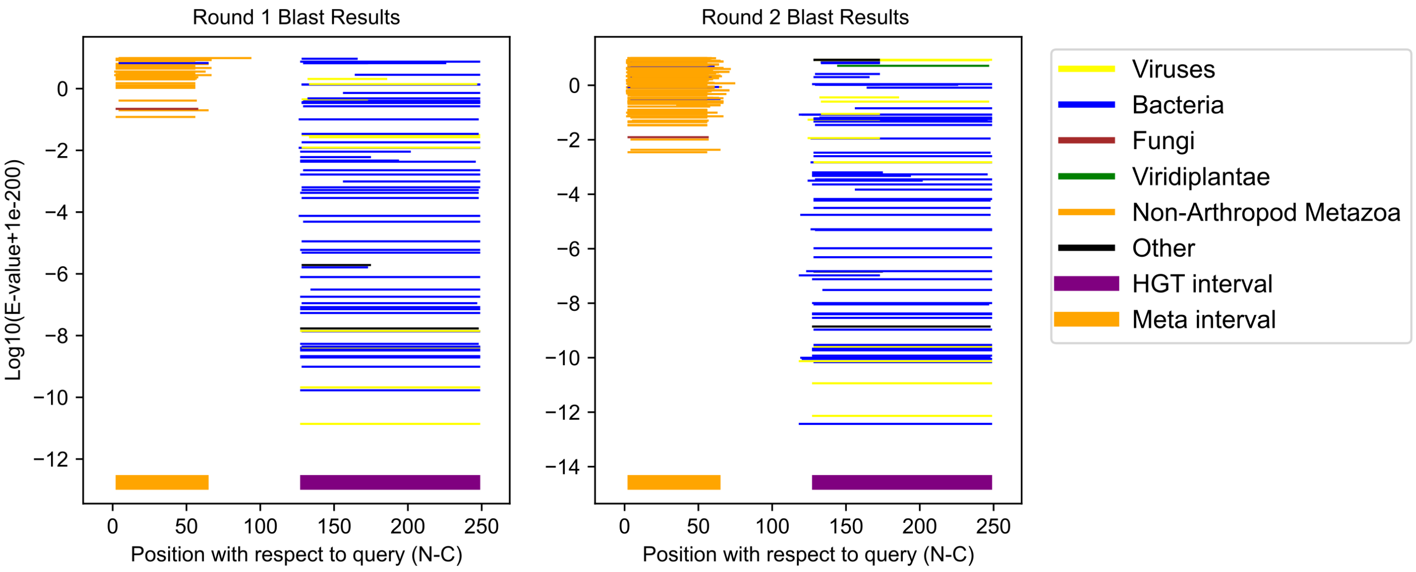

Coccinella septempunctata GCF_907165205.1;XP_044763649.1 (cluster 4)

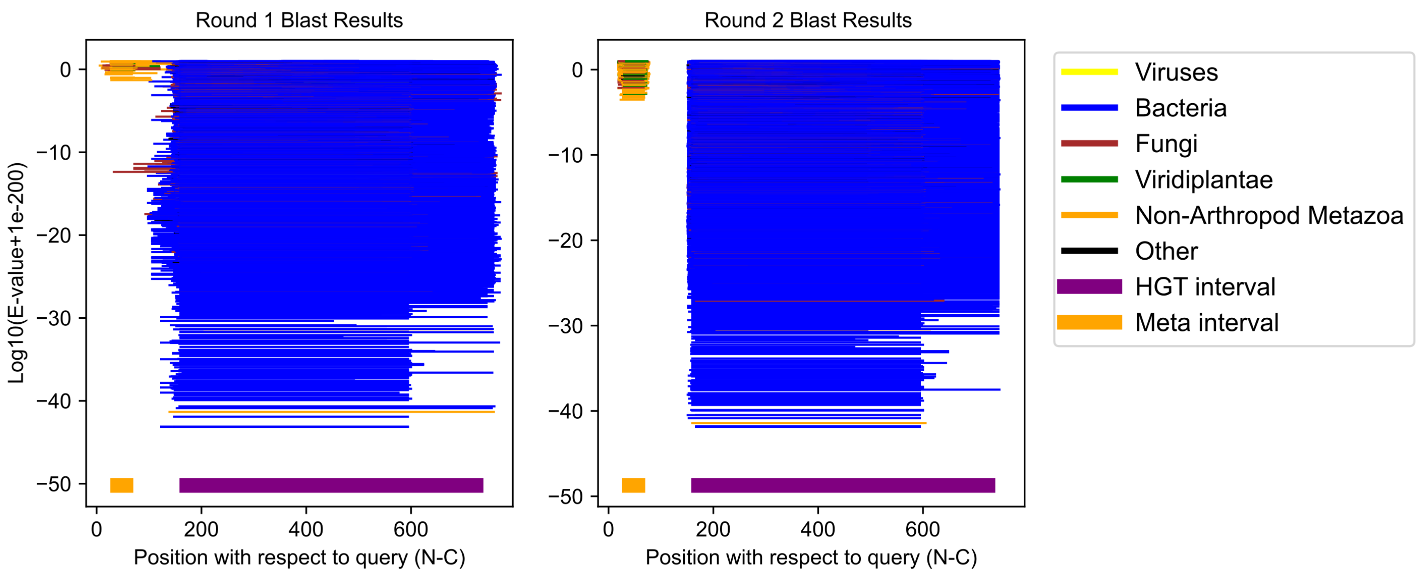

Oppia nitens GCF_028296485.1;XP_054161842.1 (cluster 5)

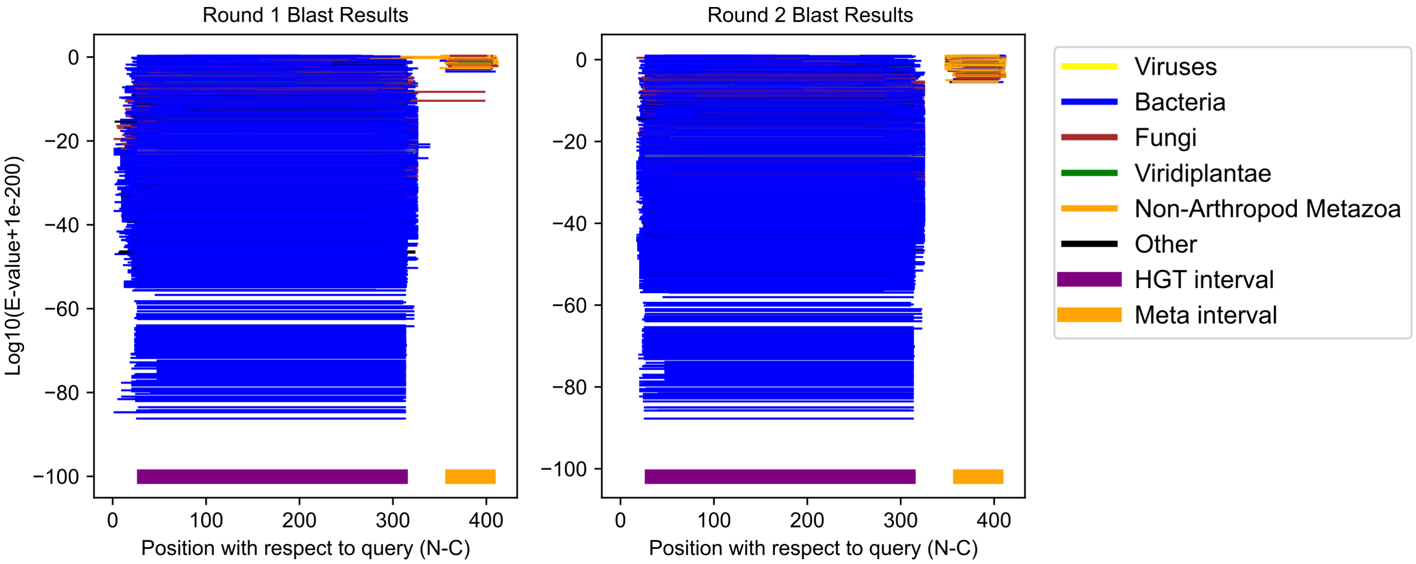

Daphnia pulicaria GCF_021234035.1;XP_046649021.1 (cluster 6)

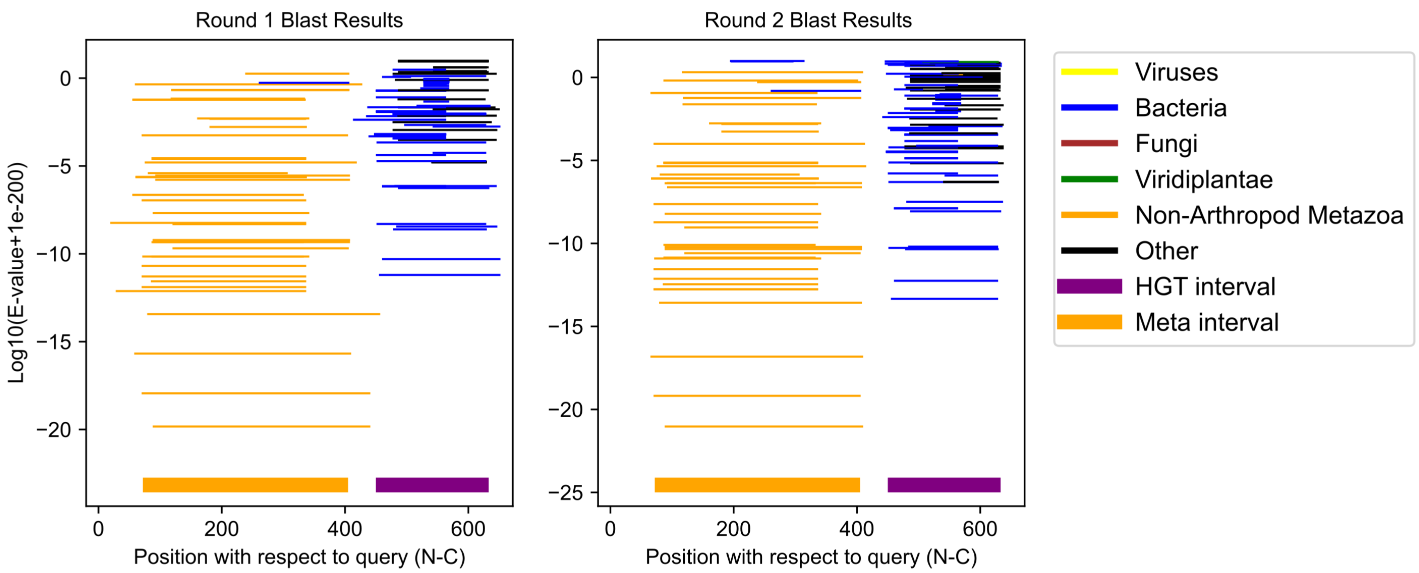

Coccinella septempunctata GCF_907165205.1;XP_044750704.1 (cluster 7)

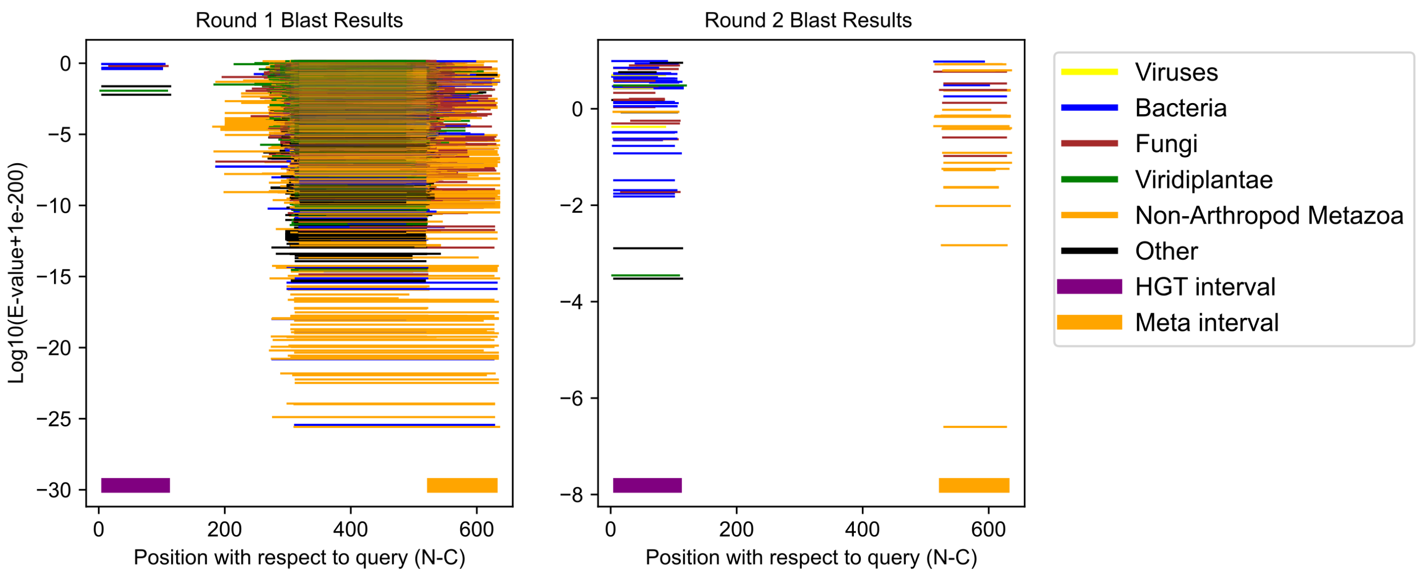

Daphnia pulex GCF_021134715.1;XP_046456339.1 (cluster 8)

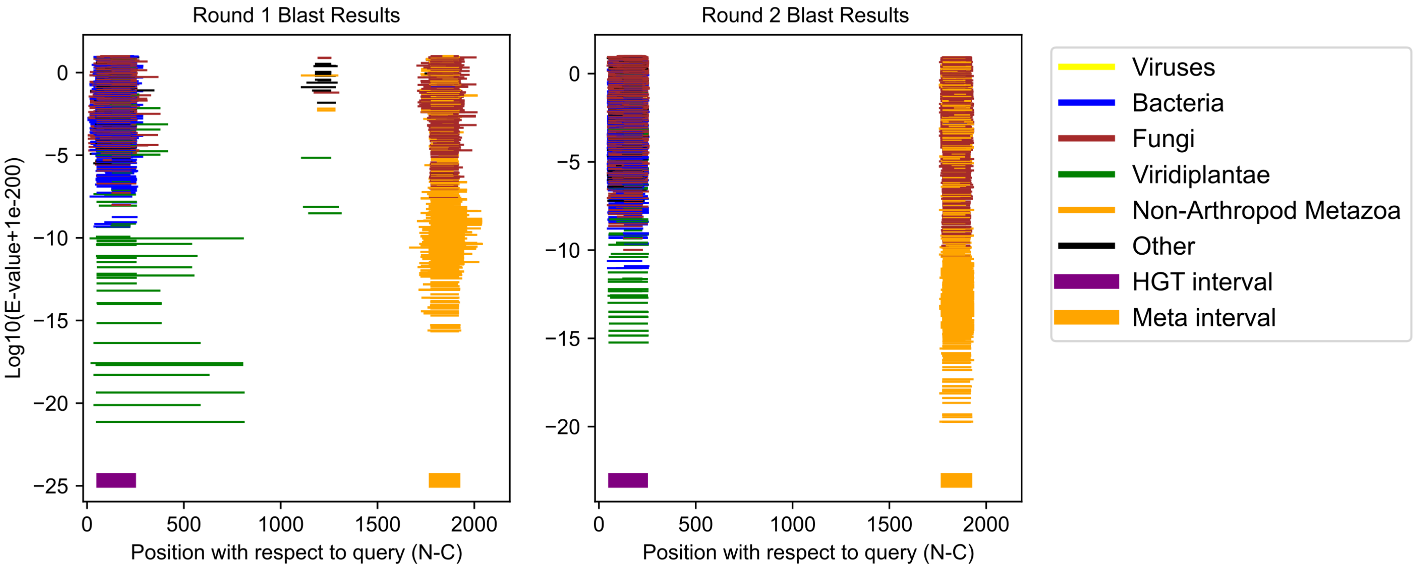

Aedes aegypti GCF_002204515.2;XP_021699539.1 (cluster 9)

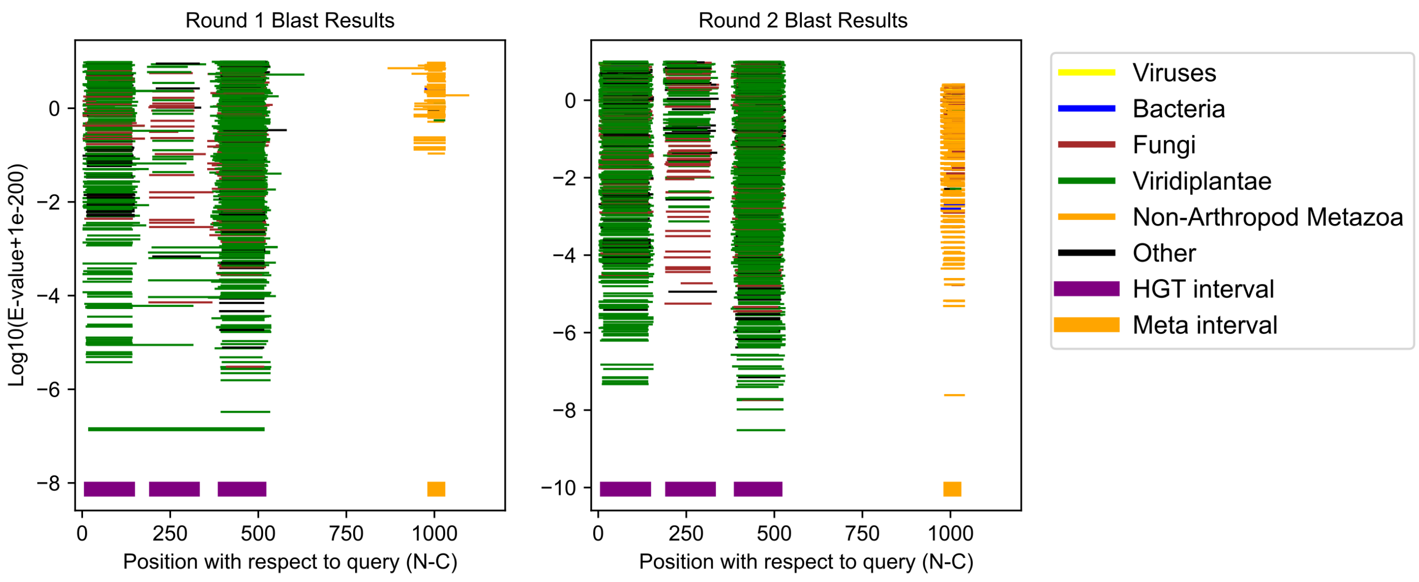

Folsomia candida GCF_002217175.1;XP_035705692.1 (cluster 10)

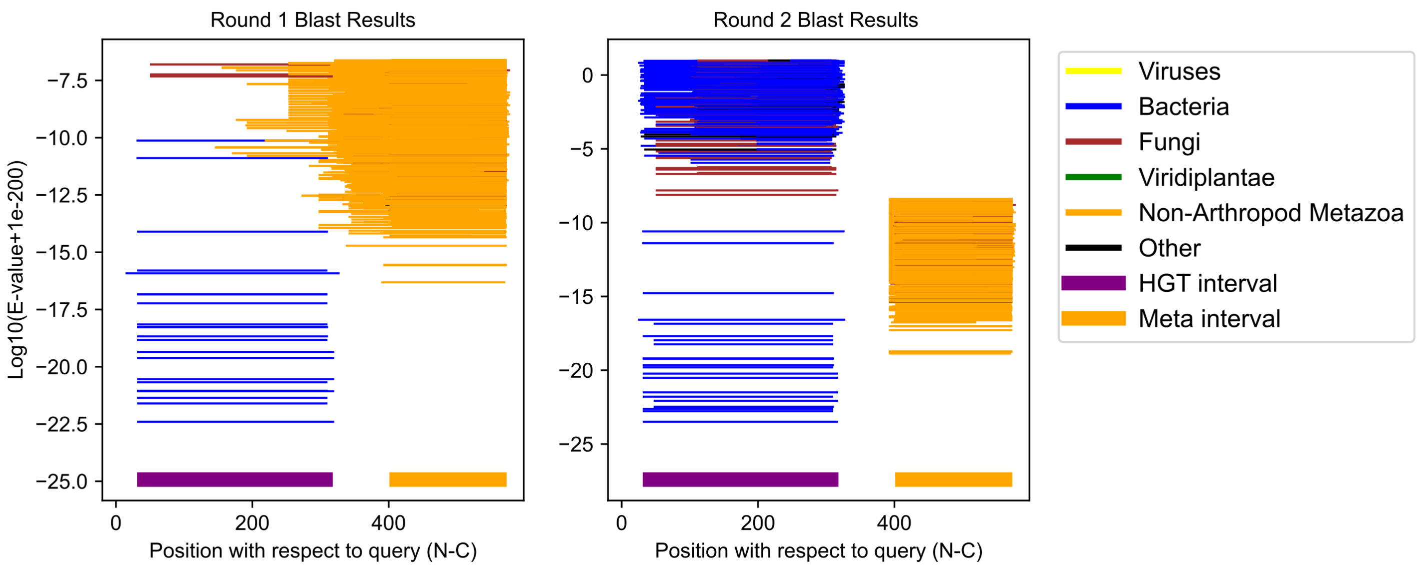

Folsomia candida GCF_002217175.1;XP_035703129.1 (cluster 11)

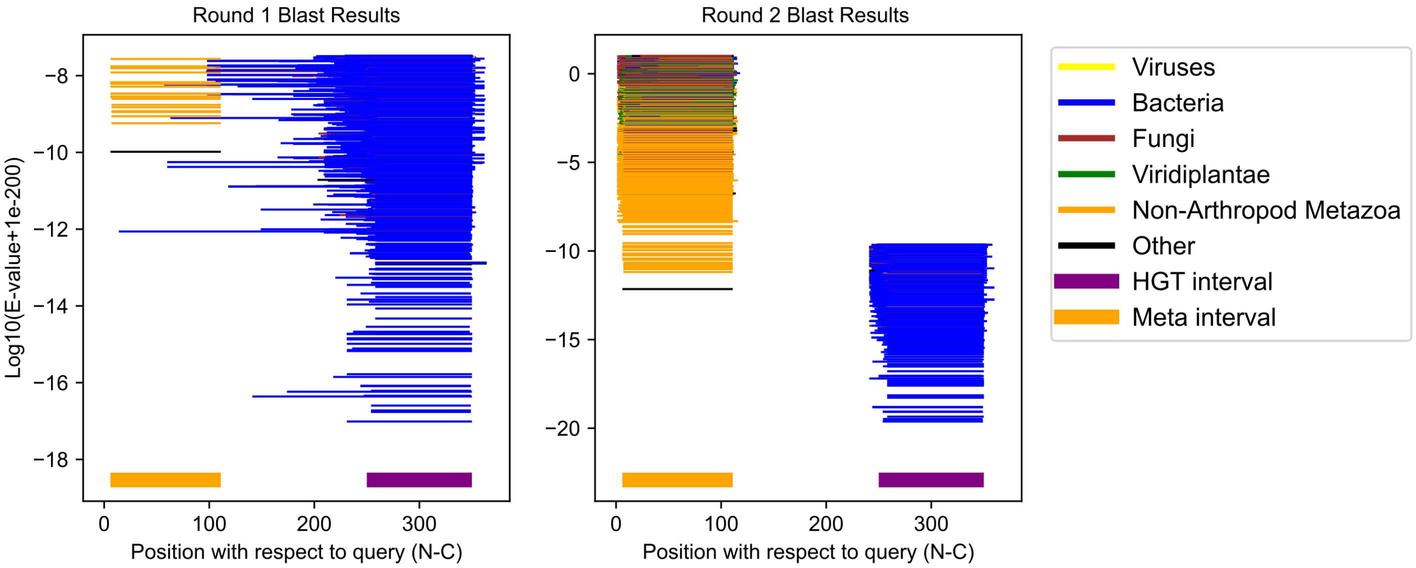

Penaeus monodon GCF_015228065.2;XP_037790819.1 (cluster 12)

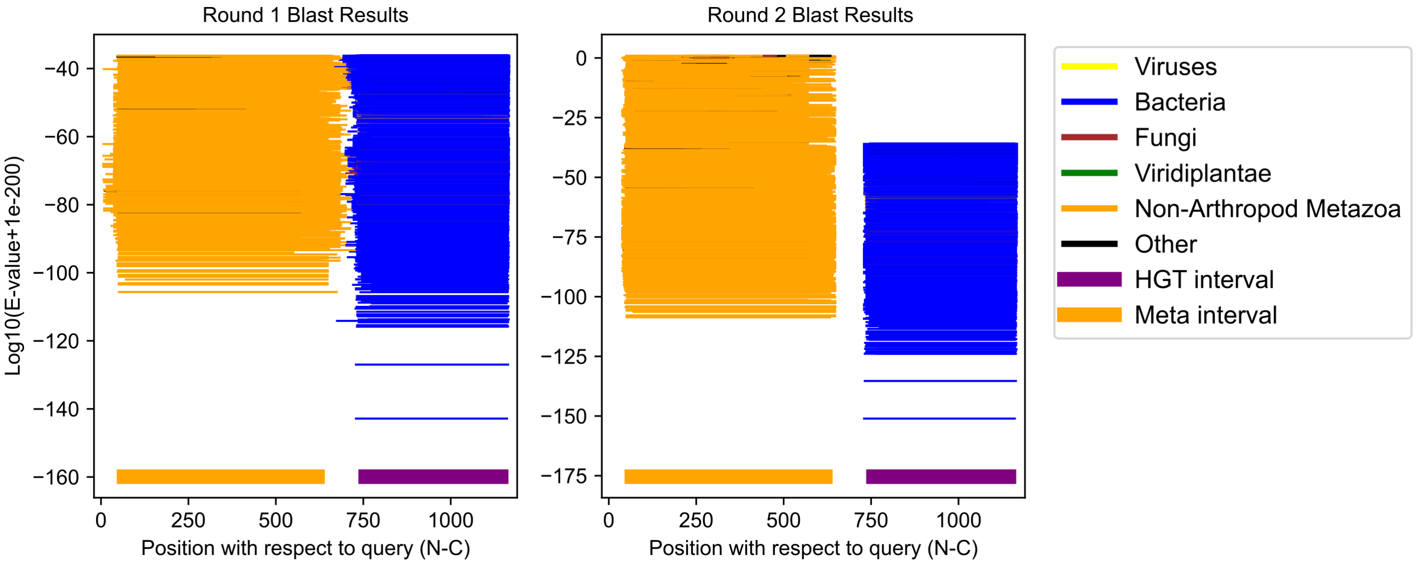

Aricia agestis GCF_905147365.1;XP_041972388.1 (cluster 13)

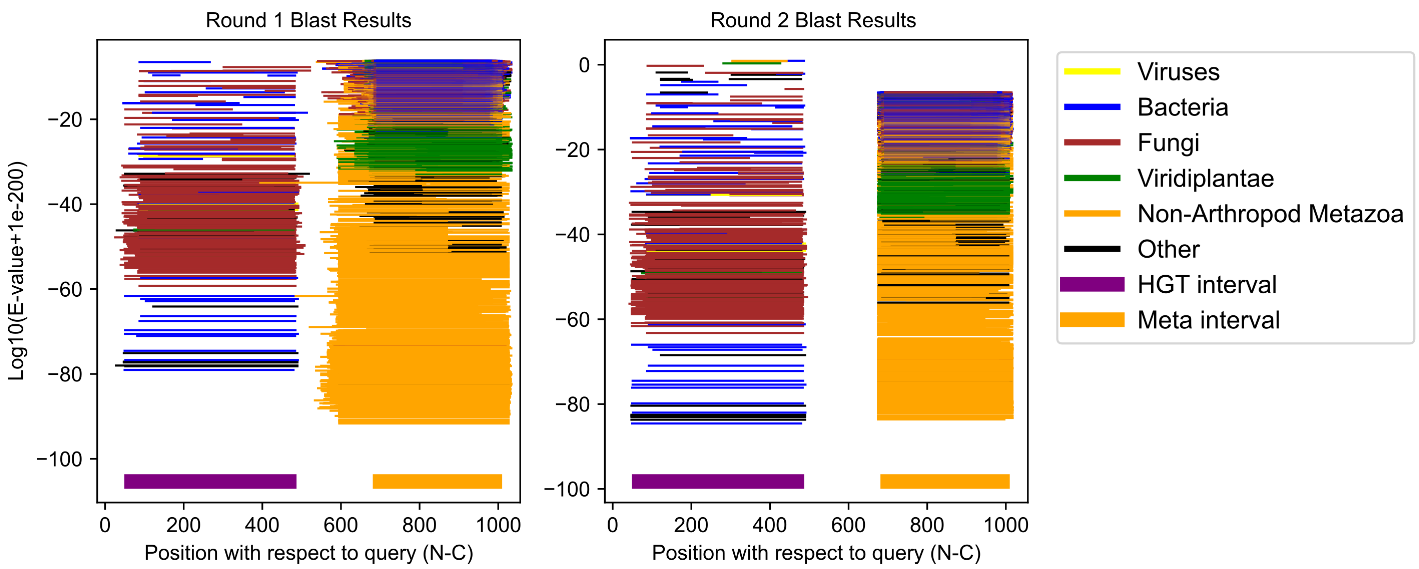

Tigriopus californicus GCF_007210705.1;XP_059092480.1 (cluster 14)

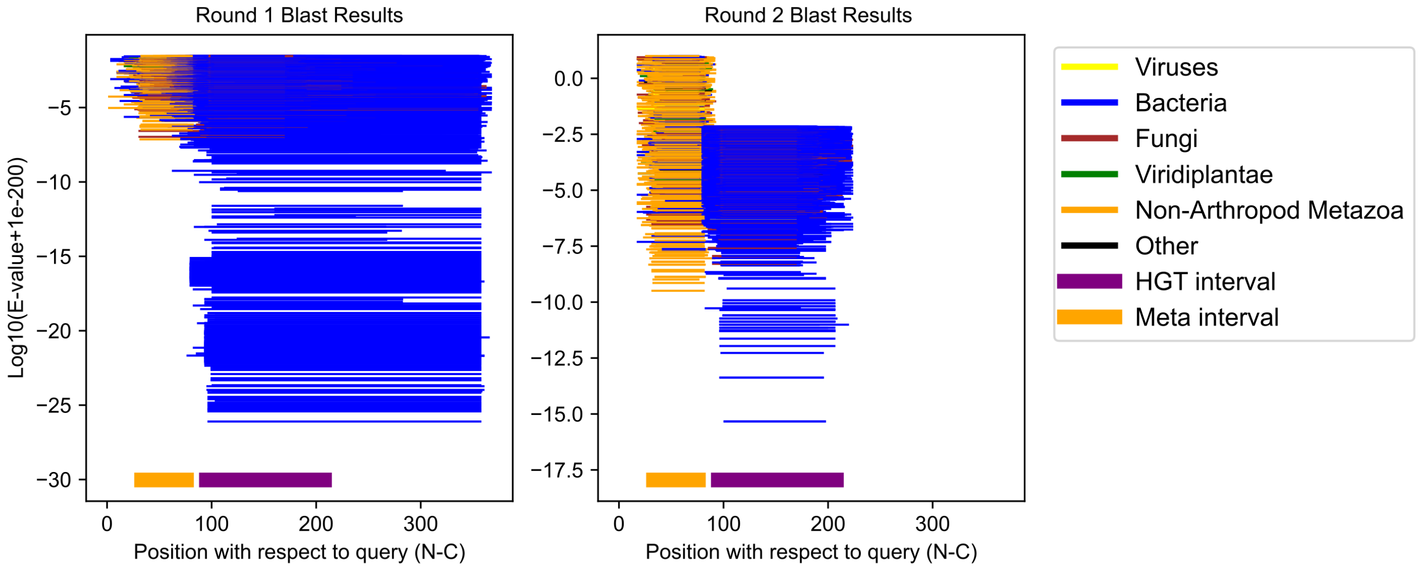

Bombyx mori GCF_030269925.1;XP_062531590.1 (cluster 15)

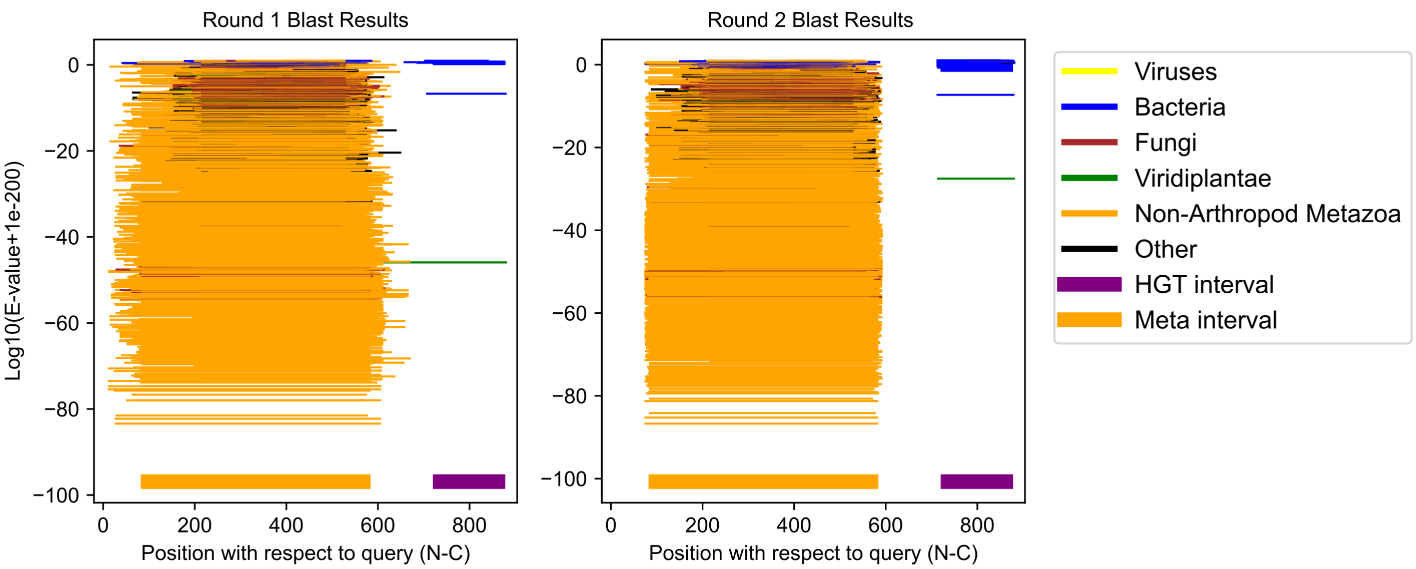

Zerene cesonia GCF_012273895.1;XP_038206242.1 (cluster 16)

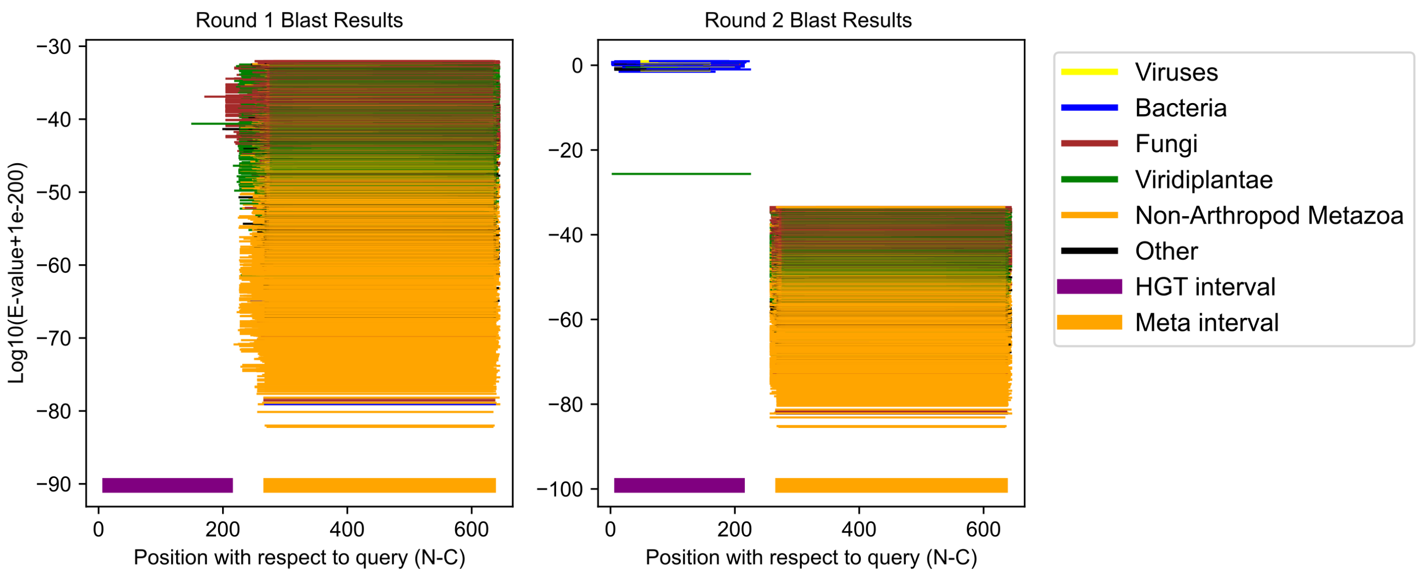

Temnothorax longispinosus GCF_030848805.1;XP_071643383.1 (cluster 17)

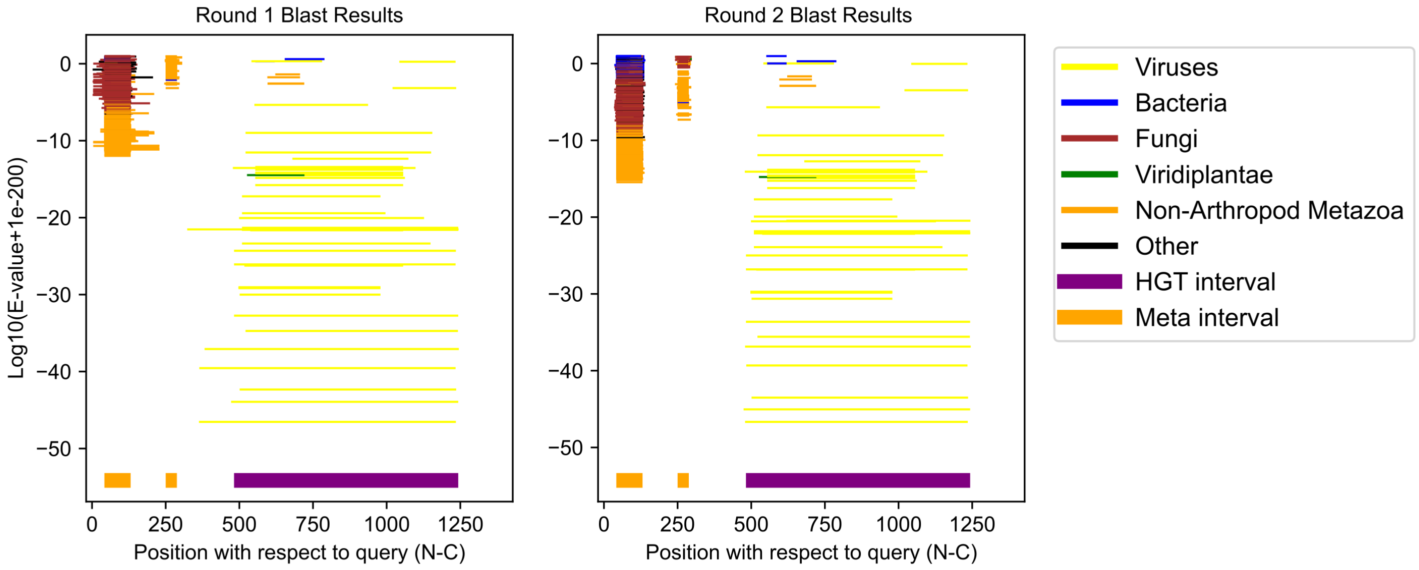

Oppia nitens GCF_028296485.1;XP_054167651.1 (cluster 18)

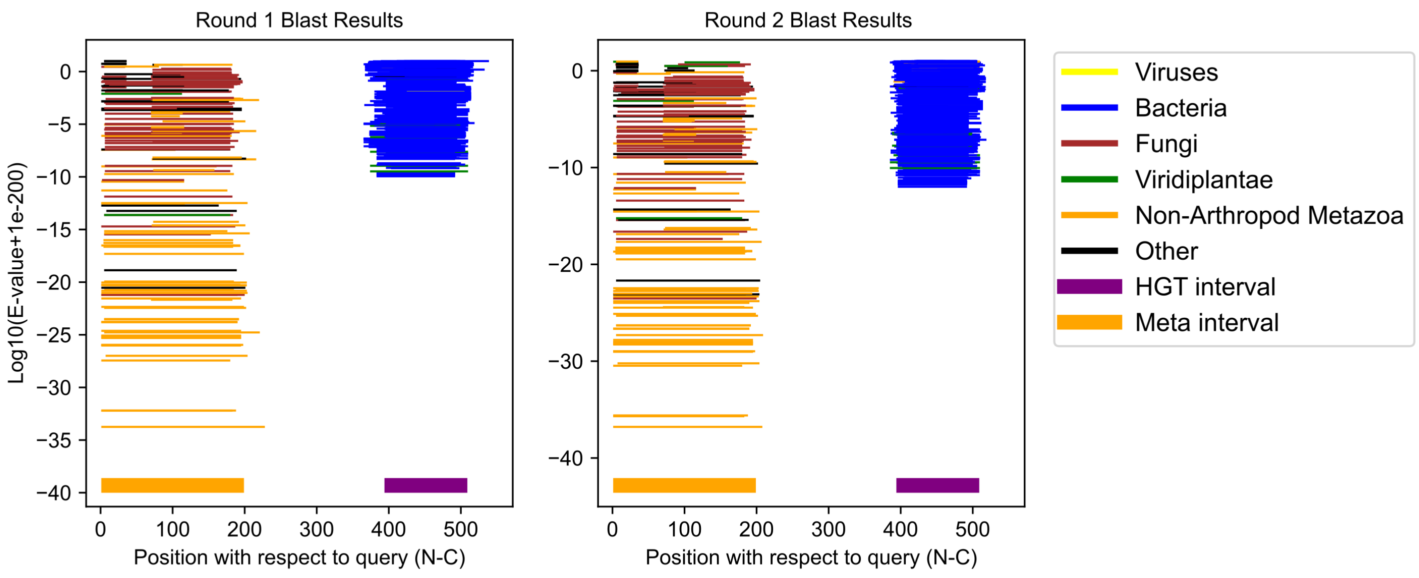

Daphnia magna GCF_020631705.1;XP_032797465.2 (cluster 19)

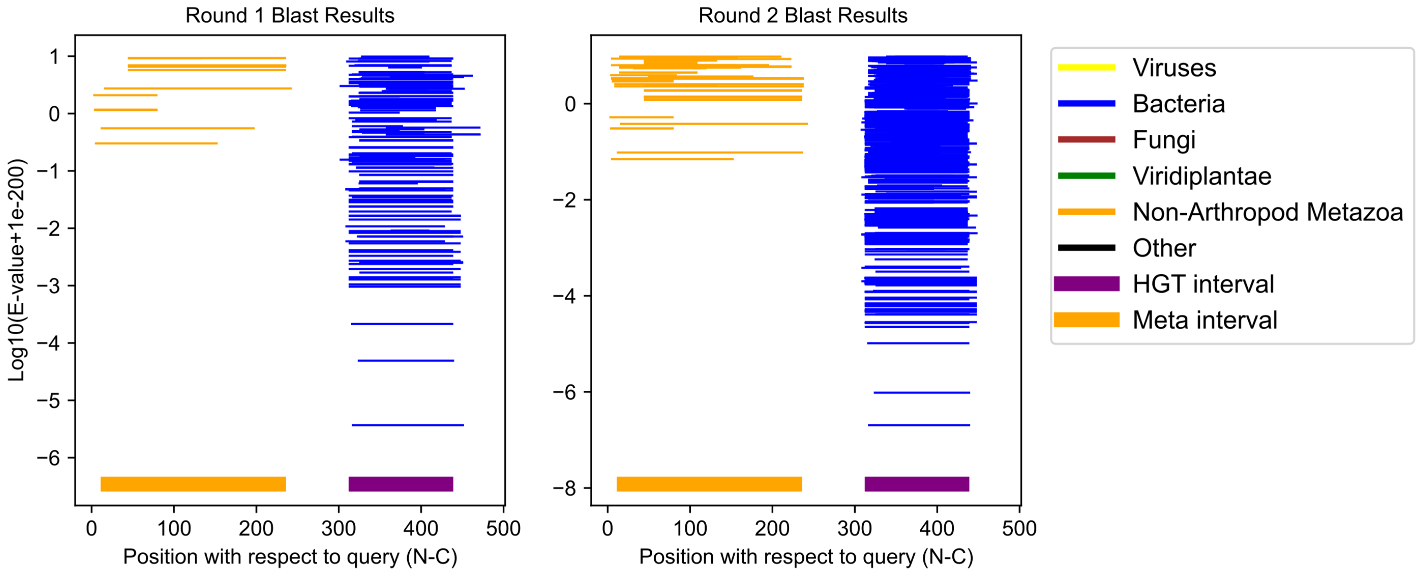

Daphnia pulex GCF_021134715.1;XP_046453153.1 (cluster 20)

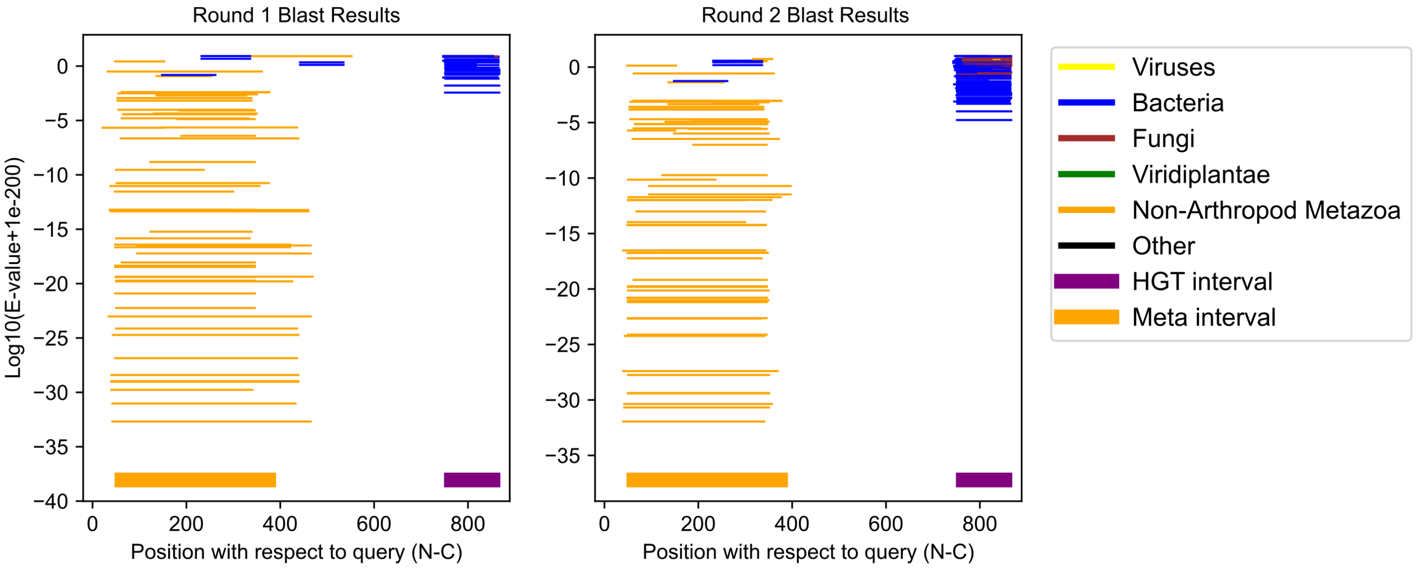

Daphnia pulex GCF_021134715.1;XP_046460947.1 (cluster 21)

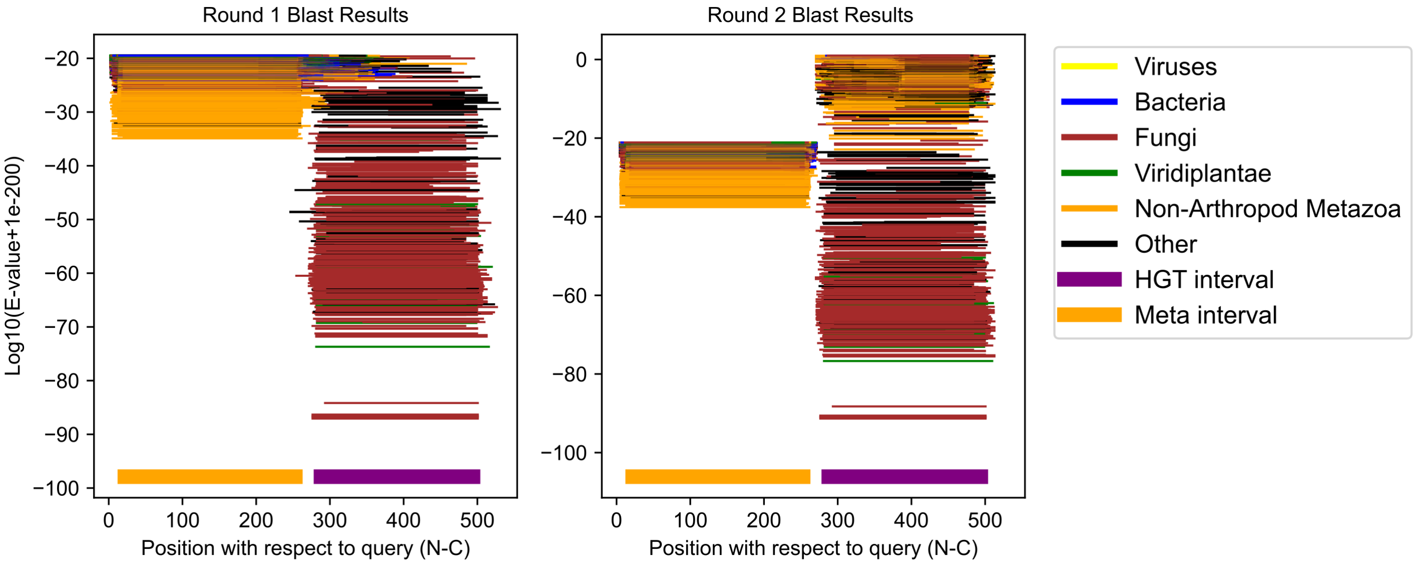

Ostrinia furnacalis GCF_004193835.3;XP_028178173.1 (cluster 22)

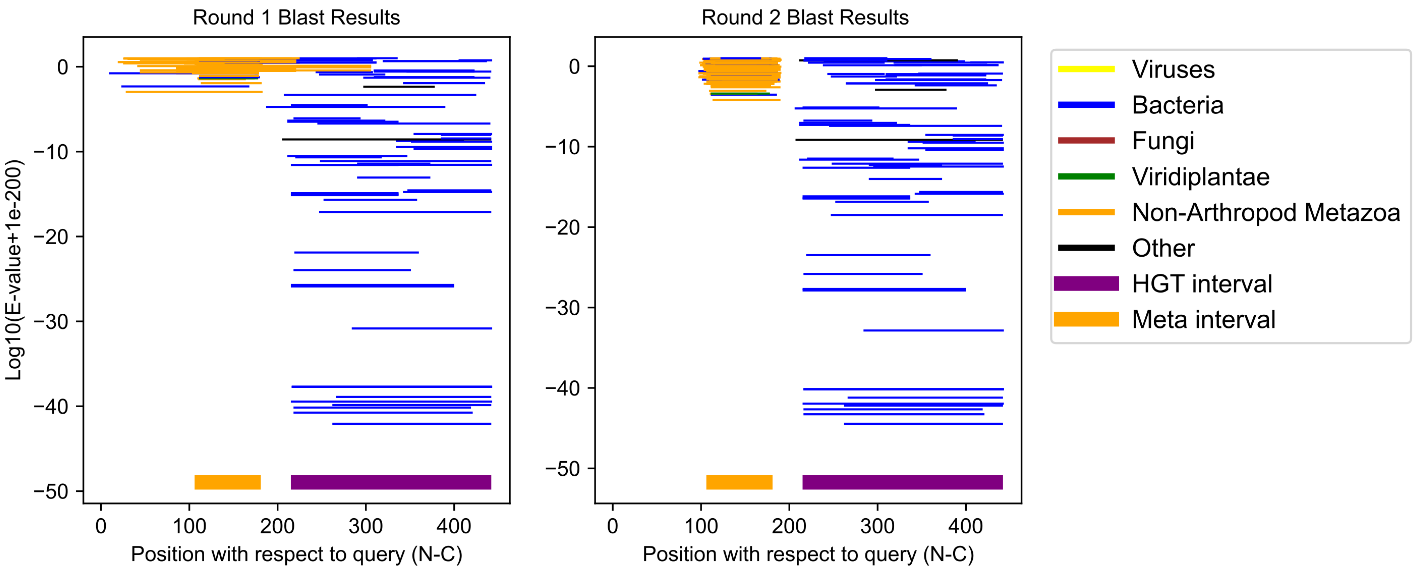

Tigriopus californicus GCF_007210705.1;XP_059086440.1 (cluster 23)

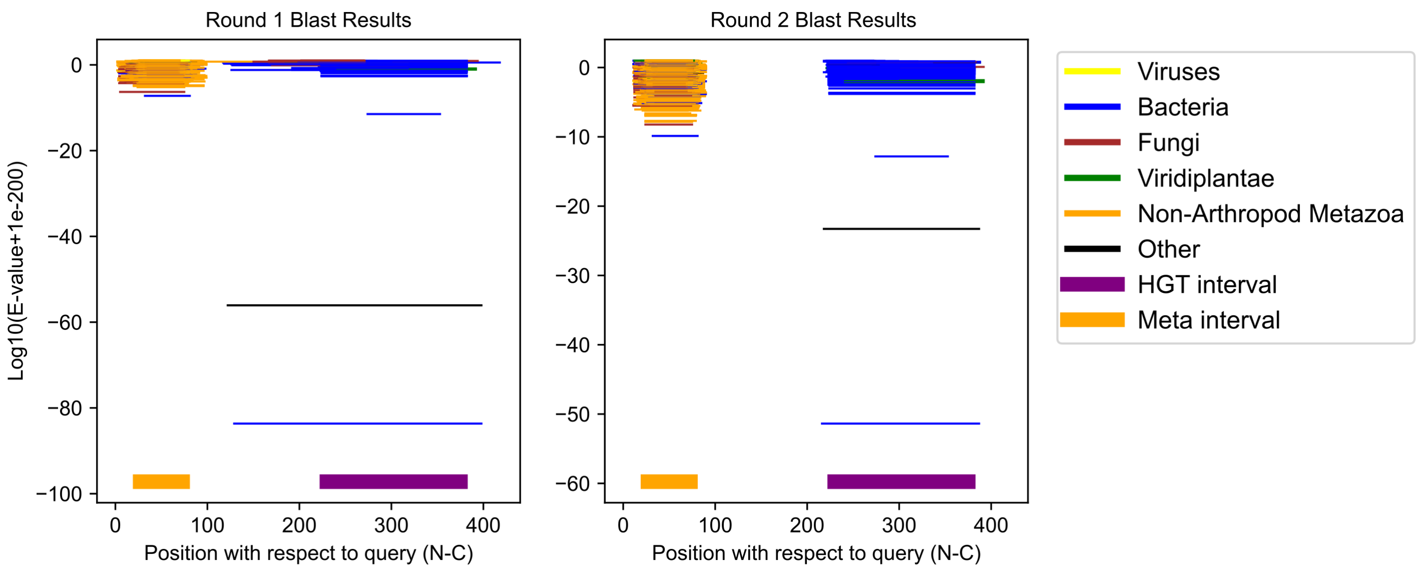

Eurosta solidaginis GCF_040869045.1;XP_067614489.1 (cluster 24)

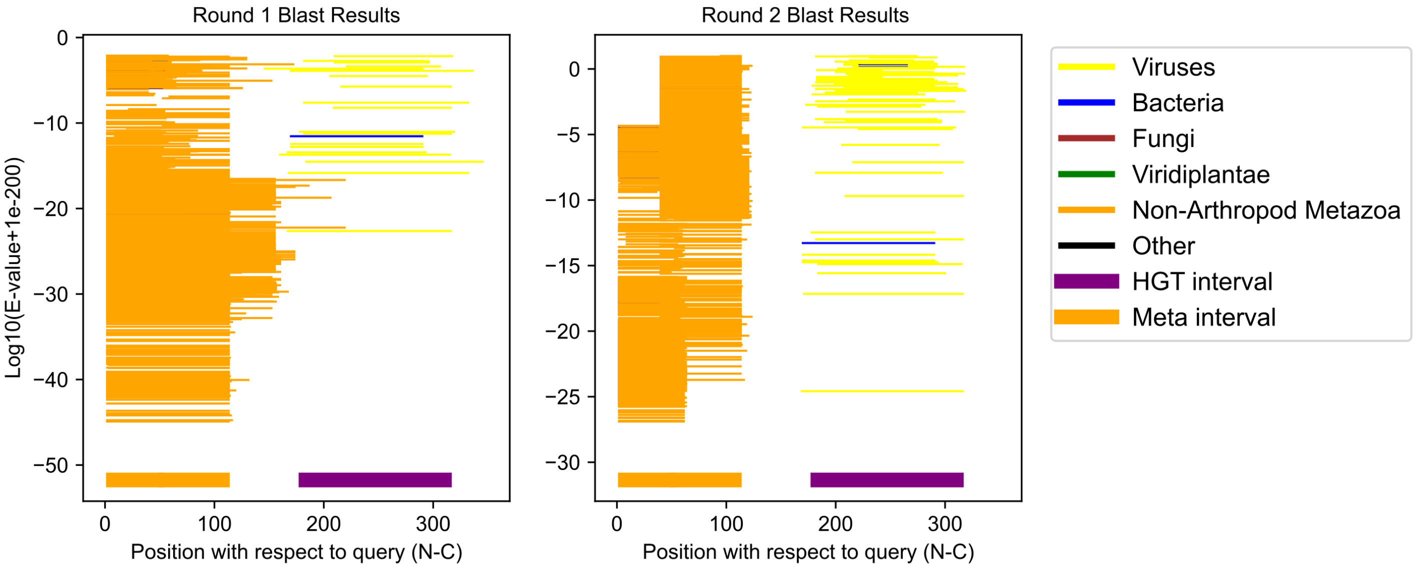

Dermacentor albipictus GCF_038994185.2;XP_070390518.1 (cluster 25)

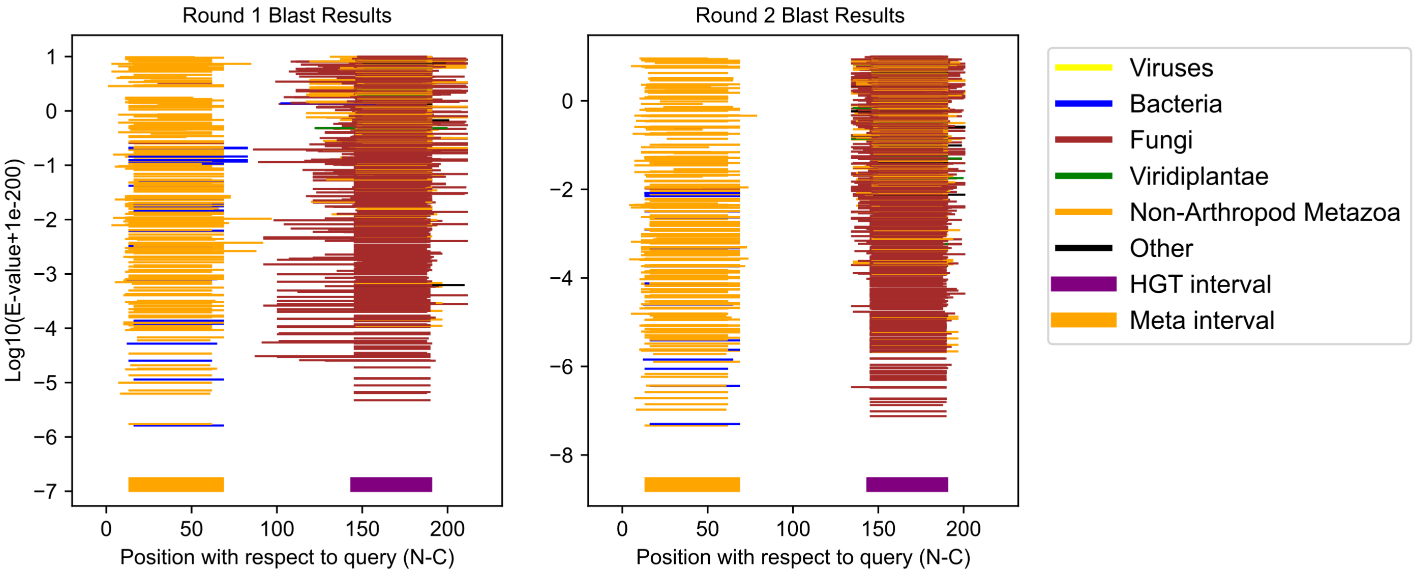

Oppia nitens GCF_028296485.1;XP_054166030.1 (cluster 26)

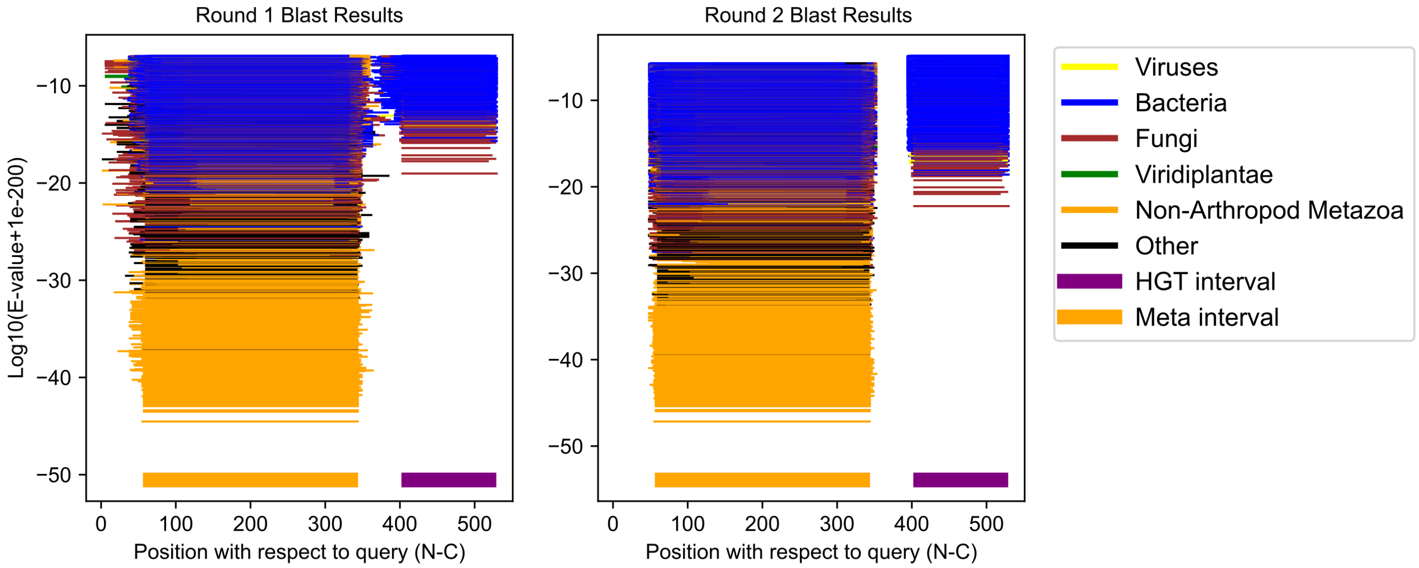

Oppia nitens GCF_028296485.1;XP_054165804.1 (cluster 27)

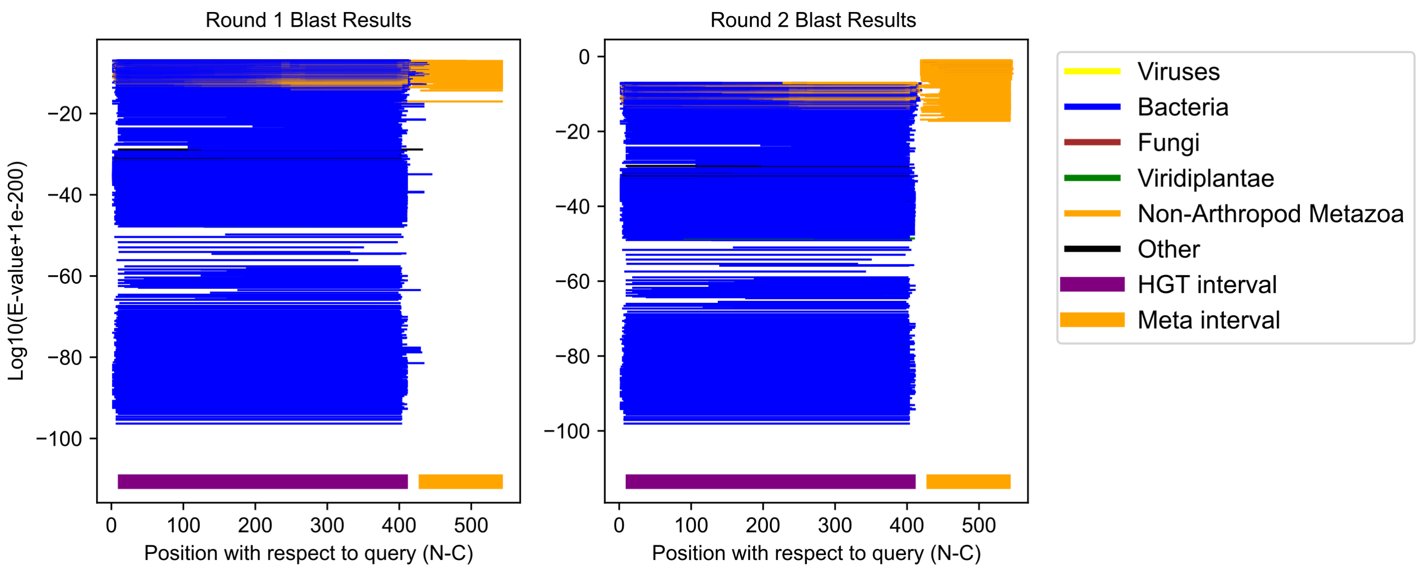

Oppia nitens GCF_028296485.1;XP_054165701.1 (cluster 28)

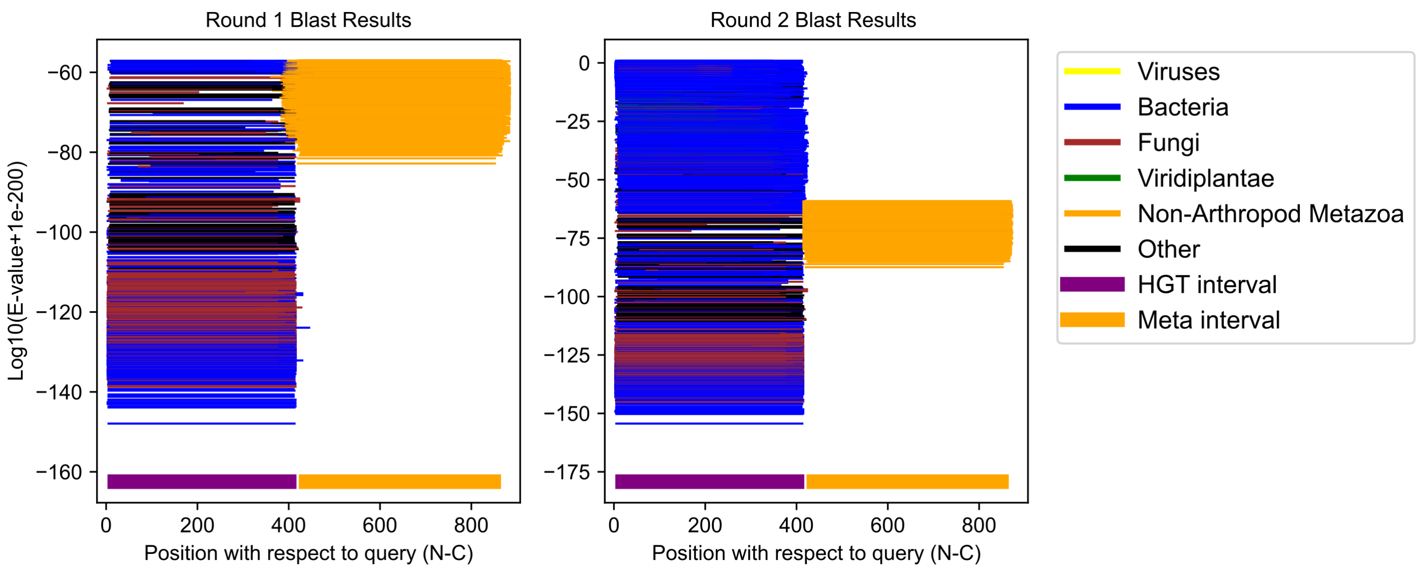

Plutella xylostella GCF_932276165.1;XP_048481436.1 (cluster 29)

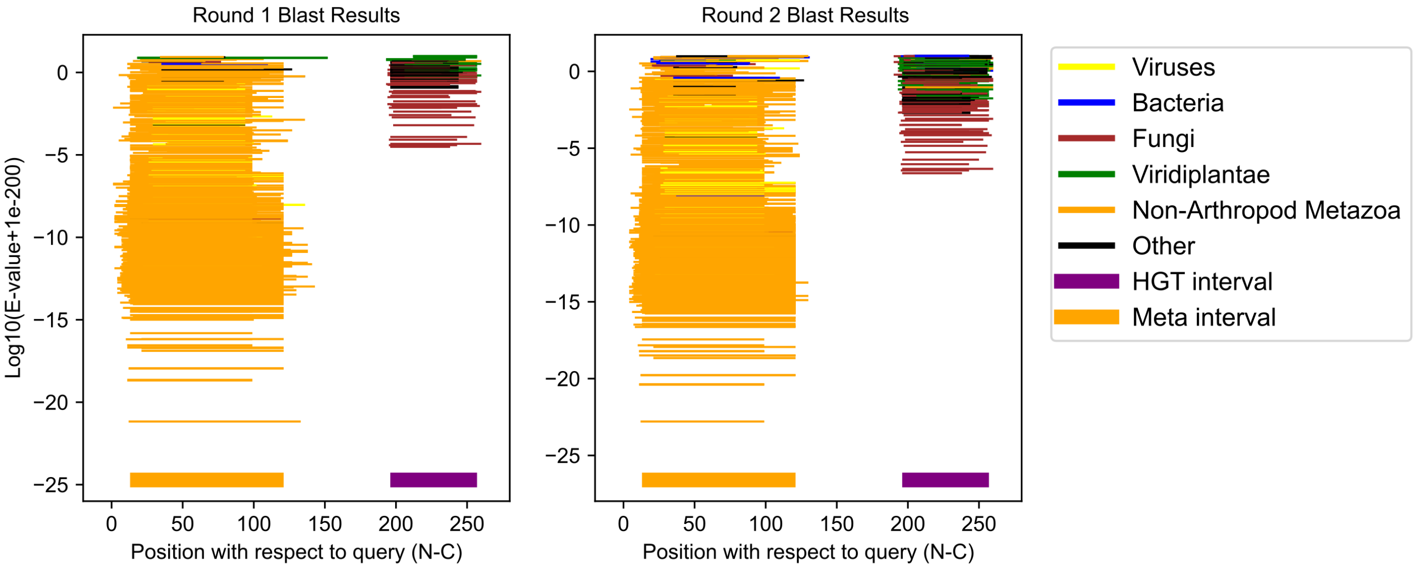

Planococcus citri GCF_950023065.1;XP_065207242.1 (cluster 30)

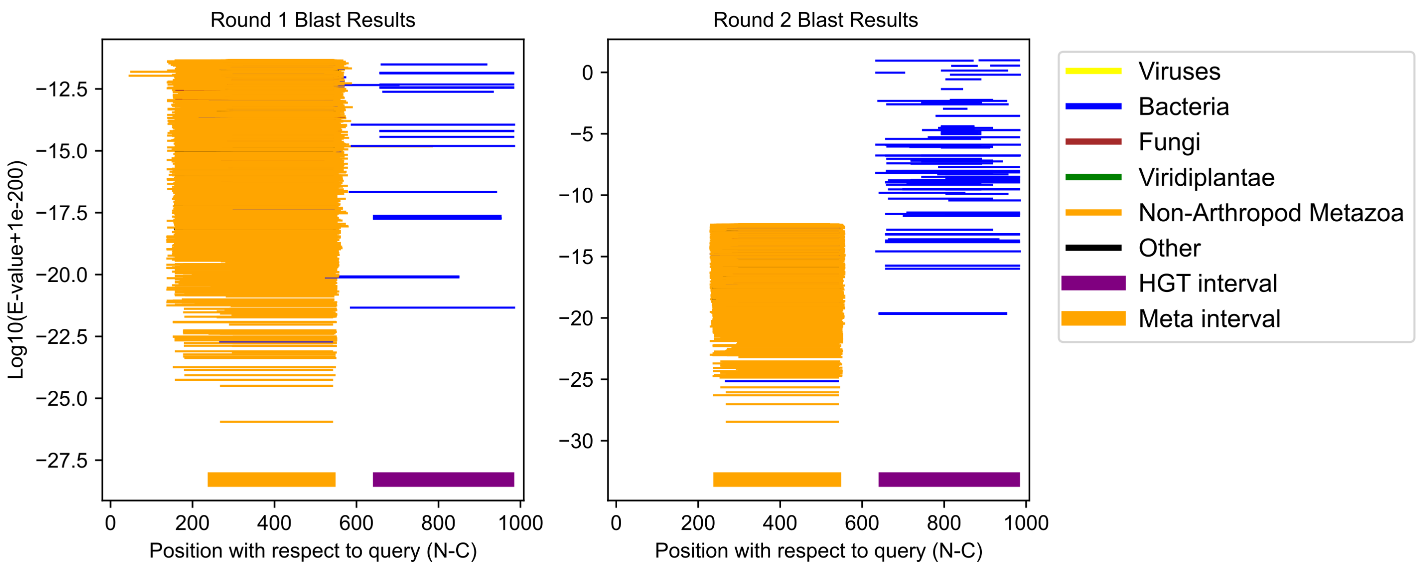

Oppia nitens GCF_028296485.1;XP_054156590.1 (cluster 31)

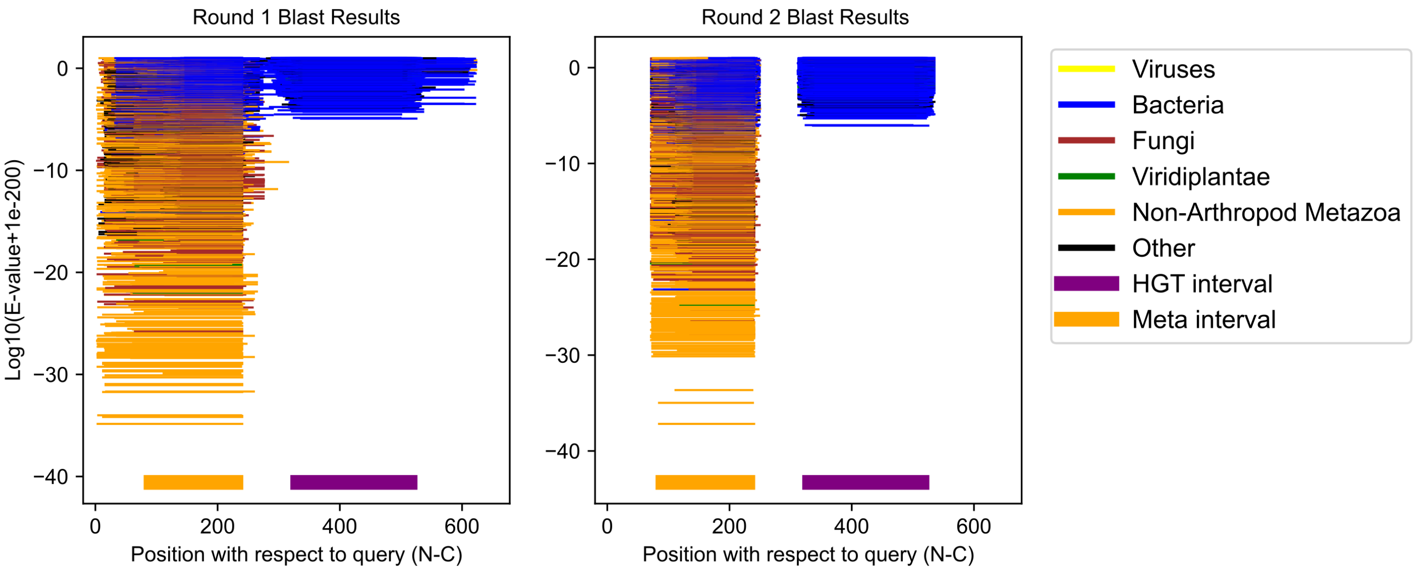

Dalotia coriaria GCF_025399875.1;XP_065173386.1 (cluster 32)

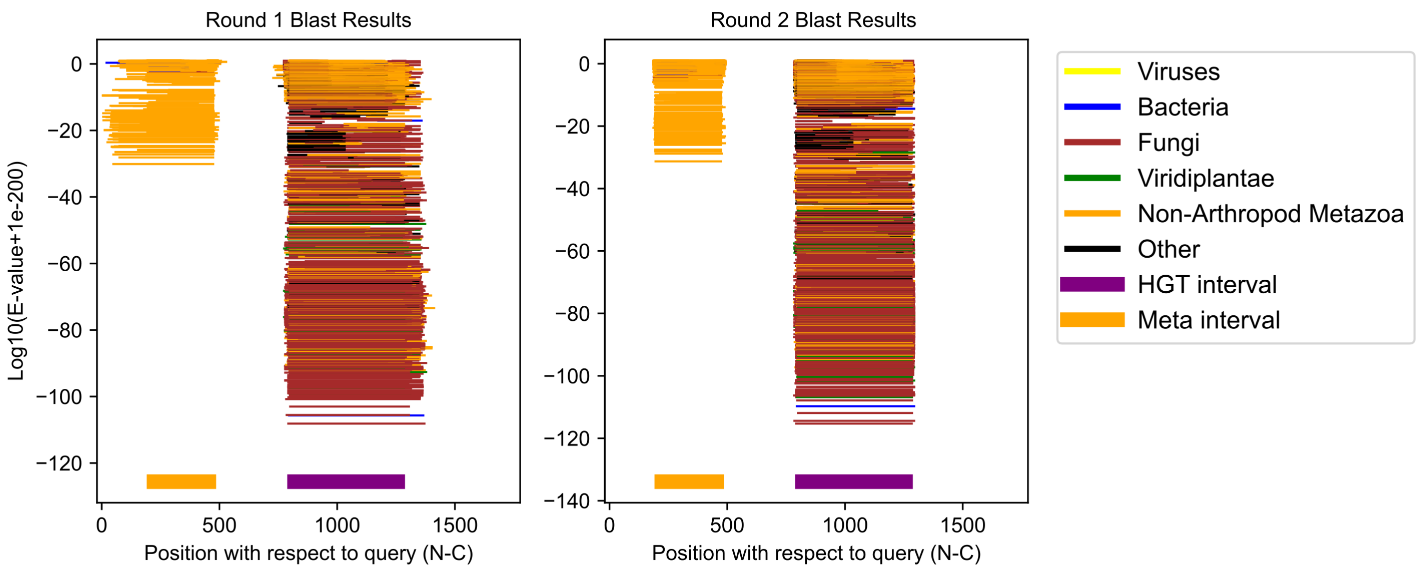

Daktulosphaira vitifoliae GCF_025091365.1;XP_050527957.1 (cluster 33)

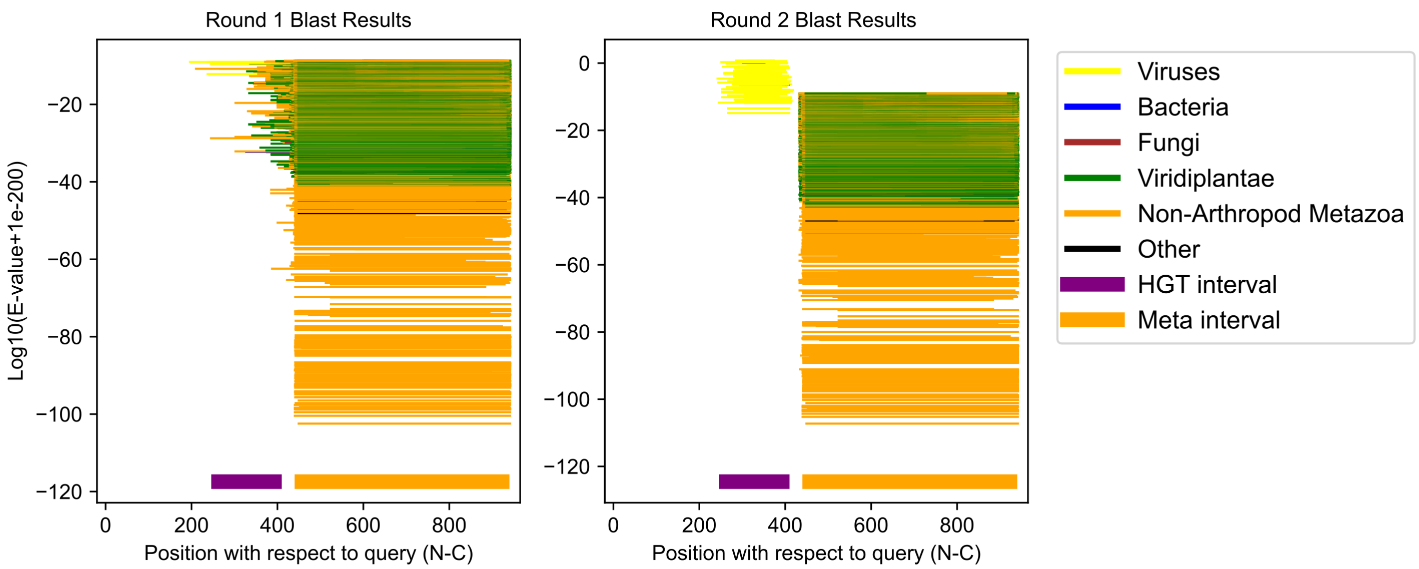

Planococcus citri GCF_950023065.1;XP_065218164.1 (cluster 34)

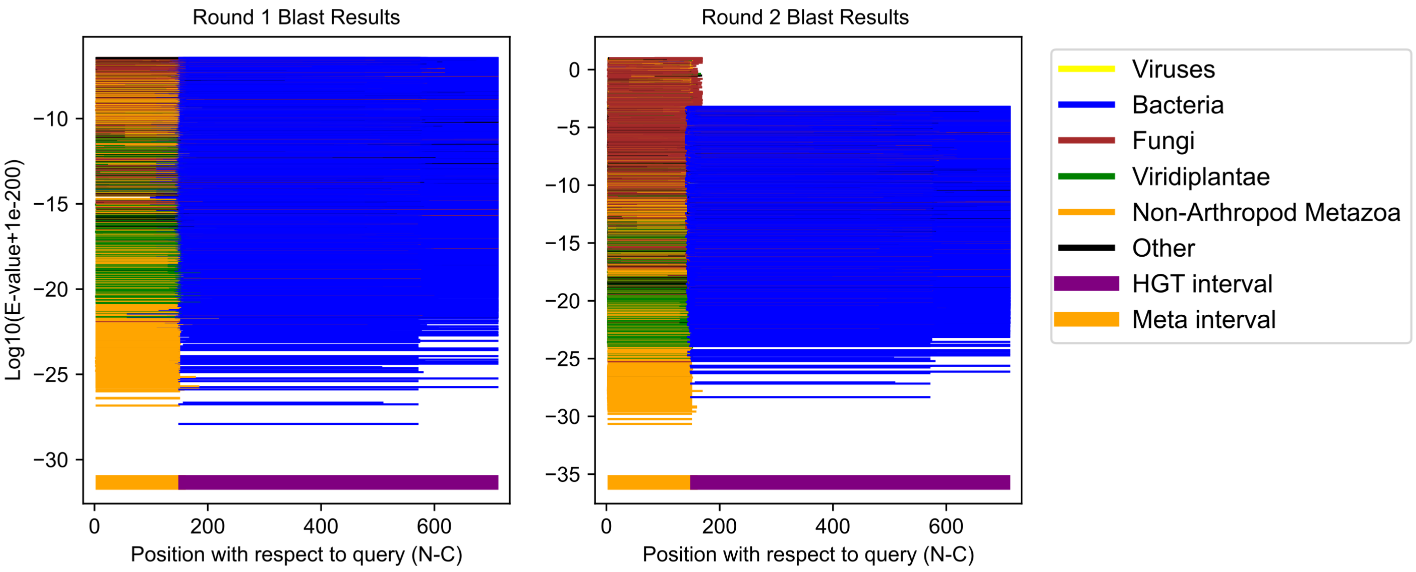

Phlebotomus papatasi GCF_024763615.1;XP_055704333.1 (cluster 35)

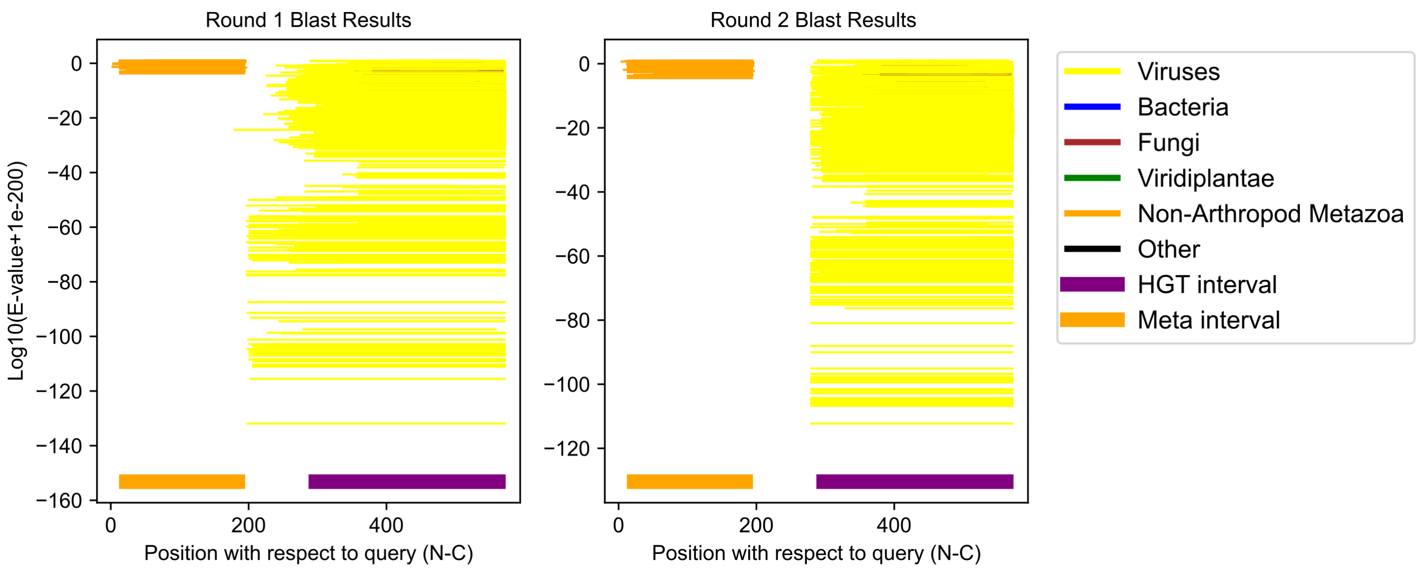

Schistocerca gregaria GCF_023897955.1;XP_049848309.1 (cluster 36)

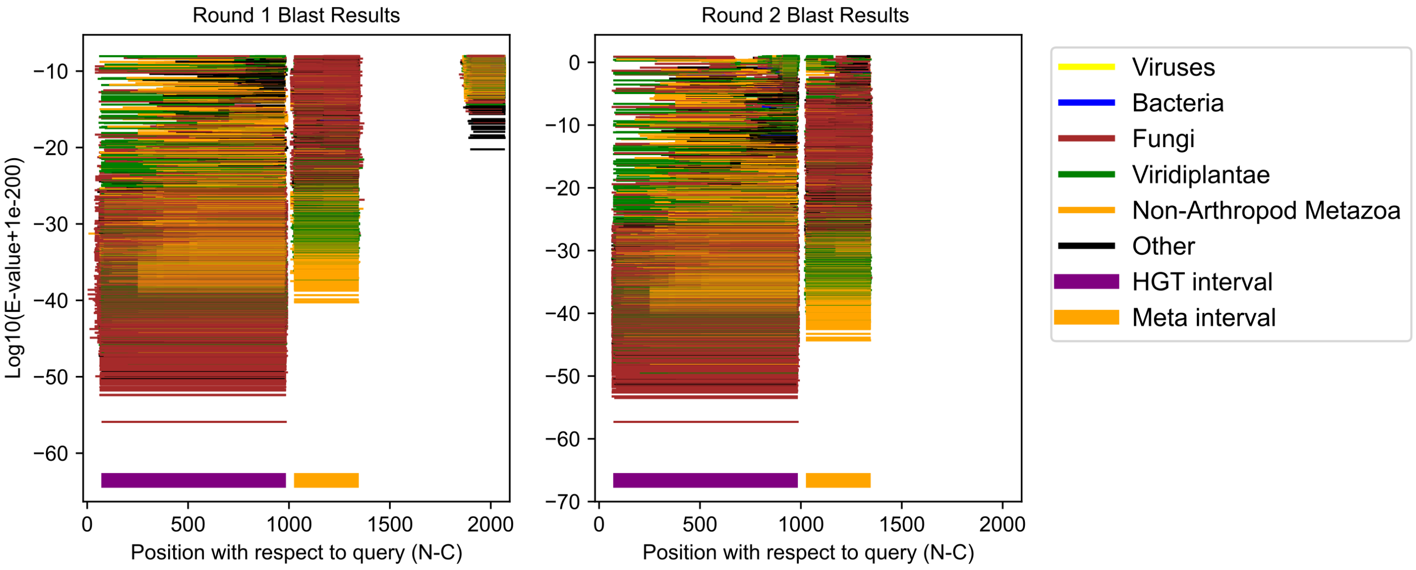

Planococcus citri GCF_950023065.1;XP_065223590.1 (cluster 37)

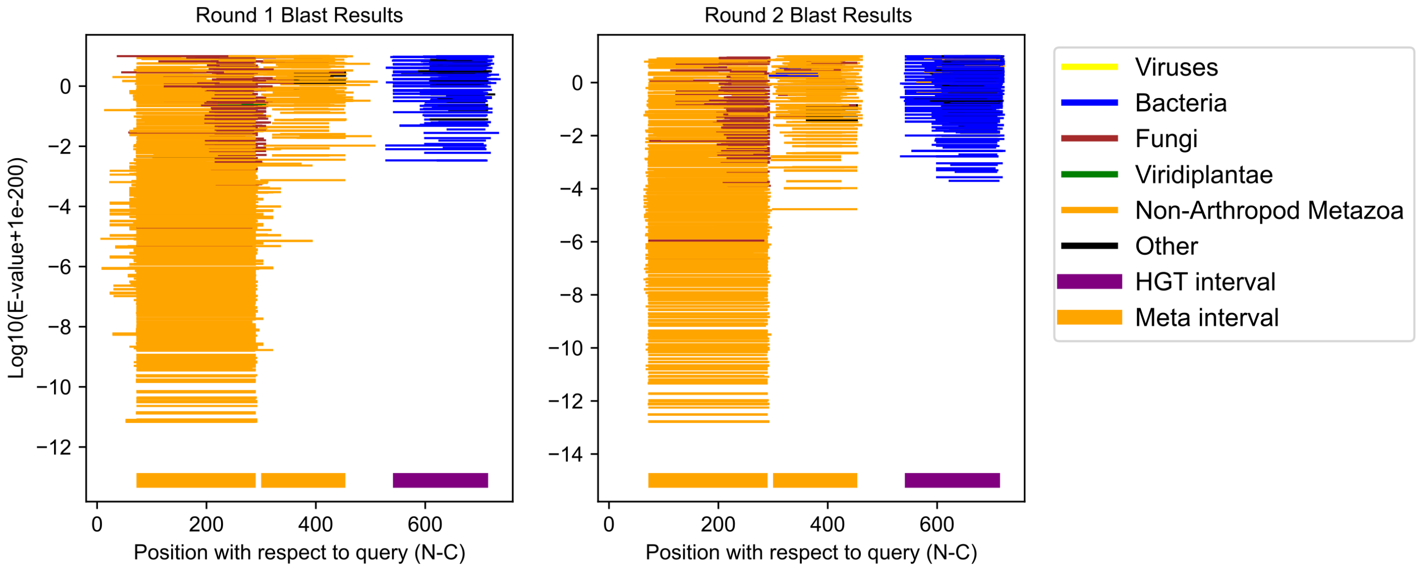

Planococcus citri GCF_950023065.1;XP_065225331.1 (cluster 38)

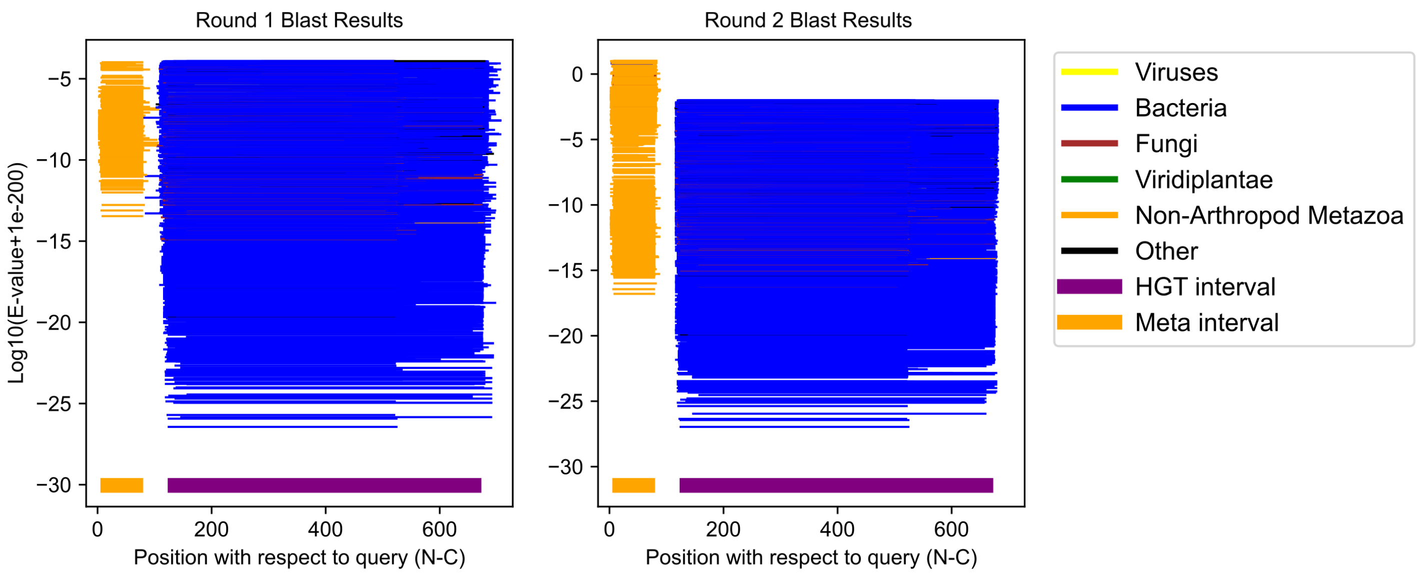

Eurytemora carolleeae GCF_000591075.1;XP_023331203.1 (cluster 39)

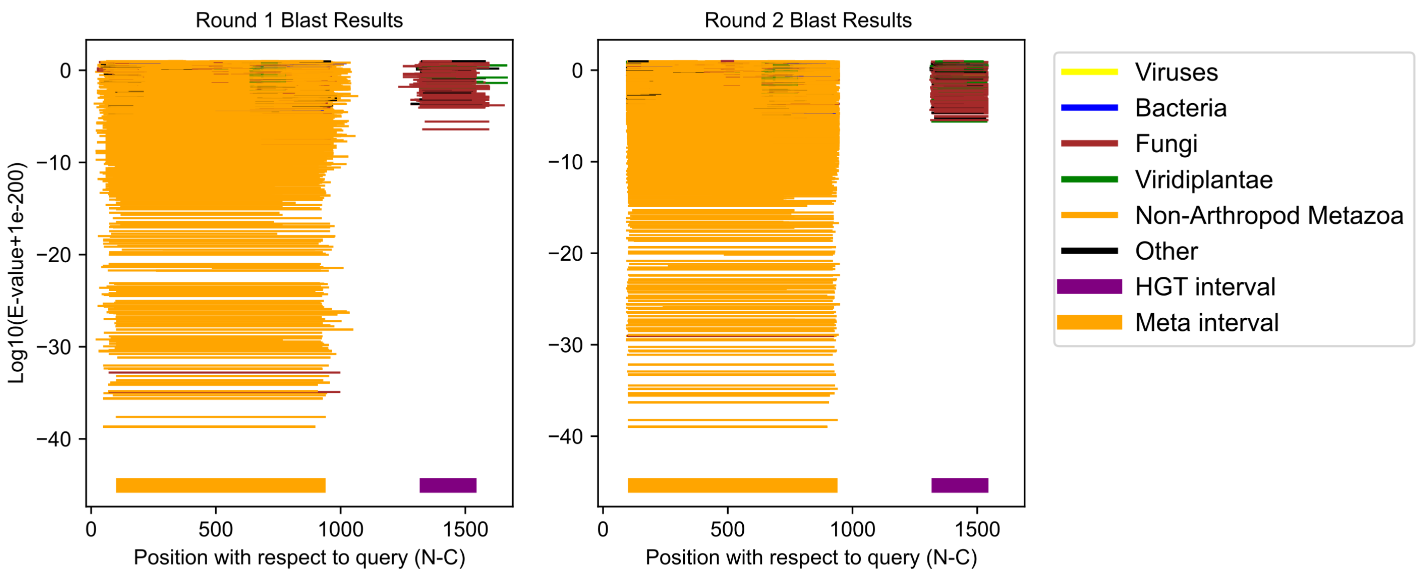

Oppia nitens GCF_028296485.1;XP_054157154.1 (cluster 40)

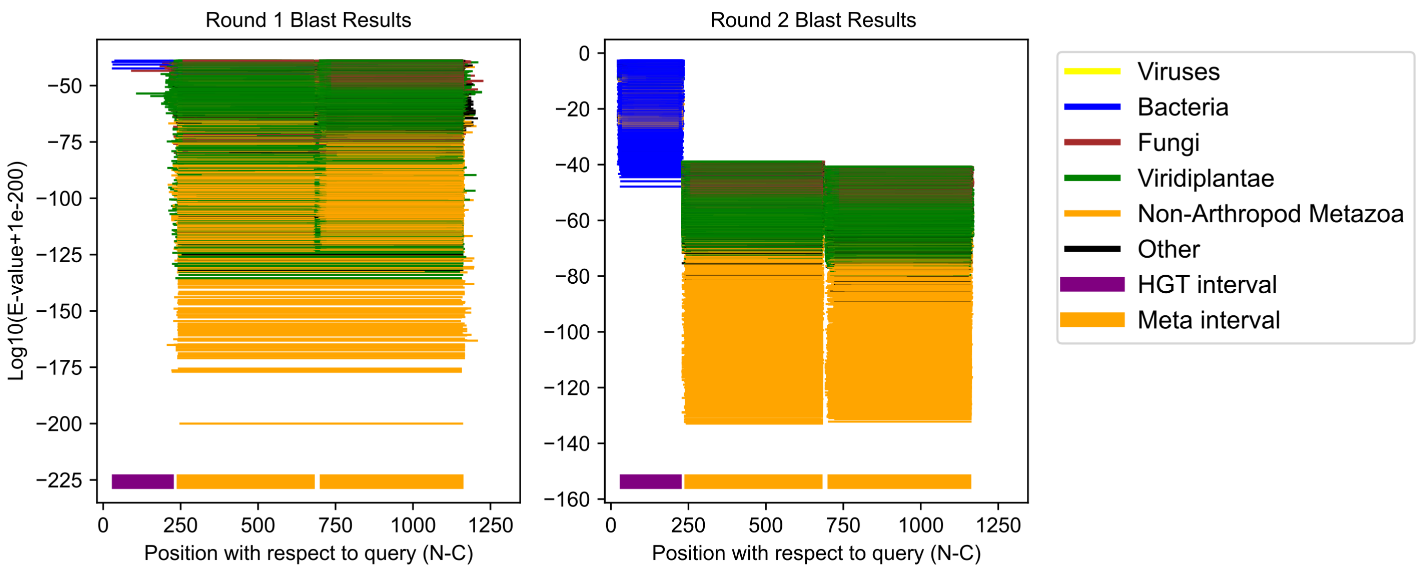

Ischnura elegans GCF_921293095.1;XP_046403459.1 (cluster 41)

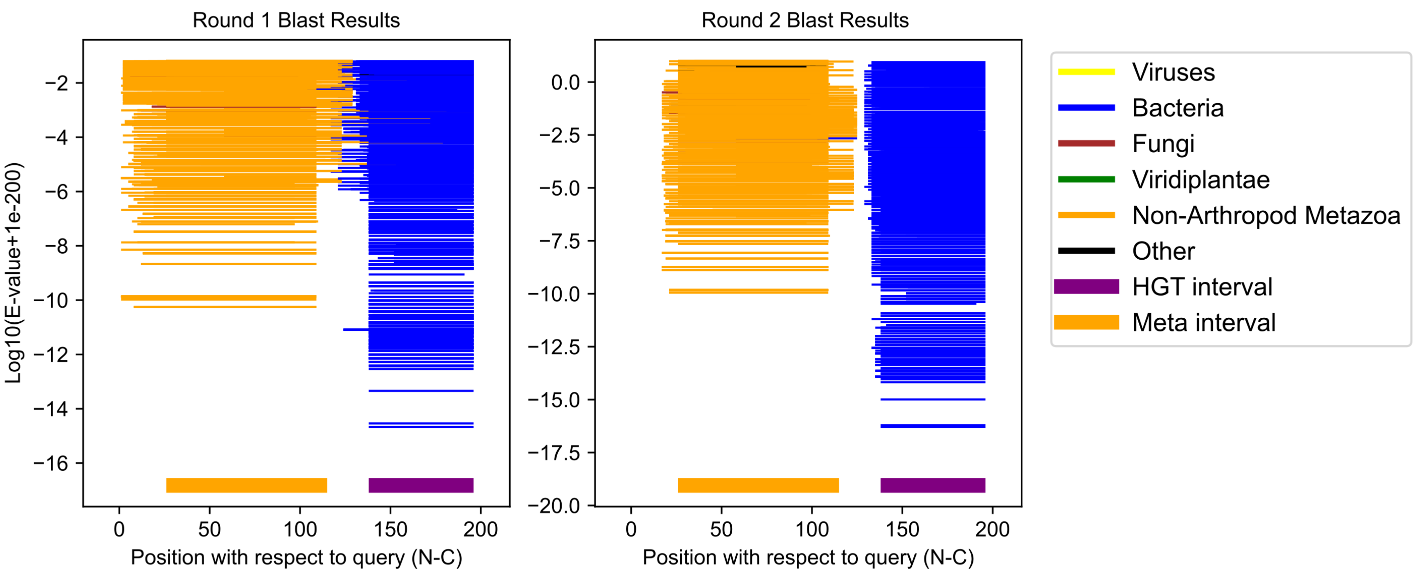

Oppia nitens GCF_028296485.1;XP_054169292.1 (cluster 42)

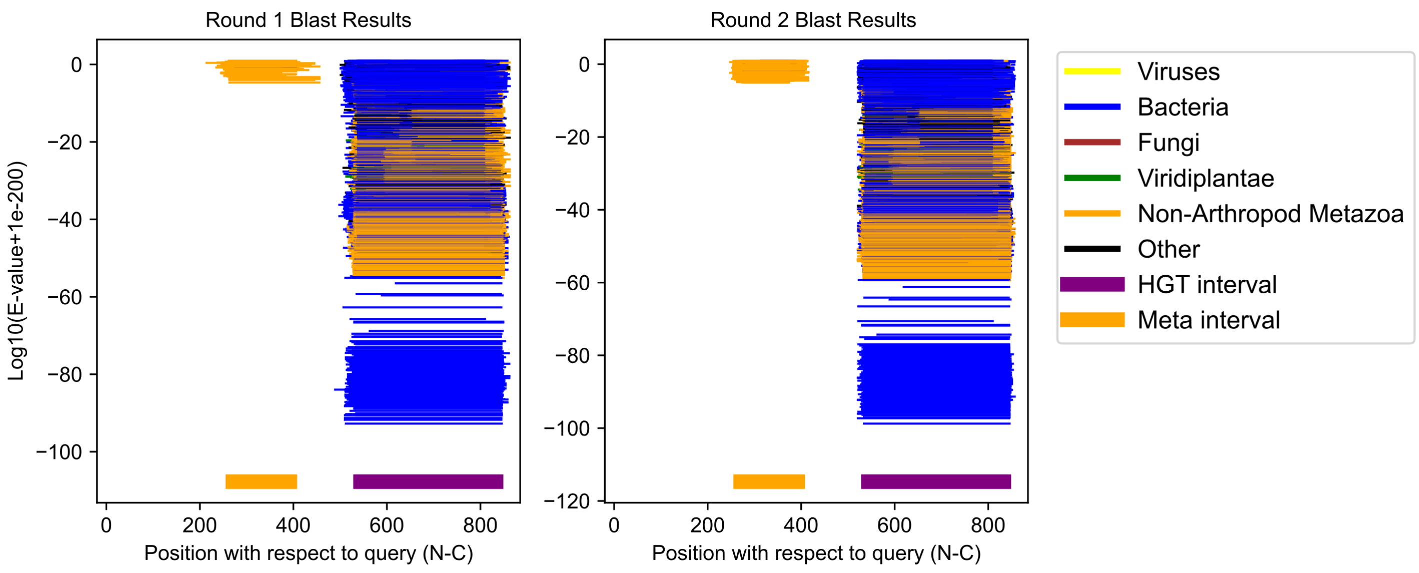

### Macrosteles quadrilineatus GCF_028750875.1;XP_054272665.1 (cluster 43)

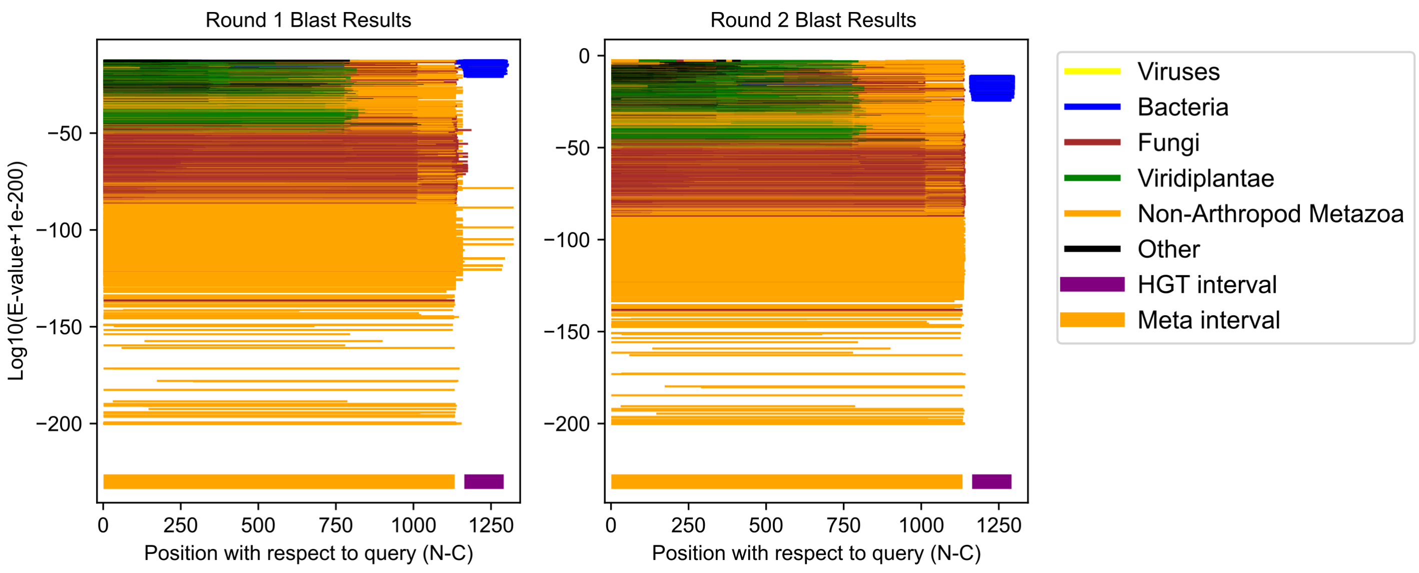

Aedes albopictus GCF_035046485.1;XP_062702136.1 (cluster 44)

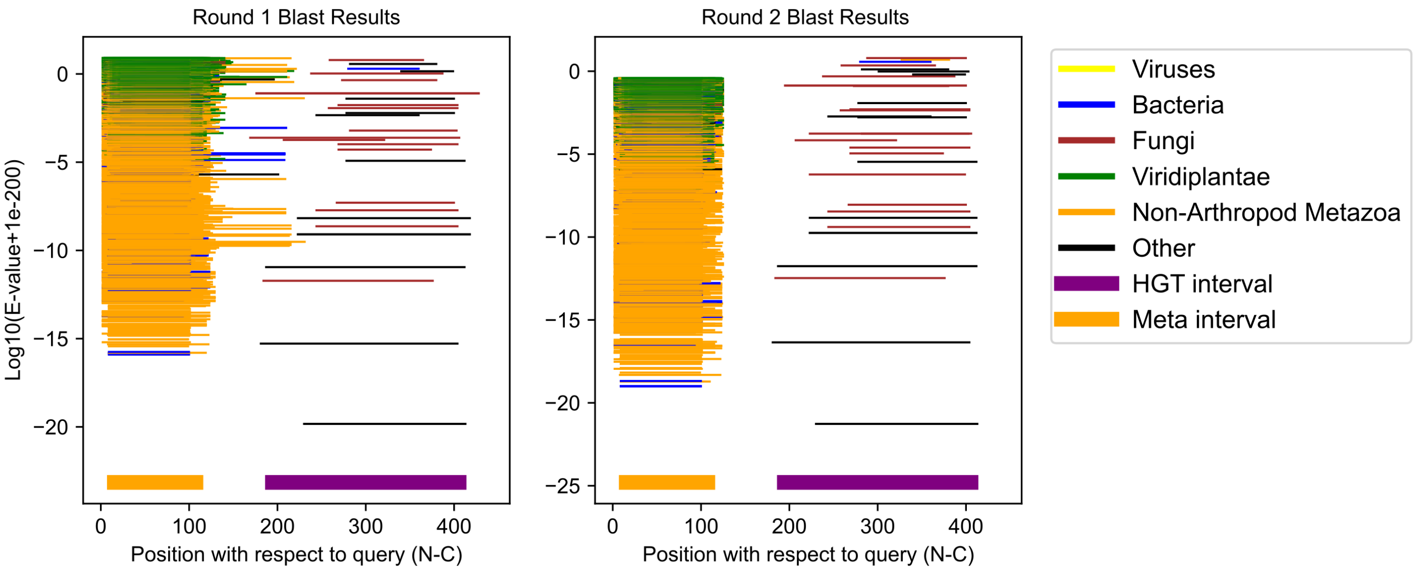

Ornithodoros turicata GCF_037126465.1;XP_064461306.1 (cluster 45)

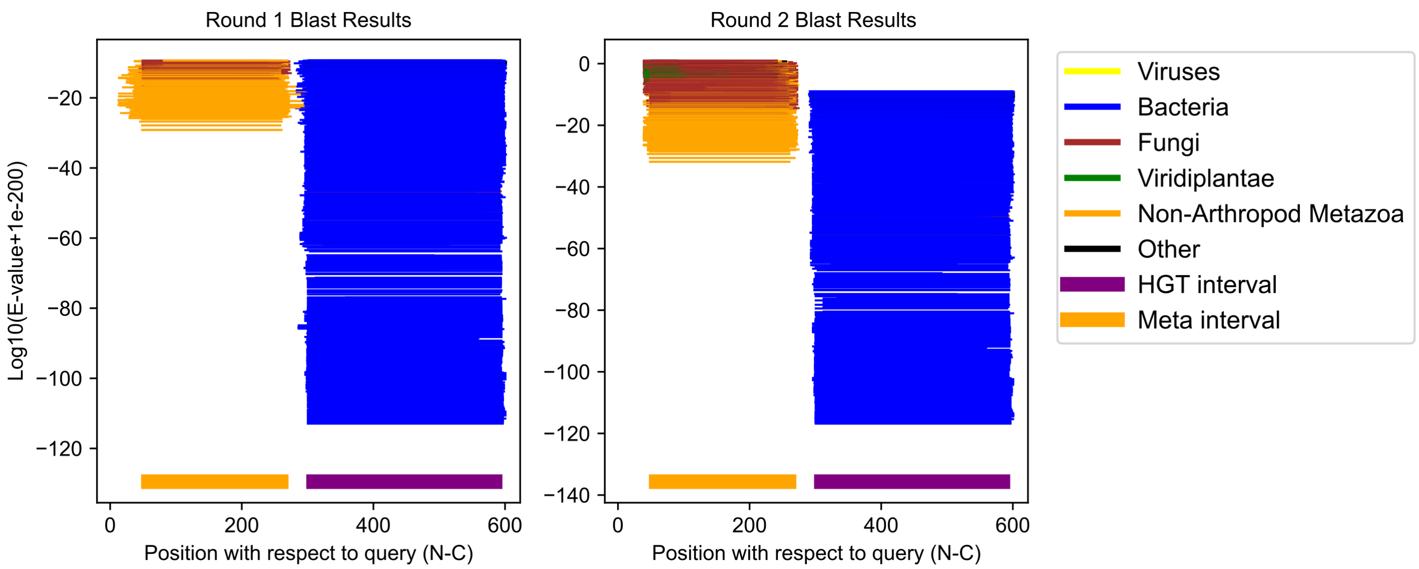

Eurosta solidaginis GCF_040869045.1;XP_067619053.1 (cluster 46)

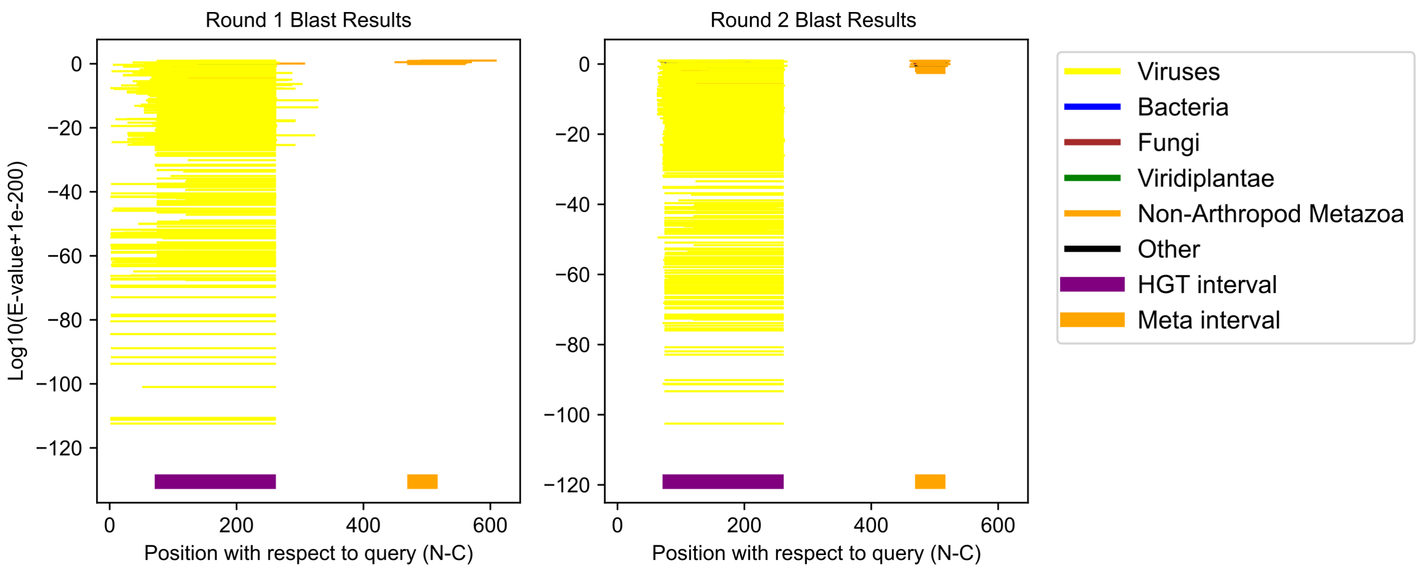

Parasteatoda tepidariorum GCF_043381705.1;XP_015919340.2 (cluster 47)

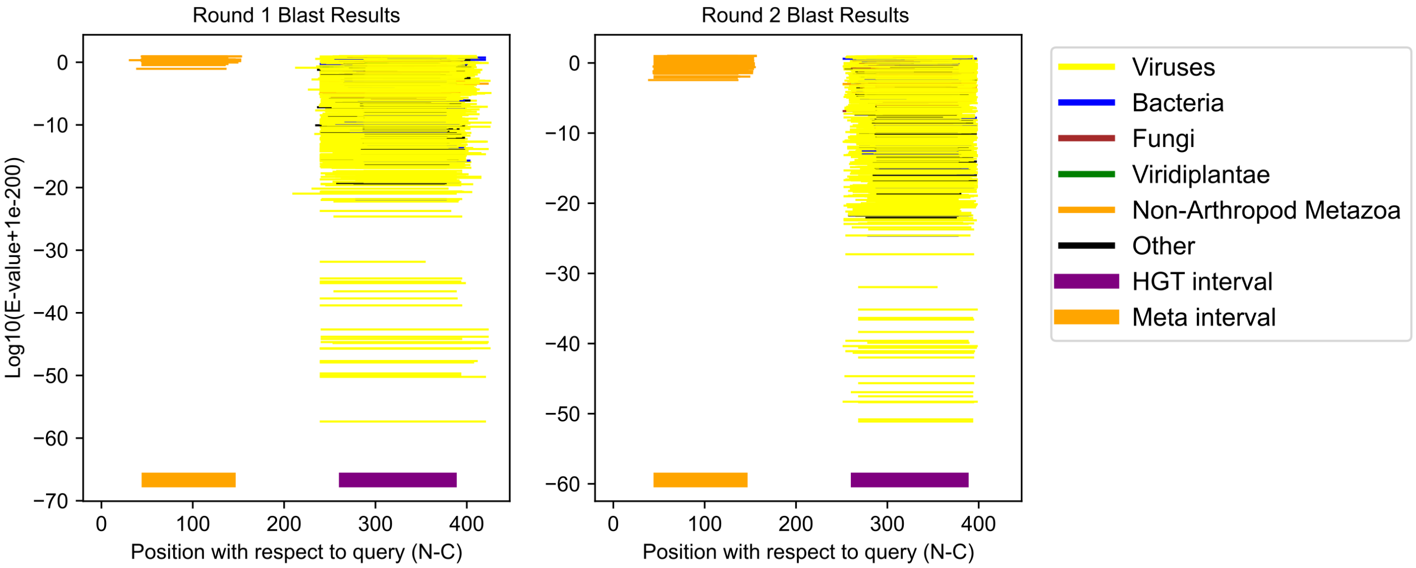

Culicoides brevitarsis GCF_036172545.1;XP_063702760.1 (cluster 48)

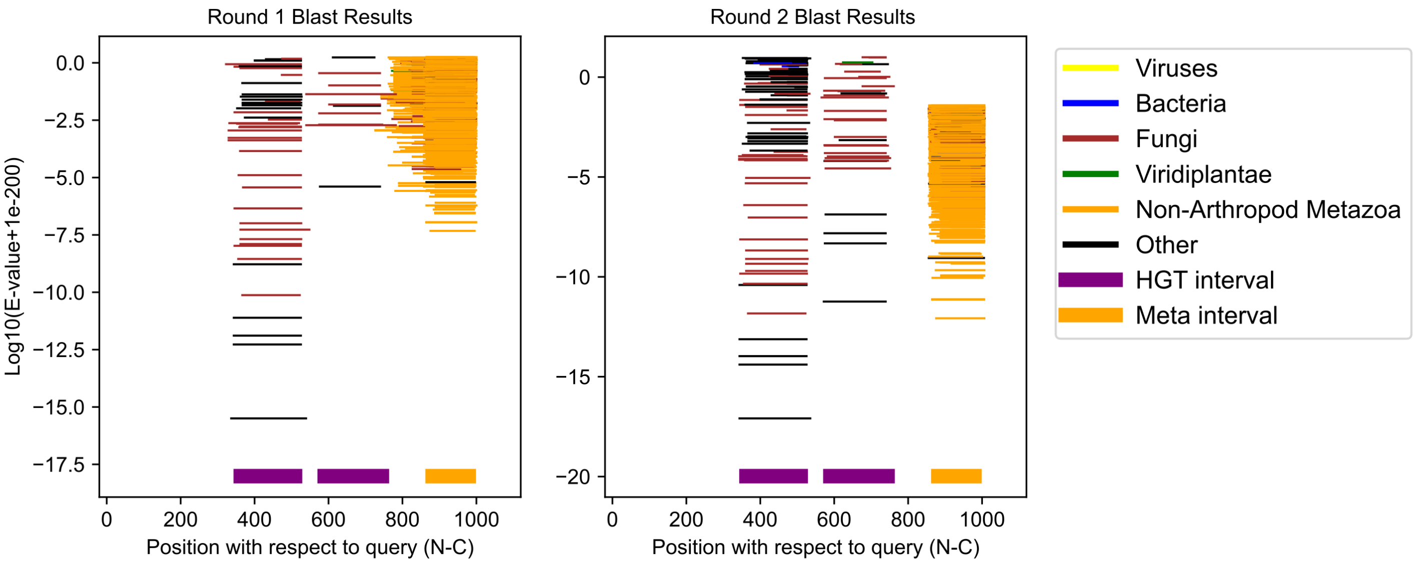

Maniola jurtina GCF_905333055.1;XP_045779580.1 (cluster 49)

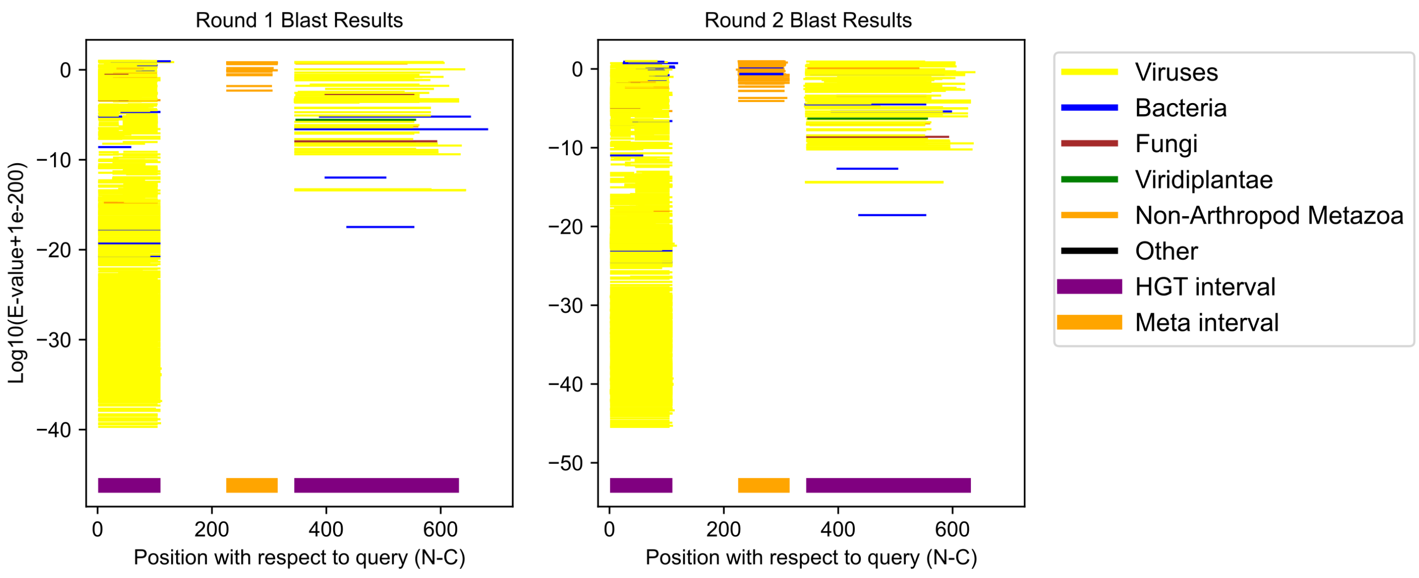

Osmia bicornis bicornis GCF_907164935.1;XP_046143733.1 (cluster 50)

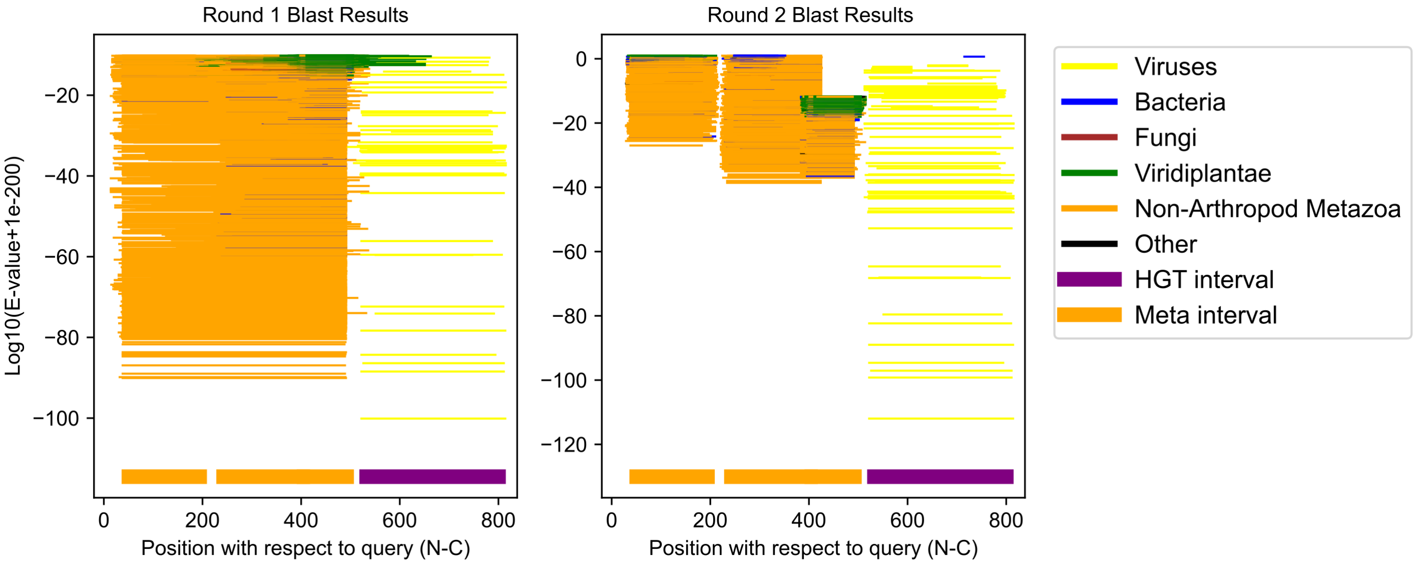

Artemia franciscana GCF_032884065.1;XP_065584755.1 (cluster 51)

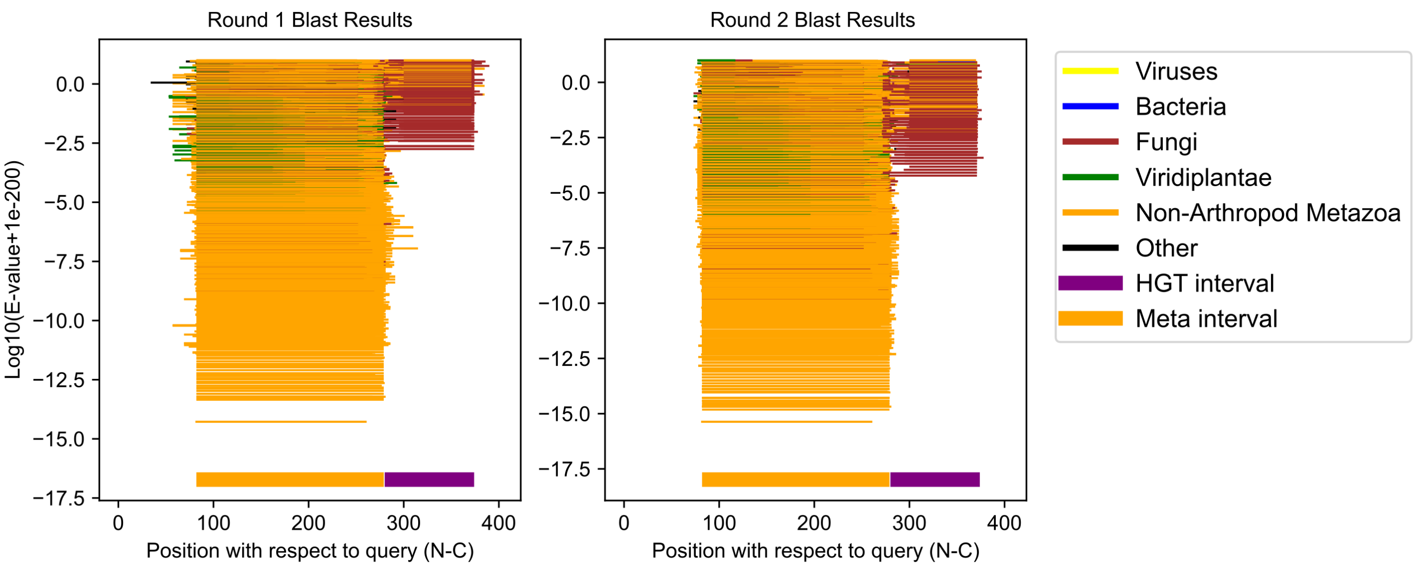

Ischnura elegans GCF_921293095.1;XP_046402901.1 (cluster 52)

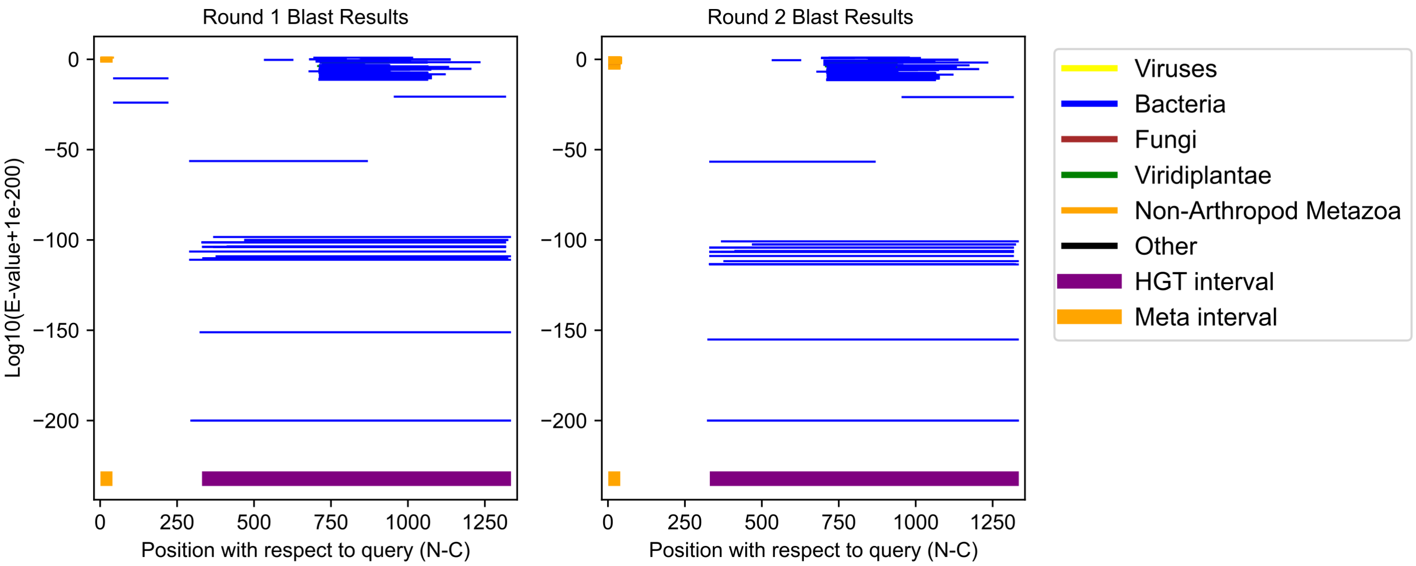

Artemia franciscana GCF_032884065.1;XP_065560524.1 (cluster 53)

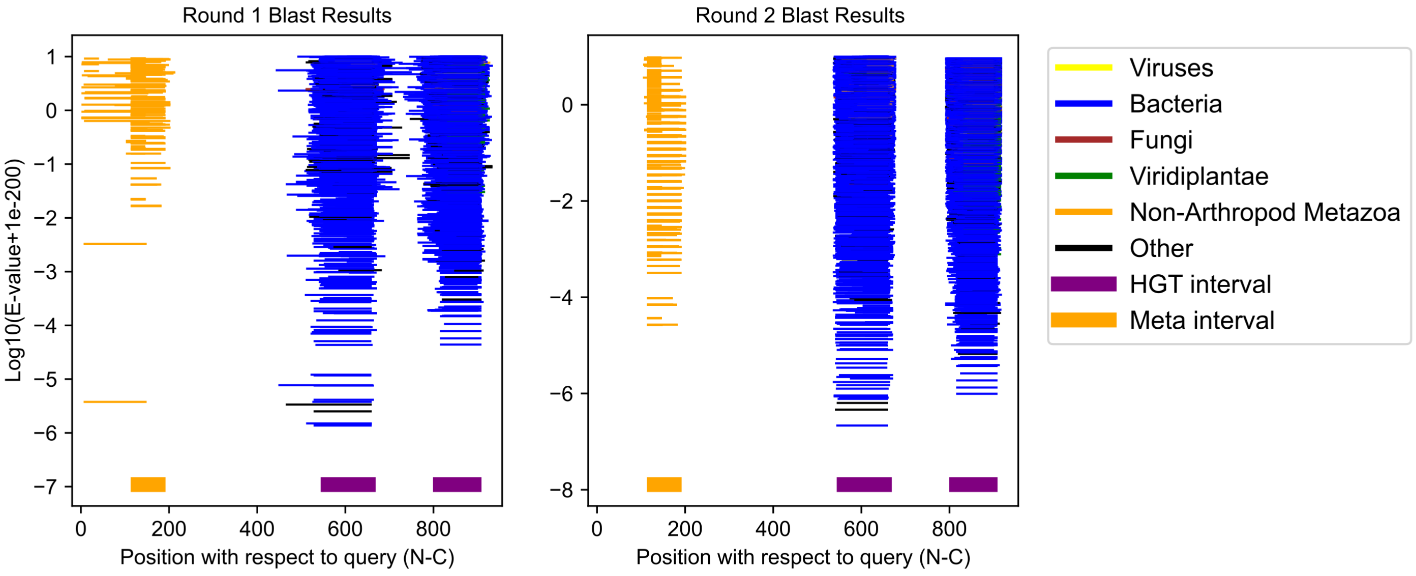

Bacillus rossius redtenbacheri GCF_032445375.1;XP_063242512.1 (cluster 54)

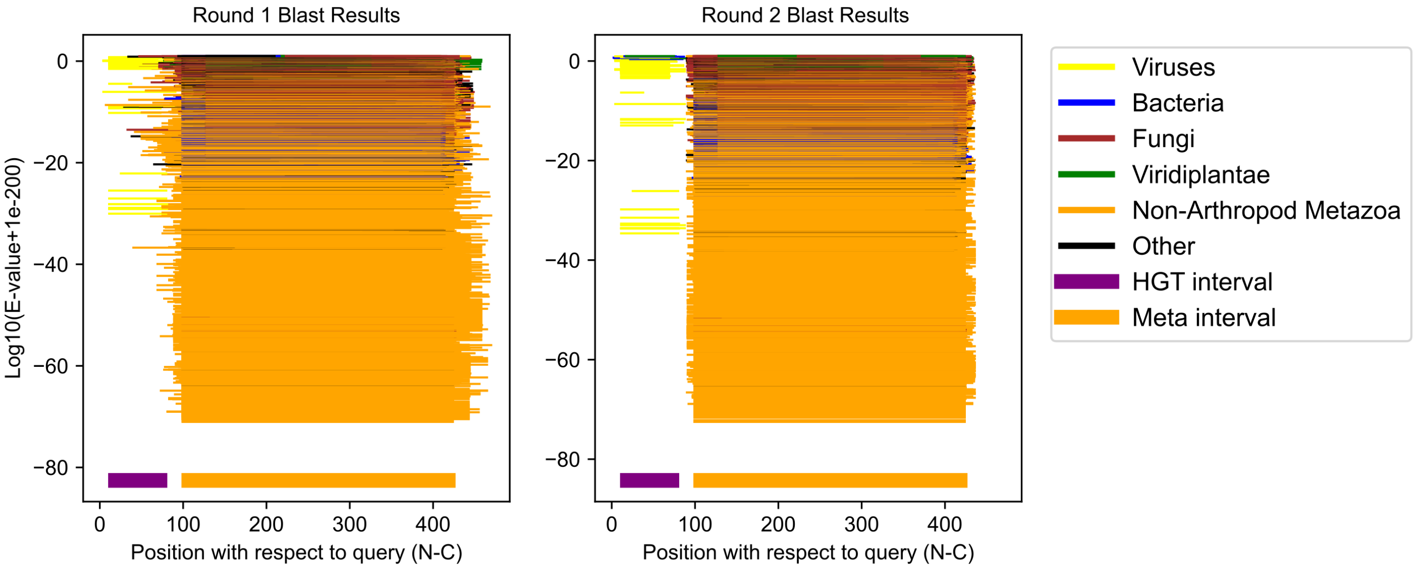

Bacillus rossius redtenbacheri GCF_032445375.1;XP_063239750.1 (cluster 55)

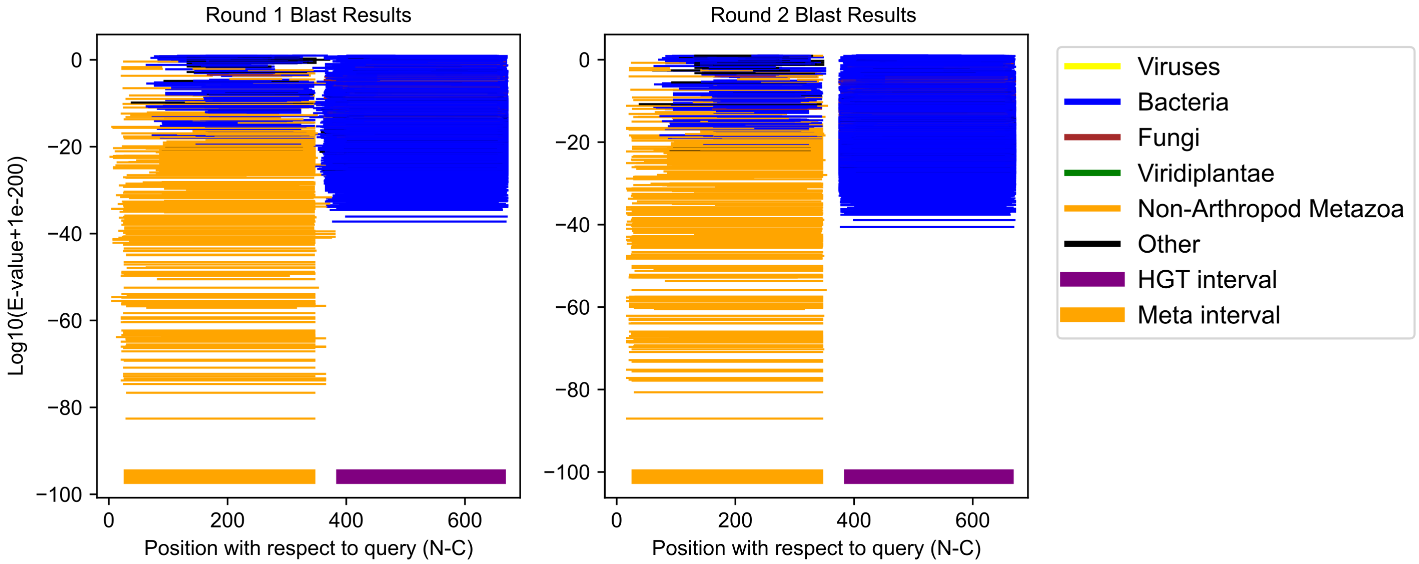

Bemisia tabaci GCF_918797505.1;XP_018903502.2 (cluster 56)

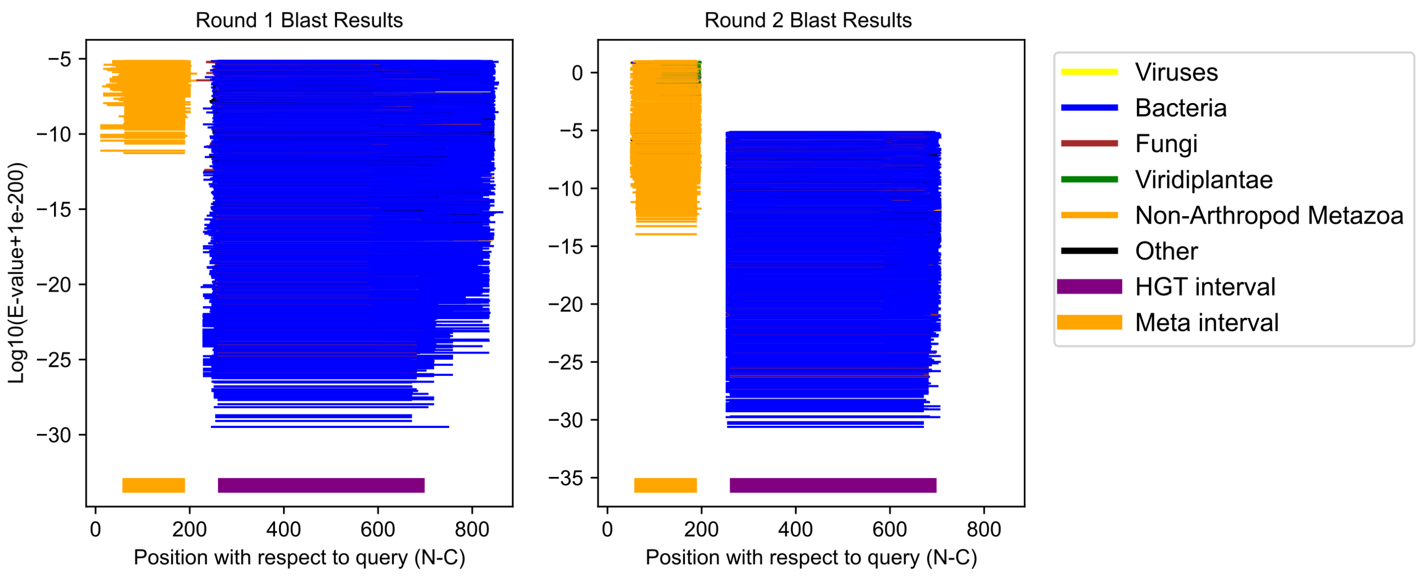

Bacillus rossius redtenbacheri GCF_032445375.1;XP_063236389.1 (cluster 57)

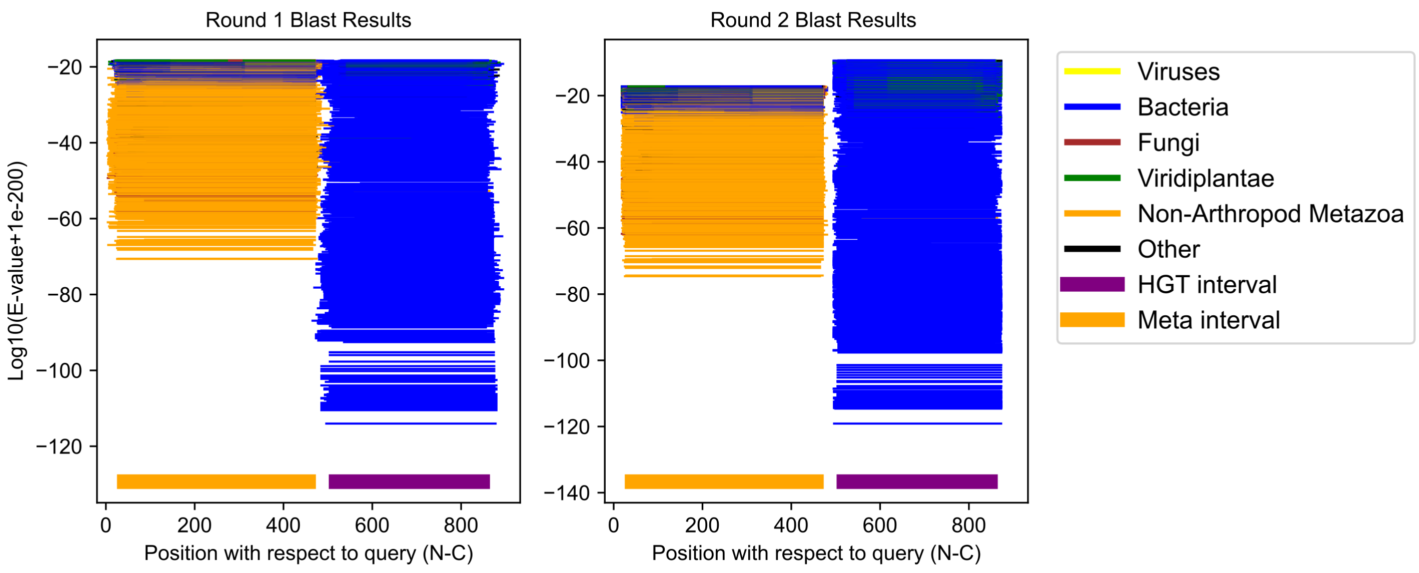

Tribolium castaneum GCF_031307605.1;XP_064214057.1 (cluster 58)

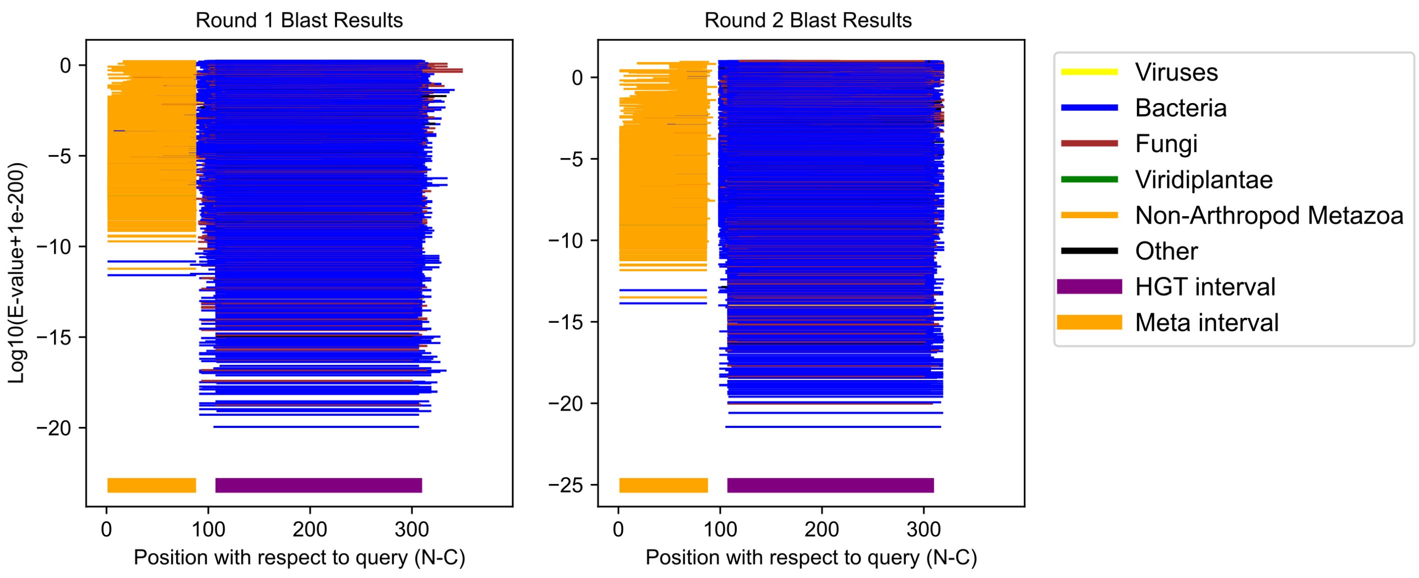

Schistocerca cancellata GCF_023864275.1;XP_049785902.1 (cluster 59)

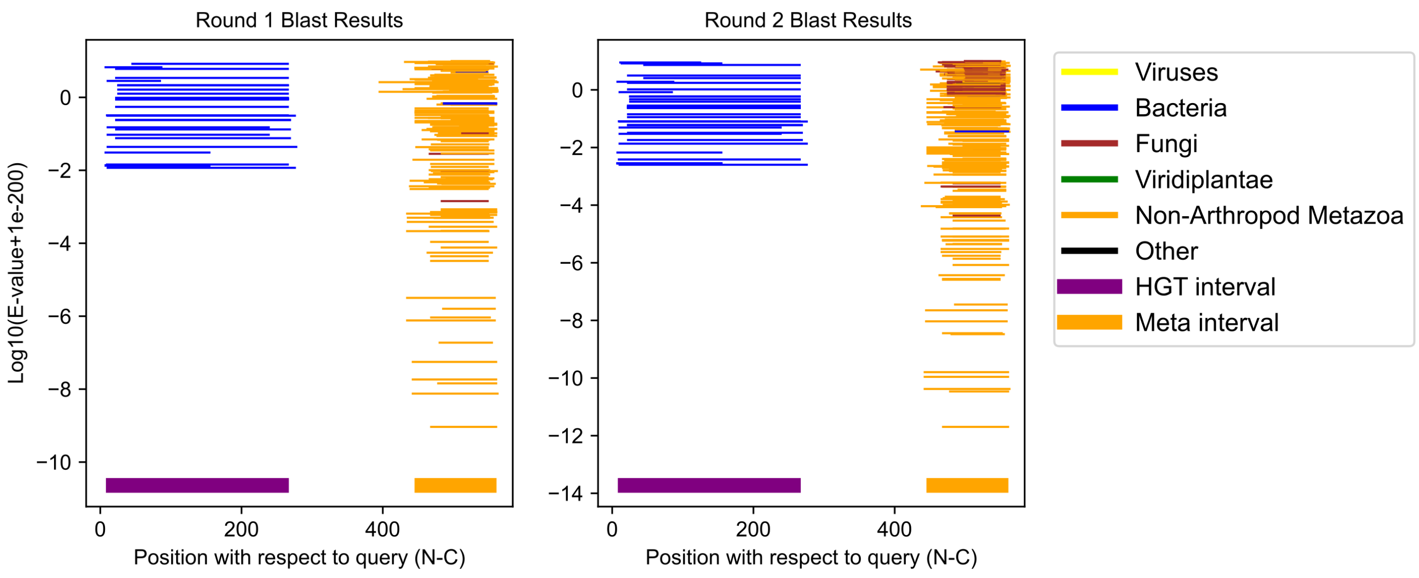

Eurytemora carolleeae GCF_000591075.1;XP_023324156.1 (cluster 60)

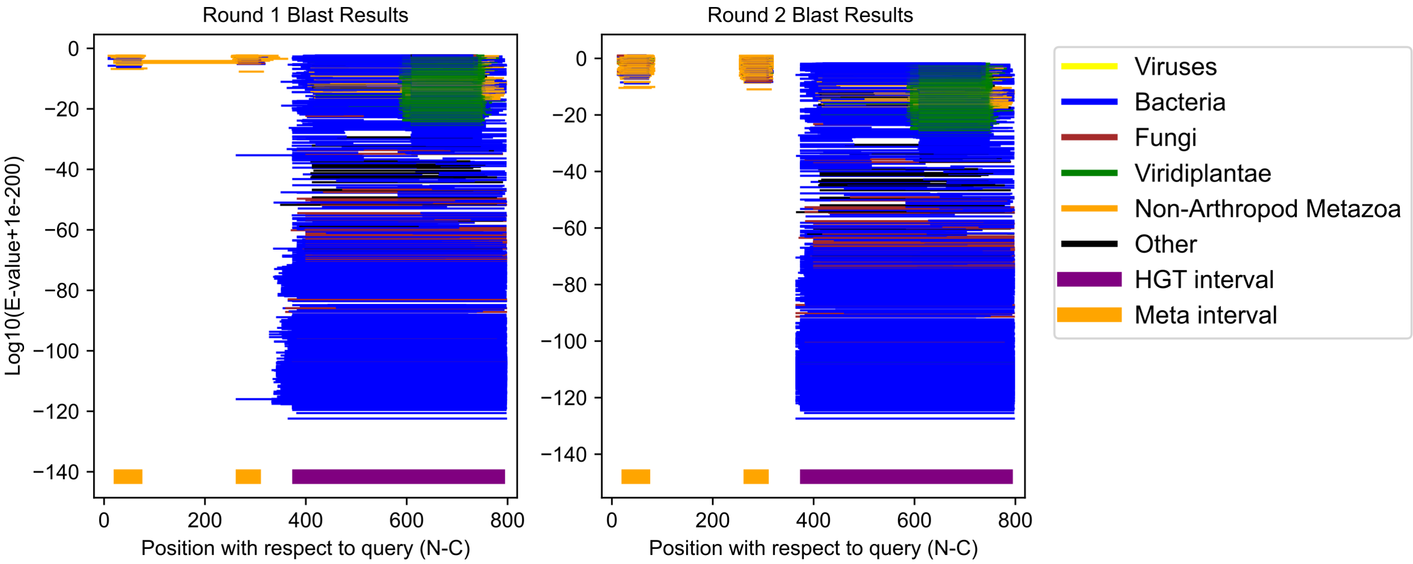

Daphnia pulicaria GCF_021234035.1;XP_046646423.1 (cluster 61)

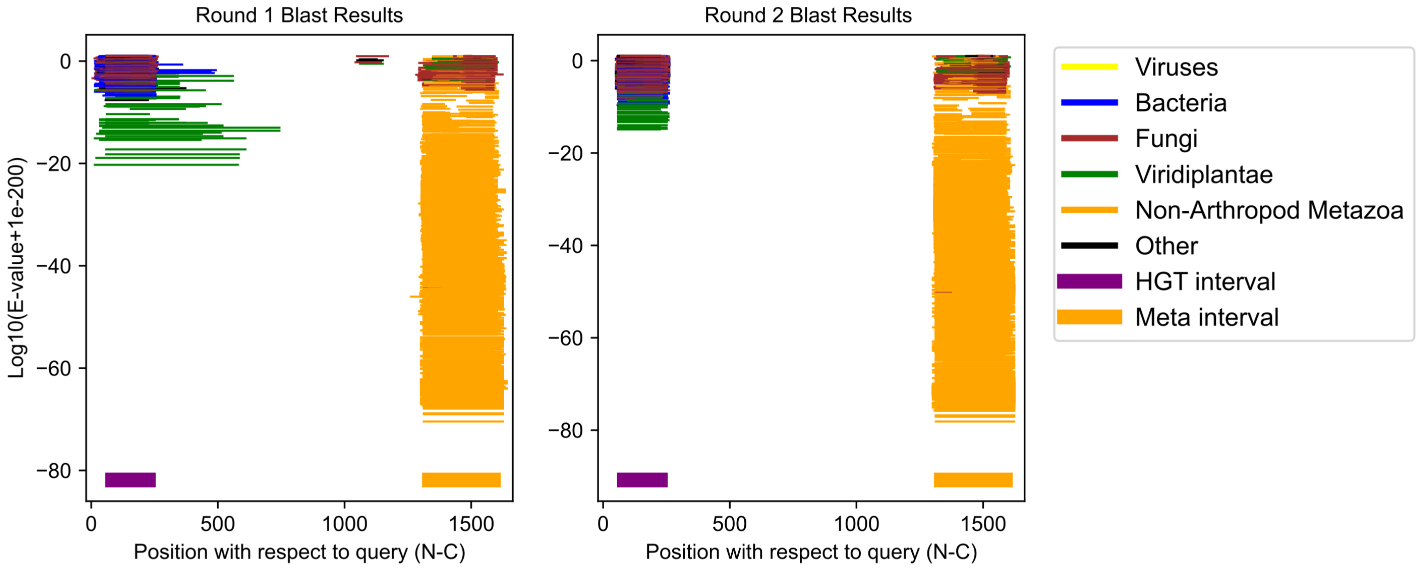

Neodiprion pinetum GCF_021155775.2;XP_068991025.1 (cluster 62)

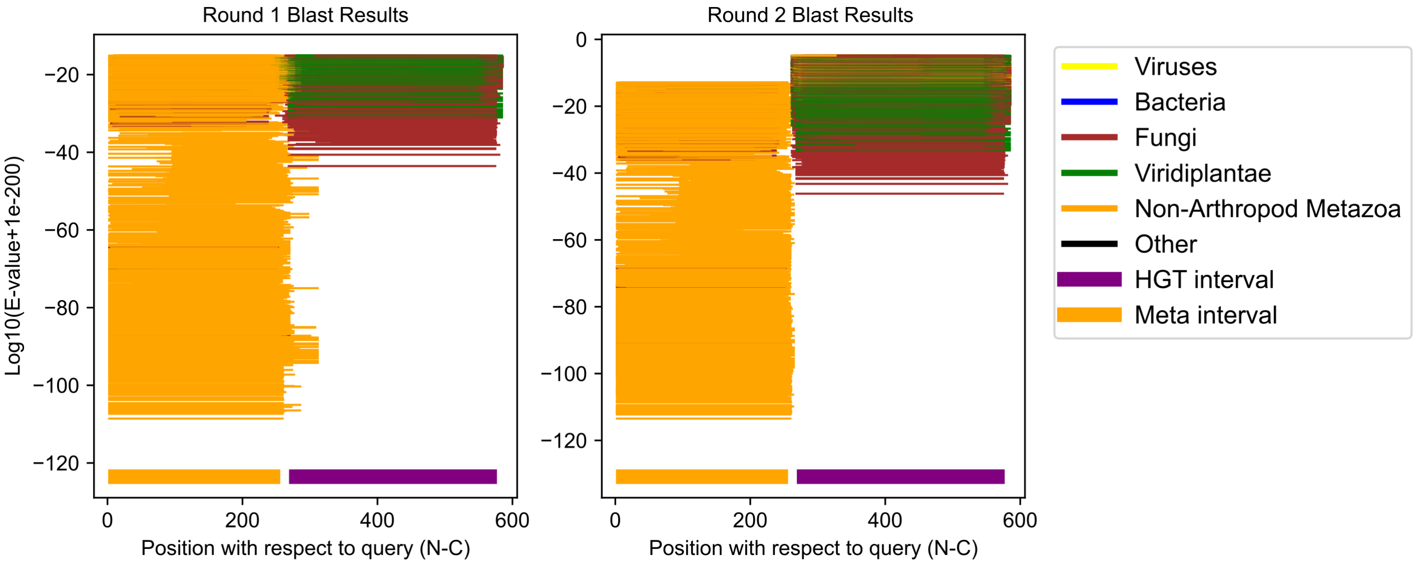

Hyposmocoma kahamanoa GCF_003589595.1;XP_026318555.1 (cluster 63)

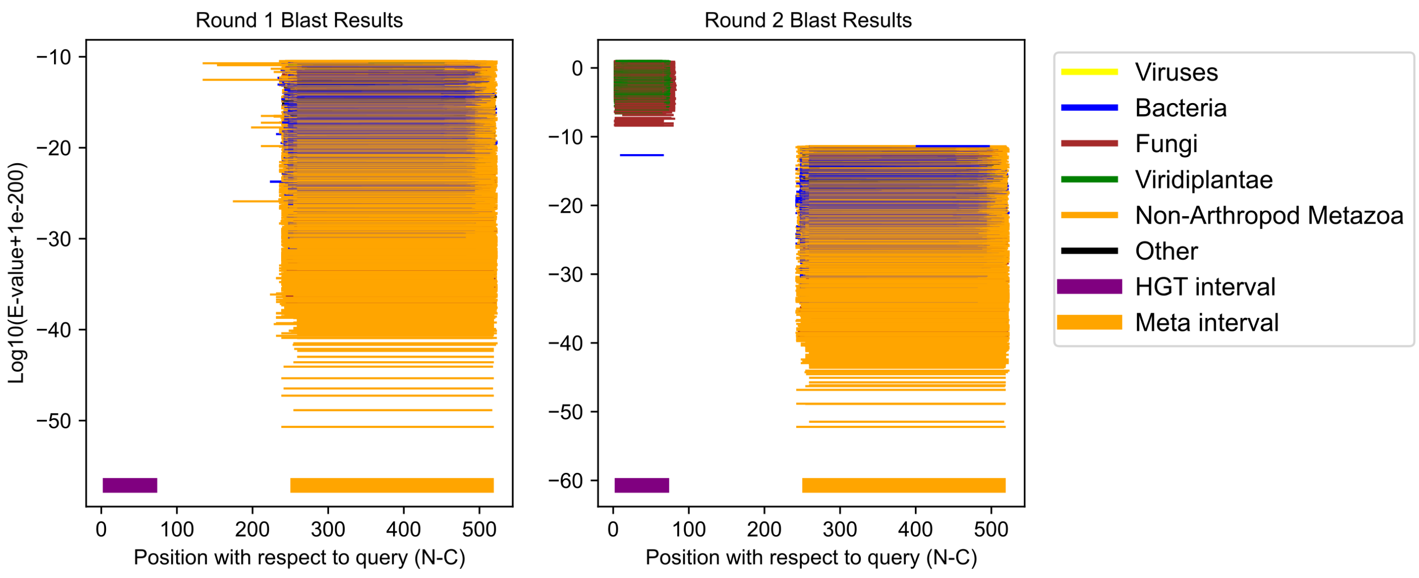

Teleopsis dalmanni GCF_002237135.1;XP_037951667.1 (cluster 64)

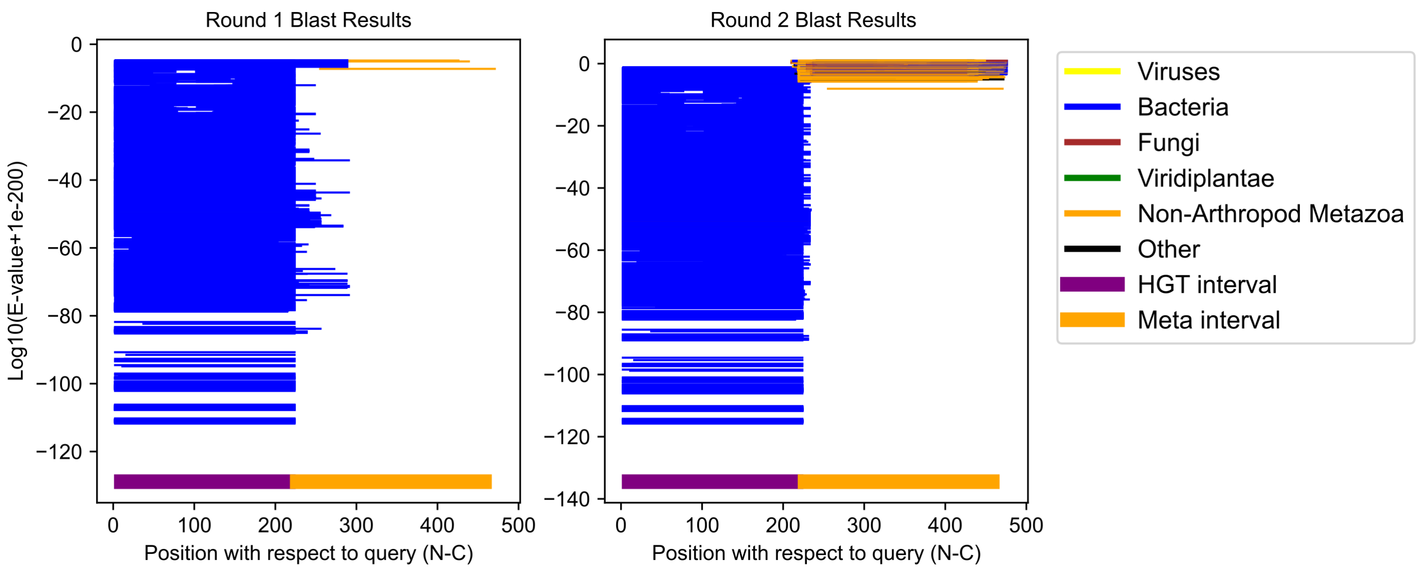

Folsomia candida GCF_002217175.1;XP_035708240.1 (cluster 65)

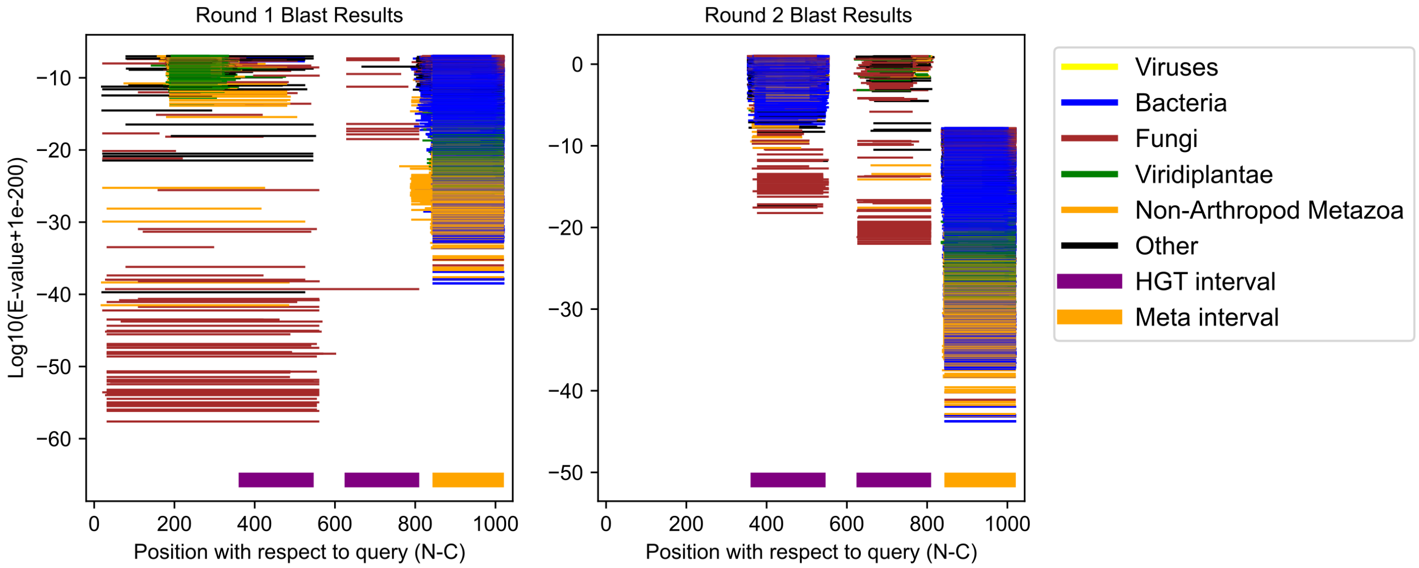

Folsomia candida GCF_002217175.1;XP_035708168.1 (cluster 66)

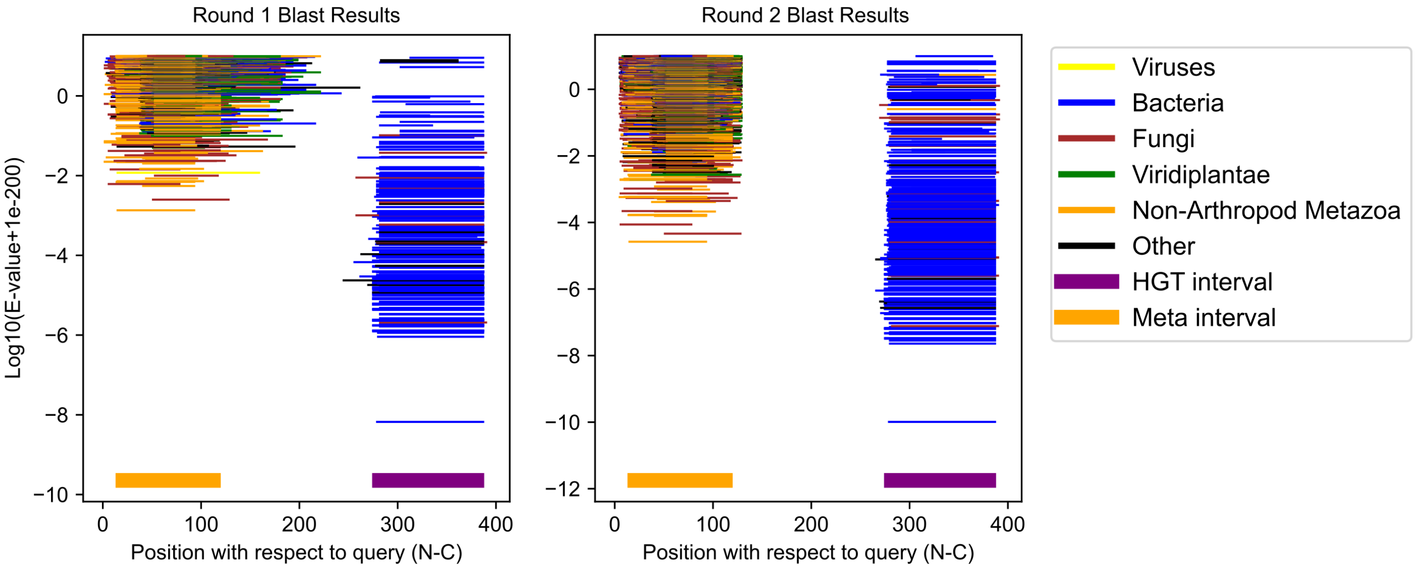

Folsomia candida GCF_002217175.1;XP_021965719.2 (cluster 67)

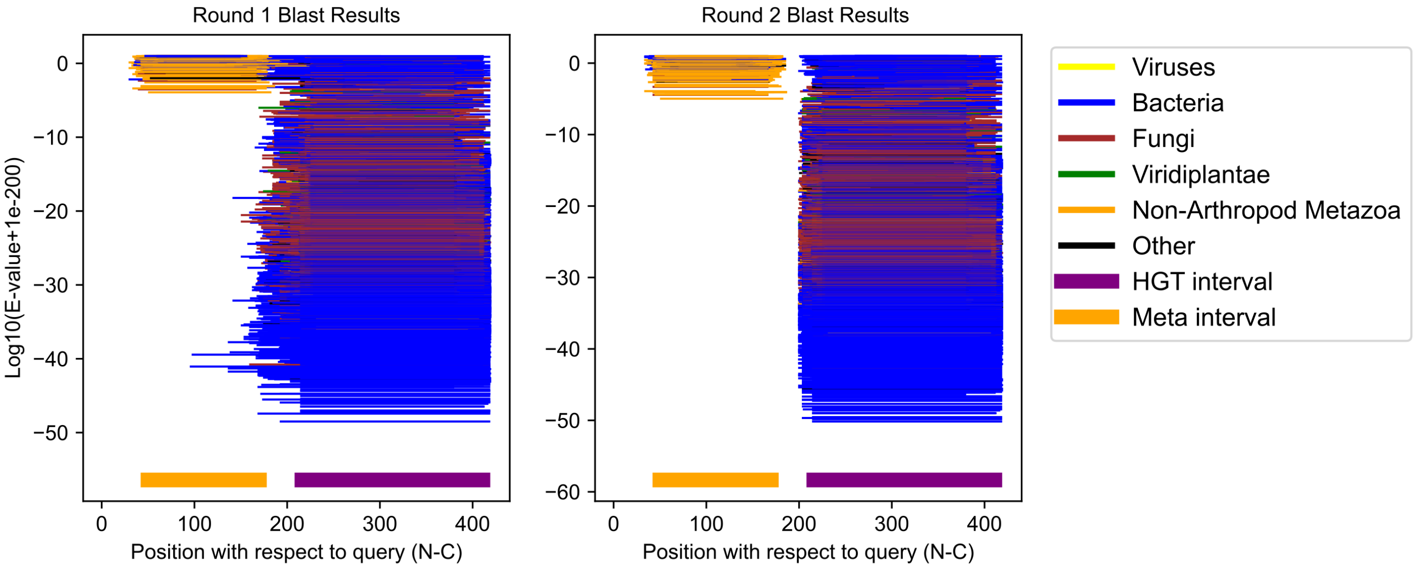

Folsomia candida GCF_002217175.1;XP_021965073.1 (cluster 68)

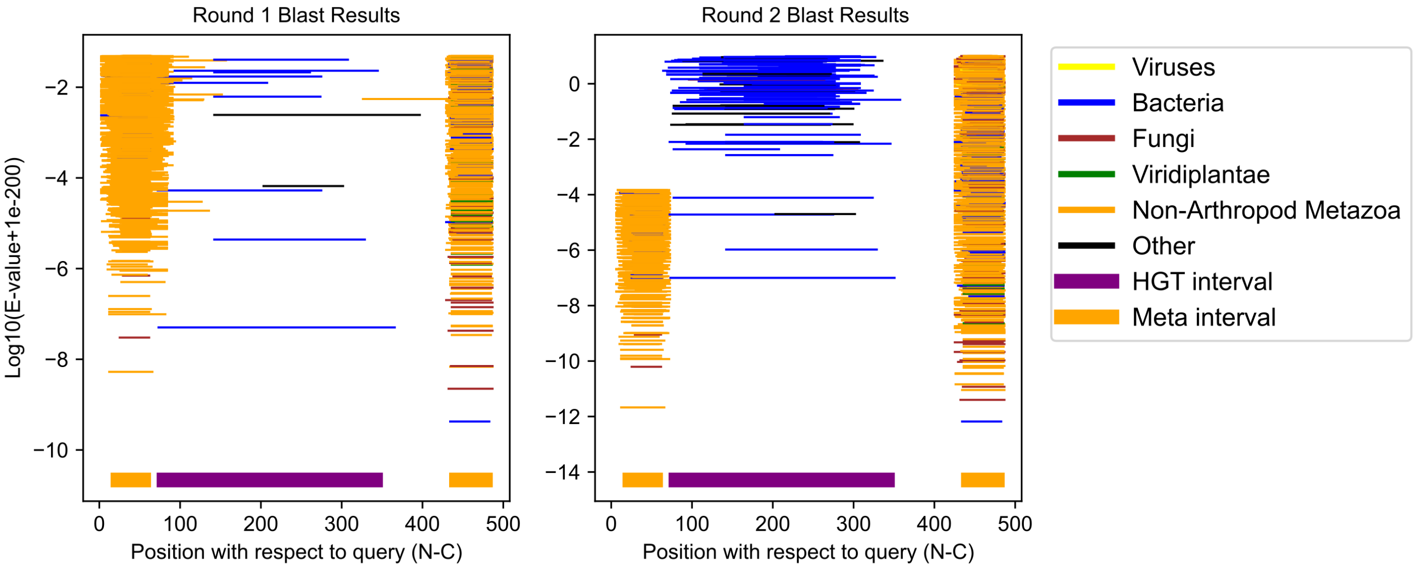

Folsomia candida GCF_002217175.1;XP_021960153.2 (cluster 69)

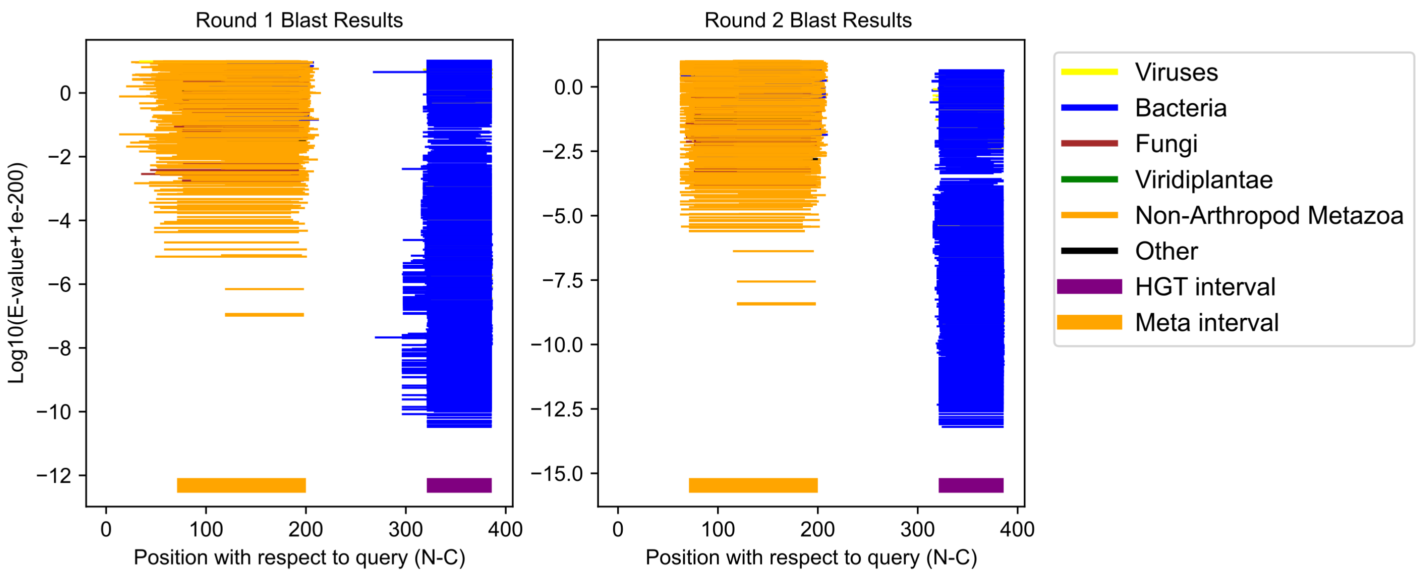

Folsomia candida GCF_002217175.1;XP_021956151.1 (cluster 70)

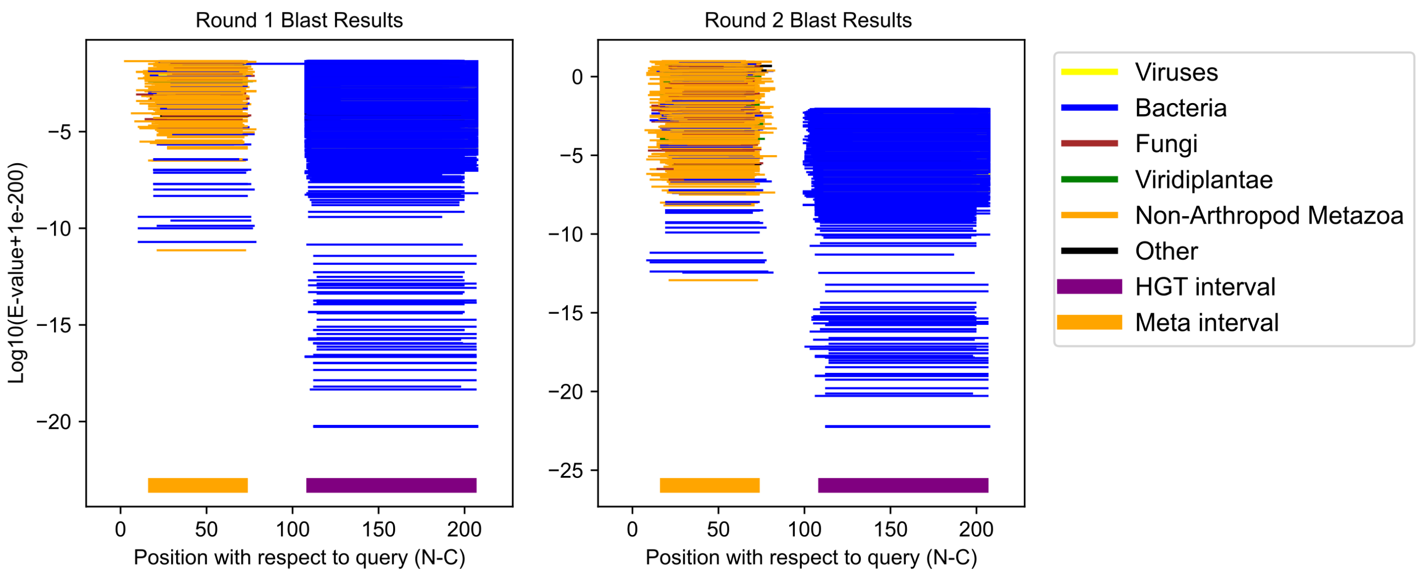

Folsomia candida GCF_002217175.1;XP_021953192.2 (cluster 71)

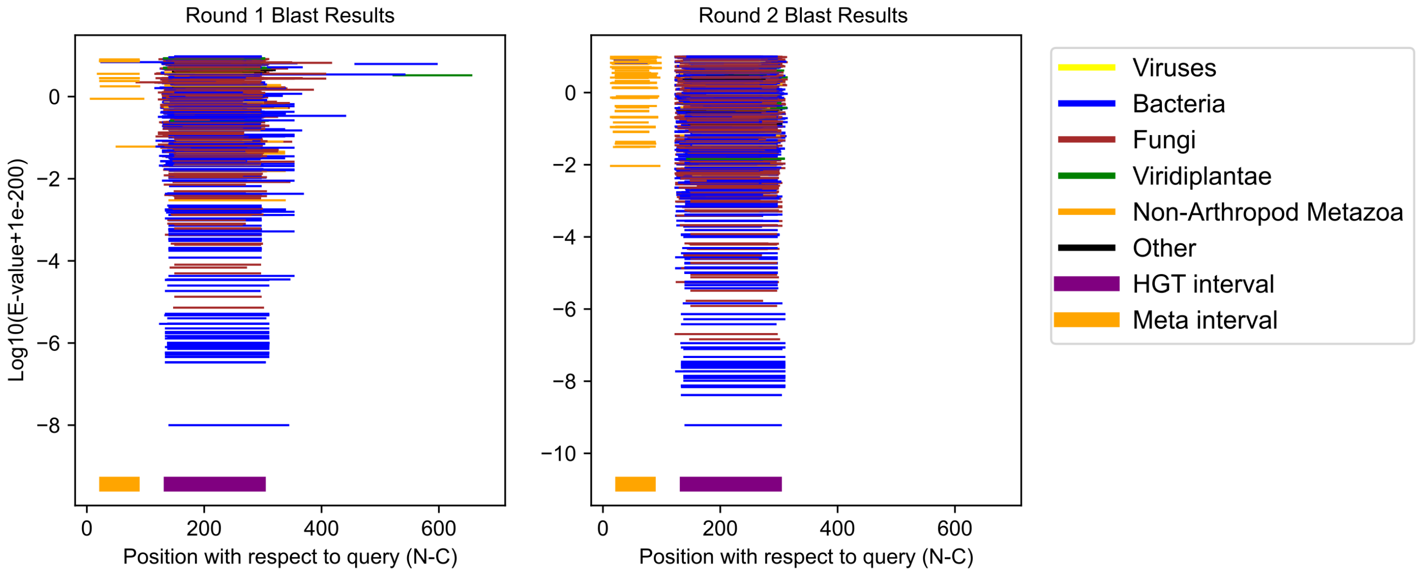

Folsomia candida GCF_002217175.1;XP_021952783.1 (cluster 72)

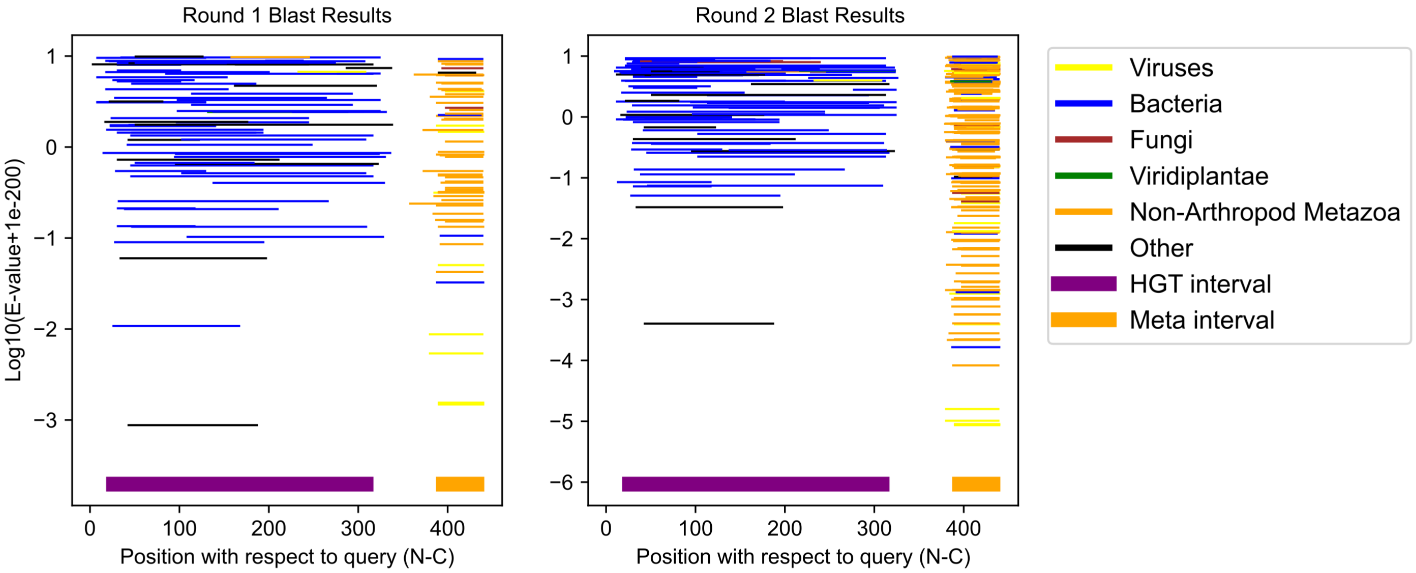

Dermatophagoides pteronyssinus GCF_001901225.1;XP_027204138.1 (cluster 73)

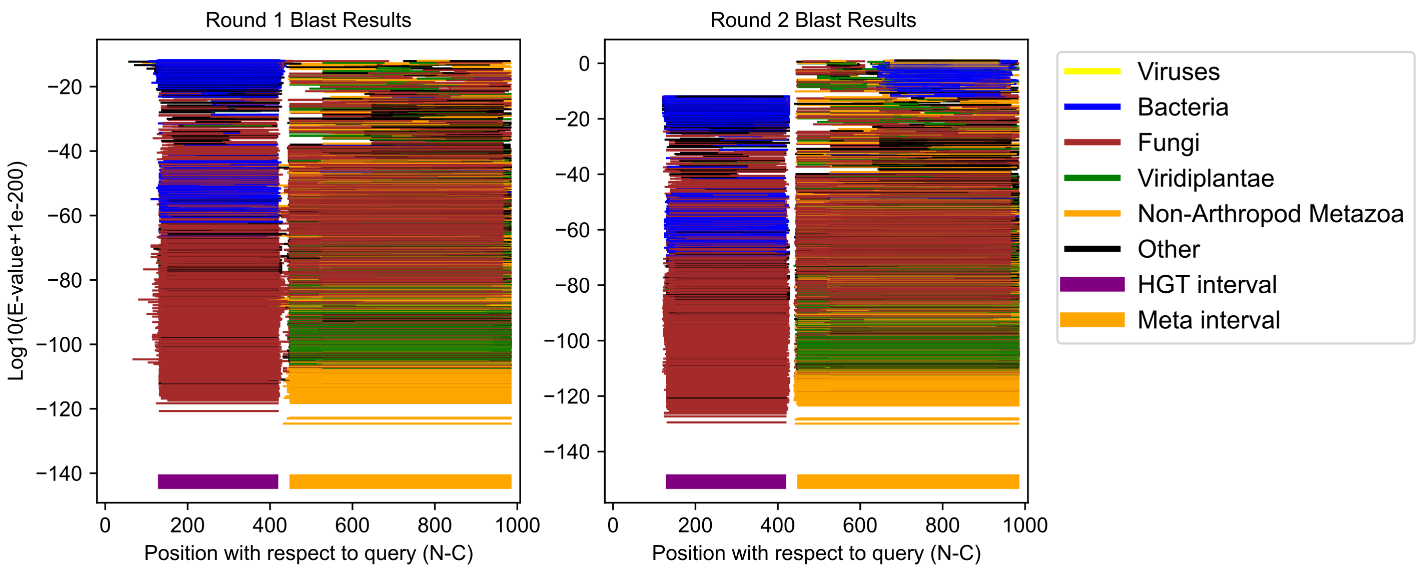

Dermatophagoides pteronyssinus GCF_001901225.1;XP_027200358.1 (cluster 74)

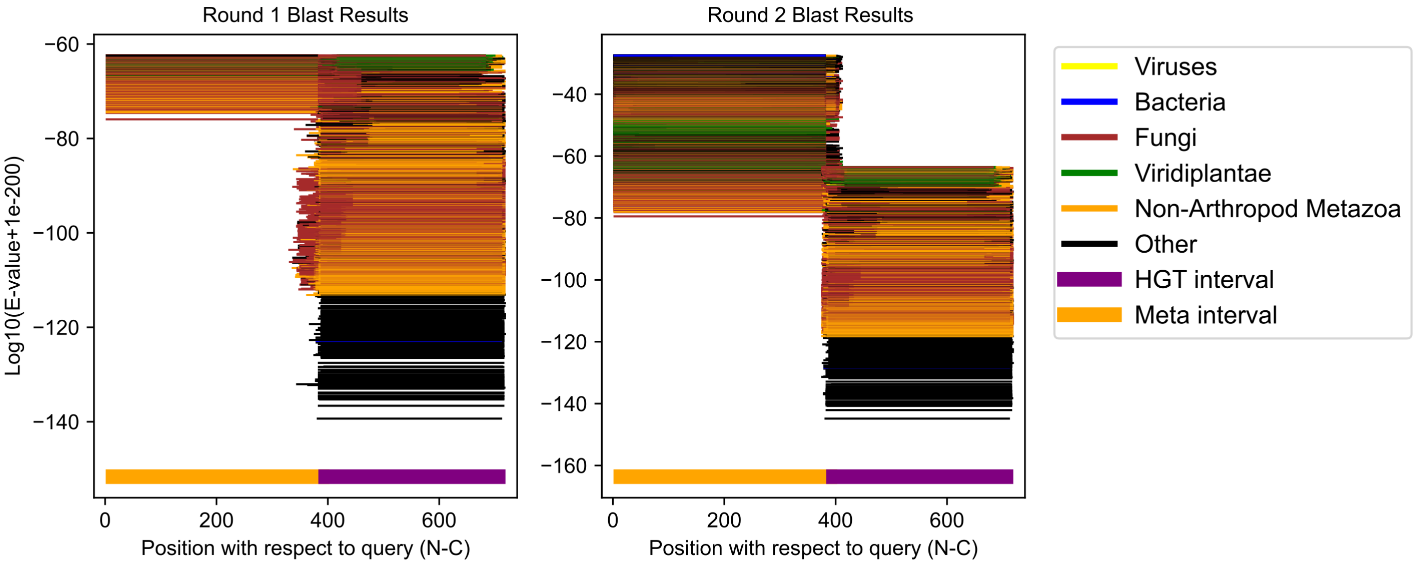

Myzus persicae GCF_001856785.1;XP_022173178.1 (cluster 75)

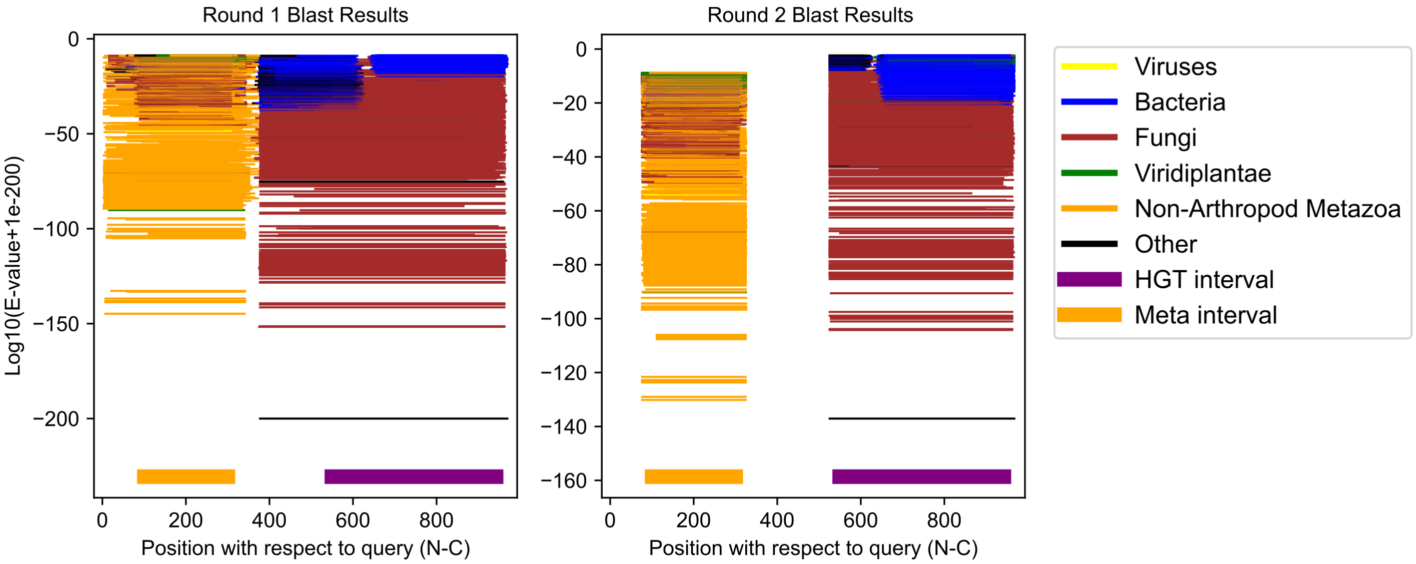

Frankliniella occidentalis GCF_000697945.3;XP_052125515.1 (cluster 76)

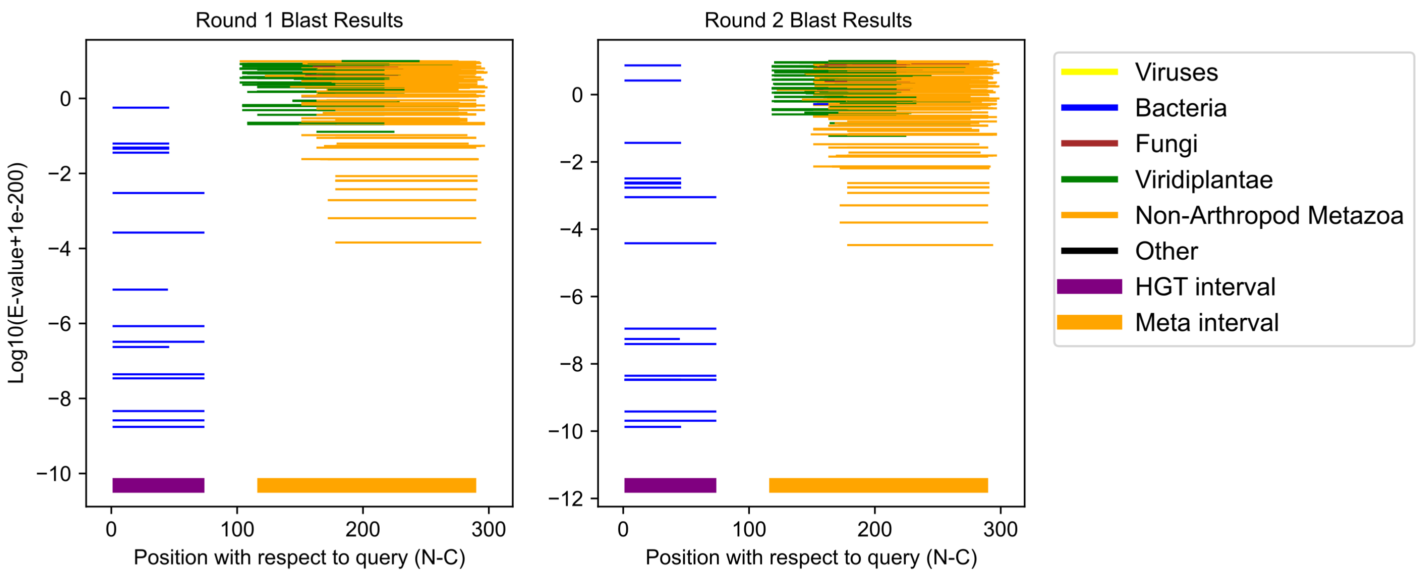

Eurytemora carolleeae GCF_000591075.1;XP_023345006.1 (cluster 77)

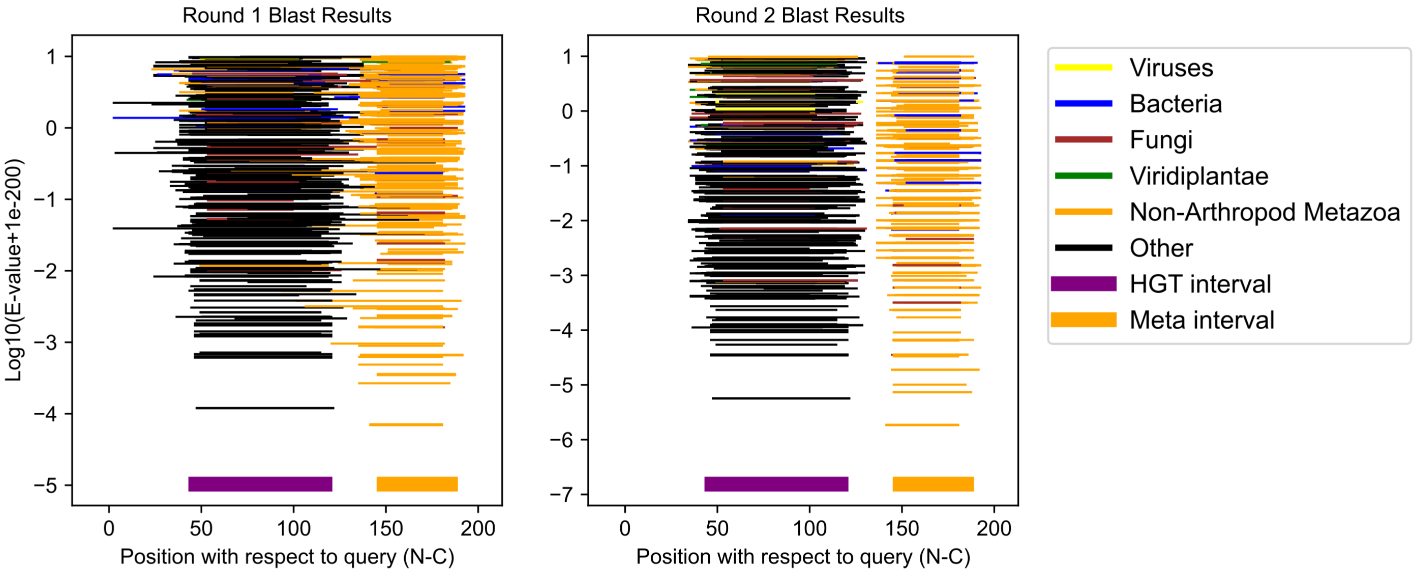

Eurytemora carolleeae GCF_000591075.1;XP_023332299.1 (cluster 78)

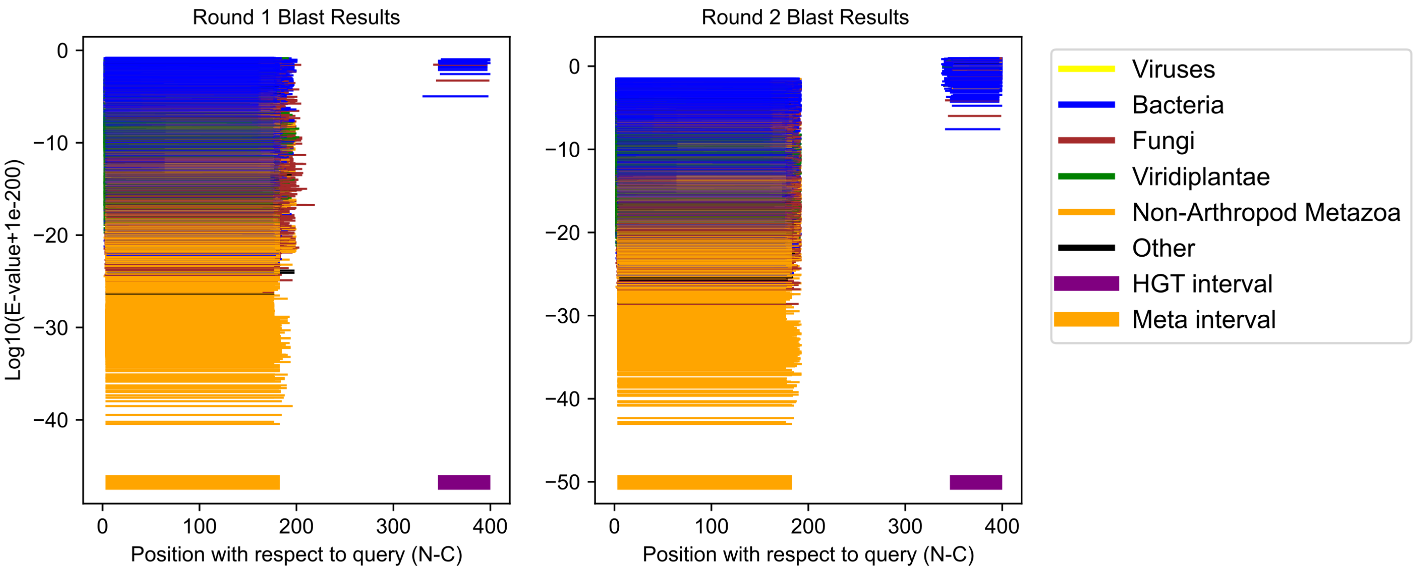

### Eurytemora carolleeae GCF_000591075.1;XP_023329593.1 (cluster 79)

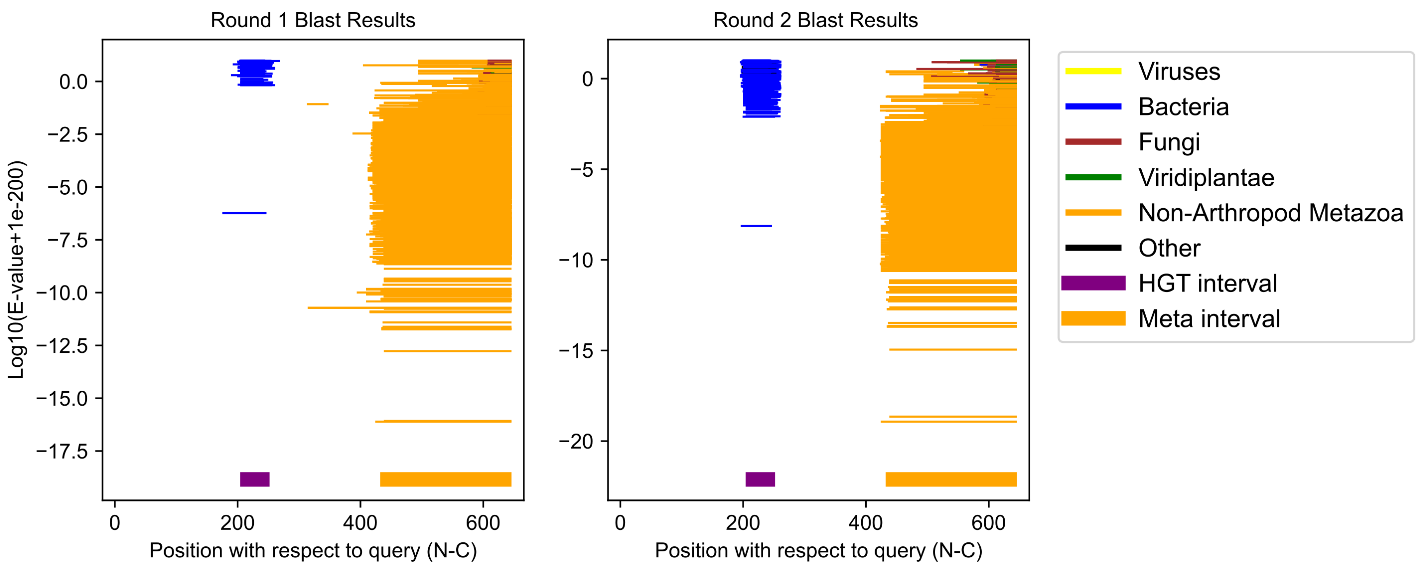

Eurytemora carolleeae GCF_000591075.1;XP_023328891.1 (cluster 80)

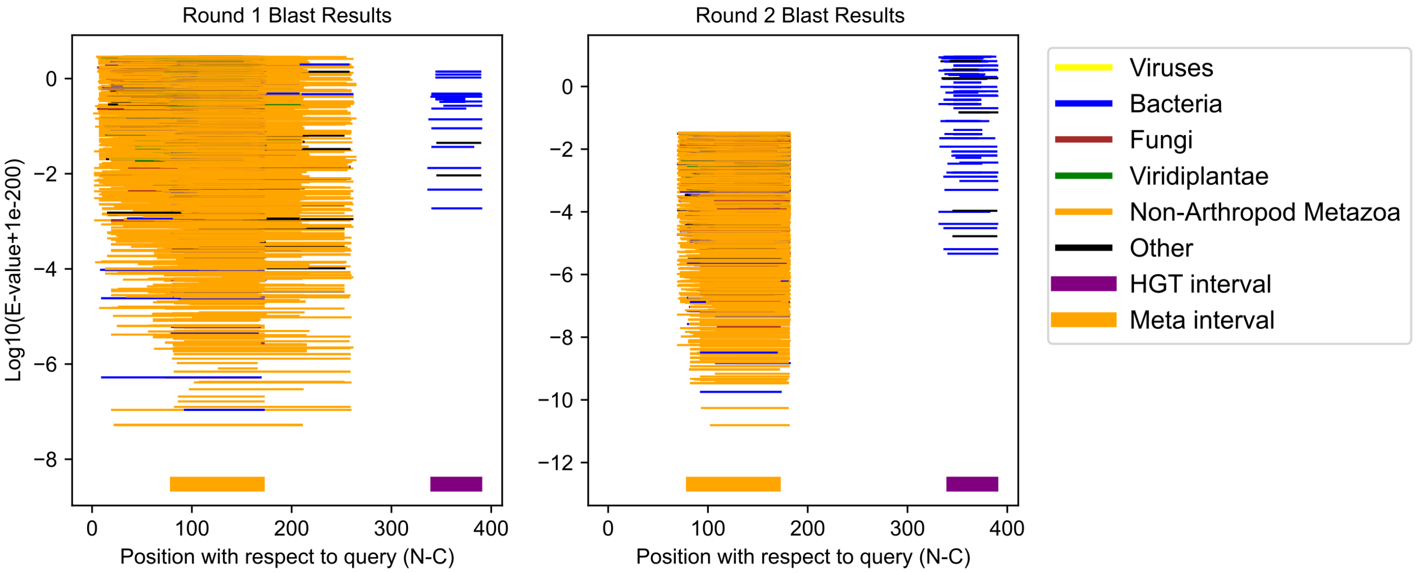

Eurytemora carolleeae GCF_000591075.1;XP_023325437.1 (cluster 81)

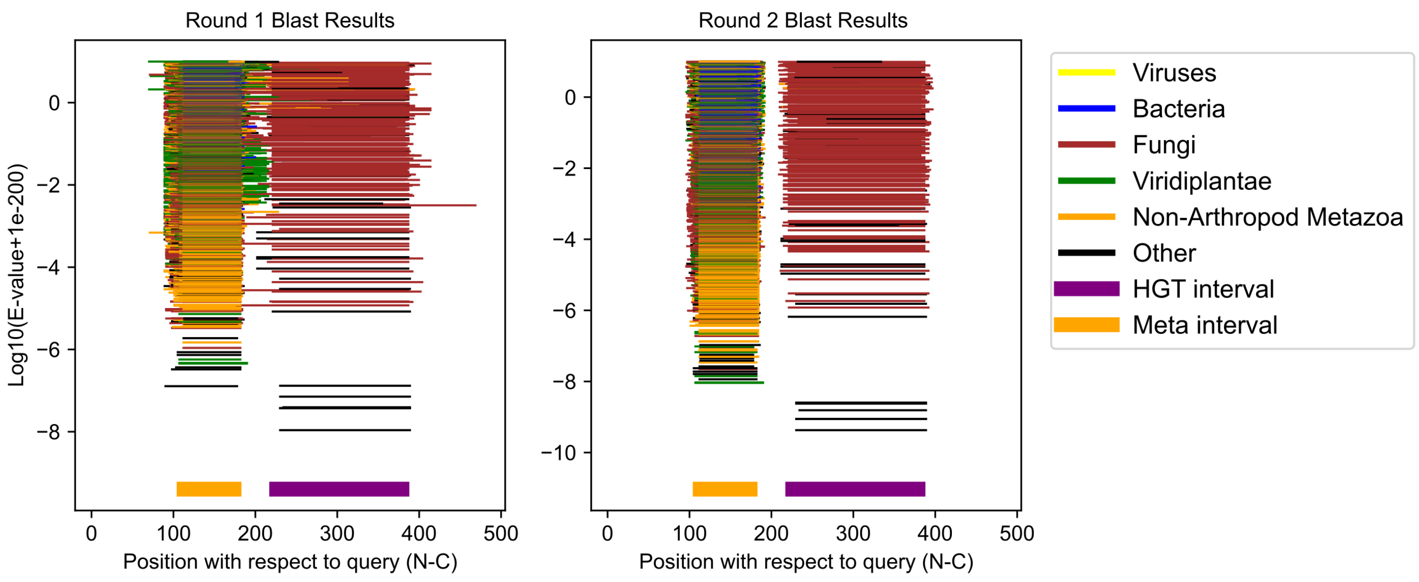

Ostrinia furnacalis GCF_004193835.3;XP_028168683.1 (cluster 82)

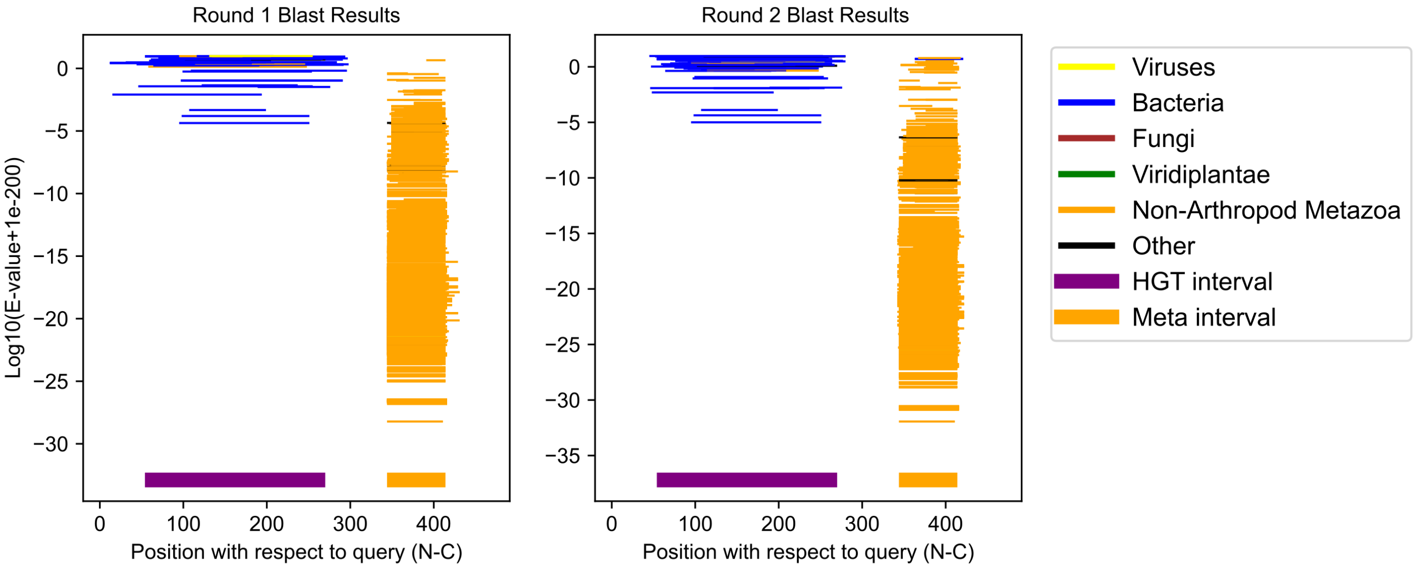

Acyrthosiphon pisum GCF_005508785.2;XP_029341141.1 (cluster 83)

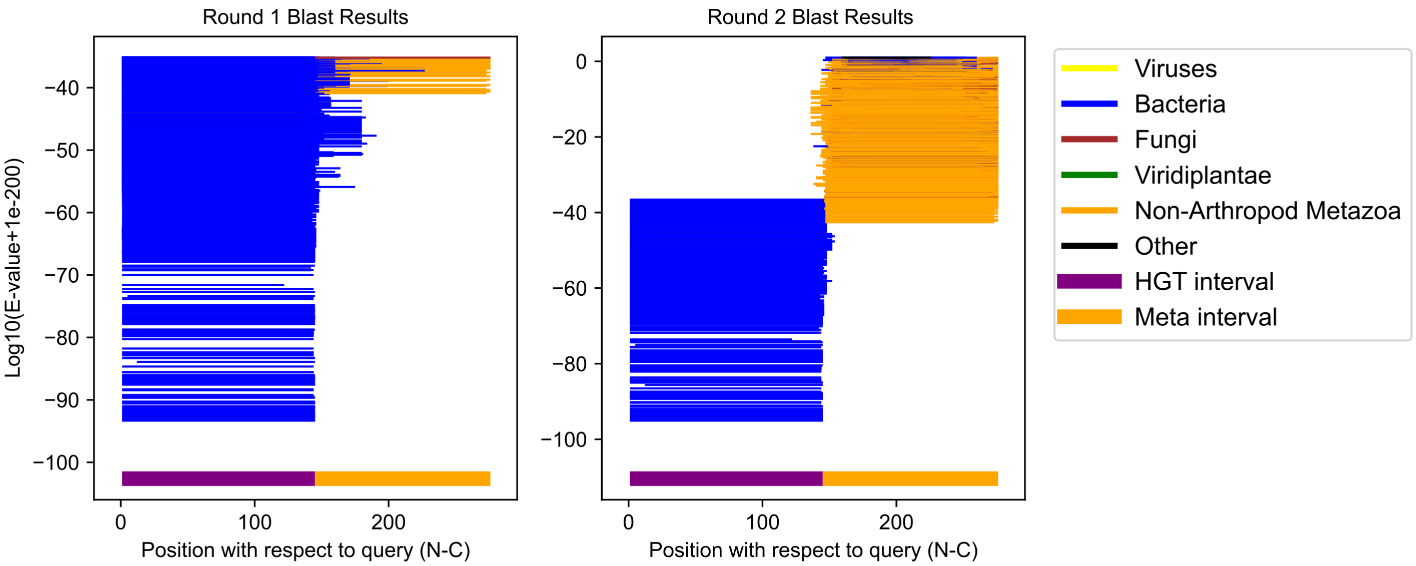

Tigriopus californicus GCF_007210705.1;XP_059093711.1 (cluster 84)

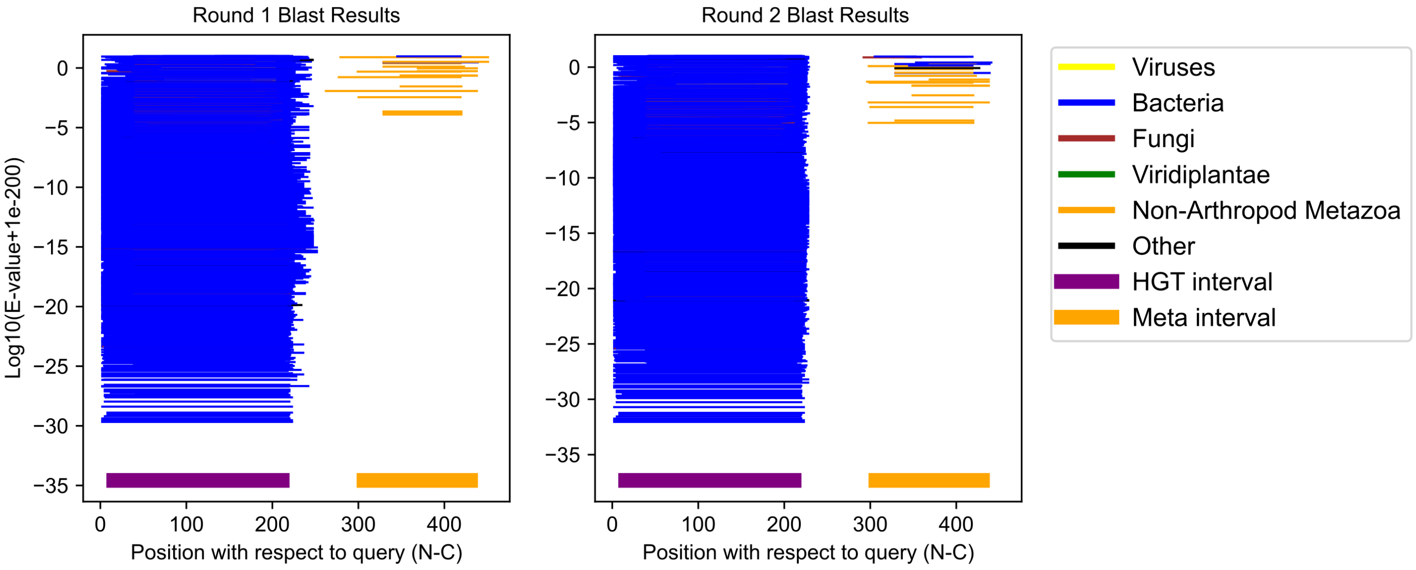

Bradysia coprophila GCF_014529535.1;XP_037041958.1 (cluster 85)

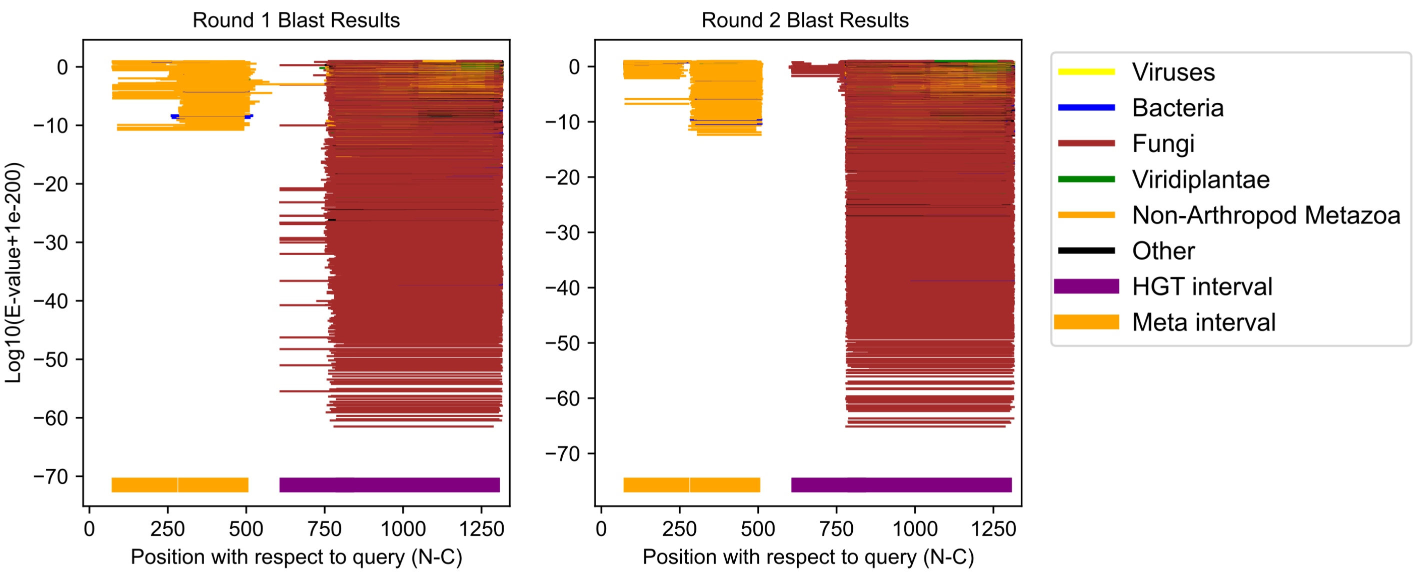

Dendroctonus ponderosae GCF_020466585.1;XP_048520150.1 (cluster 86)

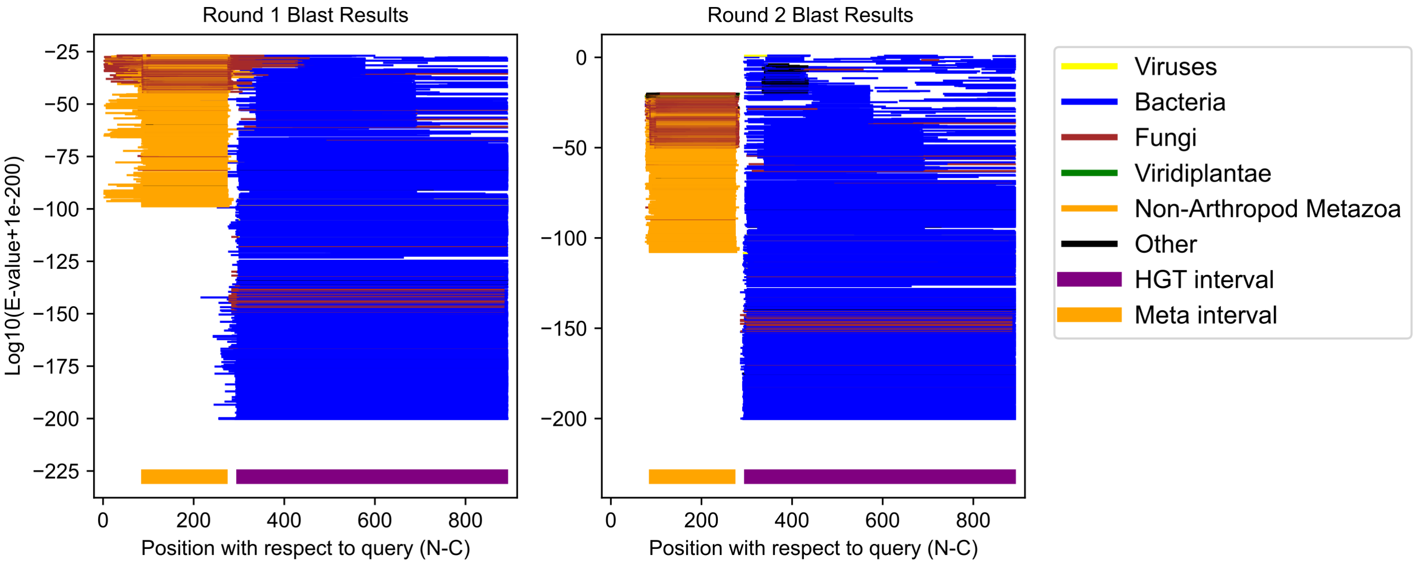

Eurytemora carolleeae GCF_000591075.1;XP_023324698.1 (cluster 87)

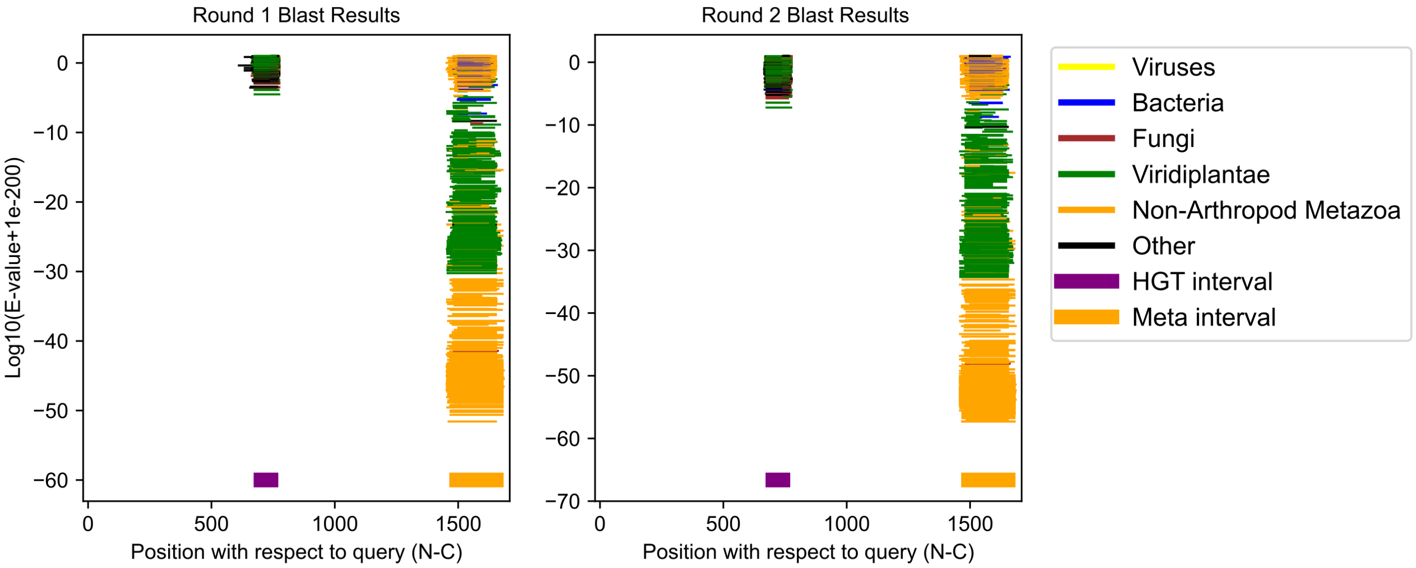

Ixodes scapularis GCF_016920785.2;XP_040063786.1 (cluster 88)

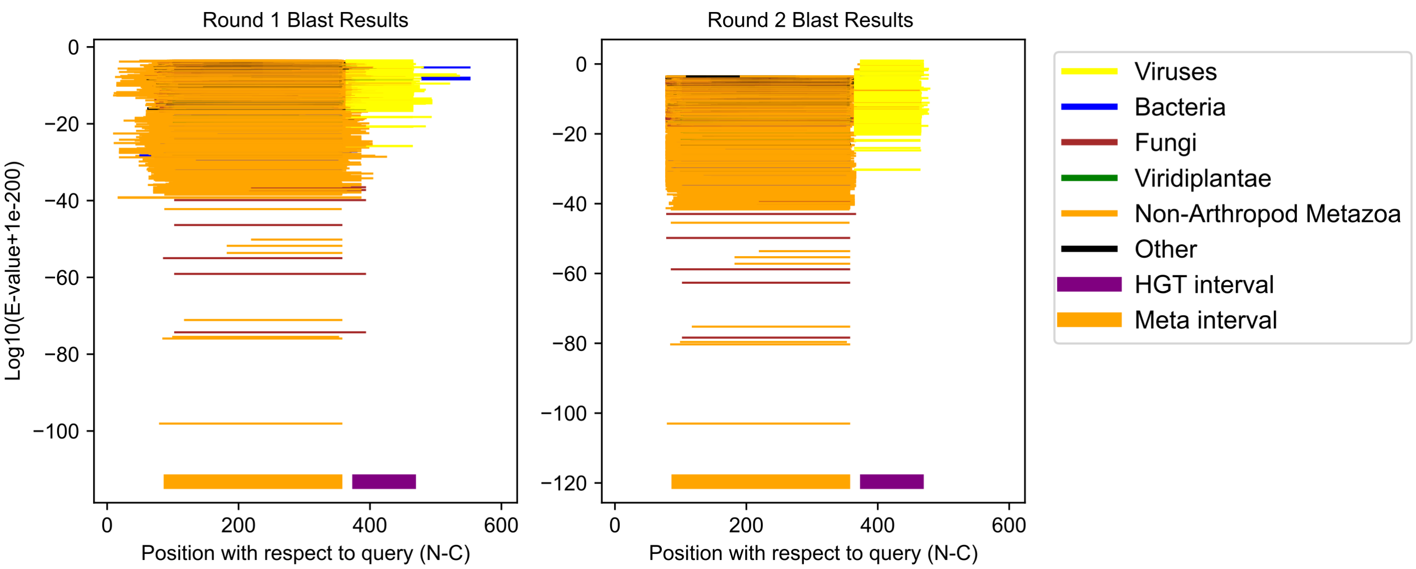

Ixodes scapularis GCF_016920785.2;XP_029821973.3 (cluster 89)

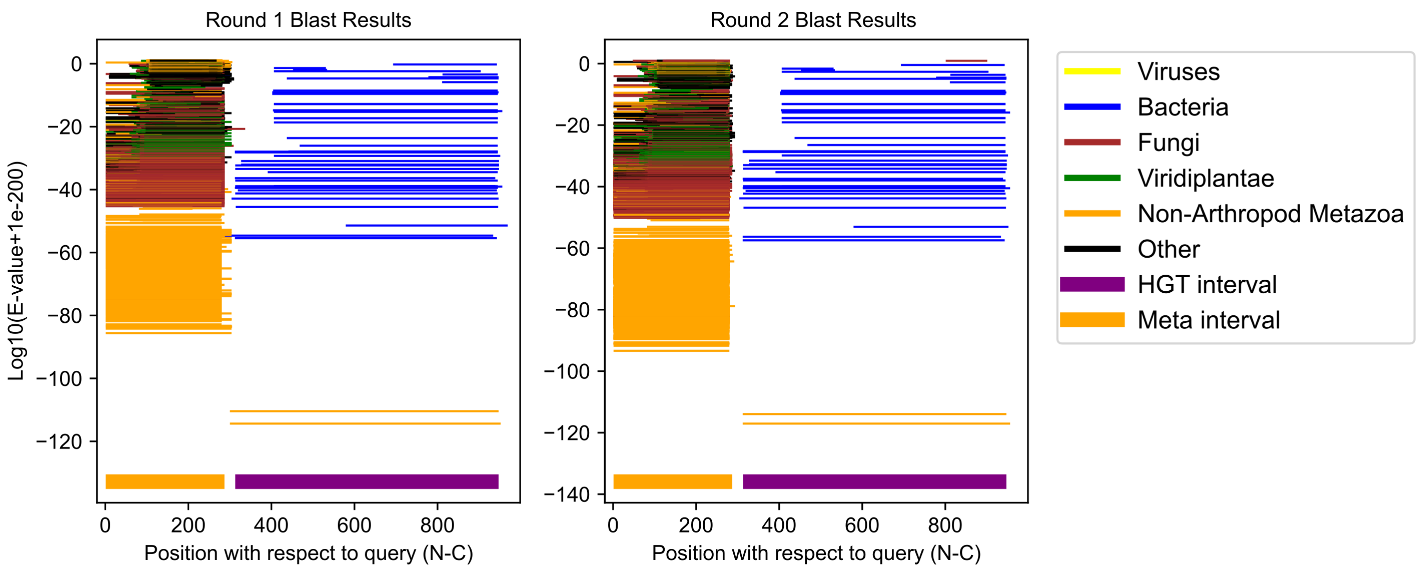

Culex pipiens pallens GCF_016801865.2;XP_052565726.1 (cluster 90)

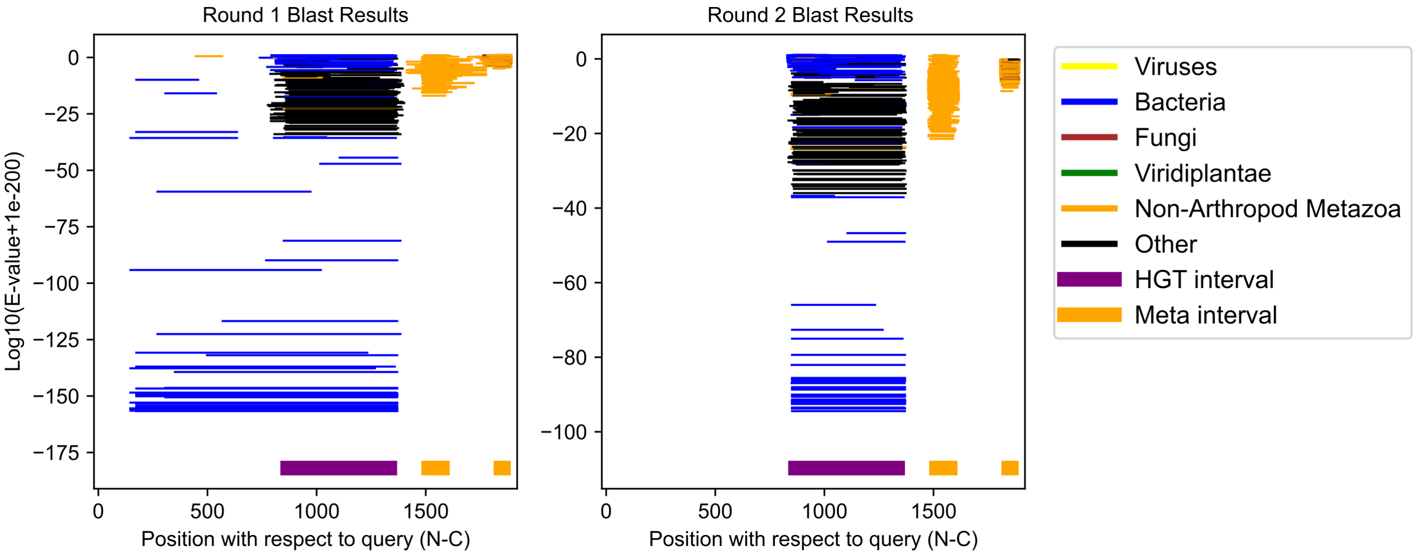

Lepeophtheirus salmonis GCF_016086655.4;XP_040568466.1 (cluster 91)

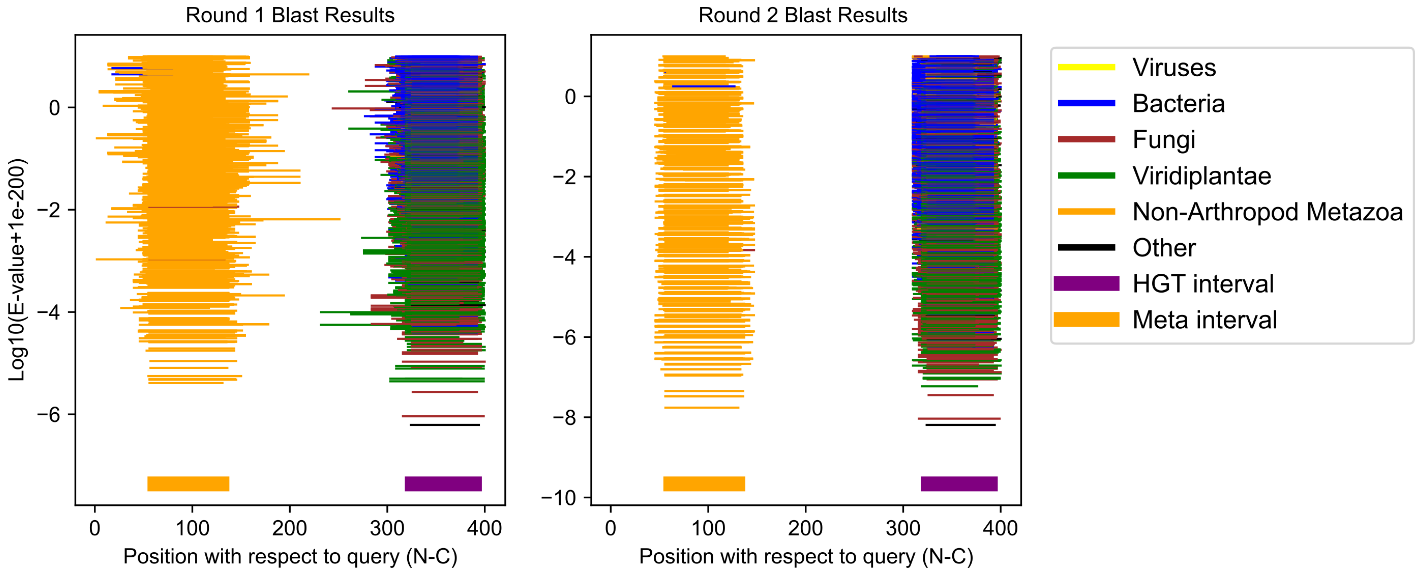

Manduca sexta GCF_014839805.1;XP_030032673.1 (cluster 92)

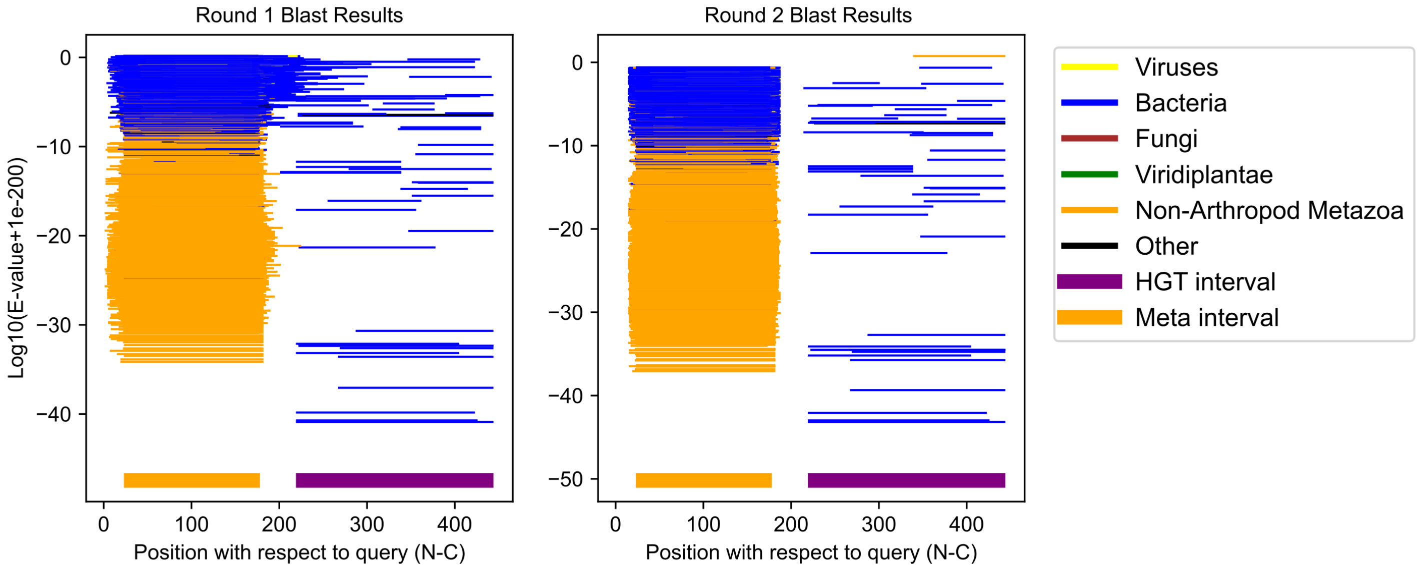

Bradysia coprophila GCF_014529535.1;XP_037049533.1 (cluster 93)

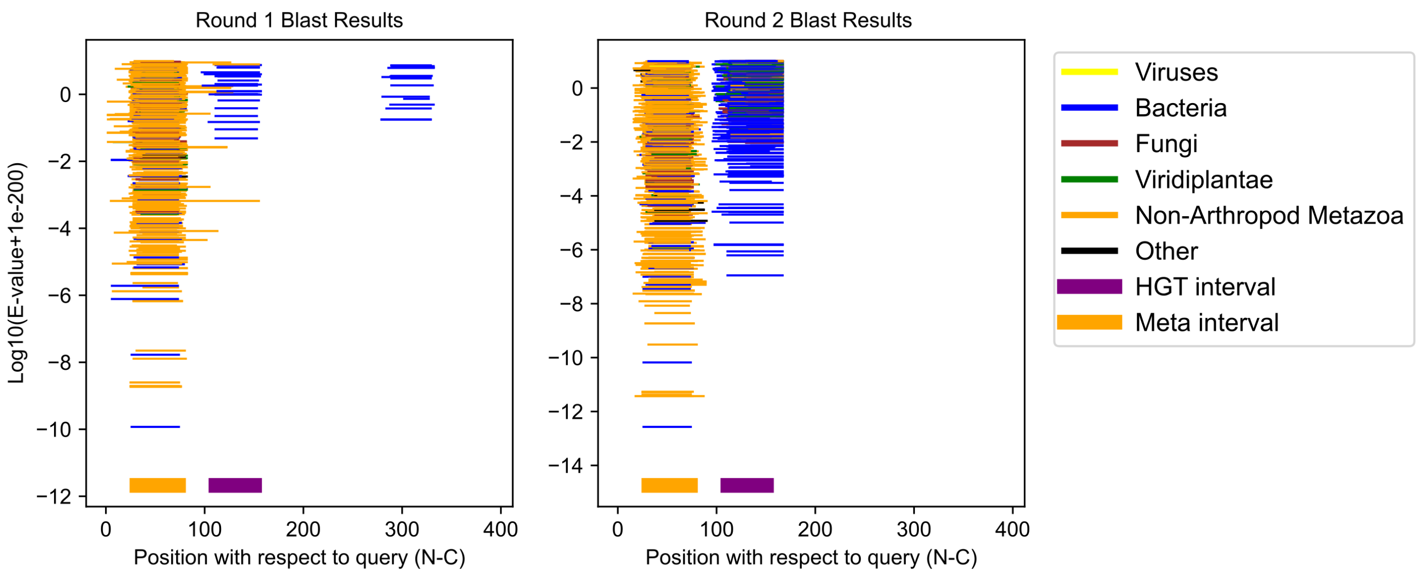

Bradysia coprophila GCF_014529535.1;XP_037030969.1 (cluster 94)

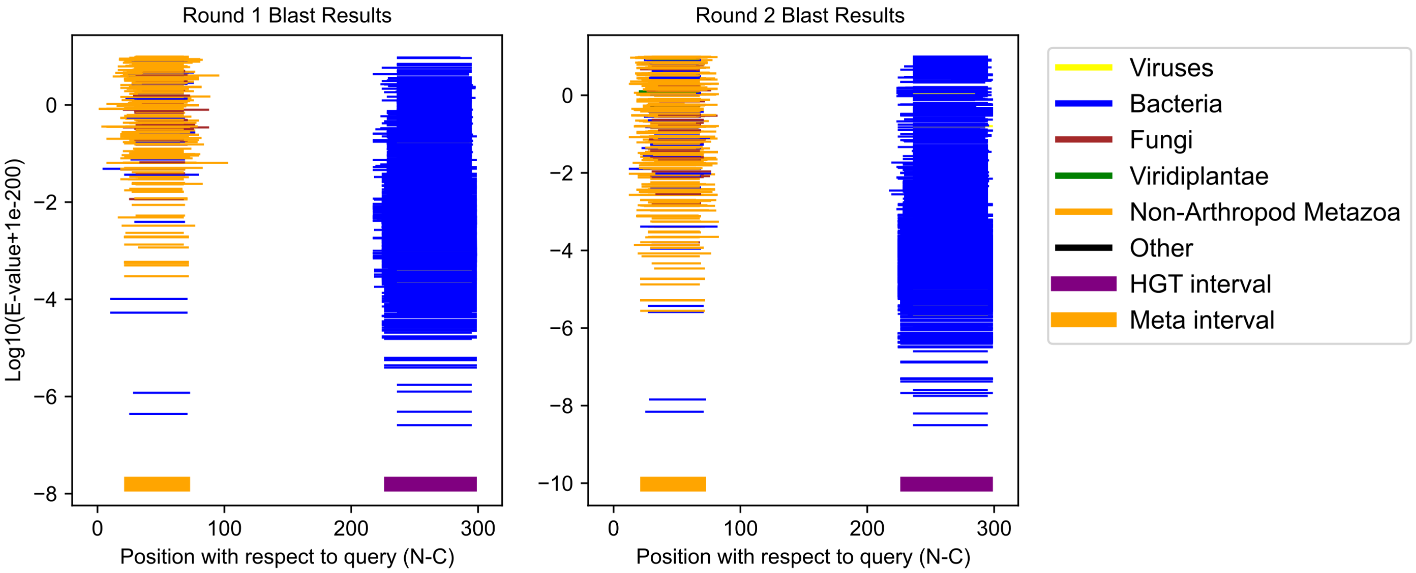

Tigriopus californicus GCF_007210705.1;XP_059099335.1 (cluster 95)

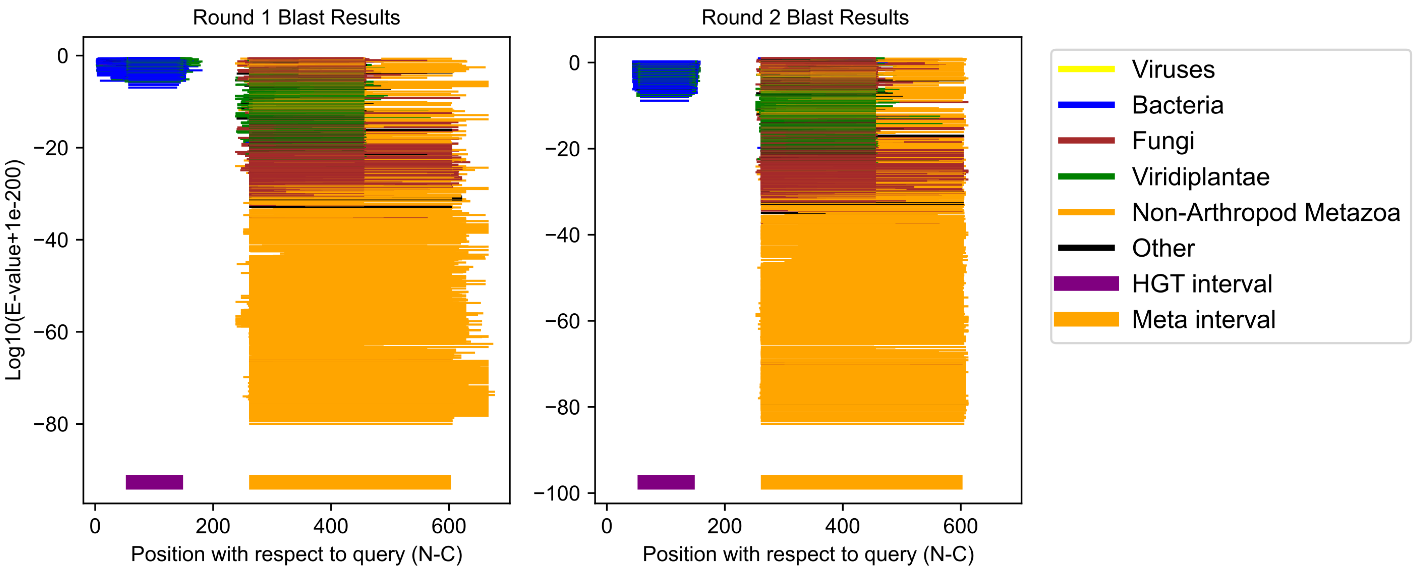

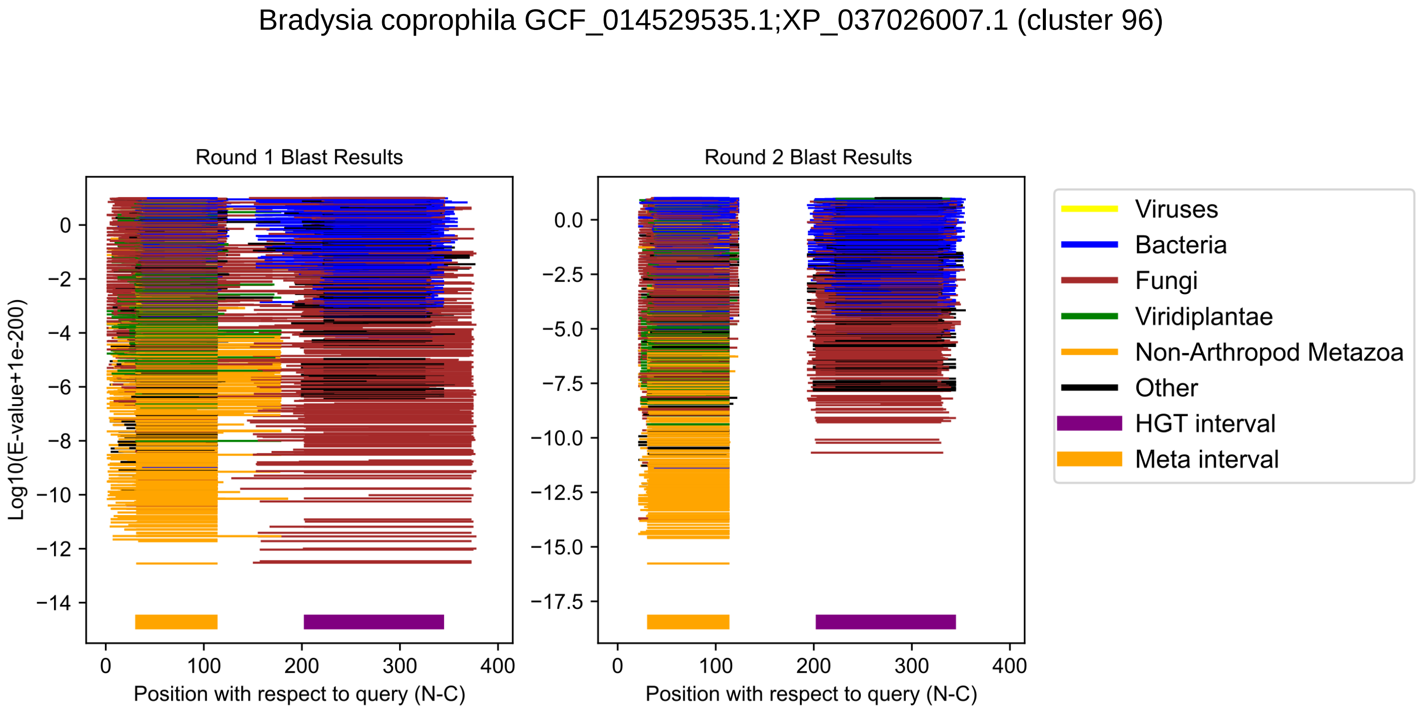
Bradysia coprophila GCF_014529535.1;XP_037026007.1 (cluster 96)
Round 1 Blast Results
Round 2 Blast Results
Log10(E-value+1e-200)
Position with respect to query (N-C)
Position with respect to query (N-C)
Viruses
Bacteria
Fungi
Viridiplantae
Non-Arthropod Metazoa
Other
HGT interval
Meta interval

Monomorium pharaonis GCF_013373865.1;XP_028045706.1 (cluster 97)

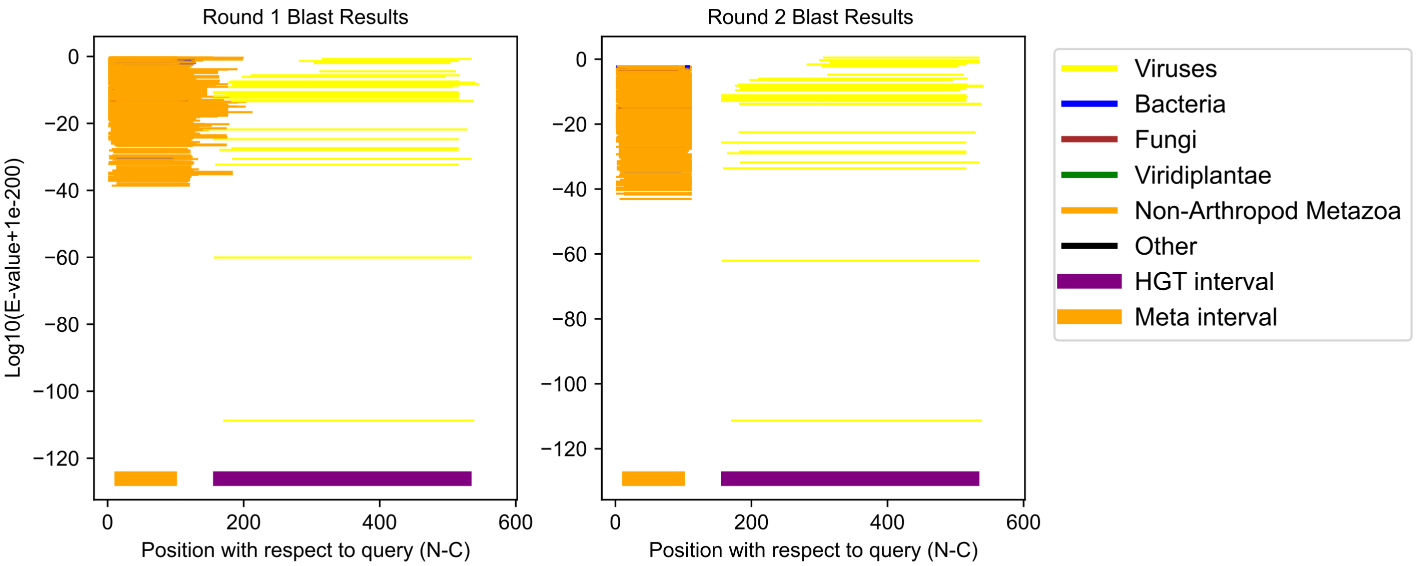

Chelonus insularis GCF_013357705.1;XP_034949802.1 (cluster 98)

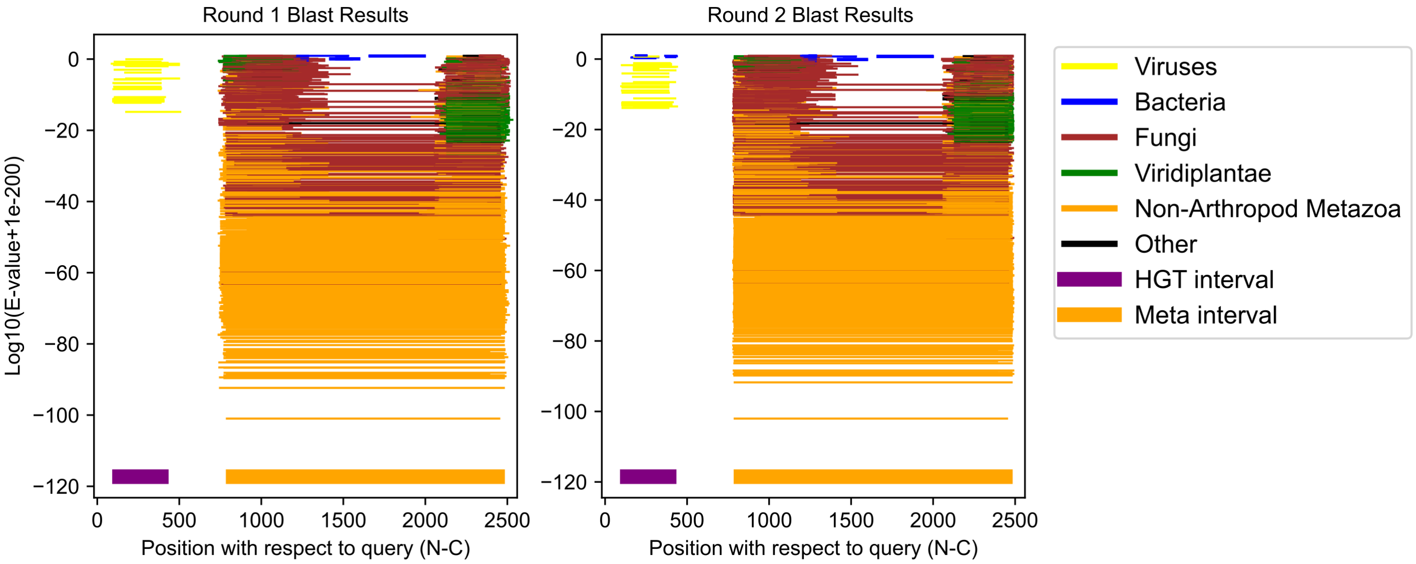

Rhipicephalus sanguineus GCF_013339695.2;XP_037517182.1 (cluster 99)

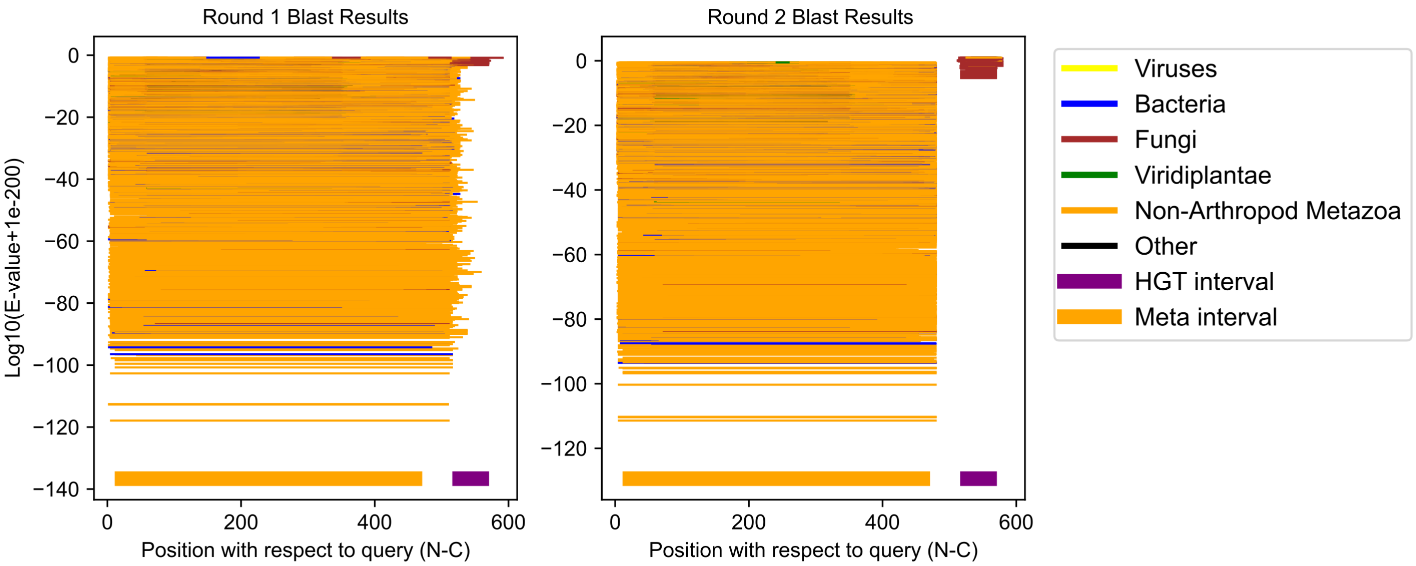

Thrips palmi GCF_012932325.1;XP_034245505.1 (cluster 100)

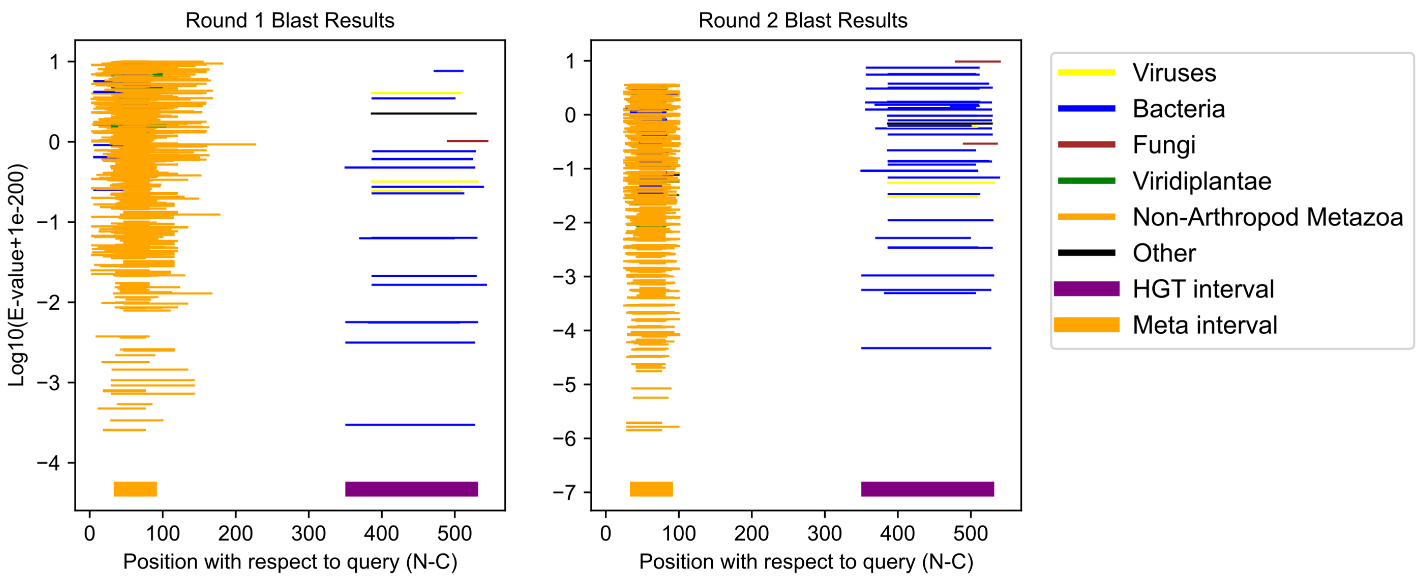

Contarinia nasturtii GCF_009176525.2;XP_031633908.1 (cluster 101)

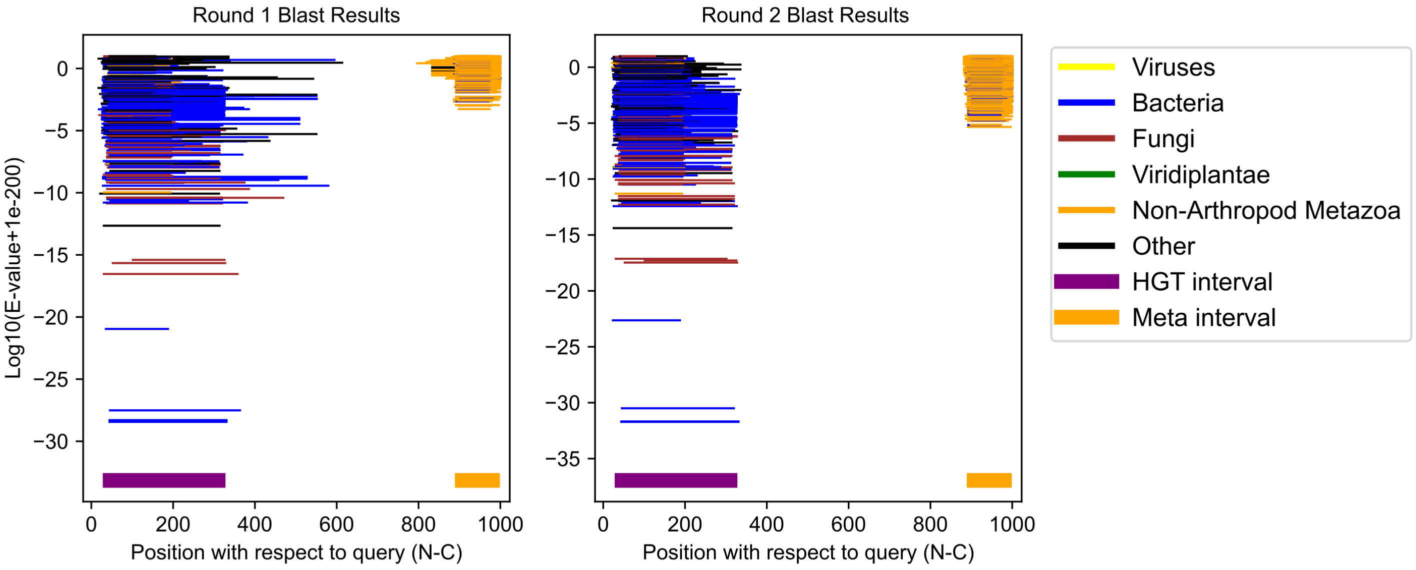

Sitodiplosis mosellana GCF_009176505.1;XP_055306475.1 (cluster 102)

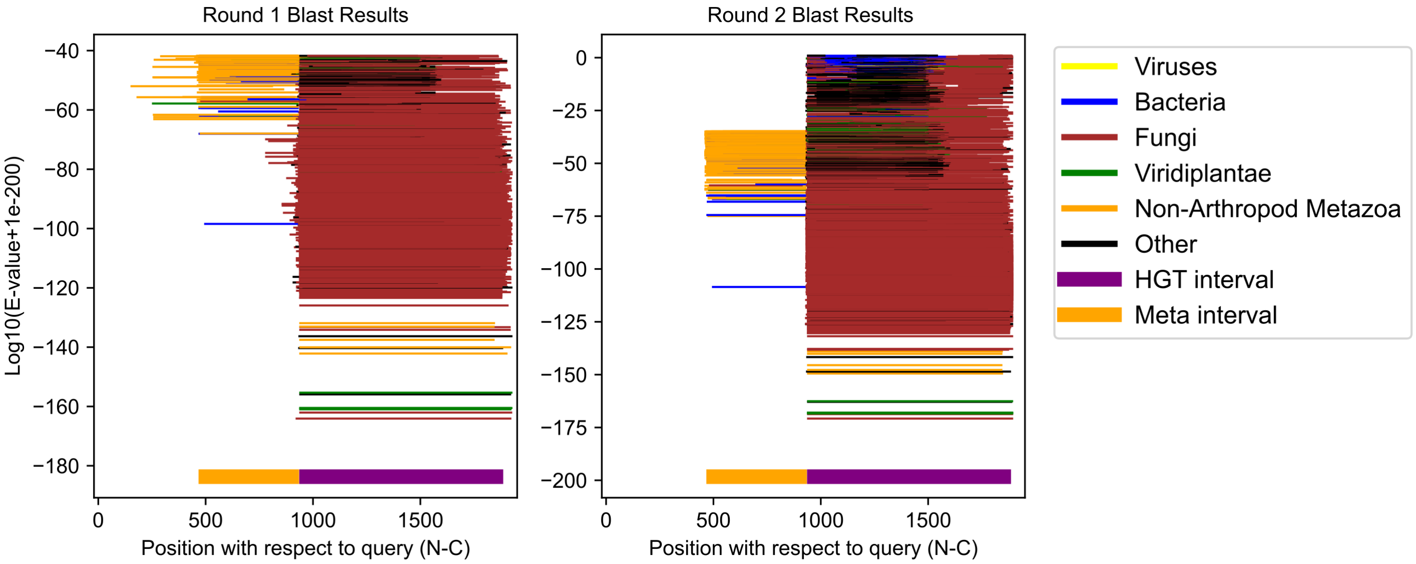

Photinus pyralis GCF_008802855.1;XP_031328047.1 (cluster 103)

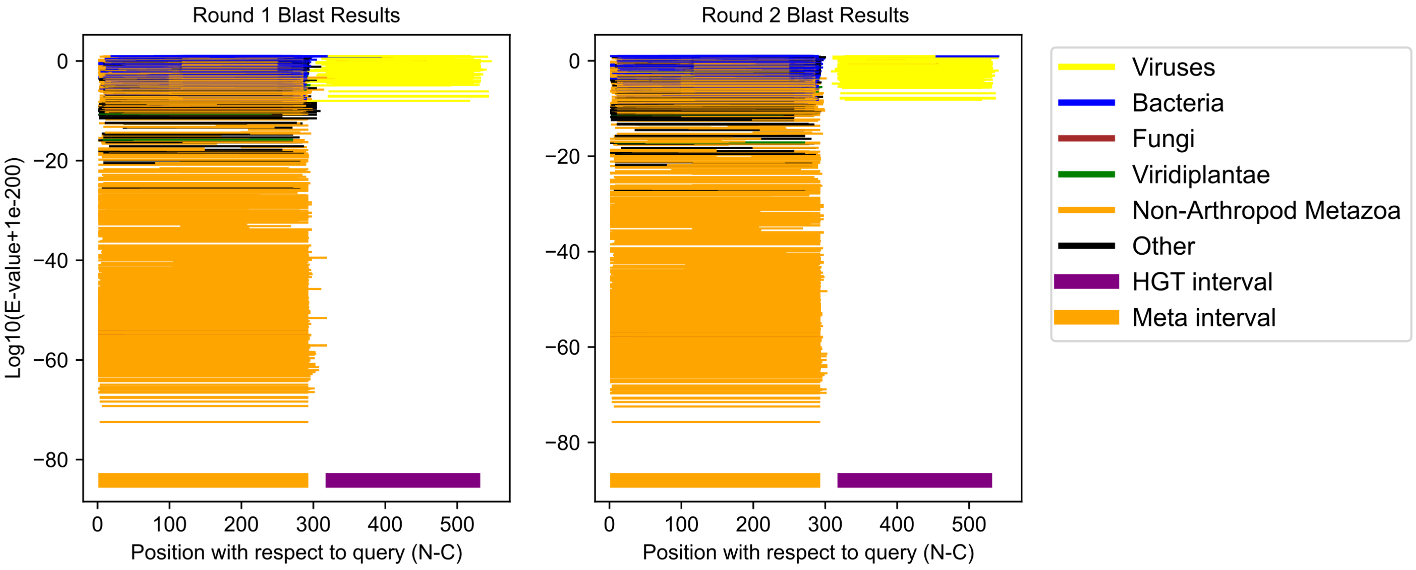

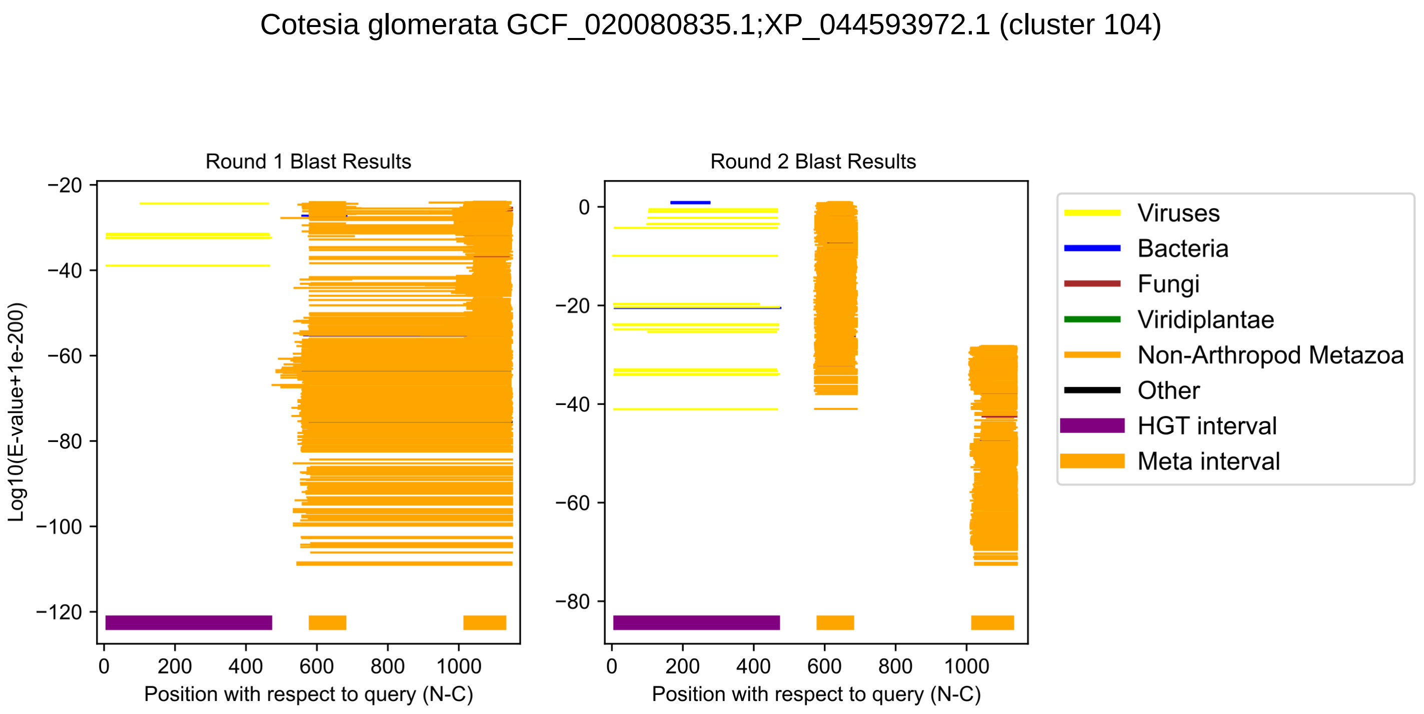
Cotesia glomerata GCF_020080835.1;XP_044593972.1 (cluster 104)
Round 1 Blast Results
Round 2 Blast Results
Log10(E-value+1e-200)
Position with respect to query (N-C)
Position with respect to query (N-C)
−20
−40
−60
−80
−100
−120
0
−20
−40
−60
−80
0
200
400
600
800
1000
Viruses
Bacteria
Fungi
Viridiplantae
Non-Arthropod Metazoa
Other
HGT interval
Meta interval
