## Supplementary material for "Evolutionary innovation through fusion of sequences from across the tree of life": SI Files 1 and 2: HGTc_SI_PDF_2_v3.pdf

# XP_042220148.1

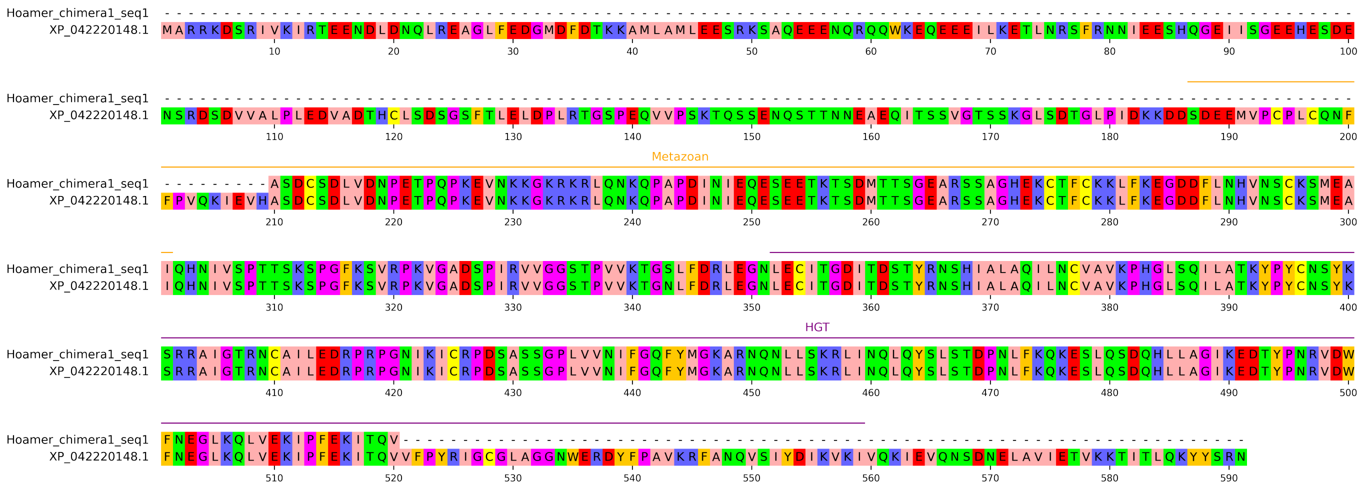

# XP_029735553.1

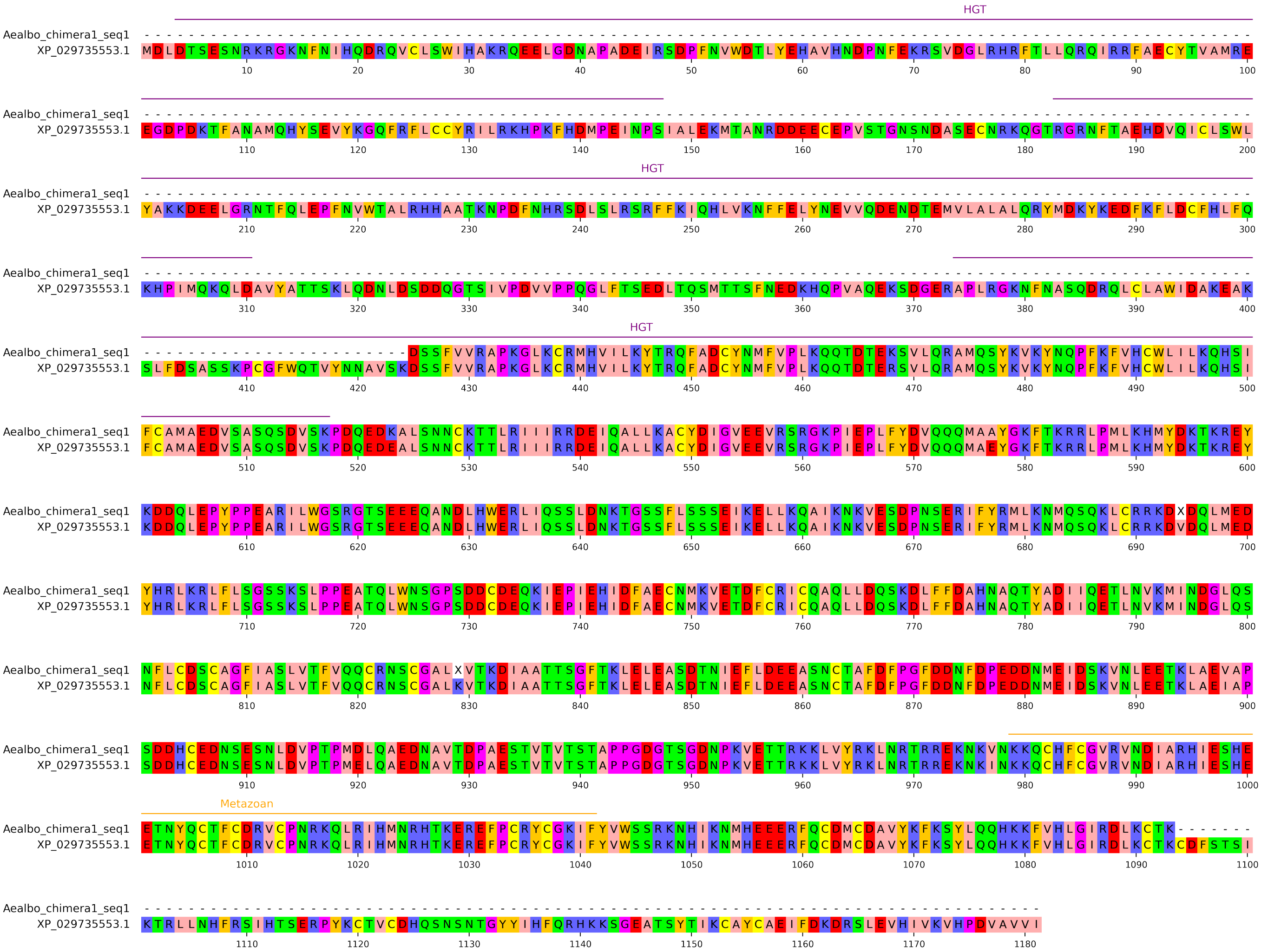

# XP_046453153.1

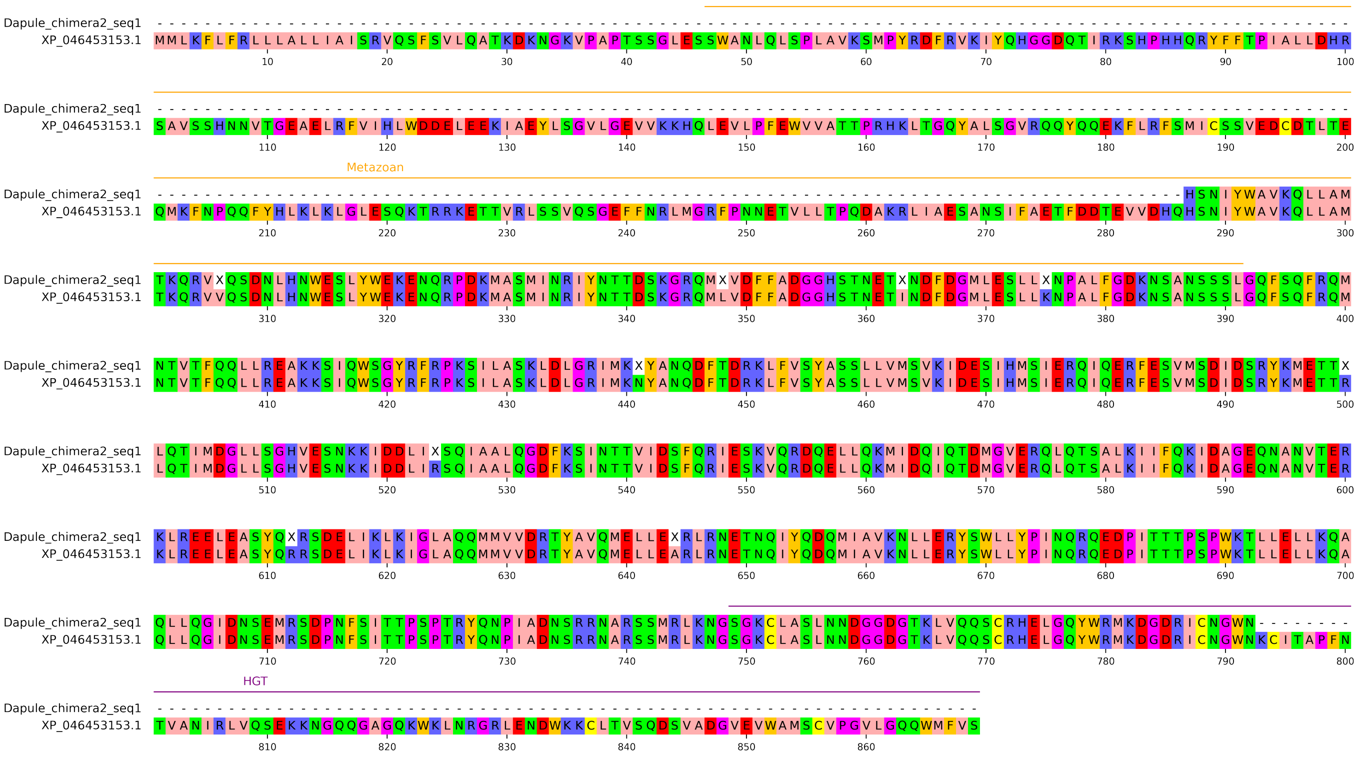

# XP_037051404.1

```
Brcopr_chimera1_seq1  ----------------------------------------------------------------------------------------VWIGAQRRSGNS
XP_037051404.1        MMRATVFCLFLIYFSITTVQGAERNSRCDTGWIPLEDEKCVRLFETFATREQAEDVCQTHGGTLVAVMSEEHQRVLTELVLNSSNAINVWIGAQRRSGNS
                               10        20        30        40        50        60        70        80        90       100

                                                   Metazoan
Brcopr_chimera1_seq1  SEFVWNDGSTVRRYTSWATGHPTNSIGRNCVQMRSELSRTQRSDMEWVDIACVIANWFICQKLQMWSPENVQQALLNLRRELRDSDYSVSNEITDMKRQL
XP_037051404.1        SEFVWNDGSTVRRYTSWATGHPTNSIGRNCVQMRSELSRTQRSDMEWVDIACVIANWFICQKLQMWSPENVQQALLNLRRELRDSDYSVSNEITDMKRQL
                               110       120       130       140       150       160       170       180       190       200

                                                   HGT
Brcopr_chimera1_seq1  EVAQTELTHLQANPVPIGFIYVQLSNEPLPSTLWPTVQWTQVTSQYAGLFFRAEGGSSAAFGQIQTEGSPRLTSVRGRIALSAATSSITVAPGVATPNLS
XP_037051404.1        EVAQTELTHLQANPVPIGFIYVQLSNEPLPSTLWPTVQWTQVTSQYAGLFFRAEGGSSAAFGQIQTEGSPRLTSVRGRIALSAATSSITVAPGVATPNLS
                               210       220       230       240       250       260       270       280       290       300

Brcopr_chimera1_seq1  VGATGATNHWGMSFTVSSAEVRPRNSAIRIWRRS-
XP_037051404.1        VGATGATNHWGMSFTVSSAEVRPRNSAIRIWRRSG
                               310       320       330
```

# XP_035715507.1

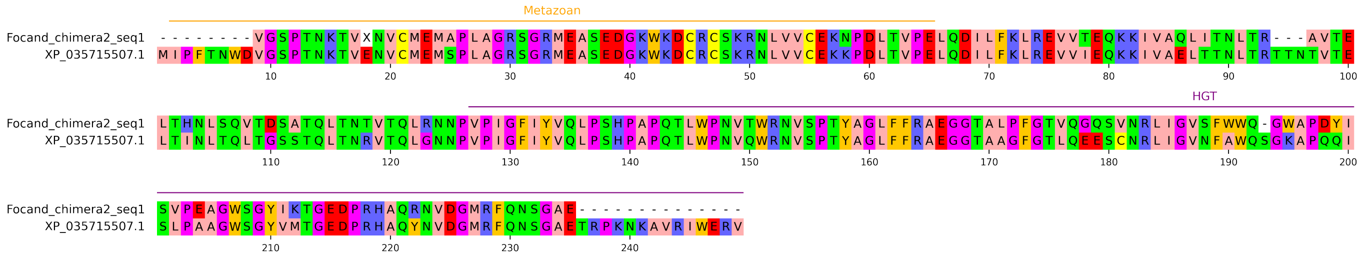

# XP_059086440.1

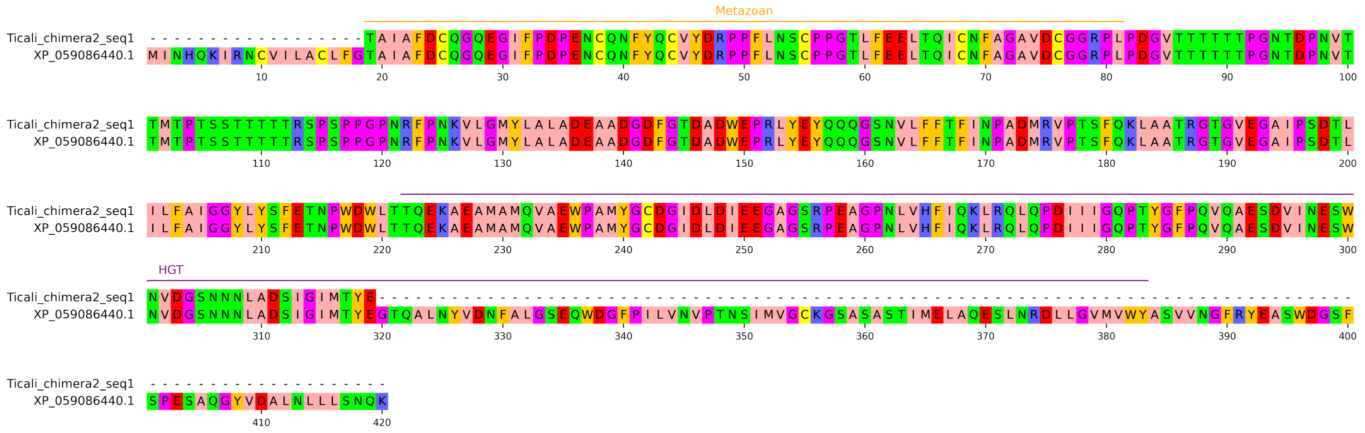

# XP_029821973.3

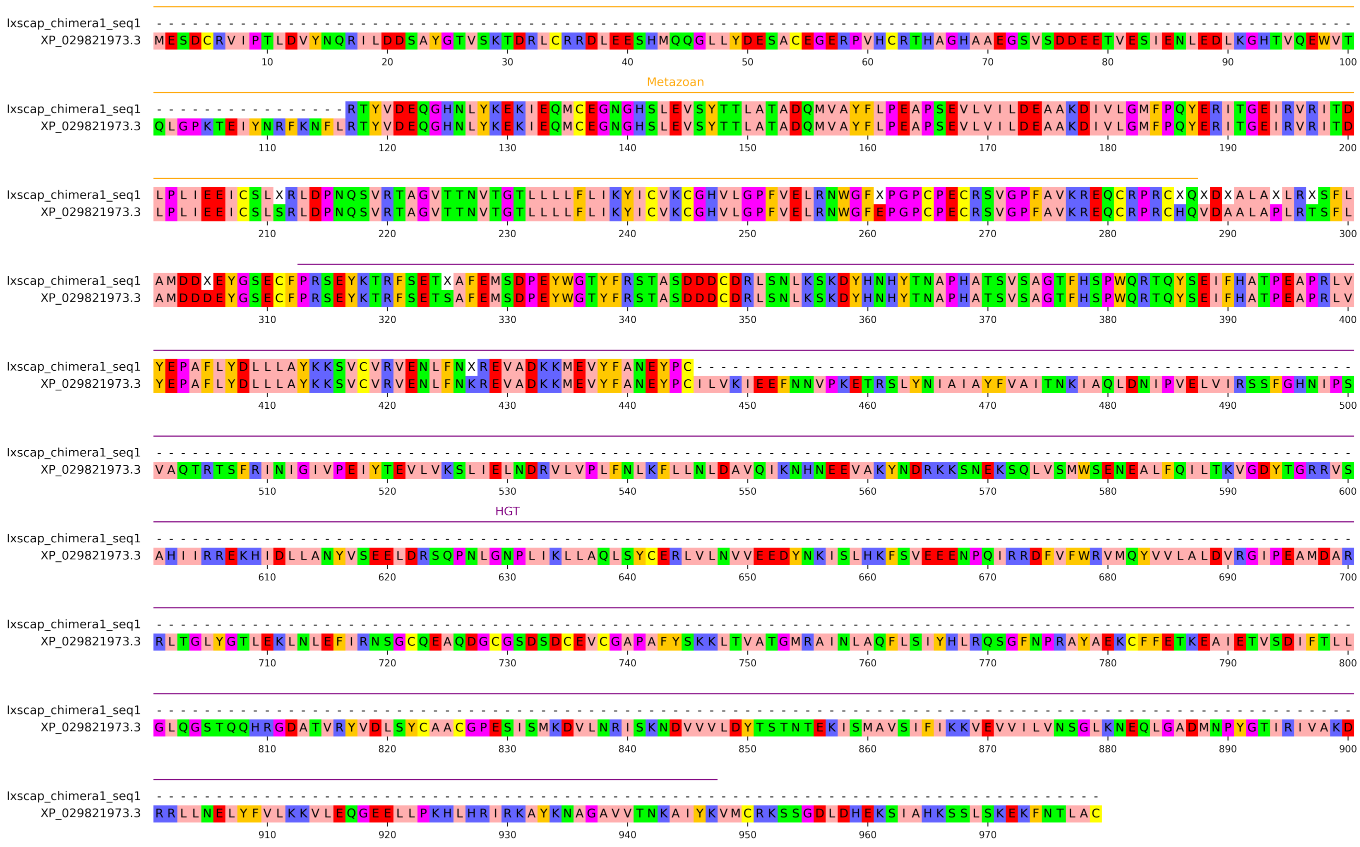

# XP_046402901.1

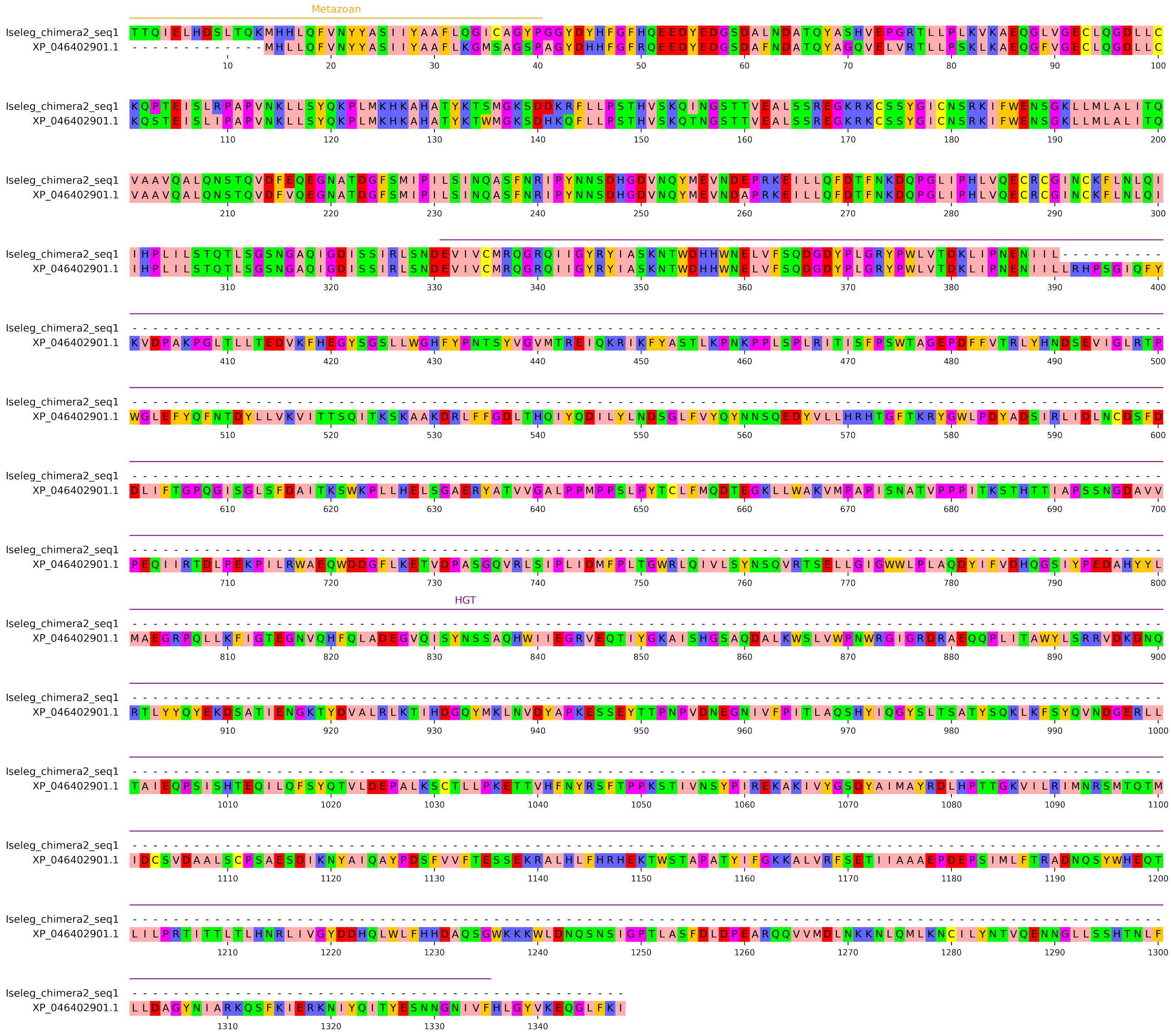

# XP_023346081.1

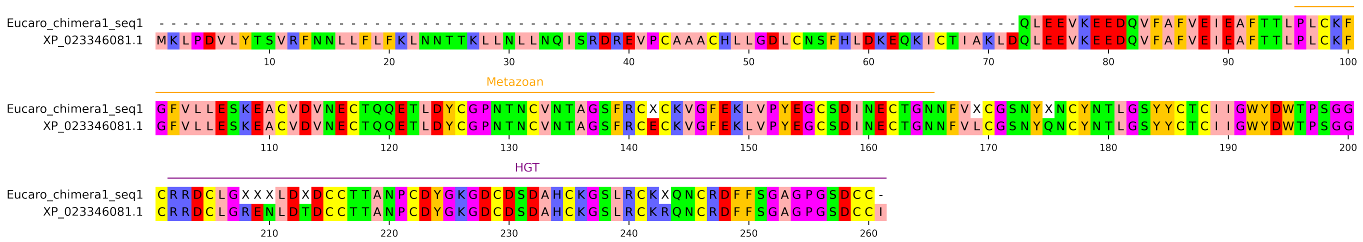

# XP_035708168.1

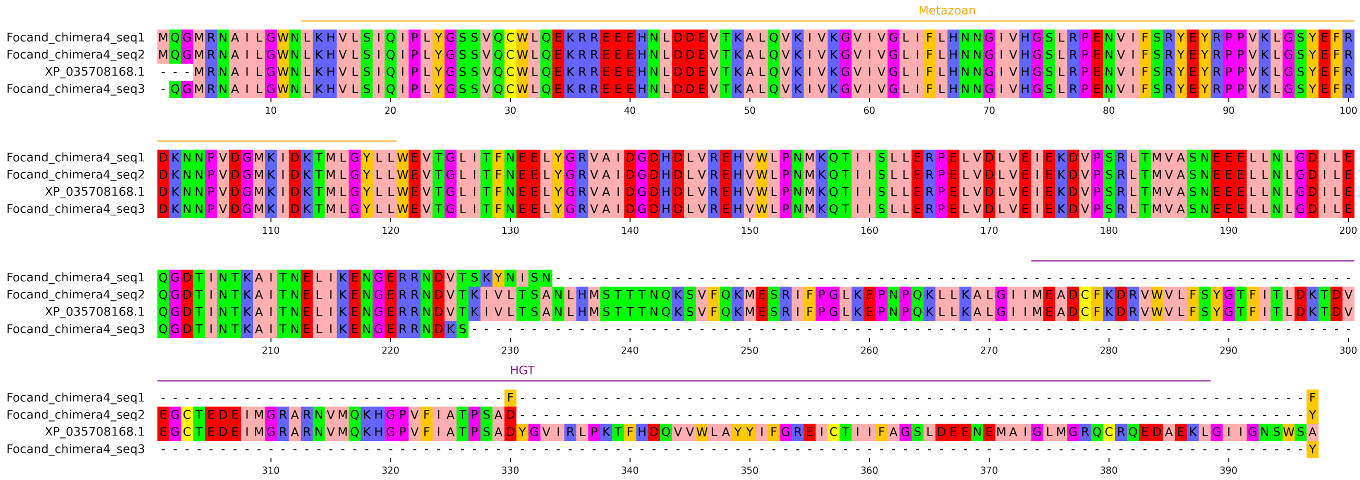

# XP_037026007.1

```
                                                                     Metazoan
Brcopr_chimera6_seq1  ---------------------------------------TAIDNGRRVAAKDSNAQTGTVGHYTDTGSRYEDASNNGVAHISENMASQGRSKRSPTDLKG
XP_037026007.1        MAYRLVKLLTNLRFSSEASNAIQRFNSCHSLVNVPATQCTAIDNGRRVAAKDSNAQTGTVGHYTDTGSRYEDASNNGVAHISENMASQGRSKRSPTDLKG
                               10        20        30        40        50        60        70        80        90       100

Brcopr_chimera6_seq1  DVVLNASYTSRDPTFRTSTFHRAYCNSKSSVPSFINMPIQNNSPKLNSISRHMSSYTSLTSVKIEQLQEIDGVNIDDVIRNEIENINEMIKVRELRMSVI
XP_037026007.1        DVVLNASYTSRDPTFRTSTFHRAYCNSKSSVPSFINMPIQNNSPKLNSISRHMSSYTSLTSVKIEQLQEIDGVNIDDVIRNEIENINEMIKVRELRMSVI
                              110       120       130       140       150       160       170       180       190       200

                                                                          HGT
Brcopr_chimera6_seq1  NEIYESLCKLISLLGVPTINTKESDKNFFVGSVLFSIAAFIRKEYQYDLSVQSEAPLKDFNIHMRGRMDFVVTDGKRKLVVIECKRHNADFGLVQCILAL
XP_037026007.1        NEIYESLCKLISLLGVPTINTKESDKNFFVGSVLFSIAAFIRKEYQYDLSVQSEAPLKDFNIHMRGRMDFVVTDGKRKLVVIECKRHNADFGLVQCILAL
                              210       220       230       240       250       260       270       280       290       300

Brcopr_chimera6_seq1  KHIYDTNNDGGDTFGFTTNGLEWRFVKYNGDSEPKVKLFEKQVLITTYMHLPV------------------------------------------
XP_037026007.1        KHIYDTNNDGGDTFGFTTNGLEWRFVKYNGDSEPKVKLFEKQVLITTYMHLPVRKERWMEHNTLILHIIFAAILSALNPTSIKLTEDTVTVTSSP
                              310       320       330       340       350       360       370       380       390
```

# XP_069990332.1

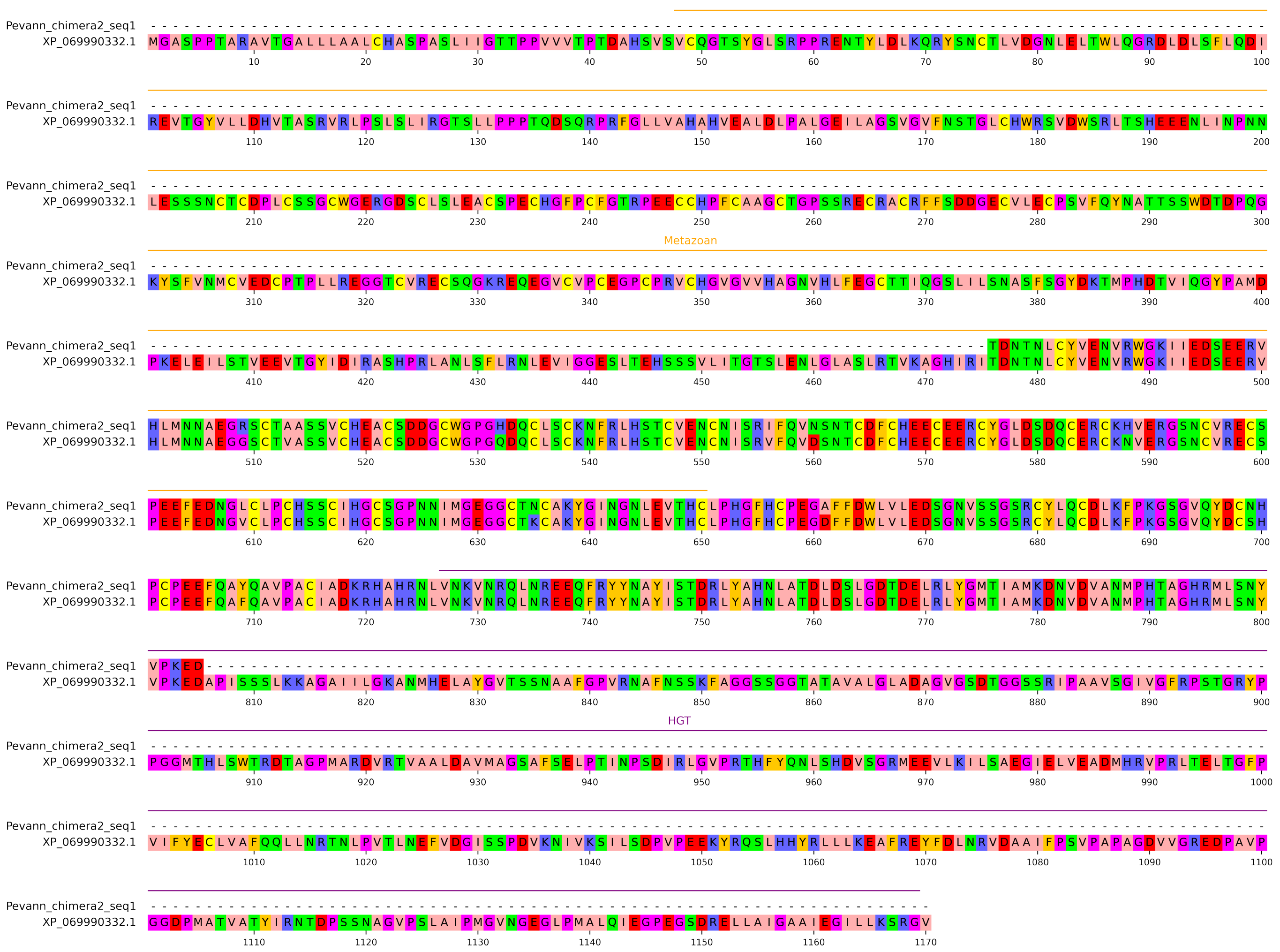

# XP_011211954.1

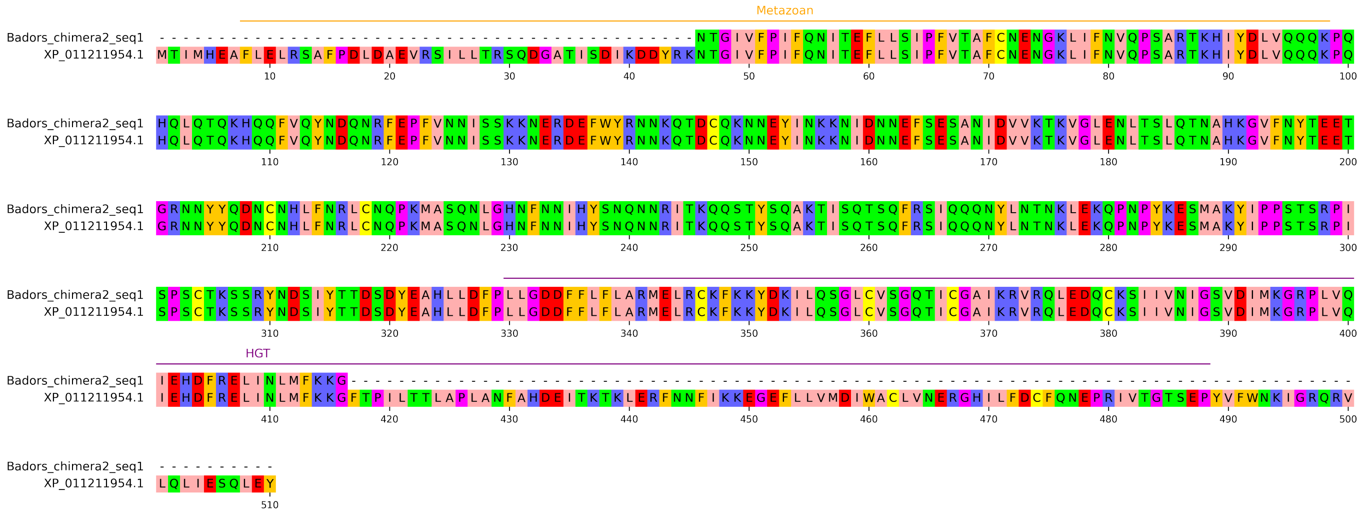

# XP_037038002.1

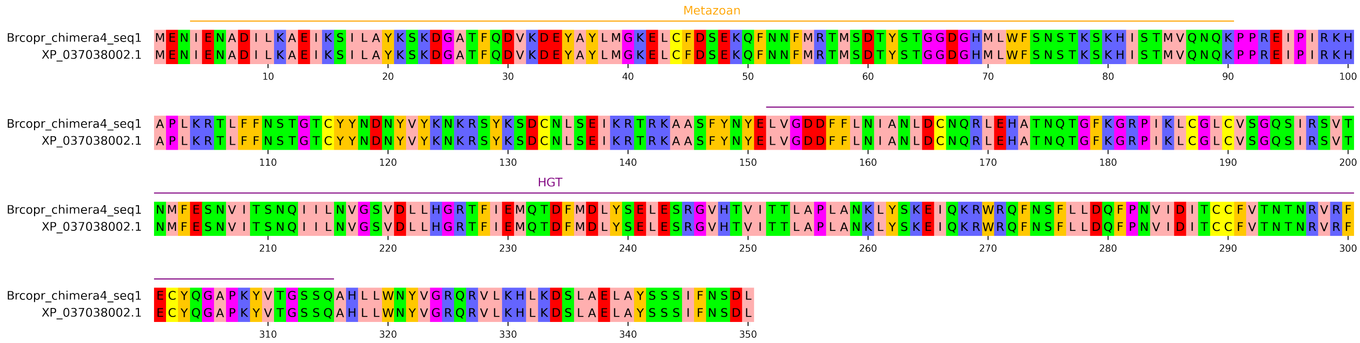

# XP_037049533.1

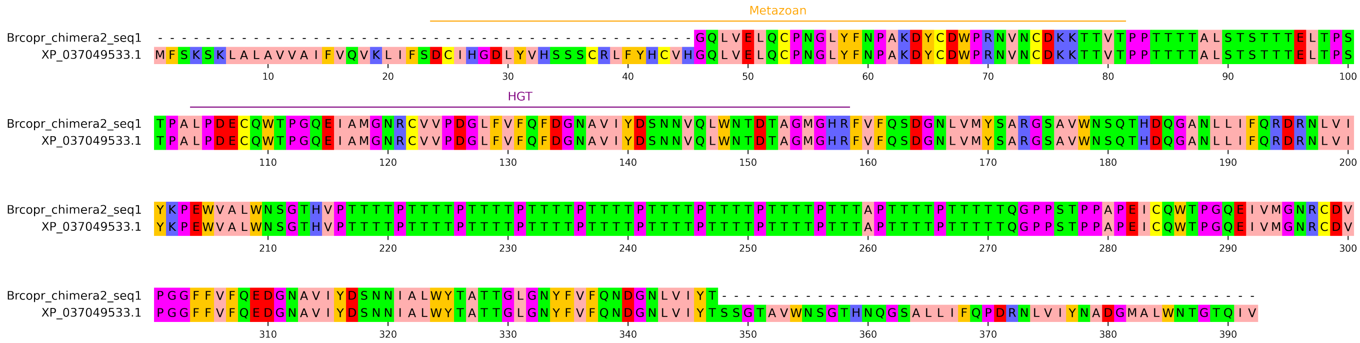

# XP_021699539.1

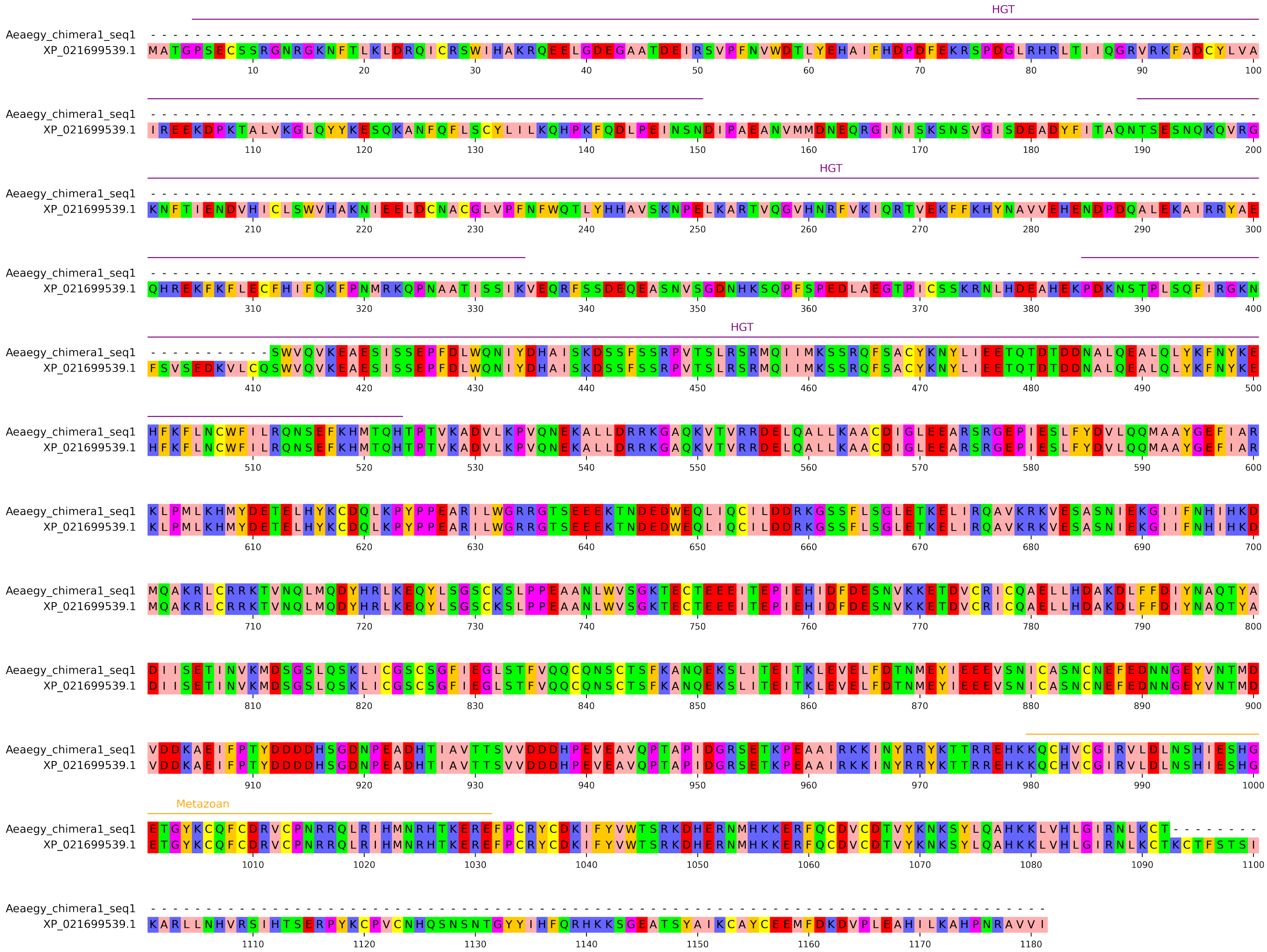

# XP_045027829.1

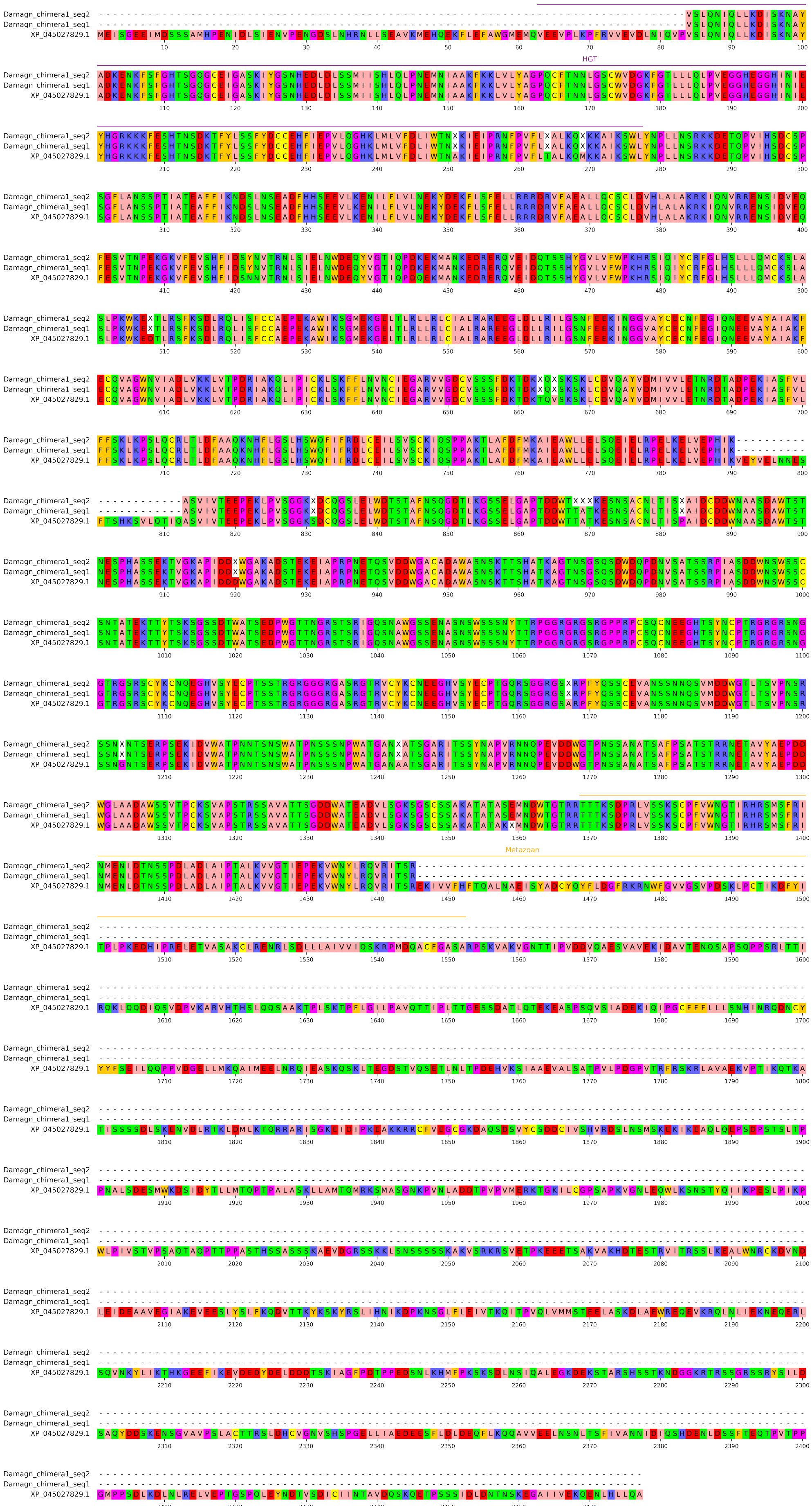

# XP_046445252.1

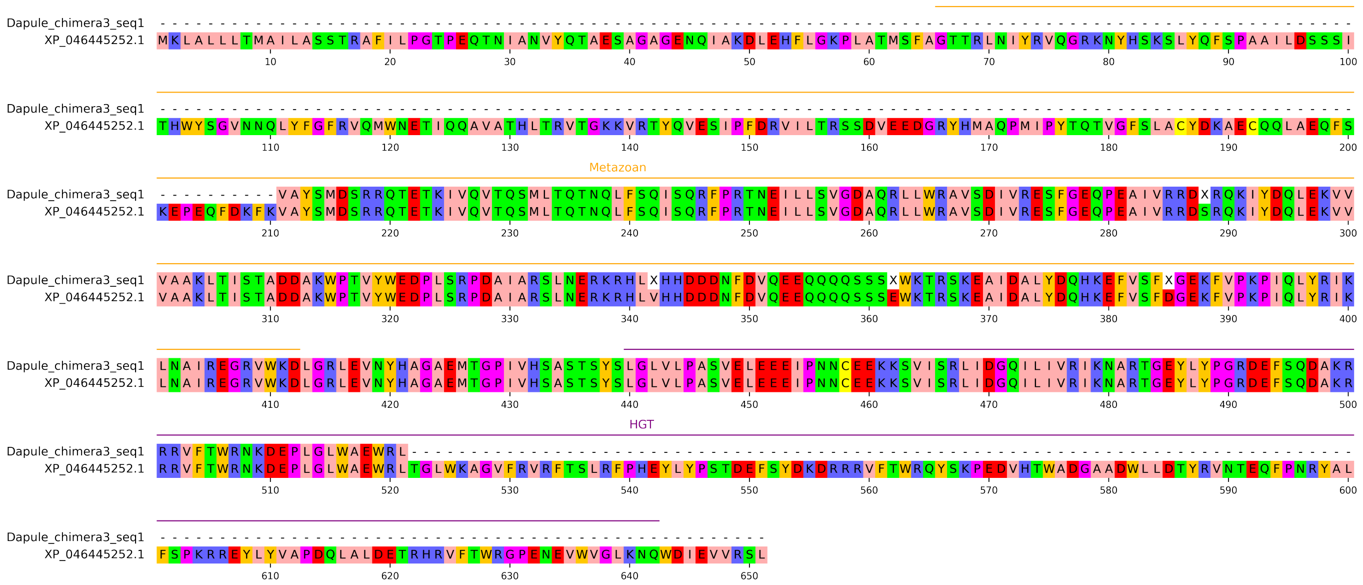

# XP_046456339.1

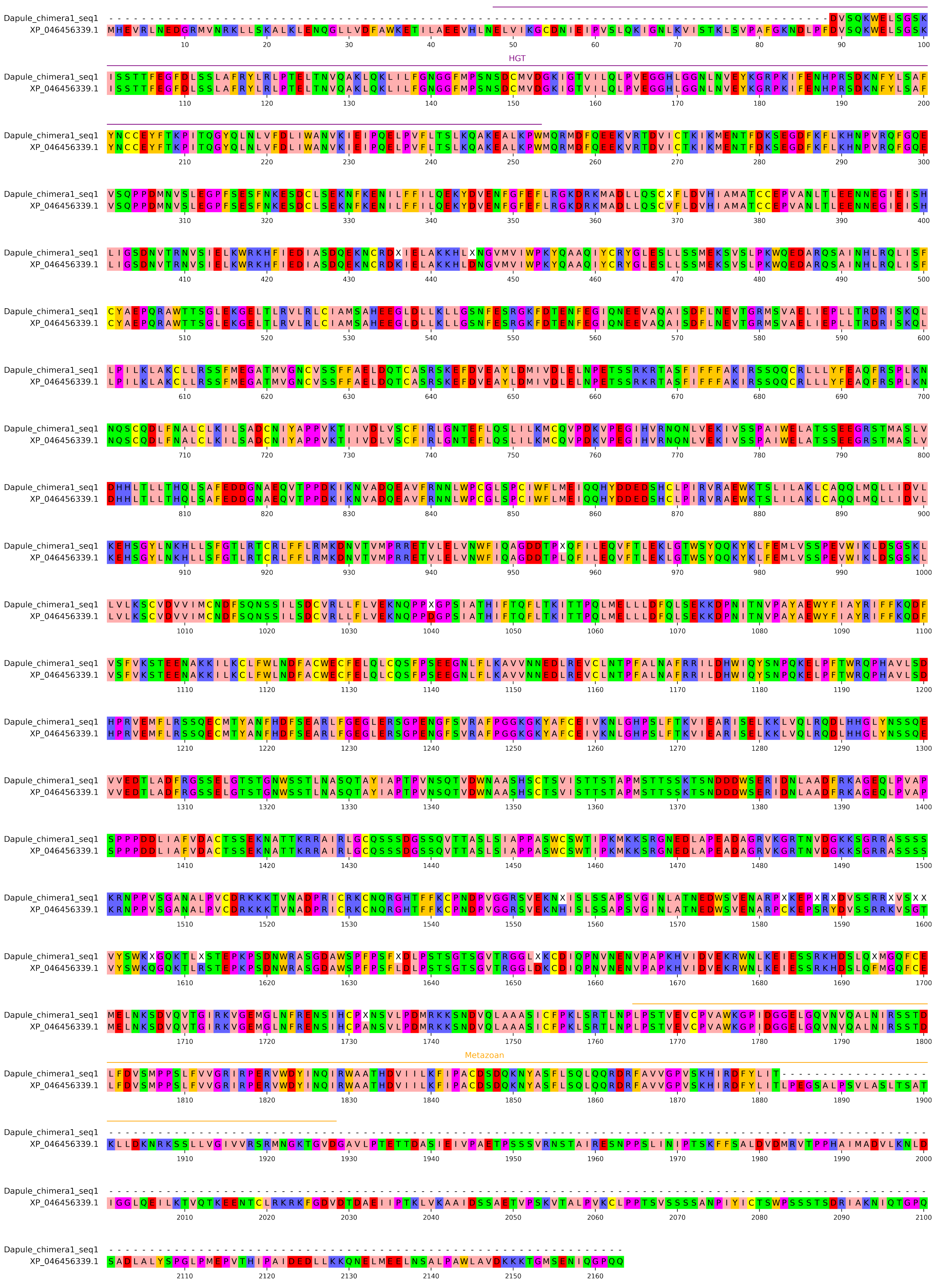

# XP_001656415.1

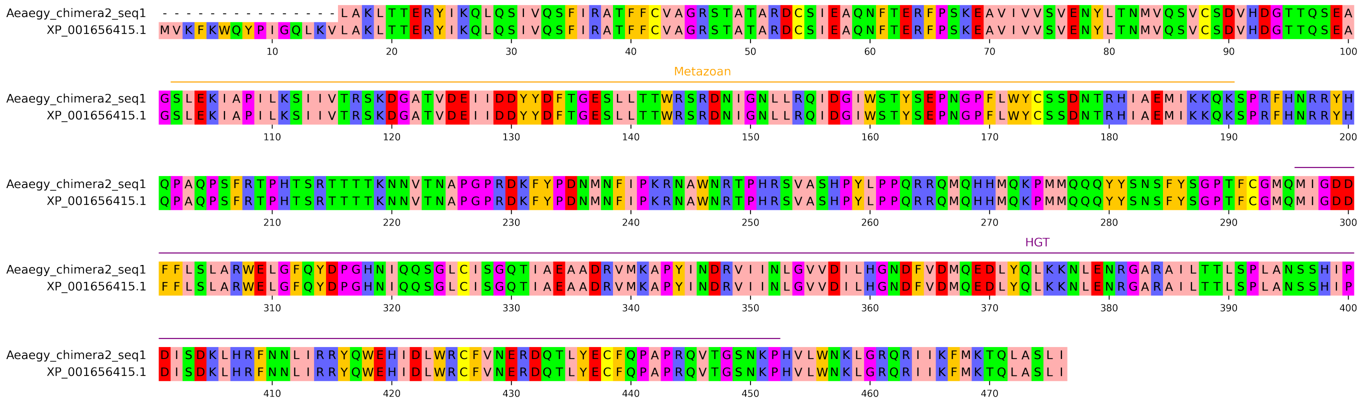

# XP_023329593.1

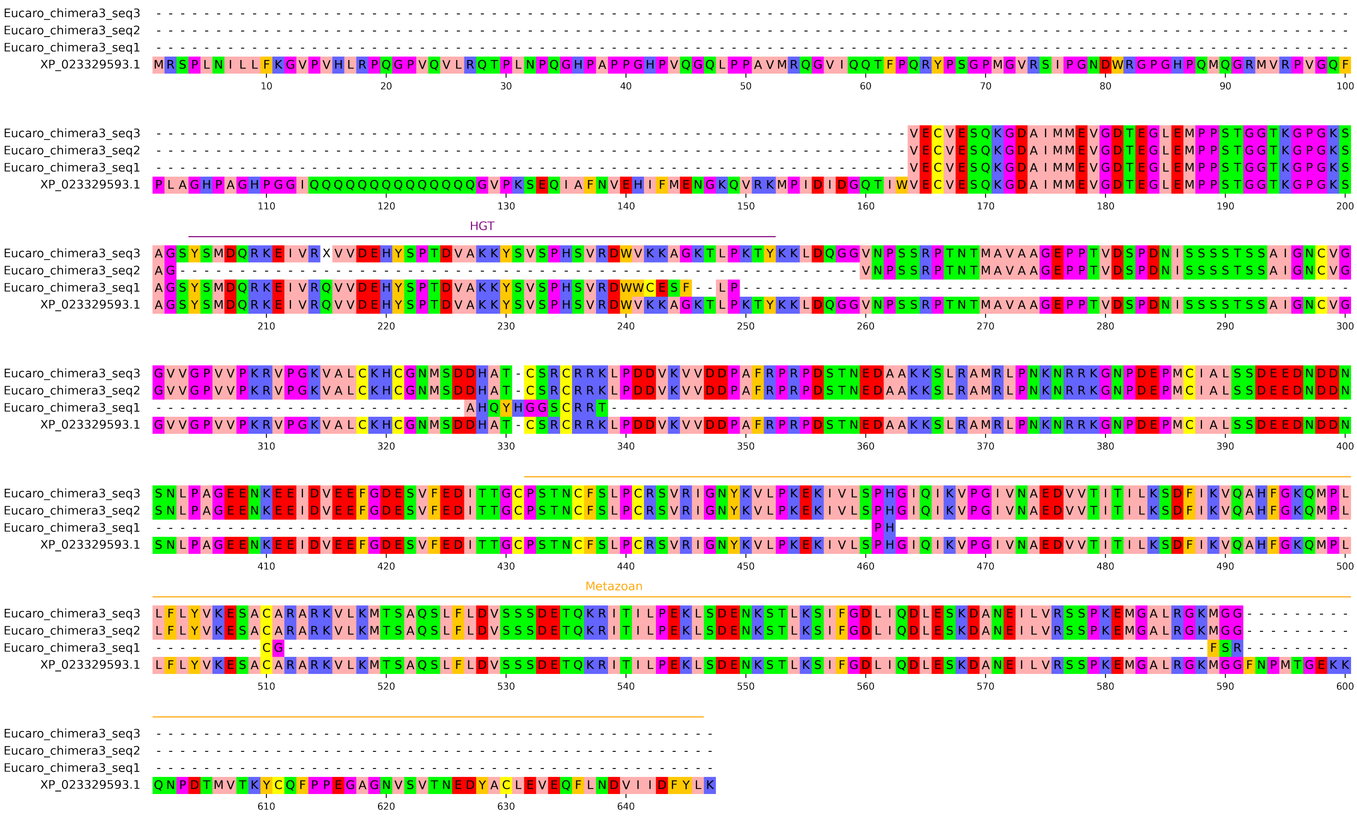

# XP_027232647.2

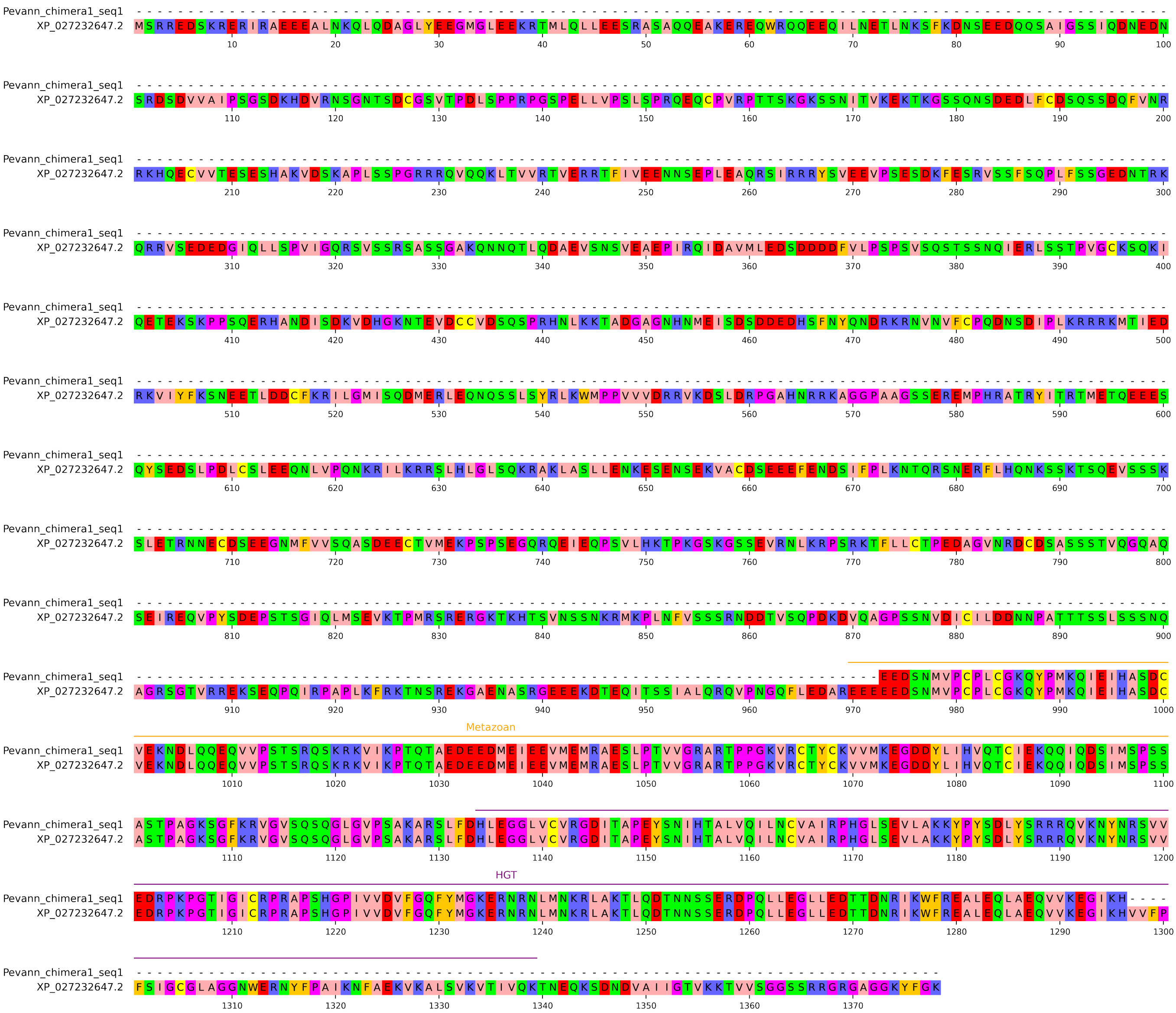

# XP_046646423.1

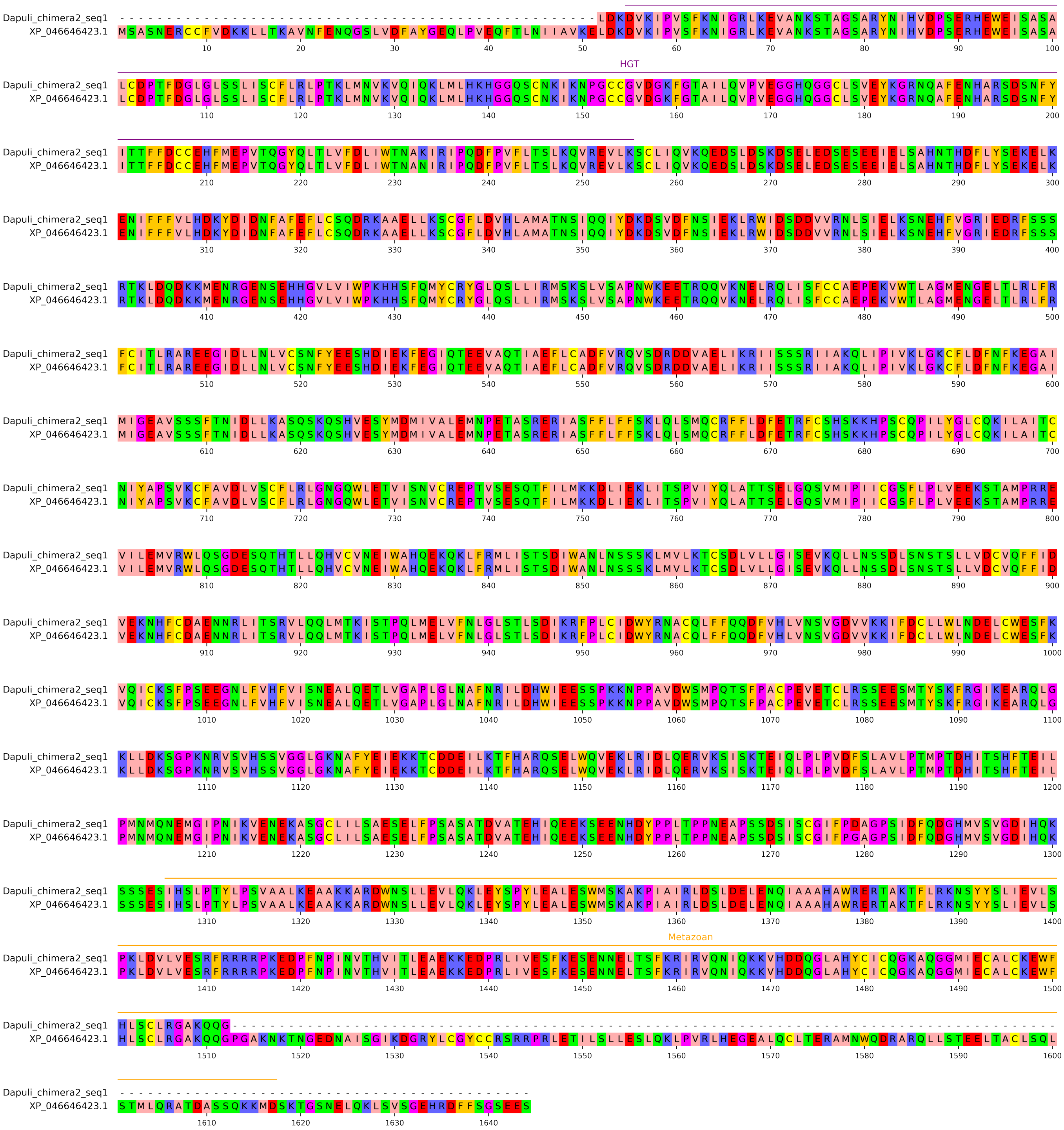

# XP_023343432.1

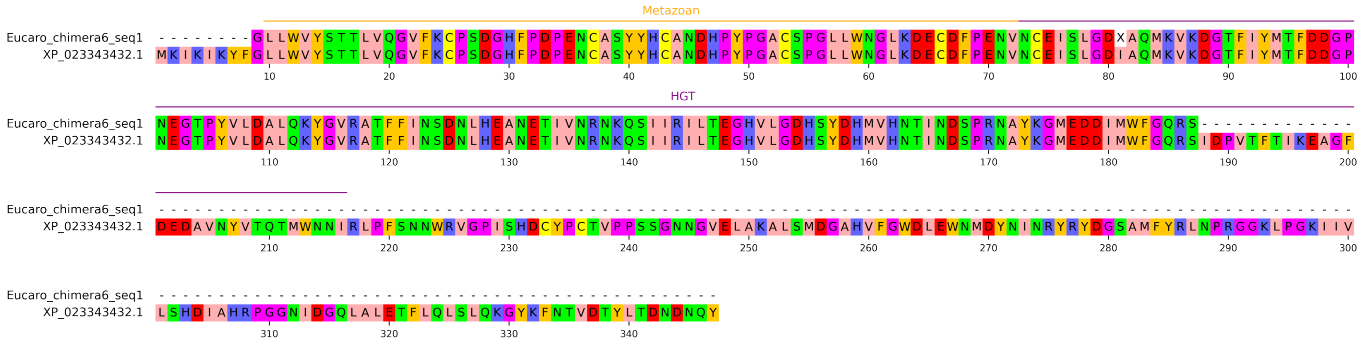

# XP_046403459.1

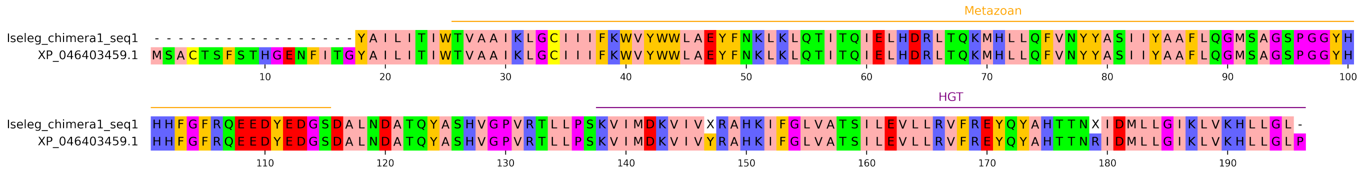

# XP_015919340.2

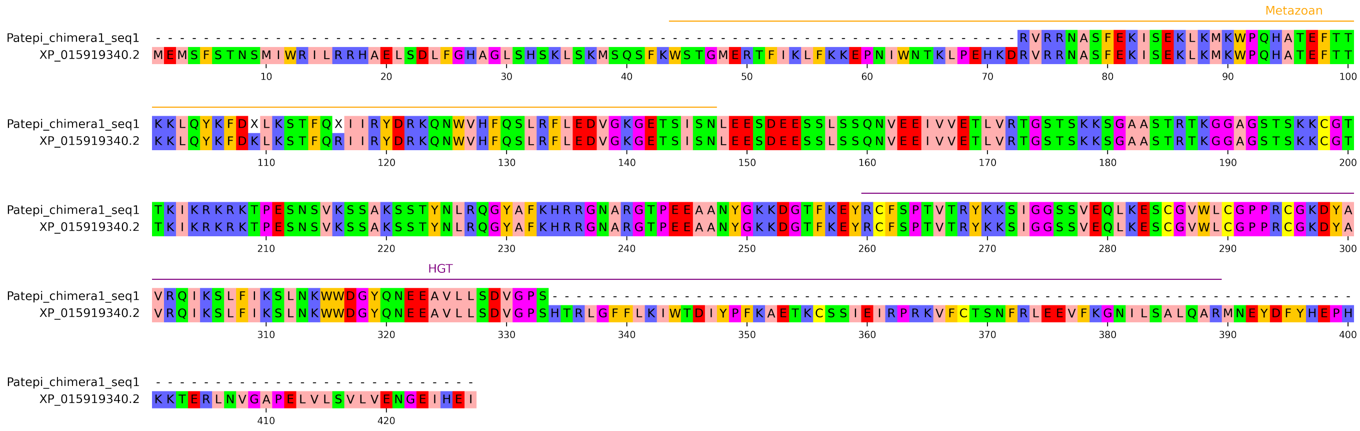

# XP_023324156.1

```
                                        Metazoan
Eucaro_chimera5_seq1  ----------------------------------------------------------------------------------------------------
XP_023324156.1        MRVGGGISLLFALFQSGVGQSVSCNHDGELLPDPENCRKYYKCAQGAPSPQSCPGTLVFDPLLKICNWPDSTTCIVDLKPKIPIREKSVISGLTEEDLFK
                               10        20        30        40        50        60        70        80        90       100

Eucaro_chimera5_seq1  ----------------------------------------------------------------------------------------------------
XP_023324156.1        GAEGIEEVPELTEAEEKRRKEAIASLFHVRDRKTNPLVRLRTSRPRTSQEPRPTARPTEVPQFSNRGSQLFRNSGDADRDRDRNSEQDENARDRDRDSFR
                              110       120       130       140       150       160       170       180       190       200

                                                                                                    Metazoan
Eucaro_chimera5_seq1  ------------------------------------------------------------------------------CESGGPSLQSCPANLIFDTKL
XP_023324156.1        DSDRDSENREEDQEKFRKQKVKEQILRDLFAESEFQESAPVEEDKPQEKKKPYFEPIHCHSGASNELYANPQNCRKYYKCESGGPSLQSCPANLIFDTKL
                              210       220       230       240       250       260       270       280       290       300

Eucaro_chimera5_seq1  KICNWPDAAECLEDPSFELPDIVSSDTHPPKQRNPSINTIDTGTFIEPEPIVISSVPSPEHANVPSFREAKERENLLTKQYPLFDKVKKTVHTLDTSIVE
XP_023324156.1        KICNWPDAAECLEDPSFELPDIVSSDTHPPKQRNPSINTIDTGTFIEPEPIVISSVPSPEHANVPSFREAKERENLLTKQYPLFDKVKKTVHTLDTSIVE
                              310       320       330       340       350       360       370       380       390       400

Eucaro_chimera5_seq1  KIKPGASDNPDNVQRVEFILTEANYEDLFPRRHPSYTYQRLLQAIGKFPAICGYVGKEDESDRICRKTLATMFAHFTQETGNHNPSDNEFEEWRQGLSVV
XP_023324156.1        KIKPGASDNPDNVQRVEFILTEANYEDLFPRRHPSYTYQRLLQAIGKFPAICGYVGKEDESDRICRKTLATMFAHFTQETGNHNPSDNEFEEWRQGLSVV
                              410       420       430       440       450       460       470       480       490       500

                                                                                                  HGT
Eucaro_chimera5_seq1  REQGCT----------------------------------------------------------------------------------------------
XP_023324156.1        REQGCTDTSPGCGYNISYPIIVQGLSVVREQGCTDTSPGLLYASKVVQILLLVVGTAYYHIIVQGLSVVREQGCTDTSPGCGYNSNCEDKDSITKKWSCG
                              510       520       530       540       550       560       570       580       590       600

Eucaro_chimera5_seq1  ----------------------------------------------------------------------------------------------------
XP_023324156.1        KDTEGKFKKYFGRGAKQLSYNFNYGQFSQAMFGESRLLLDHPDFVATTWLNLASATWFYTTPQPPKPSMLHVIDGTWIPNSVDRQNGIEPGFGATINIIN
                              610       620       630       640       650       660       670       680       690       700

Eucaro_chimera5_seq1  ---------------------------------------------------------------------------------------------------
XP_023324156.1        GGLECNTKDGKESNQALNRIAYYKQFAWYLYVDFENETLGCAKQKQFSSGGAGALPIYWDKDWSAPYSCKLVNYQTAHSALVPGEYIHCVEENFQVNIR
                              710       720       730       740       750       760       770       780       790
```

### KAH9406650.1

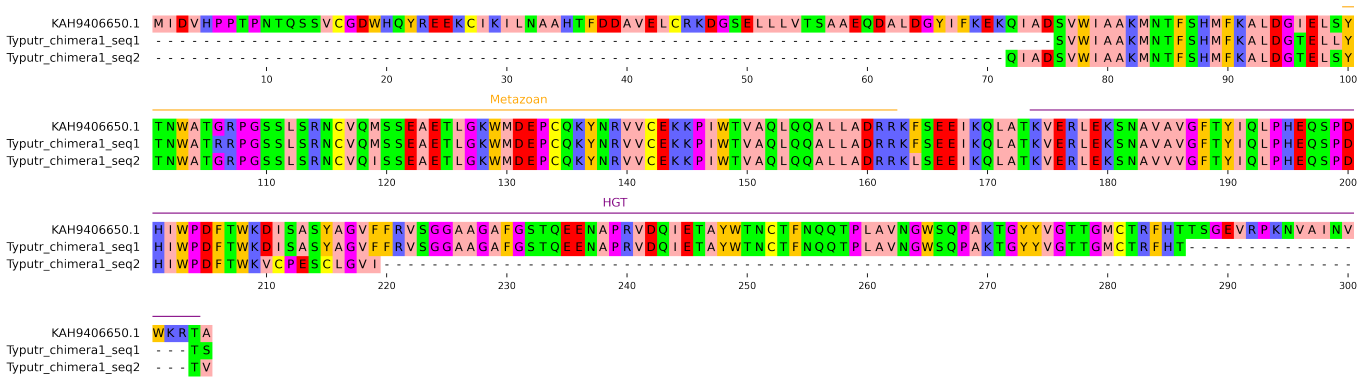

# XP_053634600.2

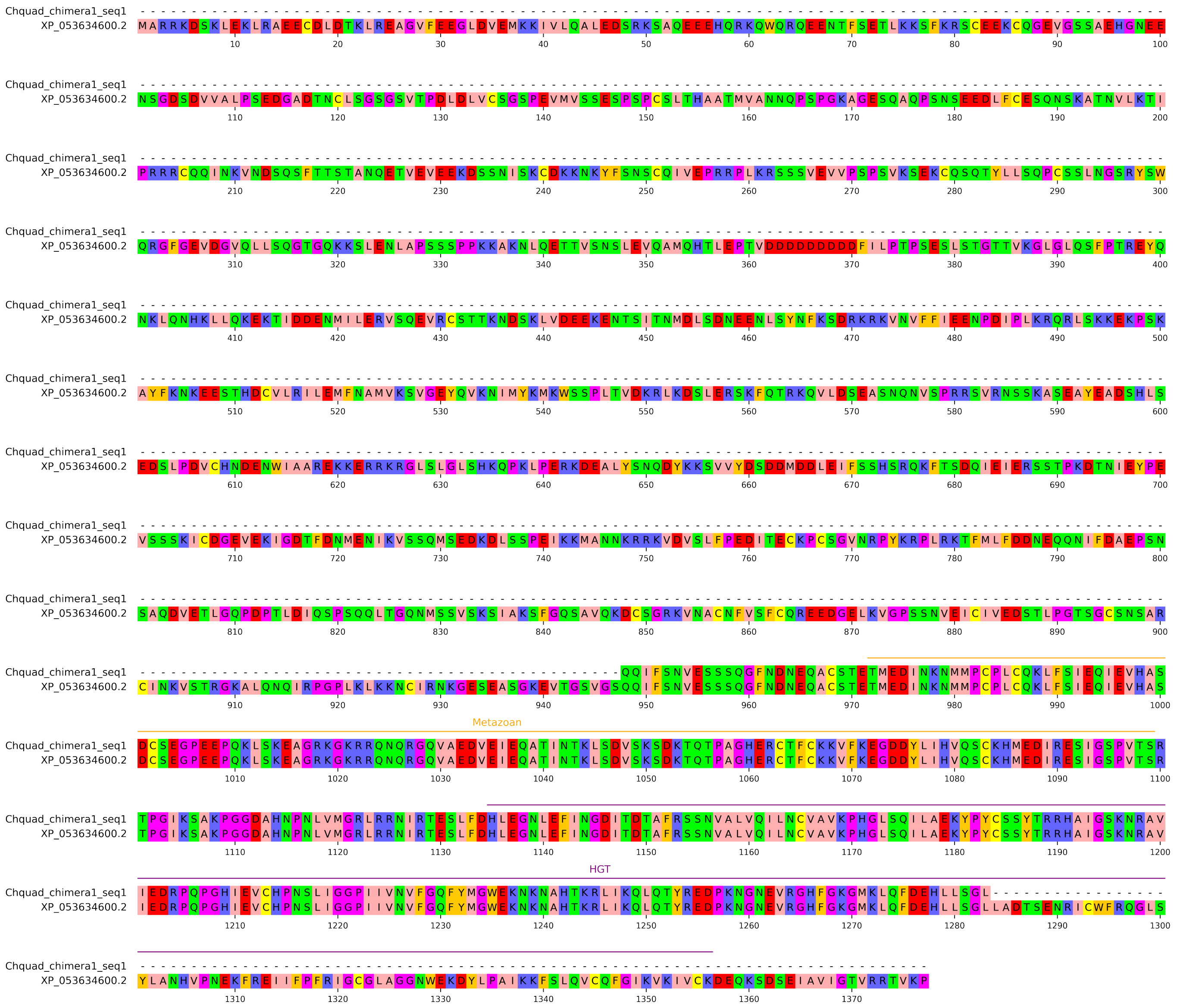

# XP_046649021.1

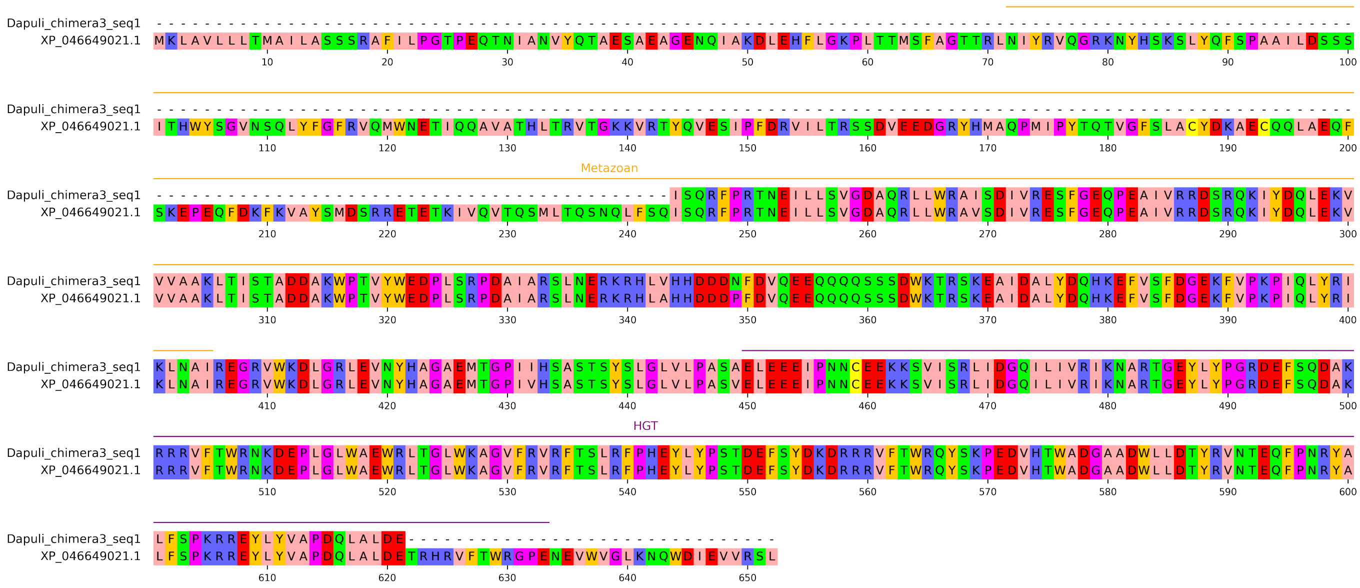

XP_037030969.1

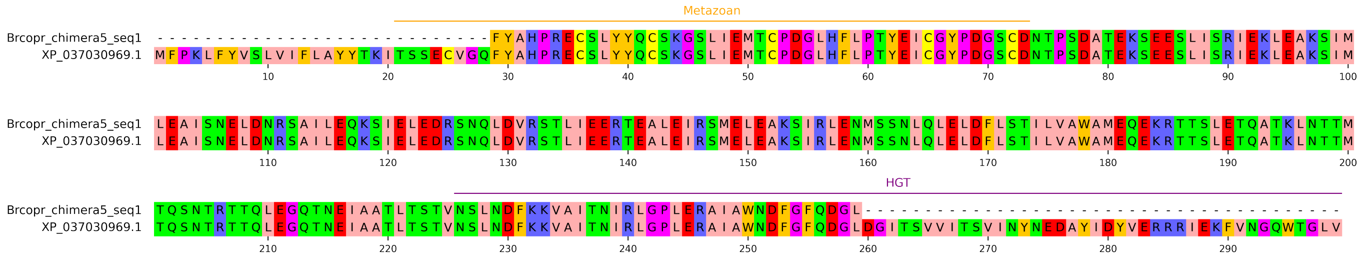

# XP_023328891.1

```
Eucaro_chimera4_seq1  --------------------------TSNTDCQGNLLCGASCLDTTYFPRTQFFGNSETFYSSSDKCCYKKCDKKYNLCLYGSTGCLTSEDCKAGLYCKT
XP_023328891.1        MWDADDDCCTQRCTAKHPCTEGQGHCTSNTDCQGNLLCGASCLDTTYFPRTQFFGNSETFYSSSDKCCYKKCDKKYNLCLYGSTGCLTSEDCKAGLYCKT
                      10 20 30 40 50 60 70 80 90 100

                      Metazoan
Eucaro_chimera4_seq1  DVASPYCEDINECDPVPRMRDGHYPGVLHCGNHATCTNTIGSFSCKCMPGFTDWQMHDGCVDIDECAIGTDNCVANSNCWNTAGSYVCTCKVGYAKNNVG
XP_023328891.1        DVASPYCEDINECDPVPRMRDGHYPGVLHCGNHATCTNTIGSFSCKCMPGFTDWQMHDGCVDIDECAIGTDNCVANSNCWNTAGSYVCTCKVGYAKNKVG
                      110 120 130 140 150 160 170 180 190 200

Eucaro_chimera4_seq1  ACVDINECAQQGTLDYCGPKATCVNTVGSFRCDCLVGFENFKPYKGCSDIDECADPSICGTGYKNCYNNIGSYTCDCTVGWLDWTPLGGCKKDFCYQDKP
XP_023328891.1        ACVDMNECAQQGTLDYCGPKATCVNTVGSFRCDCLVGFENFKPYKGCSDIDECADPRICGTGYKNCYNNIGSYTCDCMVGWTDWTPVGGCIRDFCYIDRP
                      210 220 230 240 250 260 270 280 290 300

                      HGT
Eucaro_chimera4_seq1  NCYANAKCSNTASGYICLCQPYWHYWAPNKGCLTECLGRSSDGTCCTTTEKCDYGRG----------------------------------
XP_023328891.1        NCYANAKCGNTVLGSTCLCEPYWHYWVDKKGCLTECLGRSSDATCCTKTELCDYGEGDCDSDADCKPGLTCGTDNCIQFYAGAVASYDCCI
                      310 320 330 340 350 360 370 380 390
```

# XP_050513096.1

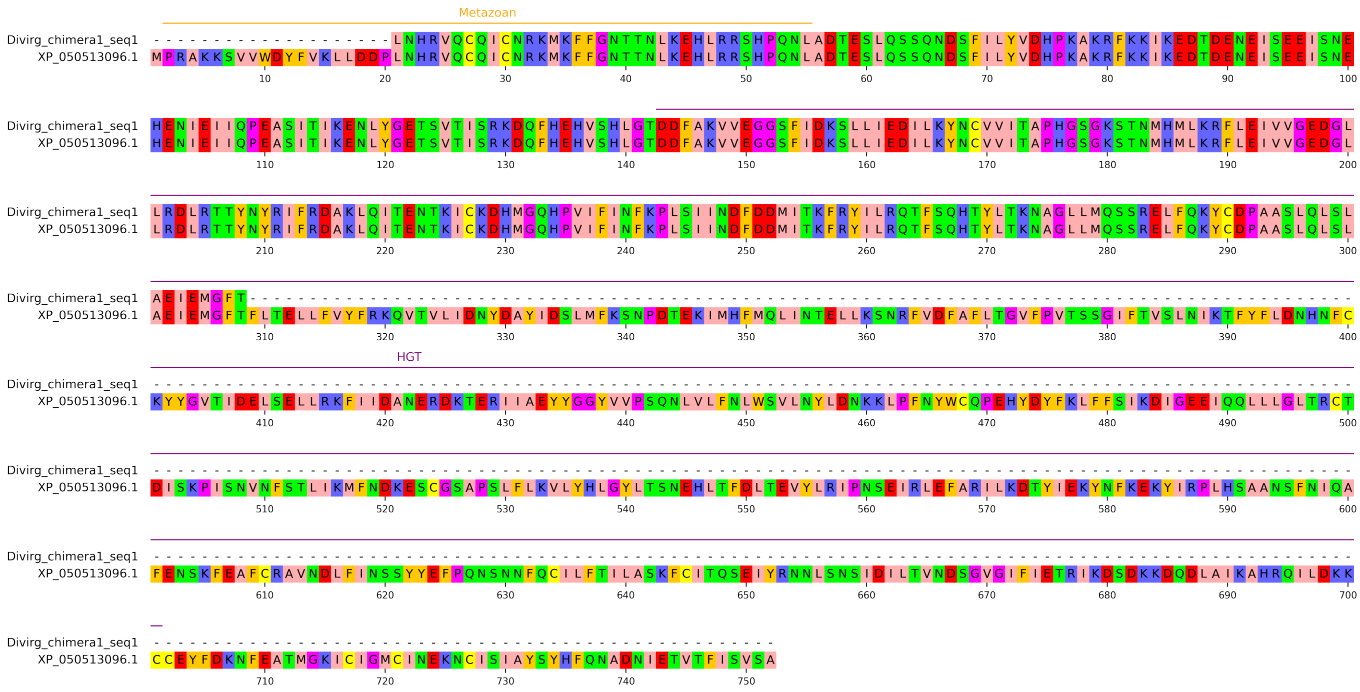

# XP_032791327.2

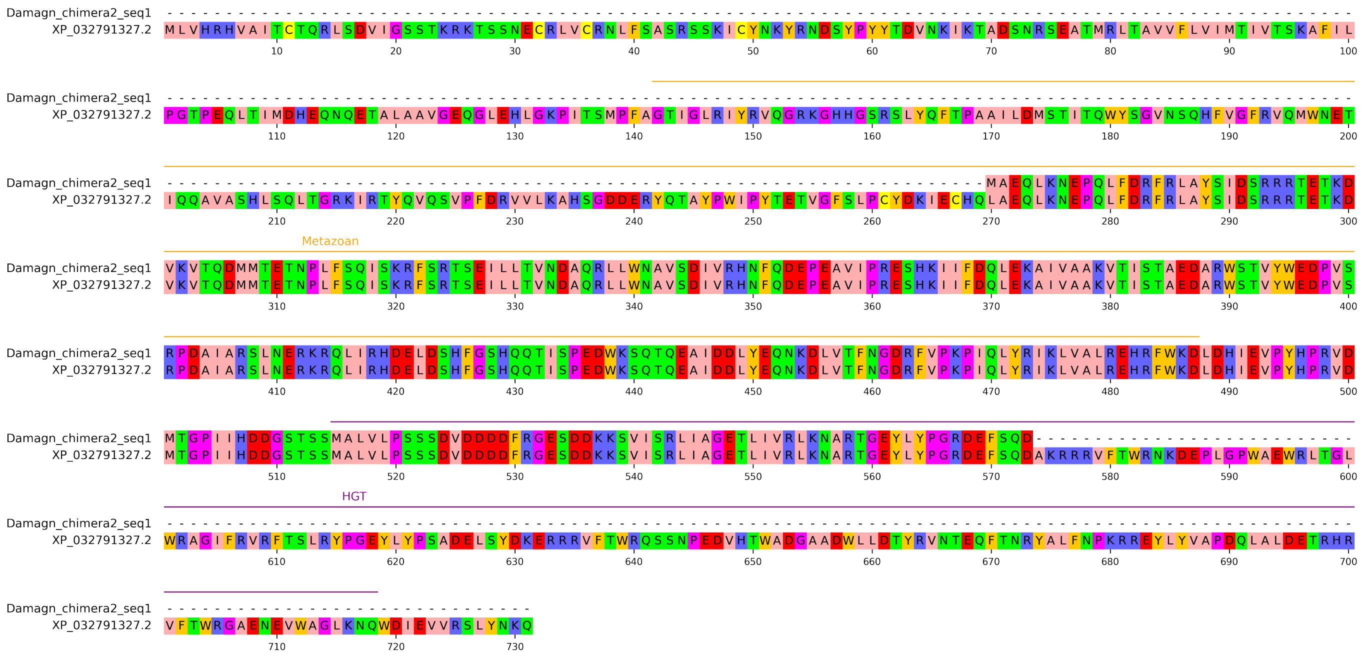

# XP_023332299.1

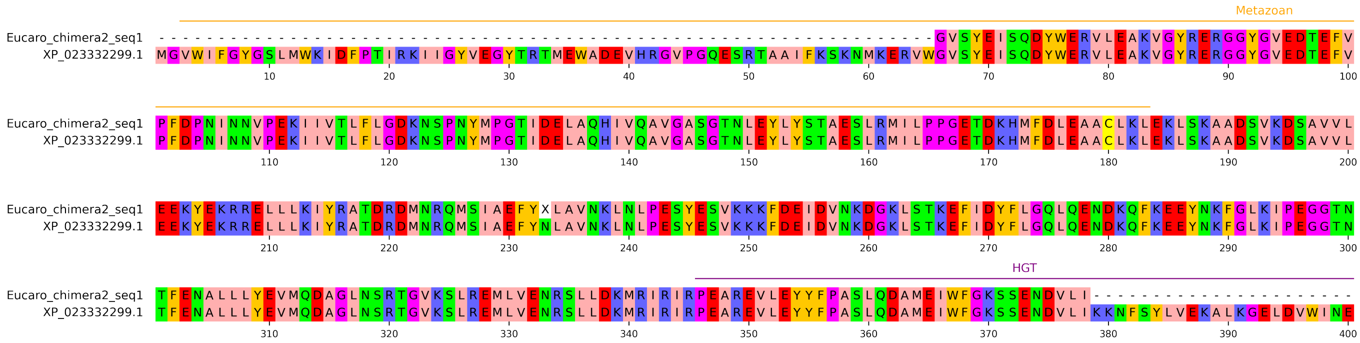

# XP_059092480.1

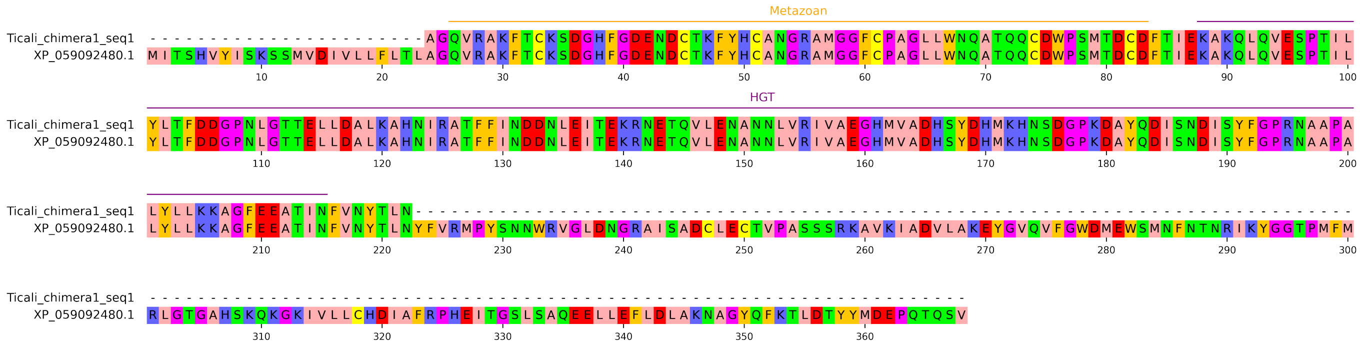
