## Supplementary Information for "Evolutionary innovation through fusion of sequences from across the tree of life"

**This PDF file includes:**

SI Text 1 to 4
SI Methods
SI Figures 1 to 13
Legends for SI Tables 1 to 21
Legends for SI Files 1 and 2
SI References

**Other supporting materials for this manuscript include the following separate documents:**

SI Tables 1 to 21
SI Files 1 and 2

### Supporting Information Text

*SI Text 1: Possible explanations for the observed excess of species-specific chimeras*
77.9% (81/104) of identified HGT-chimeras were found in only a single species (SI Figure 4C, SI Table 4). The two species with the greatest number of species-specific chimeras (N=8 for both *Folsomia candida* and *Eurytemora carolleeae*) are both separated from their closest searched relatives by hundreds of millions of years (242 MYA for *Folsomia candida* vs. *Orchesella cincta* (1); 446 MYA for *Eurytemora carolleeae* vs. *Tigriopus californicus* (2)). We therefore speculate that at least some of the observed excess of species-specific singletons reflects the poor genomic sampling of many arthropod lineages. We nonetheless uncover evidence of species-specific (SI Table 4) or young (origin <20 MYA, SI Table 5) HGT-chimeras even in densely sampled lineages such as the genus *Daphnia* and order Lepidoptera. Excesses of species-specific genes have been reported in studies of gene birth via other mechanisms (3–6), suggesting that some (though not all, see SI Table 5) newly formed genes may be short-lived (3, 6) due to weak (3, 6, 7) or evolutionarily transient (5, 8) selective constraint.

*SI Text 2: Inter-arthropod transfer is a parsimonious explanation for the sparse taxonomic distribution of HGT-chimera 3*
The 104 chimeric sequences from cluster 3 are exclusively predicted in 11 species of four divergent clades that shared a common ancestor over 500 million years ago in the last common ancestor of all extant arthropods (9), as follows: two species of springtails (subphylum Hexapoda, subclass Collembola), three species of fungus gnats (family Sciaridae, order Diptera), one tephritid fruit fly (family Tephritidae, order Diptera), and four sarcoptiform mites (order Sarcoptiformes, class Arachnida) (SI Figure 7A).

To experimentally validate this observation, we assessed gene expression and transcript sequence via RT-PCR and Sanger sequencing for a single cluster representative per genome from four species representing all four aforementioned lineages: *Bradysia coprophila* (accession XP_037051404.1, family Sciaridae), *Rhagoletis zephyria* (accession XP_017483539.1, family Tephritidae), *Folsomia candida* (accession XP_035715507.1, class Collembola), and *Tyrophagus putrescentiae* (accession KAH9406650.1, order Sarcoptiformes). We confirmed the expression and sequence of the predicted HGT-chimera in all tested species except for *R. zephyria* (SI Table 13). We propose that the presence of this predicted HGT-chimera cluster in the *R. zephyria* genome is likely an artifact of contamination of the *R. zephyria* reference genome sequence with nucleotide sequences from the mite *Tyrophagus putrescentiae*, noting that both predicted cluster representatives in the *R. zephyria* genome (XP_017483539.1 and XP_017490256.1) have >90% nucleotide identity with *T. putrescentiae* sequences, and that we failed to obtain amplicons from one of those *R. zephyria* sequences (XP_017490256.1) from cDNA. Further, there is evidence of systematic contamination of the *R. zephyria* genome with sequences from *T. putrescentiae* (10). In contrast, the substantial amino acid divergence observed among cluster representatives from the remaining ten non-Rhagoletis species (SI Figure 7B), together with the recovery of cluster representatives from independently sequenced genomes of multiple species within each of Sciaridae, Collembola, and Sarcoptiformes (SI Table 21), is inconsistent with contamination as an explanation for the distribution of this cluster across these arthropod groups (e.g. via mite infestation).

Excluding the putative contaminants in *R. zephyria*, the distribution of the remaining 102 sequences is consistent with one of three possible histories. First, similarly to our proposal for cluster 18 (see SI Figure 6), representatives of this cluster may have arisen via independent fusion events in three separate lineages of arthropods. This hypothesis predicts that the maximum likelihood trees constructed from the hmmsearch hits of HGT- and metazoan-intervals would show chimeric sequences from each independent origin group within different clades of non-metazoan or arthropod sequences, respectively. However, inconsistent with this expectation, both the HGT and metazoan intervals of this cluster from collembolans, sarcoptiform mites, and sciarids are closely related to each other despite the hundreds of millions of years of divergence separating these lineages (SI Figure 7C-D). A second hypothesis, that this HGT-chimera family formed in the last common ancestor of all extant arthropod groups, would require subsequent loss at least 19 independent times (bold branches in SI Figure 7A). Alternatively, the maximum likelihood gene tree (SI Figure 7B) for the full-length chimeric sequences of cluster 3 suggests a third hypothesis, that the HGT-chimera formed in Sarcoptiformes (mites), and was independently transferred at least twice, once to Sciaridae (fungus gnats) and once to Collembola (springtails).

Inferring the true number and sequence of inter-arthropod transfers from the maximum likelihood tree is complicated by the complex tree topology (SI Figure 7A), in which sequences of all three classes are interleaved with non-monophyletic groups. An ultimate origin in Sarcoptiformes is, however, independently supported by the observation that the divergence time of the Sarcoptiformes lineages containing this HGT-chimera (412 MYA) (11) antecedes the divergence time of crown Collembola (314 MYA) (1) and Sciaridae (82.4 MYA) (12). Thus, if the Sarcoptiformes chimeras in this cluster were inherited from a common ancestor, this chimera must have been present in sarcoptiform mites prior to its acquisition by sciarids and collembolans.

The inter-arthropod HGT scenario above is more parsimonious than the alternative of vertical descent with rampant gene loss, and is also biologically plausible due to the shared soil-dwelling niche of sarcoptiform mites, springtails, and fungus gnats (13, 14). We note that the possibility of inter-arthropod HGT between mites, springtails, and fungus gnats has also been suggested by recent studies of other gene families (13, 15). The mechanism of such inter-metazoan transfer remains unknown, although the presence of a putative bacteriophage-derived sequence in the HGT interval of this sequence (Figure 4) suggests the possibility of transduction by horizontally-transmitted bacterial endosymbionts (16–19) and/or their bacteriophages as a plausible mechanism (20–22).

*SI Text 3: Chitin-interacting copepod HGT-chimeras.*

Both constituent intervals of HGT-chimera cluster 14 are related to genes that function at the host-microbe interface. The HGT-interval is related to *Vibrio* chitin deacetylases, which are utilized by *Vibrio* to metabolize host chitin (23). The metazoan interval is annotated as a peritrophin, which is a class of chitin-binding proteins that maintains a gut barrier to bacterial infection across arthropods (24). Peritrophins and other chitin-interacting or chitin-modifying genes are transcriptionally induced in copepods in response to *Vibrio* exposure (24). Like peritrophins, HGT-chimera cluster 14 has predicted soluble and extracellular localization (DeepLoc probabilities >0.94 and >0.82, respectively; SignalP likelihood > 0.99). Notably, we detect two other copepod HGT-chimeras (clusters 23 and 60) in which both HGT and metazoan intervals are predicted to interact with chitin, raising the possibility of recurrent evolution of chimeras that function at the copepod-microbe interface.

To test the hypothesis that *Eurytemora affinis* (synonymous with *Eurytemora carolleeae*) HGT-chimeras with predicted chitin interaction could be transcriptionally responsive to exposure to symbiotic *Vibrio* bacteria, we analyzed publicly available RNA-Seq data (24) from *E. affinis* exposed to symbiotic *Vibrio* sp. F10 9ZB46, *E. affinis* exposed to free-living *Vibrio ordalii*, and *E. affinis* that was not exposed to *Vibrio*. We found that predicted HGT-Cs from clusters 14 and 23 in *E. affinis* were moderately, but not statistically significantly, upregulated upon exposure to symbiotic but not free-living *Vibrio* species in this dataset (nominal Wald test nominal p-value<0.05; not significant after transcriptome-wide FDR correction; see SI Methods). This could mean that chitin-interacting HGT-chimeras may participate in *Vibrio*-copepod interactions.

*SI Text 4: HGT-chimeras plausibly formed via duplication-mediated gene fusion*

We hypothesized that, similarly to other gene fusion events, HGT-chimera formation might be driven by duplication events, in which a duplicated copy of a HGT and/or metazoan sequence undergoes fusion with a sequence of the opposite ancestry (Figure 3C, SI Figure 9, SI Table 9) (25–28). This duplication-based hypothesis predicts that HGT-chimera sequences should have non-chimeric homologs (“relatives”) to their HGT and/or metazoan intervals elsewhere in the same genome. To test this hypothesis, we searched for evidence of such relatives via sequence similarity (DIAMOND BLASTp) and phylogenetics-based methods, both of which yielded results consistent with this prediction.

First, using BLASTp (SI Figure 9A), we considered an interval of a chimera to have a non-chimeric within-genome relative if it had a BLASTp hit to a protein encoded by a different locus (E-value <1e-10) and if the protein was not simultaneously a hit (E-value <10) for all chimera intervals of the opposite ancestry annotation (i.e. a metazoan relative should be a hit for the metazoan interval of a protein but not the HGT interval). We found that 78.8% (82/104) of HGT-chimeras had within-genome hits (E-value < 1e-10) for at least one constituent interval, and that 44.2% (46/104) of HGT-chimeras had hits for both HGT and metazoan intervals (SI Figure 9B). BLASTp-inferred HGT-chimeras additionally displayed genomic signatures consistent with gene fusion via tandem duplication (N=26 intervals) or retroduplication (N=12 intervals), two established mechanisms of gene birth via fusion (SI Figure 9C-D) (5, 26, 29–31). These data are consistent with a model in which some HGT-chimeras form via horizontal transfer followed by duplication-based gene fusion, as previously posited for a smaller number of HGT-chimeras in bacterial genomes (25).

Since the BLASTp-based strategy above may trivially detect distantly diverged proteins sharing similar domains, we additionally pursued a phylogenetic approach for relative detection. We wrote a custom ete3-based script to parse the maximum likelihood trees generated for HGT/metazoan ancestry inference for possible within-genome relatives. The script first selects the terminal branch of the tree containing a primary HGT-chimera and then identifies the closest branch with at least one non-chimeric sequence. If this neighboring branch contained a non-chimeric sequence from a different gene in the same genome (i.e. species) as the primary HGT-chimera, the non-chimeric sequence was hypothesized to be a relative (SI Figure 9E-F, SI

Table 9). By these criteria, we found that 53.8% (56/104) of HGT-chimeras had at least one constituent interval with an identified relative (SI Figure 9G, SI Table 9).

We also wrote an ete3-based script to parse the trees for non-chimeric relatives of HGT intervals in species other than the species with the primary HGT-chimera (SI Figure 3H; see main Figure 3B for a real example). Starting from the primary HGT-chimera, neighboring branches were successively added to a search set until a neighboring branch with no arthropod sequences was encountered. Any non-chimeric arthropod sequence found in this pooled set of phylogenetic relatives was reported as a possible non-chimeric relative (SI Table 10). We found that 65.1% (71/109) HGT-intervals had phylogenetic relatives in arthropod species that lacked HGT-chimeras, providing further support for the hypothesis that HGT-chimeras are preceded by non-chimeric intermediates (Figure 3C).

.

### Supporting Information Materials: Software and algorithms

| Software or algorithm name & version | Reference | URL |
|---|---|---|
| AlphaFold3 | (32) | https://alphafoldserver.com/welcome |
| Biopython v.1.83 | (33) | https://biopython.org/ |
| CENSOR web application | (34) | https://www.girinst.org/censor/ |
| cNLS Mapper | (35) | https://nls-mapper.iab.keio.ac.jp/cgi-bin/NLS_Mapper_form.cgi |
| DeepGO-SE | (36) | https://github.com/bio-ontology-research-group/deepgo2 |
| DeepLoc v.2.1 | (37, 38) | https://services.healthtech.dtu.dk/services/DeepLoc-2.1/ |
| DESeq2 v.1.46.0 | (39) | https://biocontainers.pro// |
| DIAMOND v.2.0.15 | (40) | https://biocontainers.pro/ |
| ete3 v.3.1.2 | (41) | https://pypi.org/project/ete3/ |
| fpdf v.1.7.2 | | https://pypi.org/project/fpdf/ |
| GPSite | (42) | https://bio-web1.nscc-gz.cn/app/GPSite |
| HMMER v.3.3.2 | (43) | https://biocontainers.pro/ |
| InterProScan webserver, release 5.75-106.0 | (44) | https://www.ebi.ac.uk/interpro/search/sequence/ |
| Interval demarcation algorithm | (45) | |
| IQ-TREE v.2.2.0.3 | (46) | https://biocontainers.pro/ |
| iTOL webserver and API | (47) | https://github.com/iBiology/iTOL; https://itol.embl.de/ |
| matplotlib v.3.9.2 | | https://matplotlib.org/ |
| MembraneFold v.0.0.71 | (48) | https://biolib.com/KU/MembraneFold/ |
| Minimum ancestor deviation rooting (mad.py) | (49) | https://www.mikrobio.uni-kiel.de/de/ag-dagan/ressourcen |
| MMseqs v.2:14.7e284 | (50) | https://biocontainers.pro/ |
| MUSCLE v.5.1 | (51) | https://biocontainers.pro/ |
| NCBI Conserved Domain Database (Batch CD-Search) | (52, 53) | https://www.ncbi.nlm.nih.gov/Structure/bwrpsb/bwrpsb.cgi |
| NCBI Genome Data Viewer | | https://www.ncbi.nlm.nih.gov/gdv/ |
| NetworkX v.2.8.8 | (54) | https://github.com/networkx/networkx |
| NumPy v.1.26.4 | (55) | https://numpy.org/ |
| PAL2NAL v.14.1 | (56) | https://biocontainers.pro/ |
| PAML v.4.10.6 | (57, 58) | https://biocontainers.pro/ |
| PyMol v.3.1 | (59) | https://pymol.org/ |
| pyMSAviz v.0.4.2 | | https://moshi4.github.io/pyMSAviz/ |
| Python Imaging Library (PIL/Pillow) v.9.1.1 | Andrew Clark | https://pypi.org/project/pillow/ |
| Python3.6 | | https://www.python.org/ |
| SciPy v.1.7.3 | | https://scipy.org/ |
| seaborn v.0.13.2 | (60) | https://seaborn.pydata.org/ |
| SeqKit v.2.9.0 | (61) | https://doi.org/10.1371/journal.pone.0163962 |
| SignalP-6.0 | (37) | https://services.healthtech.dtu.dk/services/SignalP-6.0/ |
| SRA toolkit v.3.1.0 | (62) | https://biocontainers.pro/ |
| STAR v.0.6.9 | (63) | https://biocontainers.pro/ |
| statsmodels v.0.14.0 | (64) | https://www.statsmodels.org/stable/index.html |
| Subread v.2.0.6 | (65) | https://biocontainers.pro/ |
| Trim Galore v.0.6.9 | (66) | https://biocontainers.pro/ |
| trimAl v.1.4.1 | (67) | https://biocontainers.pro/ |

**Supporting Information Methods**

*Phase I: Input genomes and processing*
319 Ref-Seq annotated arthropod genomes corresponding to scaffold or chromosome-level assemblies were downloaded (Table S1; access date April 19, 2025) and processed to exclude all scaffolds <100 kb in length to reduce the potential for contamination from the outset. The resulting protein database of 7,702,369 proteins was clustered according to length coverage and E-value using MMseqs2 (50) (version 14.7e284, 80% mutual coverage cutoff, 1e-3 E-value), leading to a reduced database of 610,348 proteins. 11 RT-PCR validated sequences recovered as HGT-chimeras in a previous pipeline iteration (see SI Table 20 for a comparison of pipeline iterations 1 and 2) were then appended to this database for consistency across analyses, yielding an initial search set of 610,359 proteins.

*Phase II: BLAST-based screen for HGT-chimeras*
We reasoned that the junctions between HGT and metazoan intervals might not necessarily correspond to the junctions between annotated protein domains. We therefore devised a strategy to partition proteins into intervals of possibly distinct ancestry directly from BLASTp hits. We used DIAMOND (40) BLASTp (version 2.0.15 – very sensitive, E-value < 10) to query all 610,359 proteins against a local version of the NR database (downloaded October 9, 2023). Protein query BLASTp hits were partitioned into intervals using a modified version of a previously described algorithm (45) (SI Figure 2). The algorithm iteratively performs the following tasks:

1. Identifies a global maximum in the density of BLASTp hits
2. Assigns all proteins overlapping the global maximum to the interval *i*
3. Trims the bounds of the interval *i* to 20% of the density of the global maximum

Steps 1-3 were repeated to assign all BLASTp hits to distinct intervals until ten or fewer hits remained. We considered adjacent intervals *a* and *b* to be overlapping if either >15% of the length of *a* overlapped with *b*, or >15% of the length of *b* overlapped with *a*, and consolidated overlapping intervals by discarding the interval with fewer overlapping BLASTp hits. We retained intervals of length >35 amino acids for subsequent analysis.

Next, we hypothesized preliminary ancestry state-annotations as "metazoan" or "non-metazoan HGT" for each interval through *ad hoc* Boolean rules as follows:

1. Metazoan:
   a. Greater than five non-arthropod BLASTp hits are metazoan with "metazoan index" MI>1, where $MI = log_{10} \frac{min(non-metazoan\ e-value+1e-200)}{min(non-arthropod\ metazoan\ e-value+1e-200)}$
   b. OR ≥ 50% of the NCBI taxonomic IDs of the top 300 non-arthropod BLASTp hits are metazoan; else:
2. HGT:

a. At least ten non-metazoan hits are found with "alien index" AI >5, where $AI = log_{10}\frac{min(non-arthropod\ metazoan\ e-value+1e-200)}{min(non-metazoan\ e-value+1e-200)}$ as previously defined (4, 68, 69)

OR

b. ≥ 95% of the NCBI taxonomic IDs of the top 300 non-arthropod BLASTp hits are non-metazoan.

We derived these *ad hoc* rules via iterative inspection of BLASTp alignment plots. All corresponding plots for proteins that passed the initial filters are available in our Dryad repository (https://doi.org/10.5061/dryad.t1g1jwtdz) and plots for 104 final representative chimeras are available in SI File 1. We constructed these rules according to the expectation that most protein intervals should be of metazoan ancestry. Thus, the rules for designating possible HGT ancestry were more stringent than those for designating metazoan ancestry. We aimed to make our HGT inference robust to contamination of non-arthropod genomes with arthropod sequences by requiring hits in multiple non-metazoan genomes.

Finally, we introduced two novel criteria intended to reduce false negatives of HGT inference in cases of multiple independent transfers of the same protein family to different metazoan groups, which has been reported for an increasing number of horizontally acquired genes (14, 70–75). First, we excluded from consideration any BLASTp hits to sequences from the phylum Rotifera, which has been reported to have high levels of HGT (68, 76). Given that rotifers and arthropods are part of Spiralia and Ecdysozoa, respectively, and considering their divergence more than 600 million years ago (77), we deemed it unlikely that a gene found in no other metazoan genomes would have existed in the Spiralia-Ecdysozoa last common ancestor and been inherited via vertical descent. Rotifer hits were added back to our datasets for phylogenetic screening (see below). Second, intervals that failed to meet the traditional AI-based criterion (4, 68, 69), potentially due to independent transfer events to other metazoan groups, were still considered for downstream phylogenetic inference of HGT if there was a strong non-metazoan bias in their top non-metazoan hits (criterion 2b above). Although these preliminary criteria were sufficiently flexible to detect multiple possible HGT scenarios, only 3.1% (19,179/610,359) of our query proteins had at least one putative HGT interval, in contrast to 51.8% (316,268/610,359) with at least one putative metazoan interval. Moreover, subsequent filtering and phylogenetic analysis eliminated 99.4% (19,070/19,179) of the HGT intervals that passed this step (SI Figure 1). We note that this procedure excluded intervals without detectable BLASTp hits in non-arthropod proteins, precluding detection of fusions of HGT intervals with arthropod-specific proteins.

Next, we collected all proteins that contained at least one metazoan interval and one putative HGT interval (N = 1,687 proteins). Each interval (N = 3,855) was expanded by ten residues at each end, then subjected to a second round of DIAMOND BLASTp (–very-sensitive, E-value ≤ 10; see SI Figure 3). This two-step approach was intended to improve sensitivity by focusing alignments on the candidate HGT or metazoan interval itself. We then applied the Boolean-rule framework as in the first round, with two additional criteria: (i) HGT intervals must now have BLASTp hits in > 10 unique non-metazoan NCBI taxonomic IDs (even when no non-arthropod metazoan hits were recovered) and (ii) at least one non-metazoan hit with E-value < 1 × $10^{-4}$ or bit-score > 50. We calibrated our cut-offs using the OSK domain of Oskar (78), which we previously showed returns only marginal BLASTp similarity (bitscore ~50) to bacterial sequences, yet is robustly recovered as a bacterial-origin HGT by hmmsearch and maximum-likelihood phylogenetics (see "*Phase IV: Phylogenetic inference of HGT or metazoan origin*") (79). For metazoan intervals, we imposed the same criteria as in the first round of BLAST-based inference, but with the additional requirement that at least one non-arthropod metazoan hit have an E-value < $1 \times 10^{-1}$. An identical set of criteria was used to make HGT or metazoan ancestry calls from protein-wide BLASTp hits for comparison to the results of the interval-demarcation approach (SI Table 19). Finally, we discarded any putative HGT-chimera for which adjacent metazoan and HGT intervals both returned the same non-arthropod hit (E-value < $1 \times 10^{-2}$), as this pattern is consistent with a non-chimeric origin. We recovered 525 putative HGT-chimeras at this stage.

*Phase III.1: Exclusion of ankyrin repeat proteins*

Of the 525 HGT-chimera candidates at this stage, 71 proteins (13.5% of the set) returned at least one ankyrin-repeat annotation when queried against the NCBI Conserved Domain Database (52, 53) (Batch CD-Search web-server, default parameters). The ankyrin domain is a short, repetitive motif conserved across eukaryotes, but more sparsely distributed amongst prokaryotes—most often in pathogens and endosymbionts—where its origins are ambiguous and have been attributed to HGT from eukaryotic hosts (80–82) or to convergent evolution (82). These repeats are especially common in the arthropod endosymbiont *Wolbachia* (82) and its bacteriophages (20), where they were possibly acquired from arthropods and later transferred back to arthropod genomes. However, previously reported gene trees produced for arthropod and *Wolbachia* ankyrin repeats were difficult to interpret (20, 83). We reasoned that multi-directional transfers, along with the possibility of convergent evolution and spurious homology detection in this family (83), might complicate inference of unambiguous non-metazoan to arthropod transfers, and therefore excluded all ankyrin repeat-containing proteins from downstream analyses.

*Phase III.2: Exclusion of metazoan transposable elements*

Of the 454 remaining proteins following removal of those containing ankyrin repeats, 215 (47%) returned significant matches to metazoan transposable elements (TEs) in RepBase (release 04/28/2025) via CENSOR (34). While this apparent enrichment of TEs among our HGT-chimeras may reflect the former's frequent participation in HGT and gene fusion events, we reasoned that TE-derived intervals may also confound HGT inference. Repeat masking often removes standalone TEs from genome annotations (34), which could lead to erroneous presence-absence distributions in our protein-based screen. In addition, TEs are both related to (84) and exchange segments with viruses (85–87), complicating efforts to distinguish unidirectional virus to arthropod transfers from vertically-transmitted TEs, or from transfers in the reverse direction. We therefore excluded any HGT interval overlapping a CENSOR-annotated metazoan TE.

We permitted two exceptions to this rule. First, we included hits to the second open reading frame for the Medea element from *Tribolium castaneum*, which has been shown to be of recent HGT origin from a bacterial AAA ATPase (88, 89). Second, if the DIAMOND-BLASTp bitscore of the top non-metazoan NR hit exceeded that of the best metazoan TE match by ≥ 1.7× (i.e., bitscore_non-metazoan ≥ 1.7 × bitscore_top_metazoan_TE), we retained the interval.

This 1.7× threshold was empirically chosen to preserve HGT-chimera intervals showing strong alignment to arthropod-infecting viruses, thereby favoring true HGT events over TE-fusion artifacts. This filtering process removed 89 candidates, resulting in 365 remaining candidates. We note that this procedure did not exclude proteins with non-metazoan TE annotations, nor proteins with metazoan TE annotations outside HGT intervals.

*Phase III.3 Search for HGT-chimera orthologs*

We sought to include not only the 365 proteins (henceforth "primary HGT-chimeras") identified above, but also their putative orthologs ("secondary HGT-chimeras") in other arthropod genomes in our downstream phylogenetic analysis. To that end, we searched for putative orthologs across an expanded arthropod proteome (the 319 RefSeq genomes as above + 197 GenBank genomes; SI Table 1). For each HGT-chimera interval, we built a profile Hidden Markov model (HMM); HMMER hmmbuild v.3.3.2) (43) from a MUSCLE (v.5.1) (51) alignment of all round-2 arthropod DIAMOND BLASTp hits with bitscore ≥ max(non-arthropod bit-score). We then queried these profile HMMs against the full arthropod proteome with hmmsearch (HMMER v.3.3.2, E-value < $1 \times 10^{-4}$). Since hmmbuild requires an alignment of ≥ 2 sequences, we considered DIAMOND BLASTp hits (E-value < $1 \times 10^{-4}$) directly instead of hmmsearch results for 79/807 HGT-chimera intervals lacking non-self arthropod hits with bitscore greater than that of the top non-arthropod hits. We retained as secondary HGT-chimeras those sequences that were BLASTp or hmmsearch-hits to all constituent HGT and metazoan intervals in the same linear order as their primary HGT-chimera.

*Phase III.4 Similarity clustering*

To cluster related HGT-chimeras into groups potentially reflecting single-origination events, we constructed an undirected graph (NetworkX v2.8.8) in which an edge connected two primary HGT-chimeras whenever one appeared as a secondary HGT-chimera of the other. We then defined "similarity clusters" as connected components on this graph (N=299). Within each cluster, we chose as the representative sequence the primary HGT-chimera with the greatest number of secondary hits. To maintain consistency across both pipeline iterations (SI Table 20), any HGT-chimera previously selected as a representative sequence in the first iteration was retained in that role even if a different sequence in the cluster later acquired more secondary hits in the second iteration.

We note that this clustering procedure may group proteins that evolved analogous HGT-chimera architectures via independent HGT and/or fusion events in the same cluster, and thus does not guarantee common ancestry. Indeed, some phylogenetic trees (see next section "*Phase IV: Phylogenetic inference of HGT or metazoan origin*" and SI Figure 6) would at face value suggest convergent origins of similar HGT-chimera architectures by virtue of polyphyly of the chimeric intervals relative to their non-chimeric relatives, as was previously observed for the metazoan interval of *oskar* (79). However, considering the susceptibility of phylogenies constructed from short protein alignments to reconstruction artifacts (90), as well as the rarity of both HGT (4, 91) and gene fusion (5, 6), we conservatively treated all members of each HMM/BLAST-defined cluster as descendants of a single origination event (although see SI Figure 6 for an exception inferred from the sparse taxonomic distribution and non-monophyly of both intervals of HGT-chimera cluster 18).

Next, we extracted the hmmsearch- or BLAST-aligned intervals of each secondary HGT-chimera and ran DIAMOND BLASTp (–very-sensitive, E ≤ 10) on each secondary HGT-chimera interval. We applied the same Boolean MI/AI and E-value rules from round 2 BLASTp-based inference (see above) to annotate secondary HGT-chimera intervals as “metazoan” or “HGT.” We conservatively eliminated 41 HGT-chimera candidates for which any secondary HGT-chimera interval gave an annotation conflicting with its primary designation (e.g. a primary HGT interval was annotated as metazoan in its secondary HGT-chimera or vice versa). This resulted in 258 putative HGT-chimeras. Before proceeding to phylogenetic analysis, we manually inspected the round 1 and round 2 BLASTp plots for all 258 representative primary HGT-chimeras, and eliminated 20 additional cases in which the HGT and metazoan regions appeared to be poorly resolved due sequence overlap between the intervals.

*Phase IV: Phylogenetic inference of HGT or metazoan origin*

We constructed phylogenies for each HGT interval from the remaining 238 representative primary HGT-chimeras using multiple sequence alignments of round 2 BLASTp or hmmsearch hits (both with E-value<1e-2), opting for hmmsearch hits only when a profile HMM of ≥ 2 unique arthropod sequences could be constructed (as in the secondary HGT-chimera search above). To broaden the taxonomic sampling of sequences used to construct each phylogeny, we selected a single sequence per species within each of the following categories:

- Secondary HGT-chimeras in other arthropods (as identified above)
- Non-chimeric sequences detected in our arthropod proteome database compiled from RefSeq and GenBank genomes in “*Phase III.4 Similarity clustering*” above
- Non-chimeric sequences detected in non-arthropod metazoan taxa in the NR database (maximum of 500 sequences)
- Non-chimeric sequences detected in non-metazoan taxa in the NR database (maximum of 500 sequences)

Multiple sequence alignments were generated using MUSCLE (51) with default parameters for datasets consisting of <1200 sequences or the super5 algorithm for >1200 sequences or when the default algorithm failed to complete in <48 hours. Alignments were trimmed using trimAl version 1.4.1 (67) with a threshold of 60% column occupancy (--gt 0.6). Maximum likelihood trees were inferred with IQ-TREE version 2.2.0.3 (46) using 1000 ultrafast bootstraps with automatic model selection and subsequently rooted with minimum ancestor deviation (49). Trees were visualized with superimposed taxonomic information obtained using the ete3 toolkit (41) using the iTOL python API and web browser (47).

Tree topologies were evaluated for evidence of non-arthropod to arthropod HGT or metazoan ancestry using a combination of a custom ete3-based script and manual inspection. For each representative HGT-chimera, we identified its “sisters” as the most closely related non-arthropod sequence(s), and its “cousins” as the second-most closely related sequence(s).

Metazoan ancestry was inferred when sisters and cousins were both from non-arthropod metazoan taxa (SI Figure 3A). We relaxed this requirement when sporadic non-metazoan sequences were found in sister or cousin clades if the non-metazoan sequences did not alter an overall pattern of nestedness within a clade of arthropod and non-arthropod metazoan sequences (SI Figure 3B).

HGT ancestry was inferred when both the sisters and cousins were from non-metazoan taxa, with separation from metazoan clades by at least one internal node with bootstrap support >70% (SI Figure 3E). We note that, by this criterion, the tree for the HGT-derived interval of *oskar* (GCF_004354385.1; XP_034487048.1;HGT_(412,553)) had insufficient support for a bacterial vs non-arthropod metazoan affinity (57%), but we nevertheless elected to include Oskar in our final HGT-chimera set due to its HGT-consistent tree topology and prior analyses supporting HGT origin via both Bayesian methods and formal tree topology tests (79). Bootstrap support was not considered for trees lacking multi-sequence clades of non-arthropod metazoan sequences (SI Figure 3F); in this case, no alternative tree topology would support a metazoan origin. For tree topologies in which arthropod and non-arthropod sequences were each other's sister clades, with no additional cousin clades available to polarize the direction of HGT (SI Figure 3D, H), we hypothesized non-arthropod to arthropod HGT if the non-arthropod sequences were from multiple different taxonomic phyla and numerically outnumbered the arthropod sequences in the tree. Consistent with previous a previous study of prokaryote to arthropod HGT (74), we disregarded sporadic non-arthropod metazoan sequences in inference of HGT when the majority of sister and cousin sequences were non-metazoan, such that an overall pattern of nestedness within non-metazoan sequences was maintained (SI Figure 3G). We interpret this type of topology as indicative of multiple independent transfers (or contaminations) from non-metazoan genomes to arthropod and non-arthropod metazoan genomes. By similar logic, trees in which sister or cousin sequences were non-metazoan, but non-metazoan sequences were only sporadically distributed in the tree (SI Figure 3C), were rejected as cases of non-metazoan to arthropod transfer, in favor of arthropod to non-metazoan transfers or contamination.

HGT-chimera candidates showing phylogenetic evidence of at least one HGT interval and one metazoan interval were selected for the final set of HGT-chimera candidates, for a total of 104 HGT-chimera similarity clusters.

*Phylogenetic inference of donor taxa*

Taxonomic origins of HGTs were assigned according to the kingdom or superkingdom/domain-level taxonomic labels of sister and cousin sequences. In cases with multi-sequence sisters or cousin sequences, taxonomic labels were determined by majority rule consensus. Cases of discordance between the taxonomic labels of sister and cousin sequences were resolved via consideration of the next-most closely related sequences (if available) or by majority-rule consensus of pooled sister and cousin sequences. Sister and cousin clades of intervals of inferred bacterial origin were also inspected for the presence of arthropod bacterial symbiont genera derived from the Symbiotic Genomes Database (92) (access date 6/29/2025) (SI Table 8).

*Codon use and GC content analysis*

GC content scores were calculated for all HGT and metazoan intervals within HGT-chimeric sequences using Biopython (33). The deviation in codon usage from the genomic background for these intervals was assessed using the Measure Independent of Length and Composition

(MILC) method implemented as a custom Biopython script according to the formulae in (93). This method accounts for codon bias independently of amino acid composition and sequence length. Expected (background) codon frequencies were determined by analyzing all coding sequences (CDS) in the dataset.

Within-genome background distribution of MILC and GC content values were produced for each interval by performing 10,000 iterations of sampling of in-frame intervals of the same length from CDS sequences (SI Figure 8A-B, SI Table 11). The sampled background distribution was used to generate a percentile score for each interval. Matched pairs of HGT and metazoan intervals from the same HGT-chimeras were compared in their MILC and GC content percentile scores using the Wilcoxon signed-rank test in scipy.

*Expression support from publicly available RNA-Seq data*

We queried the RefSeq annotation metadata for all predicted HGT-chimeras (SI Table 12). HGT-chimeras marked with “CDS support: full” have RNA-Seq support over the full length of the predicted CDS. Those lacking this label include *ab initio* predictions over at least some portion of their length. We determined whether the metazoan and HGT intervals were both spanned by RNA-seq reads over >50% of their length by inspecting the NCBI genome browser for the corresponding RefSeq genome annotation (SI Figure 8C).

*Expression support from mRNA isolation and cDNA sequencing*

<u>Sample receiving and storage</u>: Arthropod tissue samples were obtained as donations or collected by the authors from lab-cultured animals (SI Table 13, sheet “sample_data”). Samples (SI Table 13) were received frozen or in RNAlater (Invitrogen, catalog # AM7020) or DNA/RNA Shield (Zymo, catalog # R1100). Samples received or harvested live were submerged in TRIzol reagent prior to being flash-frozen in liquid nitrogen then transferred to storage at -80ºC. Samples that were received flash-frozen on dry ice or in RNAlater or DNA/RNA Shield were transferred directly to -80 ºC, and TRIzol was added immediately prior to RNA extraction.

<u>RNA extraction and cDNA synthesis:</u> RNA was extracted using TRIzol Reagent (Invitrogen, catalog # 15596018). Total RNA was digested with Turbo DNase (Invitrogen, TURBO DNA-free kit catalog # AM1907 and then treated with DNase Inactivation Reagent (Invitrogen, TURBO DNA-free kit catalog # AM1907) before assessing RNA quantity using the Qubit RNA Broad Range Assay Kit (Invitrogen, catalog # Q10210) using a Qubit Flex Fluorometer (Invitrogen, catalog # Q33327). 1 µg of total RNA was used to synthesize cDNA using the Maxima H Minus First Strand cDNA Synthesis Kit (Thermo Scientific, catalog # K1652) with oligo(dT) primers to selectively target eukaryotic poly-adenylated mRNAs.

<u>PCR and Sanger sequencing:</u> Primers encompassing parts or the entirety of both HGT and metazoan domains of selected HGT-chimeras were designed with Primer-BLAST (94) using standard parameters. If the mRNA interval between the HGT and metazoan regions exceeded 1.5 kb, multiple overlapping sets of primer pairs were used to assess expression, and the sequencing products were subsequently manually assembled. PCRs were run using Phusion High-Fidelity DNA Polymerase (New England BioLabs, catalog # M0530S) with annealing

temperatures and extension times listed in SI Table 13, except for one PCR for the *T. castaneum* chimera, for which the PCR was run using KOD One PCR Master Mix (Sigma Aldrich, catalog # KMM-101NVS). Resulting PCR reactions were inspected on 1.0-1.5% agarose gels, and the rest of the reactions were either PCR purified using Monarch Spin PCR & DNA Cleanup Kit (New England BioLabs, catalog # T1130L) or, in case of the presence of multiple bands, run on a preparative agarose gel and gel eluted using The Monarch Spin DNA Gel Extraction Kit (New England BioLabs, catalog # T1120L). The concentration of eluted fragments was assessed using a NanoDrop Eight (Thermo Fisher Scientific, catalog # NDE-GL), and Sanger sequenced (GENEWIZ/Azenta). In case of Sanger sequences indicating the presence of multiple PCR fragment variants, PCRs were repeated, A-tailed using 1 U of DreamTaq DNA Polymerase (Thermo Scientific, catalog # EP0702) for 15 minutes at 72°C, immediately cloned into TOPO pCR4 using the TOPO TA Cloning Kit for Sequencing (Invitrogen, catalog # 450030) and transformed into Mix & Go! DH5 alpha competent cells (Zymo Research, catalog # T3009). Transformants were screened by colony PCR using M13 Forward (-20) and M13 Reverse primers and DreamTaq DNA Polymerase (Thermo Scientific, catalog # EP0702). Plasmids were isolated using the QIAprep Spin Miniprep Kit (Qiagen, catalog # 27106) and sequenced with M13 Forward (-20) and M13 Reverse primers (GENEWIZ/Azenta). All resulting sequences were inspected, corrected and aligned using Geneious Prime version 2025.1.2.

*Selective constraint analysis*

We performed dN/dS analysis on HGT-chimeras found in more than one genome to determine whether HGT-chimeras showed selective constraint consistent with expression as functional proteins. Protein sequences for all secondary HGT-chimeras were aligned using MUSCLE version 5.1 and trimmed using trimAl version 1.4.1 as defined in "*Phase IV: Phylogenetic inference of HGT or metazoan origin.*" The resulting multiple sequence alignment was used to guide codon alignment generation using PAL2NAL version 14.1 (56). Gene trees were generated from the protein multiple sequence alignment using IQ-TREE version 2.2.0.3 with automated model selection. Maximum-likelihood dN/dS values were fitted using the M0 model with codeml (codeml model = 0, NSsites = 0, CodonFreq = 7, fix_omega = 0) in PAML version 4.10.6 (57, 58, 95). To assess whether cases with dN/dS<1 constituted a statistically significant deviation from neutrality, we re-ran M0 with dN/dS fixed at 1 (fix_omega = 1, omega = 1) and compared the likelihood under this null model to the likelihood under a model with a free dN/dS value using a likelihood ratio test implemented in python (scipy chi-squared, 1 degree of freedom). Benjamini-Hochberg correction as implemented in statsmodels was used to generate FDR-adjusted p-values.

We further assessed whether each of 23 HGT intervals from the same 21 HGT-chimeras described above displayed signatures of relaxed constraint relative to the rest of the chimeric gene with fixed-site models (96) using codeml in PAML. We first ran fixed-site models in which codons from HGT intervals were permitted to differ from the rest of the codons in their branch lengths $\mathbf{c}$, transition-transversion ratio $\boldsymbol{\kappa}$, equilibrium nucleotide frequencies $\boldsymbol{\pi}$, and dN/dS values (option G in the codon alignment file, codeml Mgene=4, CodonFreq = 2). We compared the model fit to a null model in which HGT and non-HGT sites share dN/dS and $\boldsymbol{\kappa}$ values but differ in $\boldsymbol{\pi}$ and $\mathbf{c}$ (option G in the codon alignment file, codeml Mgene=2, CodonFreq = 2) using the likelihood ratio test (2 degrees of freedom) with a Benjamini-Hochberg correction as implemented in statsmodels.

*Neofunctionalization analysis*

To determine whether the HGT and metazoan intervals of HGT-chimeras showed evidence of undergoing accelerated protein evolution following gene fusion, we compared the dN/dS values of HGT-chimeric sequences to those of their closest non-chimeric relatives using foreground/background branch models. We limited our analysis to HGT-chimeras (1) with species- or genus-restricted taxonomic span; (2) having the same relative gene via both BLASTp and tree-based approaches above; and (3) having at least three total sequences including the primary HGT-chimera and other sequences in the same taxonomic order. By these criteria, we included 28 intervals belonging to 25 HGT-chimera clusters from this analysis.

For each interval, we identified chimeric and non-chimeric phylogenetic relatives in closely related species using the same maximum likelihood interval trees used for HGT and metazoan ancestry inference described above (see "*Phase IV: Phylogenetic inference of HGT or metazoan origin*"). We selected close phylogenetic relatives on these trees with a custom ete3-based script that selects the leaf node corresponding to the query HGT-chimera and iteratively adds sequences in the same taxonomic order as the query HGT-chimera until a branch with no sequences from the same order is identified. Multiple sequence alignments, codon alignments, and gene trees were produced as described in "*Selective constraint analysis.*" Trees for HGT-chimera intervals found in multiple species in the same genus were rooted according to previously described species relationships for *Daphnia* (97) and lepidopteran (98) sequences. Foreground branches were set to include chimeric sequences and (for HGT-chimeras found in more than one species) the branch separating HGT-chimeras from non-chimeras (Figure 3E). Two-branch dN/dS models (95) were fit using codeml in PAML (codeml model = 2, NSsites = 0, CodonFreq = 7, fix_omega = 0), and a likelihood ratio test with the Benjamini-Hochberg correction was used to compare model fit to a null model with a single dN/dS value across both branches (codeml model = 0, NSsites = 0, CodonFreq = 7, fix_omega = 0).

*Functional annotation*

A fasta file of separated HGT and metazoan-ancestry intervals of all 104 representative HGT-chimeras was submitted to InterProScan (44) (web server, release 5.75-106.0), NCBI CDD-search (52, 53) (web server, default parameters), and DeepGO-SE (36) for functional annotation. DeepGO-SE molecular function and biological process predictions with reported probability ≥0.50 were tabulated. All three annotation types were considered collectively in determination of functional similarity between HGT and metazoan intervals (SI Table 18), with the additional consideration of BLASTp hits when CD-search and/or InterProScan failed to return domain hits.

We used the following tools to further characterize the structure and function of selected examples presented in Figures 2-4: GPSite (42) (for DNA and zinc binding probabilities; SI Figure 13), SignalP-6.0 ((37) for signal peptide detection; Figures 2A, C), DeepLoc v.2.1 ((38) for subcellular localization prediction), cNLS Mapper ((35) for nuclear localization sequence prediction in Figure 3), AlphaFold3 (32) with PyMol v.3.1 ((59) for structure prediction;

SI Figure 11-12), DeepTMHMM (99) for transmembrane topology prediction (Figure 11A),and MembraneFold v.0.0.71 (48) with OmegaFold (100) and DeepTMHMM (99) options (for transmembrane protein structural prediction; Figure 4C).

*Tree topology tests for HGT-chimera cluster 12*
Of the four HGT-chimera examples selected for detailed analysis (Figures 2-4), only cluster 12 contained large clade(s) of non-chimeric arthropod or non-arthropod metazoan sequences that may alter HGT inference under alternative topologies. We therefore performed two constrained ML topology searches with IQ-TREE (substitution model Q.pfam+I+I+R10, as previously selected for HGT inference) to require monophyly of cluster 12 HGT interval sequences with either of two observed arthropod clades (Figure S10). Support for constrained topologies was compared to the unconstrained maximum likelihood topology using the approximately unbiased test (101) in IQ-TREE with 10,000 RELL replicates.

*Search for representatives of HGT-chimera cluster 14 in copepod transcriptome shotgun assemblies (TSAs)*
Since HGT-chimera cluster 14 was initially detected in only two copepod genome annotations, and only three copepod genome annotations were available at the time of writing, we searched for additional orthologs in the following copepod TSAs: GCHA01 (*Tigriopus japonicus*), GHXK01 (*Platychelipus littoralis*), GJRL01 (*Calanus marshallae*), GJGX01 (*Temora stylifera*), GJAO01 (*Metridia pacifica*), GKAB01 (*Acartia tonsa*), HAHV01 (*Tisbe holothuriae*). Predicted translations were obtained with TransDecoder v. 5.7.1 then searched for simultaneous hmmsearch hits to the metazoan and HGT intervals as in similarity cluster inference above. We obtained HGT-chimera hits in six of seven examined TSAs (Figure 2D), with NCBI nucleotide accessions HAHV01008100.1, GHXK01159657.1, GCHA01003214.1, GJRL01075442.1, GJAO01071456.1, GJGX01134564.1. The orthology of the newly-detected TSA chimeras to the previously inferred genomic chimeras was verified by re-computation of the multiple sequence alignments and ML trees for the HGT and metazoan intervals with the addition of the new sequences.

*Differential expression analysis of copepod HGT-chimeras on exposure to* Vibrio
We first used the SRA toolkit v. 3.1.0 to download previously published RNA-Seq data for *Eurytemora affinis* not exposed to *Vibrio* (NCBI SRA accessions SRR1298705, SRR1298712, SRR1298714, SRR1298716), *E. affinis* exposed to symbiotic *Vibrio* sp. F10 9ZB46 (SRR1296555, SRR1297318, SRR1298386, SRR1298416), and *E. affinis* exposed to free-living *Vibrio ordalii* 1509 (SRR1298419, SRR1298422, SRR1298424, SRR1298426). We processed paired reads with Trim Galore v.0.6.9, followed by STAR v.0.6.9 to map the reads to an indexed genome (GCF_000591075.1). Finally, we tabulated exon-mapping reads per gene using featureCounts (in subread v. 2.0.6), and used DESeq2 (v.1.46.0) to perform *Vibrio* sp. F10 vs. control and *Vibrio ordalii* vs. control differential expression analysis for the HGT-chimera genes LOC111712908 (cluster 14), LOC111705930 (cluster 23), LOC111698135 (cluster 60). Two of these three genes, LOC111712908 (cluster 14) and LOC111705930 (cluster 23), had significant nominal p-values (o.026 and .001 respectively) with log2FoldChange values consistent with moderate transcriptional upregulation (0.661 and 0.436 respectively). Although neither gene

reached statistical significance after transcriptome-wide FDR correction (p-values 1.00 and 0.69 respectively), we report nominal p-values here since we restricted our hypotheses to HGT-chimeras that were *a priori* selected for predicted chitin interaction (SI Text 3).

*Software access*

Except where otherwise noted in *"Supporting Information Materials: Software and algorithms,"* all bioinformatics software was accessed via Singularity images distributed by the BioContainers project (102) on the Harvard Faculty of Arts and Sciences Research Computing cluster.

*Data visualization and analysis*

Plots were generated using the matplotlib (v.3.9.2) and seaborn (v.0.13.2) packages in python3.6, and pyMSAviz (v.0.4.2) for multiple sequence alignments of Sanger sequencing product translations with RefSeq protein sequences (SI File 2). These BLASTp plots and multiple sequence alignments were compiled into PDF files (SI Files 1 and 2, respectively) using fpdf (v.1.7.2) and the Python Imaging Library (v.9.1.1). Trees were visualized and annotated using the iTOL API and webserver tools. Statistical tests were performed with statsmodels (v.0.14.0), numpy (v.1.26.4), and scipy (v.1.7.3).

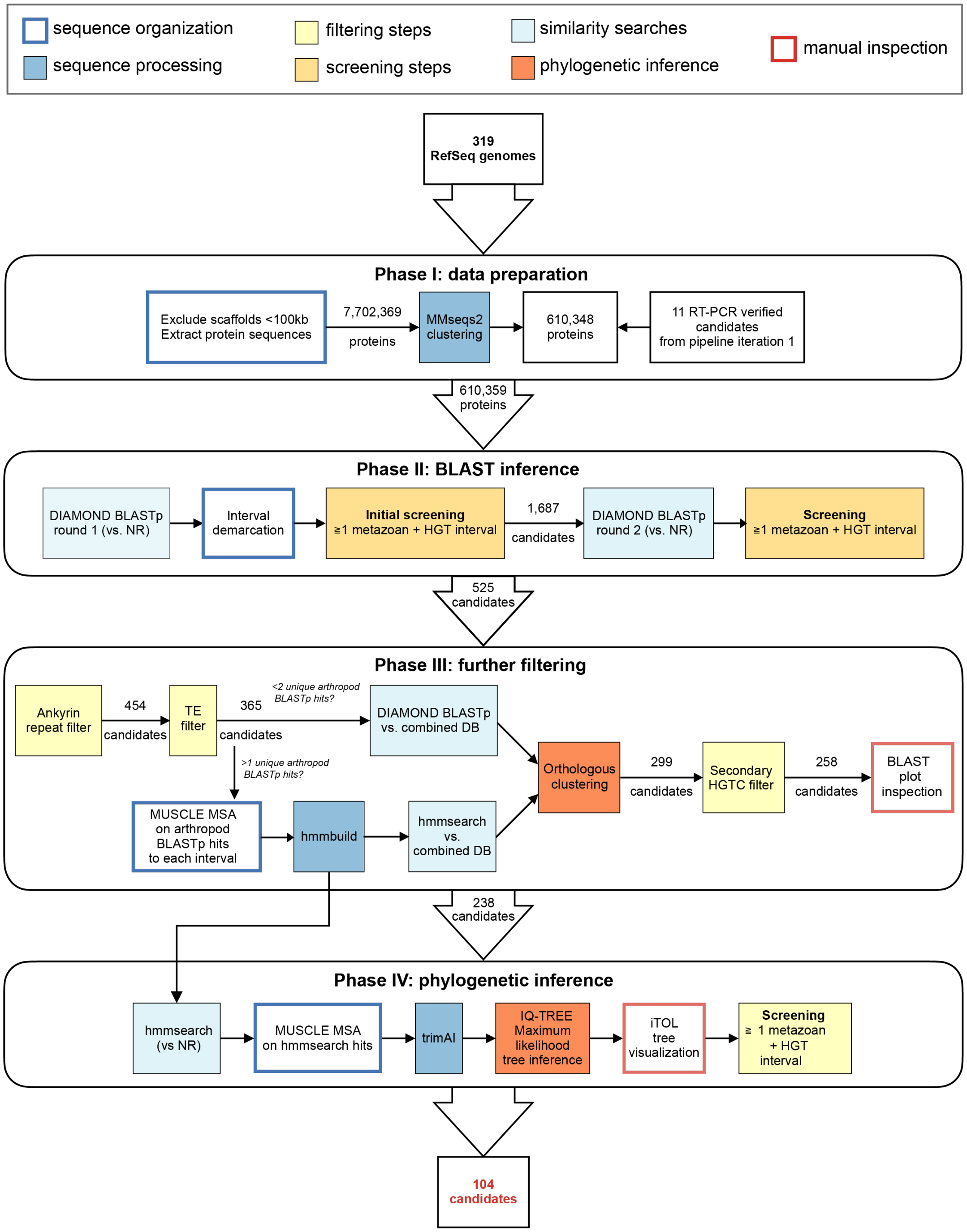
sequence organization
sequence processing
filtering steps
screening steps
similarity searches
phylogenetic inference
manual inspection
319 RefSeq genomes
Phase I: data preparation
Exclude scaffolds <100kb Extract protein sequences
7,702,369 proteins
MMseqs2 clustering
610,348 proteins
11 RT-PCR verified candidates from pipeline iteration 1
610,359 proteins
Phase II: BLAST inference
DIAMOND BLASTp round 1 (vs. NR)
Interval demarcation
Initial screening ≥1 metazoan + HGT interval
1,687 candidates
DIAMOND BLASTp round 2 (vs. NR)
Screening ≥1 metazoan + HGT interval
525 candidates
Phase III: further filtering
Ankyrin repeat filter
454 candidates
TE filter
365 candidates
<2 unique arthropod BLASTp hits?
DIAMOND BLASTp vs. combined DB
>1 unique arthropod BLASTp hits?
MUSCLE MSA on arthropod BLASTp hits to each interval
hmmbuild
hmmsearch vs. combined DB
Orthologous clustering
299 candidates
Secondary HGTC filter
258 candidates
BLAST plot inspection
238 candidates
Phase IV: phylogenetic inference
hmmsearch (vs NR)
MUSCLE MSA on hmmsearch hits
trimAl
IQ-TREE Maximum likelihood tree inference
iTOL tree visualization
Screening ≧ 1 metazoan + HGT interval
104 candidates

**Fig. S1 (previous page). Pipeline schematic for HGT-chimera detection.** Schematic illustration of methodology for HGT-chimera detection, divided into four broad phases as outlined in brief in the *Development of an HGT-chimera detection pipeline* portion of Results and Discussion and in detail in SI Methods. Colored boxes indicate computational steps described in SI Methods, with numbers outside boxes indicating the number of candidates detected or retained at each phase. Numbers of candidates after the similarity clustering step refer to HGT-chimera similarity clusters. "combined DB" refers to an expanded search database of arthropod proteins from 319 RefSeq genome annotations plus an additional 197 GenBank annotations. The italicized criteria in the bifurcation from the 365 outputs of the TE filter in Phase III indicate that we considered DIAMOND BLASTp hits (E-value < 1 × 10-4) directly instead of hmmsearch results for 79/807 HGT-chimera intervals lacking non-self arthropod hits with bitscore greater than that of the top non-arthropod hit (see SI Methods).

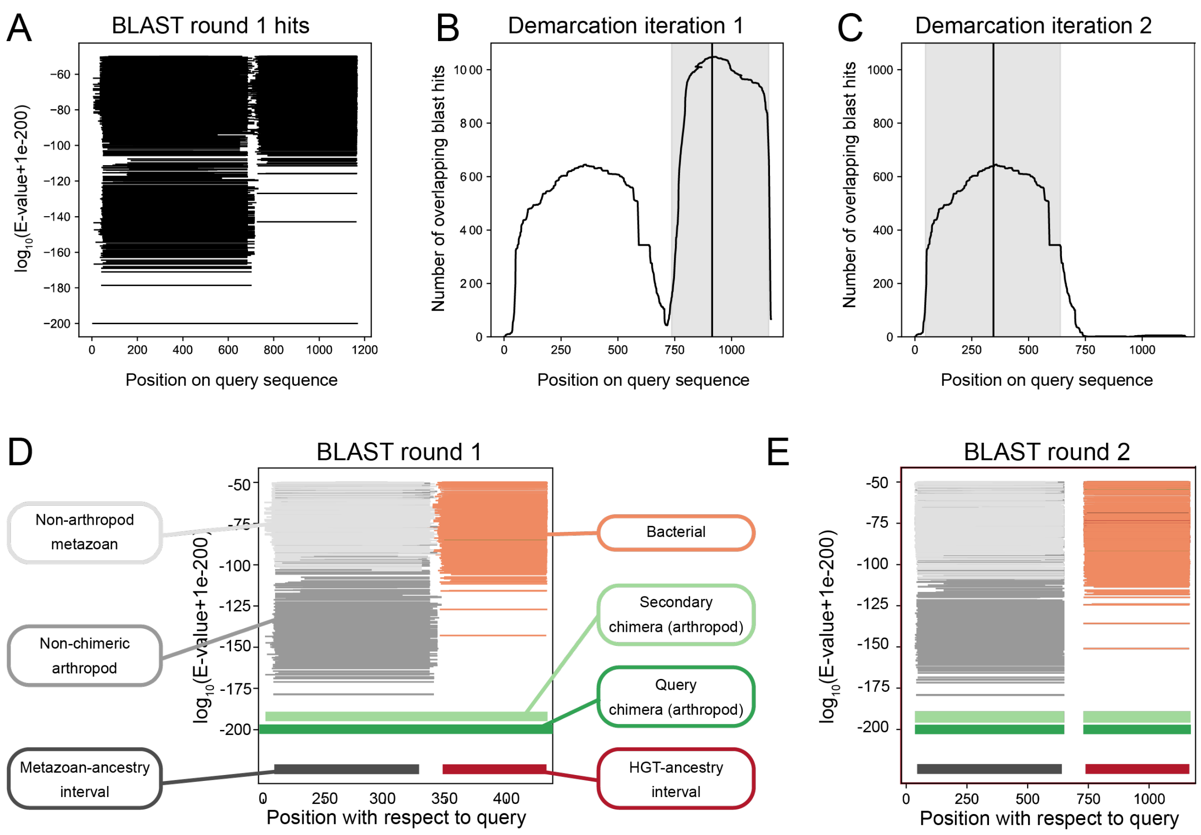


**Fig. S2. Interval demarcation algorithm**. DIAMOND BLASTp hits for a query sequence **(A)** were used to demarcate the query into intervals of possibly distinct evolutionary histories, using an algorithm adapted from Menichelli and colleagues(45). Intervals were determined using the number of BLASTp non-arthropod hits overlapping to each portion of the query. **(B-C)** Each interval was assumed to be centered on a peak in the density of BLASTp hits, and all hits overlapping a peak were assigned to its corresponding interval. Sequences were iteratively added to intervals until fewer than ten sequences remained. In the example shown, two intervals are demarcated corresponding to two peaks in hit density. After assigning putative ancestries to each demarcated interval from the first round of BLASTp **(D)** (see Figure 1A), each interval was subjected to a second round of BLASTp **(E)**. Phylogenetic datasets for each interval were then obtained by constructing a profile HMM from arthropod hits retrieved in the second round of BLASTp and querying it against NR using hmmsearch or directly from BLASTp hits in NR (see SI Methods).

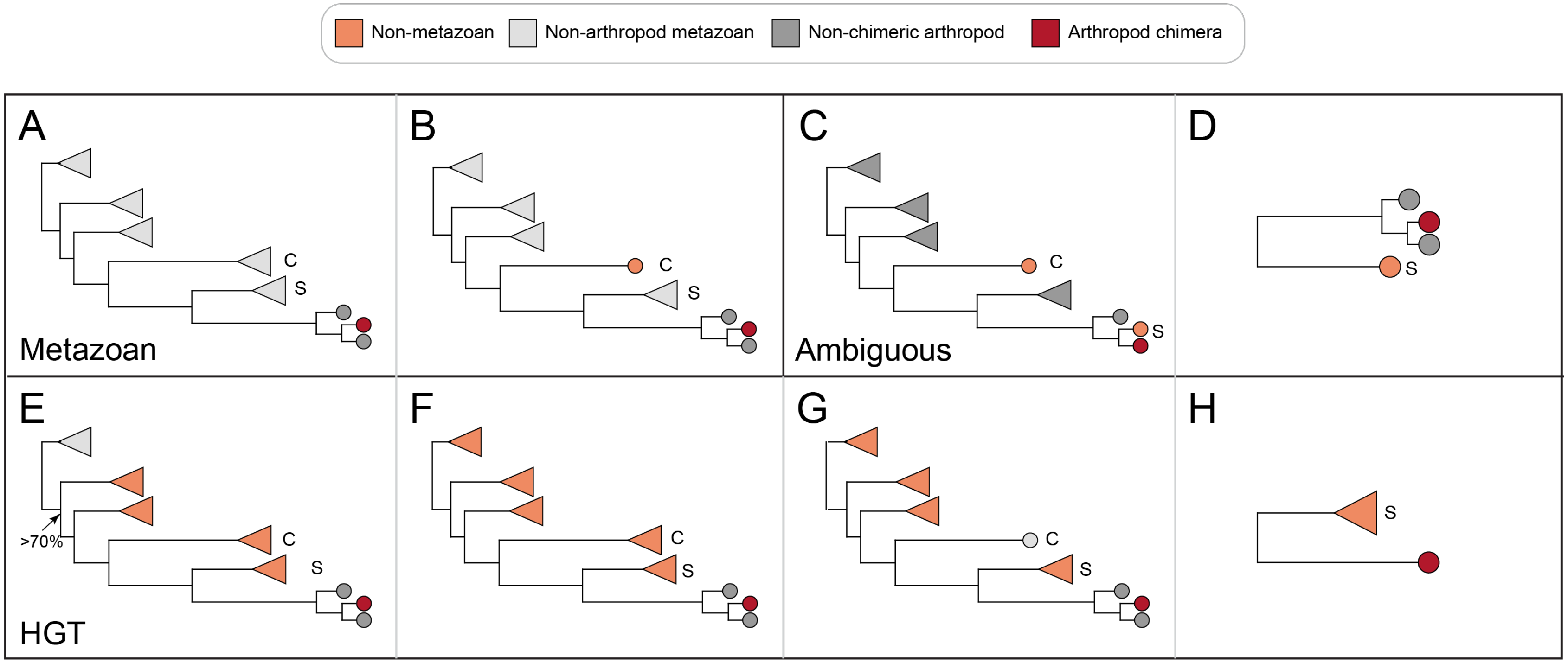


**Fig. S3. Manual tree inspection.** (**A, B**) Schematic illustration of observed types of maximum likelihood tree topologies for HGT-chimera intervals and the resulting ancestry inference. Metazoan: no evidence of HGT, sequence likely of ancient metazoan ancestry. (C, D) Ambiguous: tree topology is inconsistent with either HGT or Metazoan interpretation. **(E-H)** HGT: transfer of non-metazoan sequence to arthropod genome. Triangular leaves represent collapsed clades with more than one sequence and circles represent single sequences. Leaves are colored according to the taxonomic origin of the sequences. S: sister sequence(s); C: cousin sequence(s), which are the closest and second-closest non-arthropod sequences, respectively (see SI Methods). **(A)** Non-arthropod metazoan sister and cousin clades were interpreted as supporting metazoan ancestry. **(B)** Metazoan ancestry was also inferred in cases of sporadic non-metazoan sequences in sister or cousin clades. **(C)** Trees in which sister and cousin taxa were non-metazoan, but non-metazoan sequences were otherwise only sporadically distributed among largely arthropod sequences, were annotated as neither metazoan nor non-metazoan. **(D)** If no cousin clade was available and the non-metazoan clade contained fewer sequences than the metazoan clade, neither a metazoan nor an HGT origin was assigned. **(E)** HGT ancestry was hypothesized when sister and cousin clades were non-metazoan, and the clade containing the HGT-chimera and non-metazoan sequences was separated from non-metazoan clades by at least one node with ultrafast bootstrap support >70% (arrow). **(F)** Bootstrap support was not considered in HGT inference when no non-arthropod metazoan clades were found in the tree. **(G)** HGT ancestry was also hypothesized in cases of sporadic non-arthropod metazoan sequences in sister or cousin clades. **(H)** In the absence of a cousin clade, non-metazoan-to-arthropod HGT was hypothesized when the non-metazoan clade contained more sequences than the metazoan clade, and the non-metazoan clade contained sequences from multiple phyla.

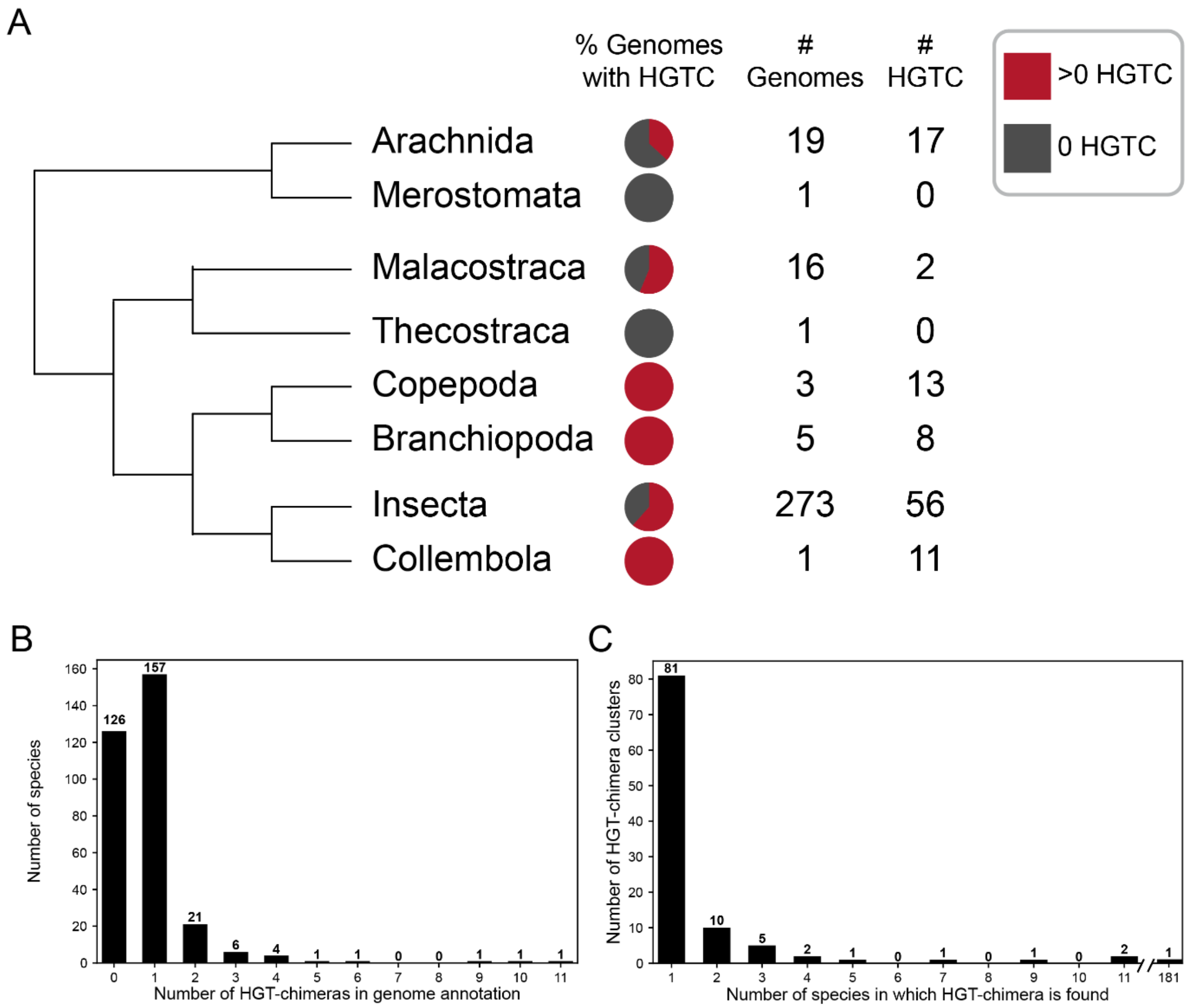


**Fig. S4. HGT-chimeras are detected across Arthropoda. (A)** The results of HGT-chimera search across 319 RefSeq genome annotation datasets by taxonomic class, shown on a phylogram (arbitrary branch lengths) constructed according to published phylogenetic relationships (103). "# Genomes" refers to the number of RefSeq genomes searched per taxonomic class (SI Table 1). "#HGTC" refers to the number of distinct HGT-chimeras found in each class after similarity clustering to a total of 104 similarity clusters. The sum of "# HGT C" across classes is 107 because cluster 3 is found in Collembola, Arachnida, and Insecta, and cluster 18 is found in Arachnida and Insecta. Note that the class "Hexanauplia" in SI Table 1, derived from the NCBI taxonomy database, is now outdated (104) and was substituted for "Copepoda" here. **(B)** Distribution of the number of HGT-chimeras detected per genome for the 319 RefSeq reference genome annotations in the primary search dataset, with a maximum of one member of each HGT-chimera similarity cluster tabulated per species (total N=274 genes). **(C)** The number of species possessing each of 104 HGT-chimera similarity clusters, including both RefSeq and GenBank genomes (source data in SI Table 4). Species range is examined by inspecting the species distribution of arthropod proteins that are hmmsearch or BLASTp hits to both the HGT and metazoan intervals of the representative HGT-chimeras (see SI Methods).

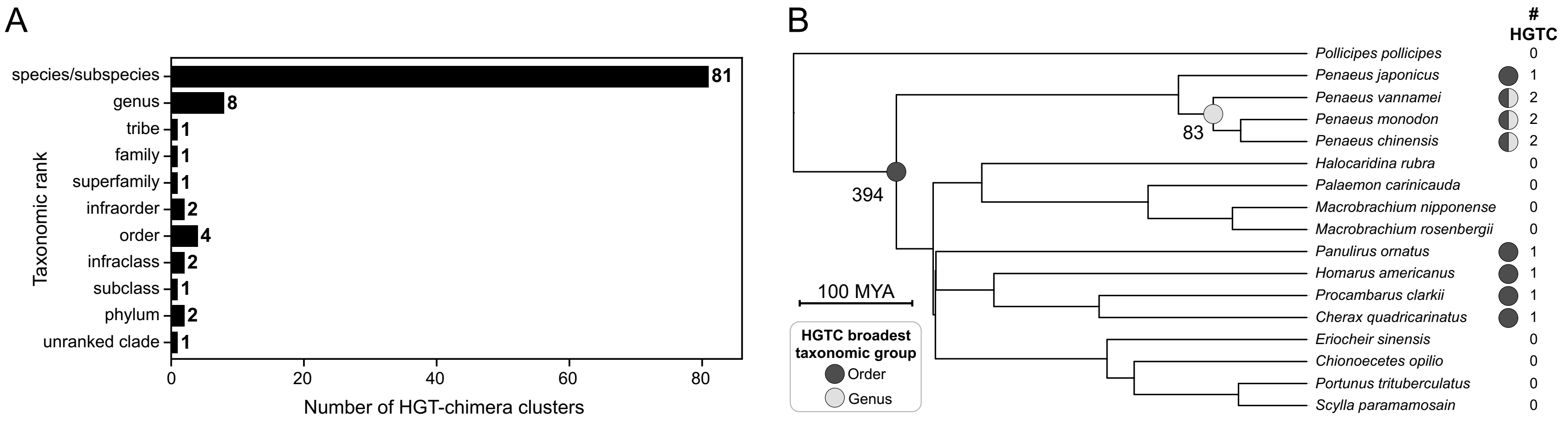


**Fig. S5. Taxonomic range of HGT-chimeras**. **(A)** The lowest taxonomic group encompassing all species containing each HGT-chimera cluster. **(B)** Example phylostratigraphic dating of HGT-chimeras for the order Decapoda. Time-resolved decapod tree obtained using TimeTree (105) with 16 decapod species in our full set of RefSeq and GenBank genomes and one thecostracan (*Pollicipes pollicipes*) included as an outgroup (103). We note that three decapod species (*Penaeus indicus*, *Petrolisthes cinctipes*, *Petrolisthes manimaculis*) found to have representatives of the HGT-chimera with an "Order" label (cluster ID = 2) were omitted from this phylogeny because TimeTree (105) lacked data for these species at the time of writing. Internal nodes labeled with circles correspond to the inferred earliest possible origination of each of two decapod HGT-chimera clusters. Circles and numbers to the right of species names indicate the proportion and number of HGT-chimeras found in each species' genome, colored to match their node of origin. The HGT-chimera with an order label (cluster ID = 2; dark grey) is found in 50% (8/16) of decapod species in this phylogeny, and is hypothesized to have originated in their last common ancestor (394 MYA). The HGT-chimera with a "genus" label (cluster ID = 12; light grey) is found in *Penaeus monodon*, *Penaeus vannamei*, and *Penaeus chinensis* and is hypothesized to have evolved in their last common ancestor (83 MYA).

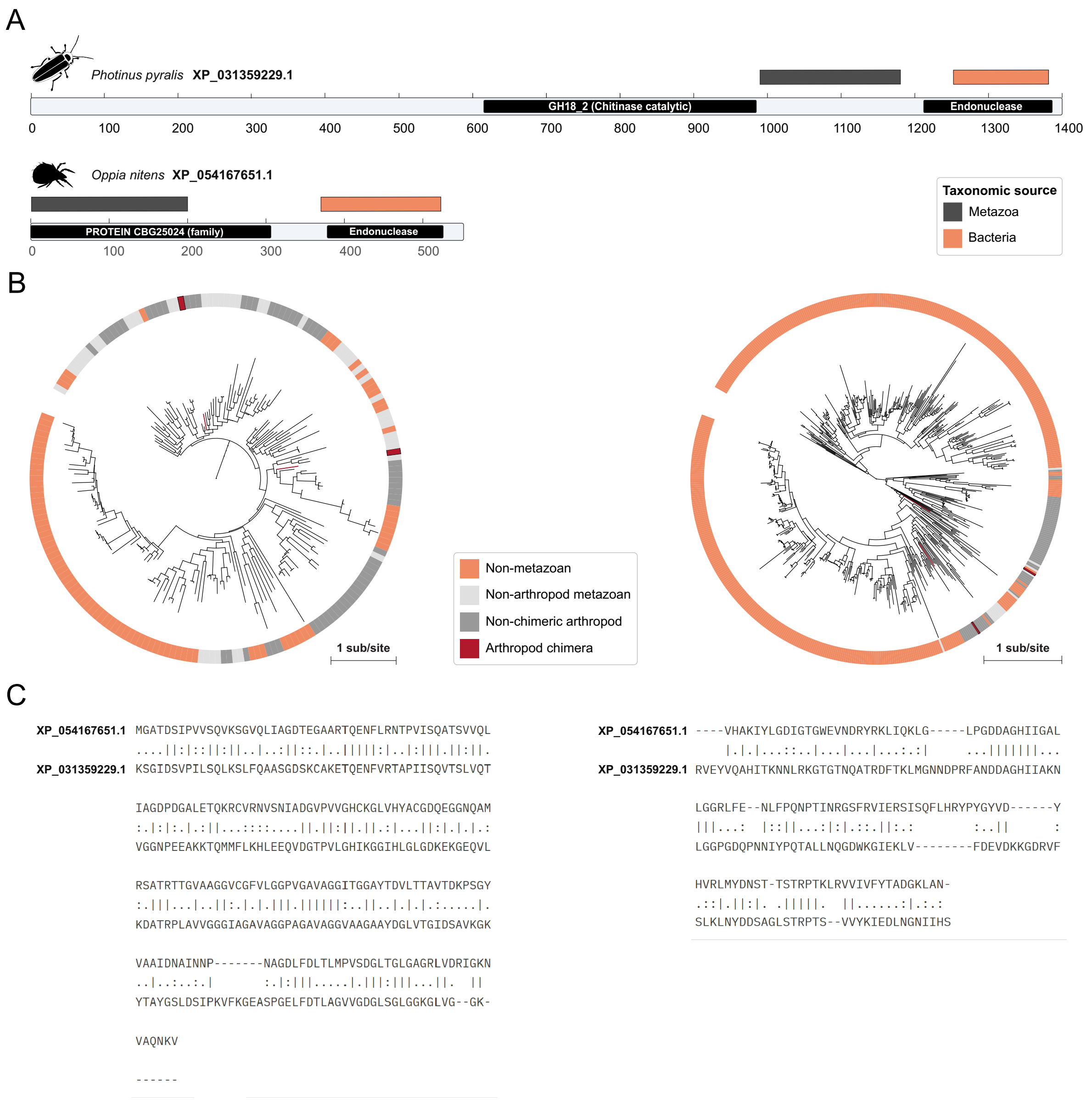

A
Photinus pyralis XP_031359229.1
GH18_2 (Chitinase catalytic)
Endonuclease
0
100
200
300
400
500
600
700
800
900
1000
1100
1200
1300
1400
Oppia nitens XP_054167651.1
PROTEIN CBG25024 (family)
Endonuclease
0
100
200
300
400
500
Taxonomic source
Metazoa
Bacteria
B
1 sub/site
Non-metazoan
Non-arthropod metazoan
Non-chimeric arthropod
Arthropod chimera
1 sub/site
C
XP_054167651.1 MGATDSIPVVSQVKSGVQLIAGDTEGAARTQENFLRNTPVISQATSVVQL
....||:|::||:||..|..:||::.|..|||||:|..|:|||.||:||.
XP_031359229.1 KSGIDSVPILSQLKSLFQAASGDSKCAKETQENFVRTAPIISQVTSLVQT
IAGDPDGALETQKRCVRNVSNIADGVPVVGHCKGLVHYACGDQEGGNQAM
:.|:|:.|.:||...::::....||.||:||.||.:|...||:|.|.|.:
VGGNPEEAKKTQMMFLKHLEEQVDGTPVLGHIKGGIHLGLGDKEKGEQVL
RSATRTTGVAAGGVCGFVLGGPVGAVAGGITGGAYTDVLTTAVTDKPSGY
:.|||...|..||:.|.|.|||.||||||:..||..|.|.|.:.....|.
KDATRPLAVVGGGIAGAVAGGPAGAVAGGVAAGAAYDGLVTGIDSAVKGK
VAAIDNAINNP-------NAGDLFDLTLMPVSDGLTGLGAGRLVDRIGKN
..|..:..:.|       :.|:|||.....|.|||:|||...||.  ||
YTAYGSLDSIPKVFKGEASPGELFDTLAGVVGDGLSGLGGKGLVG--GK-
VAQNKV
------
XP_054167651.1 ----VHAKIYLGDIGTGWEVNDRYRKLIQKLG-----LPGDDAGHIIGAL
|.|.|...::..|...|...|...:.:|     ...|||||||...
XP_031359229.1 RVEYVQAHITKNNLRKGTGTNQATRDFTKLMGNNDPRFANDDAGHIIAKN
LGGRLFE--NLFPQNPTINRGSFRVIERSISQFLHRYPYGYVD------Y
|||...:  |::||...:|:|.::.||:.:        :..||      :
LGGPGDQPNNIYPQTALLNQGDWKGIEKLV--------FDEVDKKGDRVF
HVRLMYDNST-TSTRPTKLRVVIVFYTADGKLAN-
.::|.||:|. .|||||.  ||......:|.:.:
SLKLNYDDSAGLSTRPTS--VVYKIEDLNGNIIHS

**Fig. S6 (previous page). Potentially convergent originations of HGT-chimeras in cluster 18. (A)** InterPro (44) domain architectures of the two protein sequences in HGT-chimera cluster 18, one from the firefly *Photinus pyralis* (order: Coleoptera, class: Insecta) and the other from the mite *Oppia nitens* (order: Sarcoptiformes, class: Arachnida). HGT and metazoan-derived intervals are indicated by orange and dark grey colored rectangles respectively. HGT-derived intervals in both proteins return an endonuclease annotation. Consistent with the hypothesis of independent origins, we note that outside of the HGT and metazoan-derived intervals, the sequence from *P. pyralis* (top) has a long unannotated N-terminal region and a chitinase region not found in the sequence from *O. nitens* (bottom). **(B)** Maximum-likelihood amino acid phylogenetic trees of the metazoan-derived (left) and HGT-derived (right) intervals for HGT-chimera cluster 18. Colored wedges outside the tree indicate the taxonomic origin of each sequence. We note that, in both trees, the HGT- and metazoan-derived intervals (red) do not group in a single clade, consistent with the hypothesis of independent fusion events leading to analogous domain architectures in *P. pyralis* and *O. nitens*. Trees have been pruned and leaf labels omitted for clarity, but full tree topologies are available in the iTOL project "Arthropod HGT-chimera interval trees 8/27/2025" (https://itol.embl.de/shared/rkapoor). Scale bars indicate one substitution per site. **(C)** Pairwise global alignments via the EMBOSS Needle tool (https://www.ebi.ac.uk/jdispatcher/psa/emboss_needle) for the HGT- and metazoan-derived intervals of the two proteins in this alignment. Vertical lines show identical amino acids, while two dots indicate substitution for amino acids with similar physiochemical properties. Although the metazoan-derived regions of the two proteins (left) do not return the same InterPro domain, they align with 40.8% identity and 55.8% similarity,

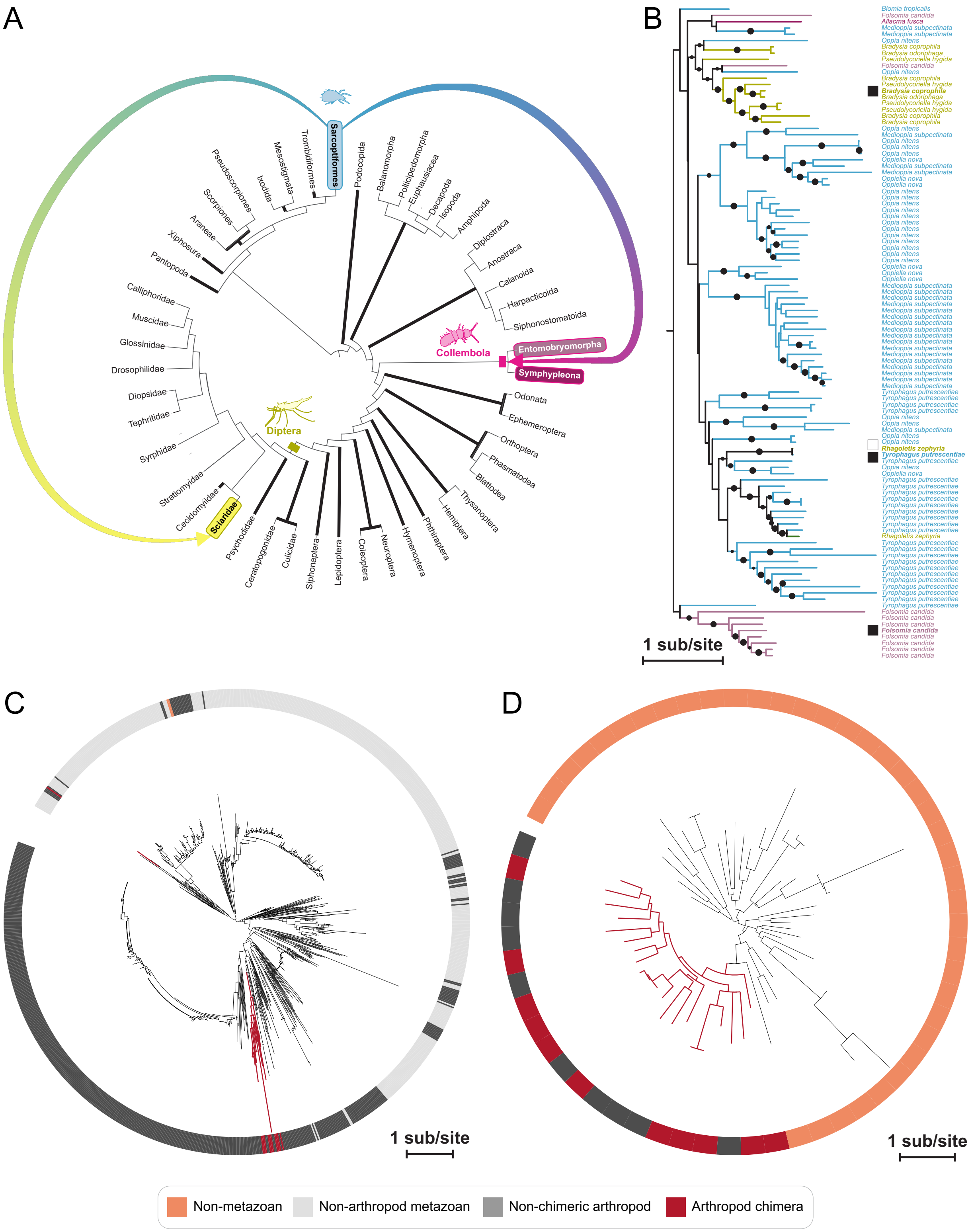
A
B
C
D
Sarcoptiformes
Trombidiformes
Mesostigmata
Ixodida
Pseudoscorpiones
Scorpiones
Araneae
Xiphosura
Pantopoda
Podocopida
Balanomorpha
Pollicipedomorpha
Euphausiacea
Decapoda
Isopoda
Amphipoda
Diplostraca
Anostraca
Calanoida
Harpacticoida
Siphonostomatoida
Collembola
Entomobryomorpha
Symphypleona
Odonata
Ephemeroptera
Orthoptera
Phasmatodea
Blattodea
Thysanoptera
Hemiptera
Phthiraptera
Hymenoptera
Neuroptera
Coleoptera
Lepidoptera
Siphonaptera
Culicidae
Ceratopogonidae
Psychodidae
Sciaridae
Cecidomyiidae
Stratiomyidae
Syrphidae
Tephritidae
Diopsidae
Drosophilidae
Glossinidae
Muscidae
Calliphoridae
Diptera
Blomia tropicalis
Folsomia candida
Allacma fusca
Medioppia subpectinata
Oppia nitens
Bradysia coprophila
Bradysia odoriphaga
Pseudolycoriella hygida
Oppiella nova
Tyrophagus putrescentiae
Rhagoletis zephyria
1 sub/site
1 sub/site
1 sub/site
Non-metazoan
Non-arthropod metazoan
Non-chimeric arthropod
Arthropod chimera

**Fig. S7 (previous page) Potential post-formation inter-arthropod HGT of cluster 3. (A)** The phylogenetic distribution of HGT-chimera similarity cluster 3 mapped onto a phylogram depicting relationships among the taxa in which the cluster is found. The phylogram includes all orders represented in our combined database of RefSeq and GenBank genomes (SI Table 1), and was obtained via concatenation of published relationships for hexapods (106), chelicerates (107), non-hexapod Pancrustacea (103), and Diptera (108). Leaf nodes indicate taxonomic orders except for Diptera, which was further expanded to include families as necessary to show the distribution of the HGT-chimera. Taxa in which the chimera was found are highlighted with a unique color per taxon. Strong black internal branches indicate a hypothesis of independent loss, under the most parsimonious gene history in which the chimera was found in the last common ancestor of all lineages in the tree followed by repeated losses with no HGT, for a total minimum number of losses of N=19 under this vertical descent hypothesis. In contrast, the external arrows indicate a more parsimonious series of two horizontal HGT events following an origination in Sarcoptiformes (supported by SI Figure 7B). **(B)** Maximum likelihood amino acid phylogenetic tree for the 104 HGT chimeras across 11 species in cluster 3, limited to at most one isoform per gene. Multiple sequence alignments of protein sequences were generated by MUSCLE and trimmed to 90% column occupancy with trimAl. The maximum likelihood tree was generated using IQ-TREE with automated model selection and 1000 iterations of ultrafast bootstrapping (bootstrap support indicated by internal node support values). Tree is arbitrarily rooted for ease of visualization. Sequence names are colored to correspond to taxa in (A). Black boxes to the left of species names indicate sequences validated via RT-PCR and Sanger sequencing. The white box indicates that RT-PCR was attempted but failed to produce any bands. Scale bar represents 1 amino acid substitution per site. Dots at internal nodes indicate bootstrap support >70%. **(C)** Maximum likelihood tree for the metazoan interval produced for chimera 3, constructed with at most one chimera representative per genome as for metazoan/HGT ancestry inference (see SI Methods). Colors represent the taxonomic origin of the sequence intervals. Note that all but one of the intervals derived from HGT-chimeric sequences are clustered in a clade (red branches) at the bottom of the tree labeled with red branches (100% bootstrap support). The non-chimeric sequences within this clade are all from the same taxonomic groups as the chimeric sequence (Collembola, Sciaridae, Sarcoptiformes), consistent with a secondary loss of the HGT. The remaining chimeric sequence groups with non-arthropod metazoan sequences with low bootstrap support (11%), consistent with the hypothesis that this apparent non-monophyly is artifactual. Scale bar represents 1 amino acid substitution per site. **(D)** Maximum likelihood tree for the HGT interval produced for chimera 3, constructed with at most one chimera representative per genome as for metazoan/HGT ancestry inference (see SI Methods). Note that all arthropod chimeric and non-chimeric intervals form a single clade (red branches) to the exclusion of non-metazoan (bacterial and viral) sequences, consistent with the hypothesis that all HGT-chimera intervals descend from a single non-metazoan transfer event. As in (C), the non-chimeric arthropod sequences in the red clade are all from Collembola, Sarcoptiformes, and Sciaridae.

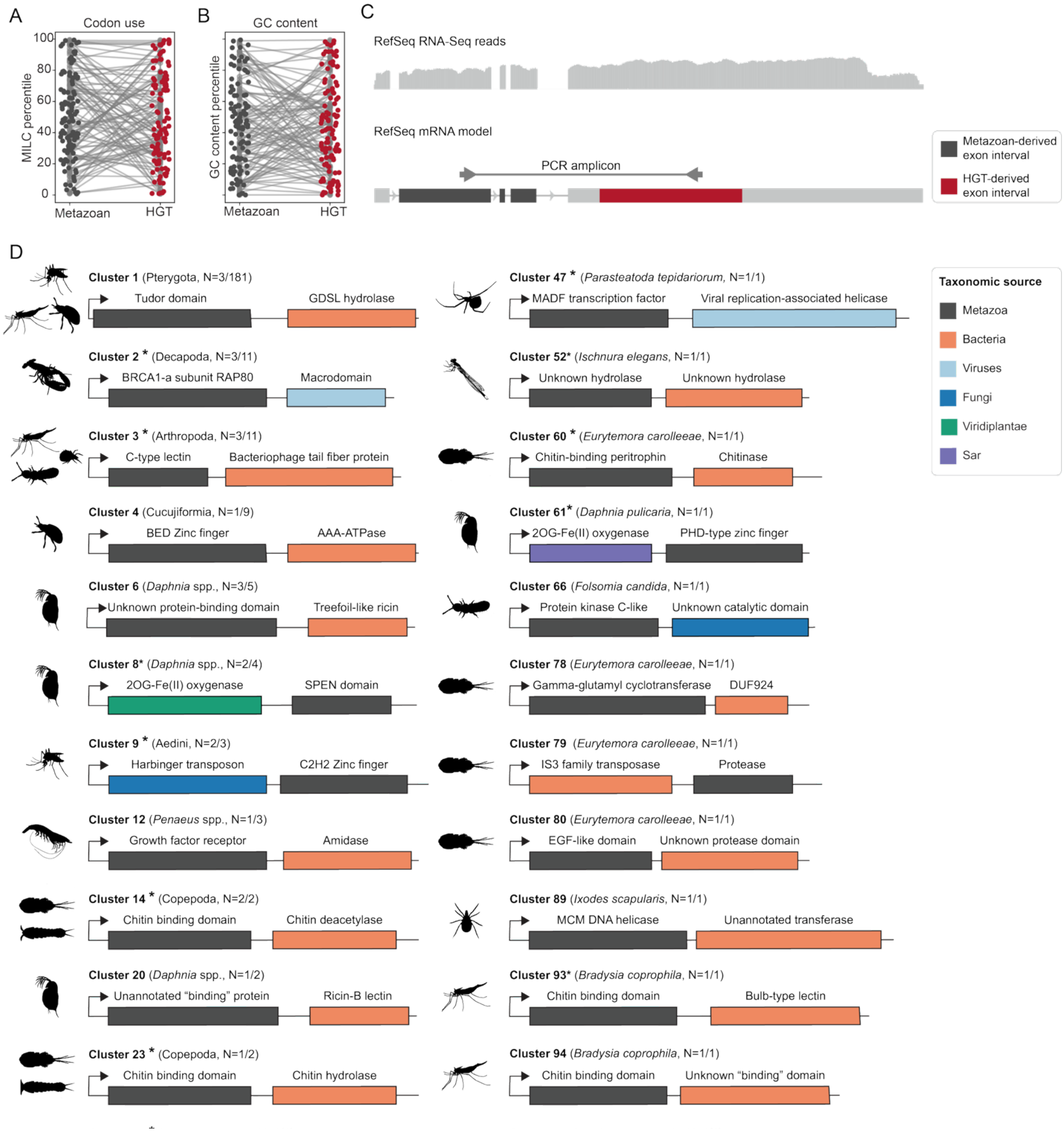
A
Codon use
MILC percentile
100
80
60
40
20
0
Metazoan
HGT
B
GC content
GC content percentile
Metazoan
HGT
C
RefSeq RNA-Seq reads
RefSeq mRNA model
PCR amplicon
Metazoan-derived exon interval
HGT-derived exon interval
D
Cluster 1 (Pterygota, N=3/181)
Tudor domain
GDSL hydrolase
Cluster 2 * (Decapoda, N=3/11)
BRCA1-a subunit RAP80
Macrodomain
Cluster 3 * (Arthropoda, N=3/11)
C-type lectin
Bacteriophage tail fiber protein
Cluster 4 (Cucujiformia, N=1/9)
BED Zinc finger
AAA-ATPase
Cluster 6 (Daphnia spp., N=3/5)
Unknown protein-binding domain
Treefoil-like ricin
Cluster 8* (Daphnia spp., N=2/4)
2OG-Fe(II) oxygenase
SPEN domain
Cluster 9 * (Aedini, N=2/3)
Harbinger transposon
C2H2 Zinc finger
Cluster 12 (Penaeus spp., N=1/3)
Growth factor receptor
Amidase
Cluster 14 * (Copepoda, N=2/2)
Chitin binding domain
Chitin deacetylase
Cluster 20 (Daphnia spp., N=1/2)
Unannotated "binding" protein
Ricin-B lectin
Cluster 23 * (Copepoda, N=1/2)
Chitin binding domain
Chitin hydrolase
Cluster 47 * (Parasteatoda tepidariorum, N=1/1)
MADF transcription factor
Viral replication-associated helicase
Cluster 52* (Ischnura elegans, N=1/1)
Unknown hydrolase
Unknown hydrolase
Cluster 60 * (Eurytemora carolleeae, N=1/1)
Chitin-binding peritrophin
Chitinase
Cluster 61* (Daphnia pulicaria, N=1/1)
2OG-Fe(II) oxygenase
PHD-type zinc finger
Cluster 66 (Folsomia candida, N=1/1)
Protein kinase C-like
Unknown catalytic domain
Cluster 78 (Eurytemora carolleeae, N=1/1)
Gamma-glutamyl cyclotransferase
DUF924
Cluster 79 (Eurytemora carolleeae, N=1/1)
IS3 family transposase
Protease
Cluster 80 (Eurytemora carolleeae, N=1/1)
EGF-like domain
Unknown protease domain
Cluster 89 (Ixodes scapularis, N=1/1)
MCM DNA helicase
Unannotated transferase
Cluster 93* (Bradysia coprophila, N=1/1)
Chitin binding domain
Bulb-type lectin
Cluster 94 (Bradysia coprophila, N=1/1)
Chitin binding domain
Unknown "binding" domain
Taxonomic source
Metazoa
Bacteria
Viruses
Fungi
Viridiplantae
Sar

**Fig. S8 (previous page). HGT-chimeras are transcribed. (A)** Measure Independent of Length and Composition (MILC) is a measure of codon use bias (109). Higher MILC values indicate greater deviation from genome-wide expected codon use frequencies. To enable comparison of codon use between HGT and metazoan intervals from the same protein as well as comparison to non-chimeric transcripts, we percentile normalized MILC values with respect to a distribution obtained via random sampling of annotated coding sequences from the same genome as each HGT-chimera. This plot shows percentile-normalized MILC across HGT (red) and metazoan-derived (dark grey) intervals across all 104 representative chimeras, with light grey lines connecting HGT and metazoan intervals from the same chimera (see SI Methods). Percentile-normalized MILC values were roughly uniformly distributed over [0,1], as would be expected for any genomic interval. HGT and metazoan intervals did not differ significantly in their percentile-normalized MILC values (Wilcoxon rank-sum p-value > 0.10). **(B)** GC values by interval type, with percentile scores produced as in (A). HGT and metazoan intervals do not differ significantly in their percentile-normalized GC values (Wilcoxon rank-sum p-value > 0.10). Collectively, A and B show that HGT-ancestry intervals do not systematically deviate in codon use or GC content from the background of expressed transcripts from the same genome, nor from metazoan intervals in the same protein. Source data for A-B are available in SI Table 11. **(C)** Schematic showing evidence of transcription for HGT-chimeras. Top: First, for all HGT-chimeras, RefSeq mRNA models and supporting aggregate RNASeq read data were visualized using the NCBI genome data viewer. Sequences were checked to determine whether RNASeq exon-spanning reads mapped to both HGT and metazoan-annotated regions of the mRNA transcript. 62.5% (65/104) of HGT-chimeras had RNA-Seq support over the full length of their transcripts in the RefSeq genome browser, while 89.4% (93/104) of HGT-chimeras had RNA-seq support over >50% of the length of all identified HGT and metazoan intervals (SI Table 12). Bottom: Next, for a subset of 41 HGT-chimeras, RT-PCR and Sanger sequencing were performed using primer pairs that produce amplicons spanning the junction between HGT and metazoan regions (SI Table 13). Multiple overlapping amplicons were sequenced and assembled when it was not possible to obtain a single amplicon <1.5 kb in length that spanned the HGT-metazoan junction. **(D)** Predicted domain architectures via NCBI CDD (53), InterProScan (44), DeepGO-SE (36), DIAMOND BLASTp, and CENSOR for 24 HGT-chimeras with expression validated by RT-PCR and Sanger sequencing (SI Table 13). The unique cluster identifier of each HGT-chimera is indicated, along with its taxonomic range in parenthesis. Asterisks indicate a predicted functional commonality between the metazoan and HGT intervals (SI Table 18). Note that cluster 3 may have undergone inter-arthropod HGT (SI Text 1). Representative silhouettes obtained from phylopic.org, or generated by Isobel Ronai for *Ixodes scapularis*. Fractions next to taxon names indicate the number of total species in the cluster for which we confirmed chimera expression via RT-PCR (SI Table 13). Domain length is arbitrary and not to scale.

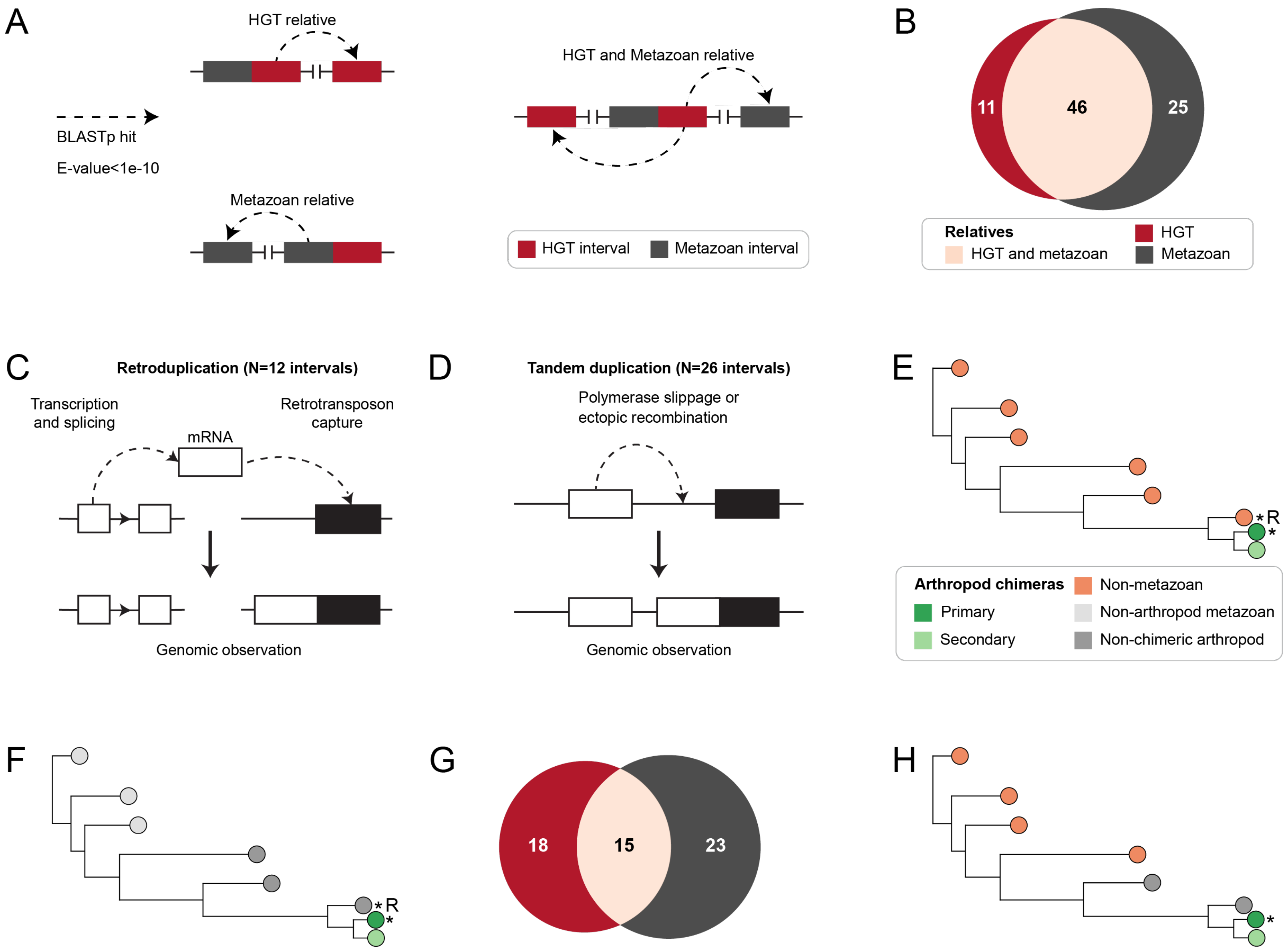
A
HGT relative
BLASTp hit
E-value<1e-10
Metazoan relative
HGT and Metazoan relative
HGT interval
Metazoan interval
B
11
46
25
Relatives
HGT and metazoan
HGT
Metazoan
C
Retroduplication (N=12 intervals)
Transcription and splicing
mRNA
Retrotransposon capture
Genomic observation
D
Tandem duplication (N=26 intervals)
Polymerase slippage or ectopic recombination
Genomic observation
E
*R
*
Arthropod chimeras
Primary
Secondary
Non-metazoan
Non-arthropod metazoan
Non-chimeric arthropod
F
*R
*
G
18
15
23
H
*

**Fig. S9 (previous page). HGT-chimera relative detection. (A)** Illustration of DIAMOND BLASTp-based relative detection, using separated HGT- and metazoan intervals as queries against proteins from other genes in the same genome as the reference chimera. **(B)** The number of HGT-chimeras with non-chimeric parents (DIAMOND BLASTp E-value <1e-10) detected at a different locus within the same genome, for HGT intervals, metazoan intervals, or both. Venn diagram includes data for the 82/104 HGT-chimeras for which we found evidence of a relative of at least one constituent interval. **(C)** HGT-chimera intervals were hypothesized to be products of retroduplication if the interval was spanned by one exon in the HGT-chimera and more than one exon in the BLASTp-defined relative (right) or **(D)** products of tandem duplication if HGT-chimeras were located on the same chromosome or genomic scaffold as their BLASTp-defined relatives with ≤ 2 intervening genes (left). **(E)** Illustration of tree-based relative inference for an HGT-origin interval. Asterisks indicate sequences from the same species as the primary interval. R: relatives hypothesized as the closest non-chimera phylogenetic relative of the primary HGT-chimera, if the sequence is from the same species as the primary HGT-chimera. Note that relatives are hypothesized even when they are part of a multi-sequence neighboring clade (not shown). **(F)** Illustration of tree-based parent inference for a metazoan interval. **(G)** Venn diagram shows HGT (red) and metazoan (grey) relatives detected for a total of 56/104 chimeras with at least one detected relative. (**H)** Illustration of phylogenetic search for non-chimeric HGT-chimera relatives in arthropod species that lack the HGT-chimera, returning the dark grey sequences as relatives.

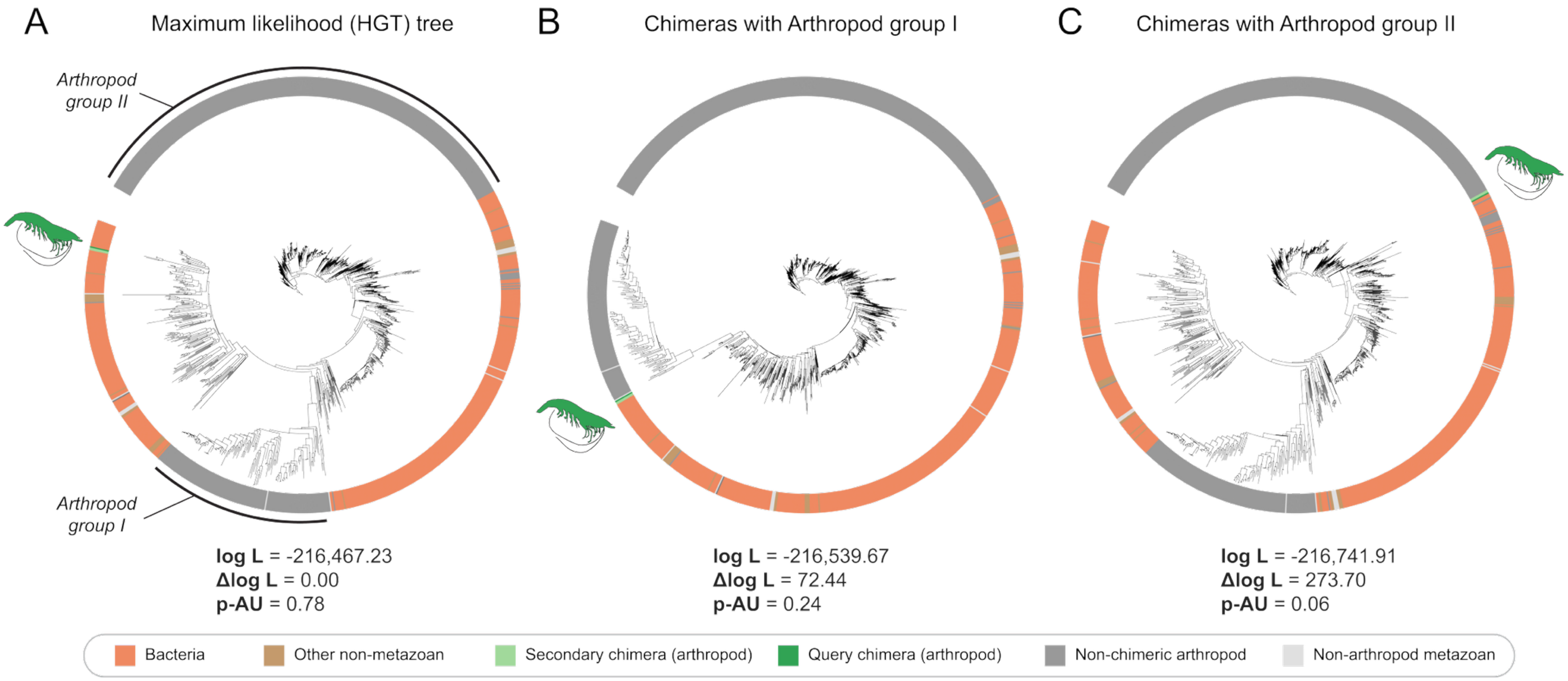


**Fig. S10. Tree topology tests support HGT inference for *Penaeus* HGT-chimera cluster 12. (A)** Full maximum-likelihood topology (unrooted) for the HGT interval of HGT-chimera cluster 12, placing *Penaeus* chimera sequences within a large clade of bacterial sequences. Two additional, distantly related arthropod clades (arthropod groups I and II) possibly represent endogenous (non-HGT) amidases. To assess alternative placements of the *Penaeus* HGT-chimeras, we imposed constraints forcing them to group with arthropod group I **(B)** or arthropod group II **(C)**, to the exclusion of the remaining sequences. Topology support was evaluated with the approximately unbiased (AU) test. The ML (HGT-supporting) topology received the highest support. Constraining *Penaeus* HGT-chimeras with arthropod group II was rejected (p-AU < 0.05), whereas constraining them with arthropod group I had a lower likelihood than the ML tree but was not rejected (p-AU = 0.24). Notably, under the group-I constraint the *Penaeus* HGT-chimeras are placed as the outgroup to all other arthropod sequences in this clade. We view this as biologically implausible given the presence of early-diverging pancrustacean and arachnid lineages within group I. We instead hypothesize that this placement reflects the strong affinity of the *Penaeus* HGT-chimeras to bacterial sequences.

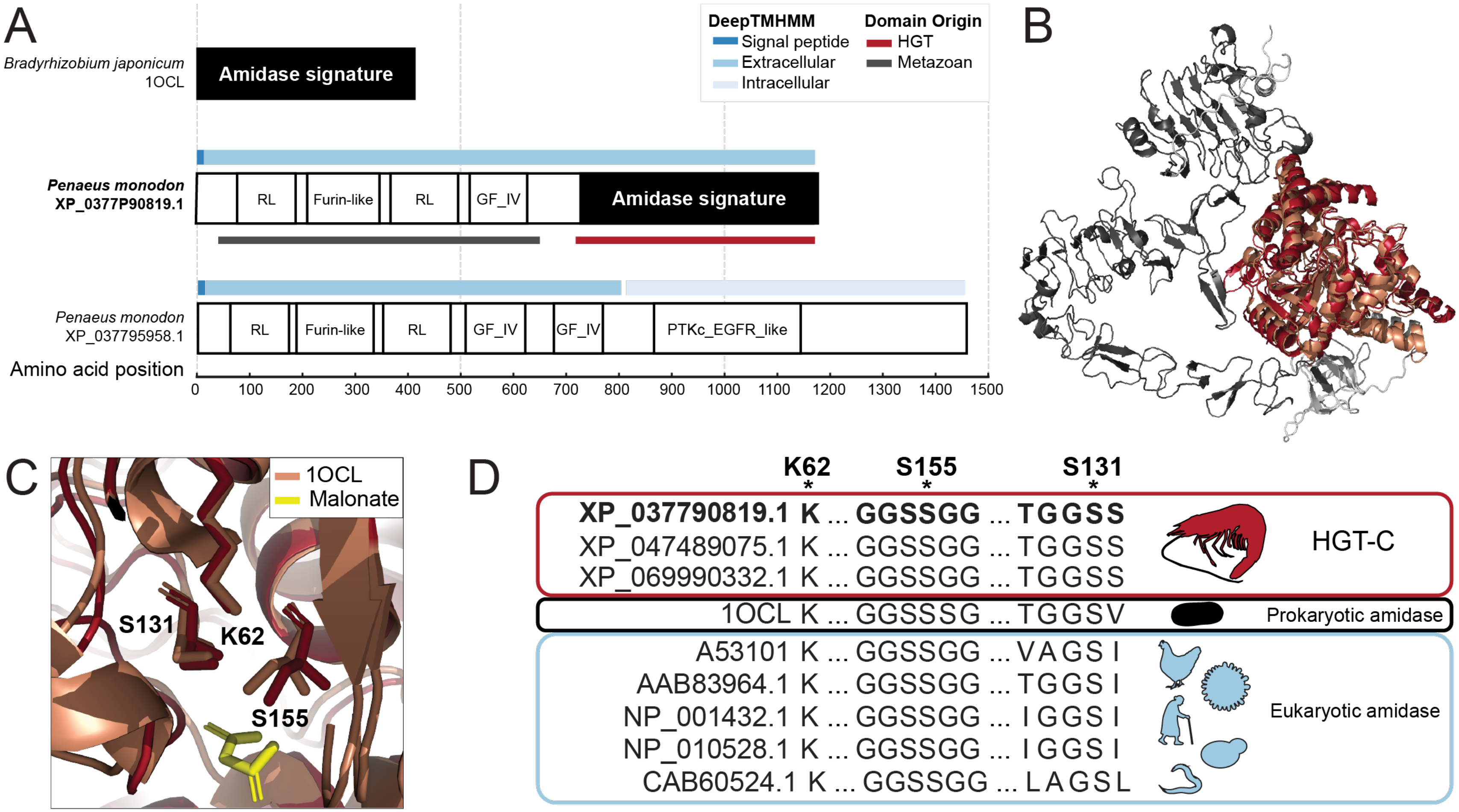
A
Bradyrhizobium japonicum
1OCL
Amidase signature
DeepTMHMM
Signal peptide
Extracellular
Intracellular
Domain Origin
HGT
Metazoan
Penaeus monodon
XP_0377P90819.1
RL
Furin-like
RL
GF_IV
Amidase signature
Penaeus monodon
XP_037795958.1
RL
Furin-like
RL
GF_IV
GF_IV
PTKc_EGFR_like
Amino acid position
0 100 200 300 400 500 600 700 800 900 1000 1100 1200 1300 1400 1500
B
C
1OCL
Malonate
S131
K62
S155
D
K62 S155 S131
XP_037790819.1 K ... GGSSGG ... TGGSS
XP_047489075.1 K ... GGSSGG ... TGGSS
XP_069990332.1 K ... GGSSGG ... TGGSS
HGT-C
1OCL K ... GGSSSG ... TGGSV
Prokaryotic amidase
A53101 K ... GGSSGG ... VAGSI
AAB83964.1 K ... GGSSGG ... TGGSI
NP_001432.1 K ... GGSSGG ... IGGSI
NP_010528.1 K ... GGSSGG ... IGGSI
CAB60524.1 K ... GGSSGG ... LAGSL
Eukaryotic amidase

**Figure S11 (previous page). Evidence of amidase catalytic activity in the HGT-derived interval of HGT-chimera cluster 12 from *Penaeus spp.*** **(A)** Domain architecture (inferred by searching the NCBI Conserved Domain Database (CDD) using CD-Search (52, 53), and  (44) Scan) for the representative HGT-chimera with NCBI accession XP_037790819.1 from *P. monodon*, its closest non-chimeric metazoan phylogenetic relative in the same genome (epidermal growth factor receptor-like XP_037795958.1; NCBI BLASTp E-value: 0), and a bacterial amidase (Protein DataBase (PDB) accession 1OCL) with a solved crystal structure and experimentally validated biochemical activity that aligns with the HGT-interval (NCBI BLASTp E-value: 6E-18) (110). Annotated domains include Recep L (NCBI CDD Receptor L domain), Furin-like (NCBI CDD Furin-like cysteine rich region), GF_IV (NCBI CDD growth factor receptor domain IV), PTKc_EGFR_like (NCBI CDD catalytic domain of epidermal growth factor receptor-like protein tyrosine kinases), and the amidase signature domain (InterProScan). Predicted ancestries of HGT-chimera intervals are indicated as colored bars below XP_037790819.1. DeepTMHMM (99) membrane topology predictions are indicated as colored lines above XP_037790819.1 and XP_037795958.1. We note that the chimera has preserved the N-terminal signal peptide and predicted extracellular region of its metazoan relative but is predicted to lack any intracellular domains and to be soluble. **(B)** AlphaFold3 (32) structural predictions for *P. monodon* XP_037790819.1 aligned with the crystal structure of *B. japonicum* malonamidase (PDB 1OCL) in complex with substrate analog malonate. 1OCL is shown in orange, malonate is yellow, and XP_037790819.1 is colored according to predicted ancestry as in (A). Light grey indicates protein interval of undefined ancestry, meaning that this interval was assigned to neither metazoan nor HGT ancestry. As predicted from domain architectures (A), the chimera and *B. japonicum* proteins align in the amidase domain (RMSD: 1.741). **(C)** The structural alignment in (B), zoomed in to the previously characterized active site of 1OCL(110) . All previously identified catalytic triad residues in 1OCL (labeled) are conserved in XP_037790819.1. **(D)** MUSCLE multiple sequence alignment of amidase signature family proteins, including HGT intervals from HGT-chimera cluster 12 representatives in three *Penaeus* species (*P. monodon* XP_037790819.1, *P. chinensis* XP_047489075.1, *P. vannamei* XP_069990332.1), *B. japonicum* 1OCL_1, *Bacillus subtilis subtilis* AAB83964.1, *Saccharomyces cerevisiae* NP_010528.1, *Caenorhabditis elegans* CAB60524.1, *Gallus gallus* A53101, *Homo sapiens* NP_001432.1. HGT-chimera sequences contain conserved catalytic triad residues (indicated with asterisks), along with flanking residues that support catalytic activity through orientation of the catalytic residues (dashed line) or formation of an oxyanion hole (solid line) (110).

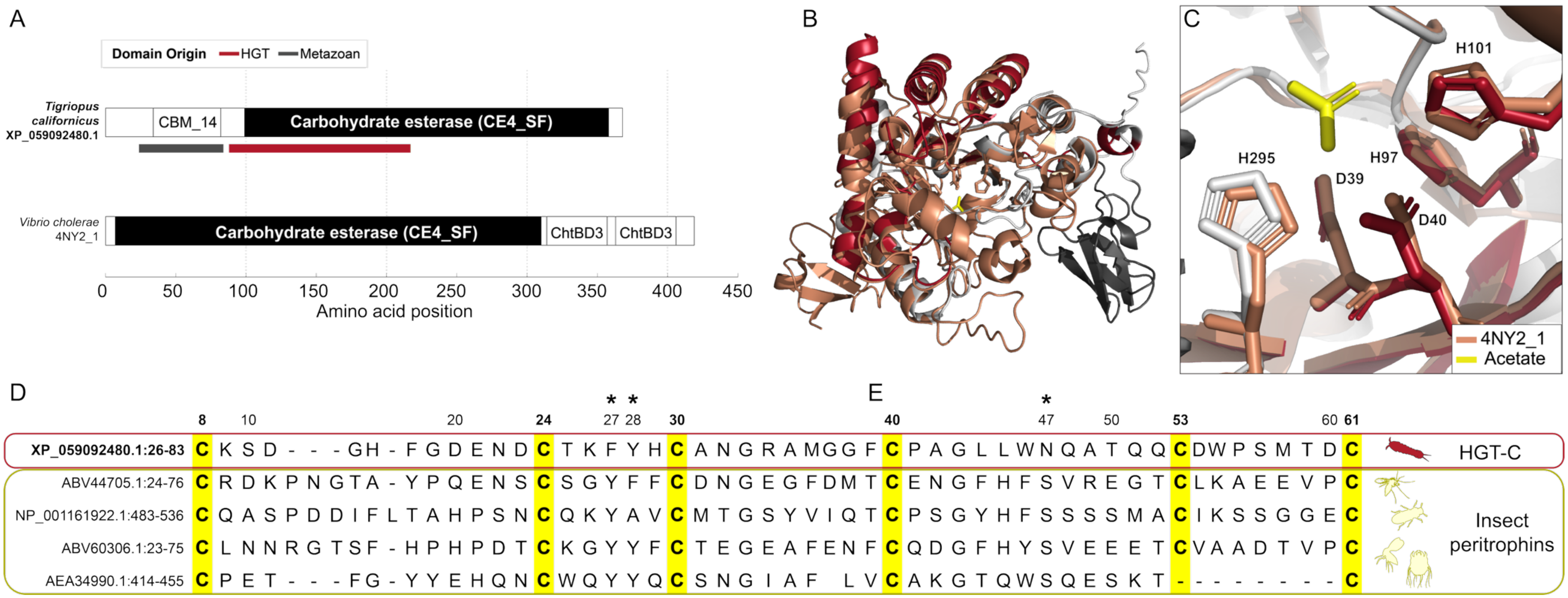
A
Domain Origin
HGT
Metazoan
Tigriopus californicus
XP_059092480.1
CBM_14
Carbohydrate esterase (CE4_SF)
Vibrio cholerae
4NY2_1
Carbohydrate esterase (CE4_SF)
ChtBD3
ChtBD3
0 50 100 150 200 250 300 350 400 450
Amino acid position
B
C
H101
H295
H97
D39
D40
4NY2_1
Acetate
D
E
8 10 20 24 27 28 30 40 47 50 53 60 61
XP_059092480.1:26-83 C K S D - - - G H - F G D E N D C T K F Y H C A N G R A M G G F C P A G L L W N Q A T Q Q C D W P S M T D C
ABV44705.1:24-76 C R D K P N G T A - Y P Q E N S C S G Y F F C D N G E G F D M T C E N G F H F S V R E G T C L K A E E V P C
NP_001161922.1:483-536 C Q A S P D D I F L T A H P S N C Q K Y A V C M T G S Y V I Q T C P S G Y H F S S S S M A C I K S S G G E C
ABV60306.1:23-75 C L N N R G T S F - H P H P D T C K G Y Y F C T E G E A F E N F C Q D G F H Y S V E E E T C V A A D T V P C
AEA34990.1:414-455 C P E T - - - F G - Y Y E H Q N C W Q Y Y Q C S N G I A F L V C A K G T Q W S Q E S K T - - - - - - - - C
HGT-C
Insect peritrophins

**Figure S12 (previous page). Evidence of chitin-interacting functionality in the HGT and metazoan intervals of XP_059092480.1 (cluster 14) from *Tigriopus californicus*. (A)** Domain architecture (inferred by NCBI CD-Search (52, 53)) for the HGT-chimera with NCBI accession XP_059092480.1, and a *Vibrio cholerae* chitin deacetylase with a solved crystal structure and experimentally validated biochemical activity (Protein DataBase (PDB) accession 4NY2) (111). The esterase/deacetylase domain (CE4_Mll8295_like; CES4 superfamily) of the HGT-chimera partially overlaps the inferred HGT interval, and the chitin-binding peritrophin domain (CTBM4) overlaps the metazoan interval. Given that the entire deacetylase/esterase domain of the chimera aligns with the corresponding domain of the *V. cholerae* de-acetylase (NCBI BLASTp E-value 3E-27), we hypothesize that this entire domain is likely of horizontal origin and was excluded by the interval demarcation procedure due to a lower density of BLASTp hits outside of the core catalytic region. We further note that the *V. cholerae* sequence contains two C-terminal chitin-binding domains (ChtBD3) absent from the HGT-chimera, which instead contains an N-terminal metazoan chitin-binding peritrophin domain. **(B)** AlphaFold3 (32) structural predictions for *T. californicus* XP_059092480.1 aligned with the crystal structure of *V. cholerae* chitin deacetylase (PDB 4NY2) in complex with acetate. 4NY2 is shown in orange, acetate is yellow, and XP_059092480.1 is colored according to predicted ancestry as in (A). Light grey indicates protein interval of undefined ancestry, meaning that this interval was assigned to neither metazoan nor HGT ancestry. . As predicted from domain architectures (A), the chimera and *V. cholerae* structures align in their catalytic deacetylase domains (RMSD: 1.486), but not in the N-terminal peritrophin of the chimera or the C-terminal chitin binding domains of the *V. cholerae* protein. Active site residues are shown as sticks. **(C)** The structural alignment in (B), zoomed in to the previously characterized active site of 4NY2, with catalytic residues of 4NY2 labeled (111) . All previously identified catalytic residues in 4NY2 are conserved in XP_059092480.1, including those that coordinate a catalytic zinc ion (H101, D40, H97) and those that participate directly in the acid/base catalytic mechanism (H295, D39) (111) . **(D)** MUSCLE multiple-sequence alignment of the chitin-binding peritrophin domain of *T. californicus* XP_059092480.1 with chitin-biding domains from the arthropod peritrophins ABV44705.1 (*Phlebotomus papatasi*, class: Insecta, order: Diptera), NP_001161922.1 (*Tribolium castaneum*, class: Insecta, order: Coleoptera), ABV60306.1 (*Lutzomyia longipalpis*, class: Insecta, order: Diptera), and AEA34990.1 (*Sarcoptes scabiei*, class: Insecta, order: Sarcoptiformes). We note that the chitin-binding abilities of ABV44705.1 (112) and AE34990.1 (113) were previously experimentally validated. The peritrophin domain of the chimera contains all six conserved disulfide bridge-forming cysteine residues (shown in yellow) as well as all aromatic residues predicted to bind chitin monomers (designated with asterisks) (112).

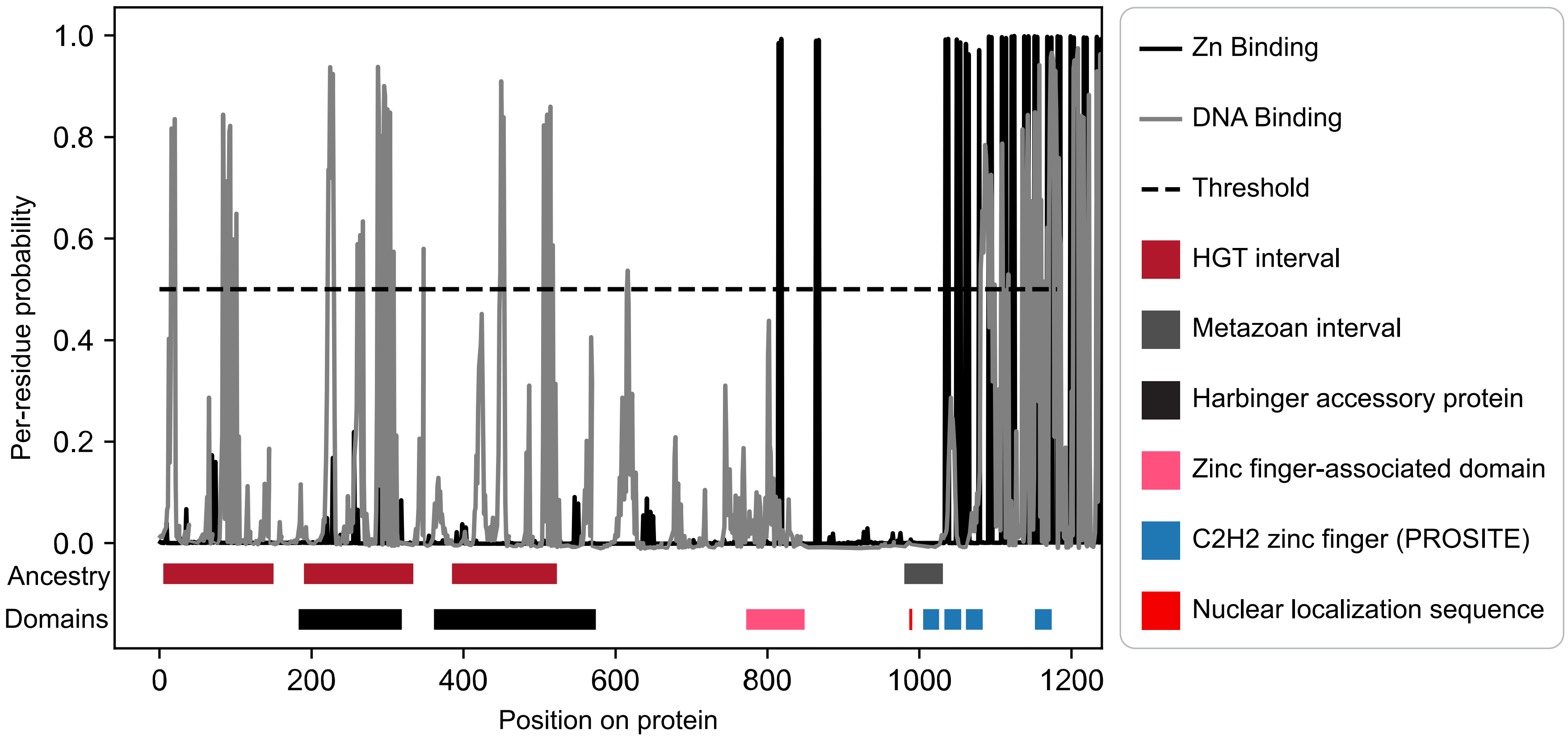


**Figure S13. Evidence of DNA-binding functionality in the HGT and metazoan intervals of XP_021699539.1 (cluster 9) from *Aedes aegypti*.** The protein sequence of XP_021699539.1 was submitted to the GPSite (42) webserver, split into two regions (1-1024 and 1024-1181) due to sequence length limits. Both regions returned high protein-wide probabilities of DNA binding (0.90 and 0.97, respectively) and zinc ion binding (0.99 and 1.00, respectively). Plot shows ancestry inferences, domain architecture predictions, and the per-residue GPSite binding probability for zinc and DNA (significant probabilities > 0.50). We find significant probabilities of DNA-binding in both metazoan and HGT intervals, along with significant zinc binding probabilities at the C-terminus. Consistently, domain architecture predictions via InterProScan revealed intact zinc finger motifs in the C-terminal region, alongside an accessory domain often found in metazoan zinc finger transcription factors (zinc finger-associated domain) (114) . We hypothesize that the entire C-terminus (from 600 amino acids onwards) is of metazoan origin but was trimmed by the interval demarcation procedure due to a low density of BLASTp hits. The N-terminal HGT intervals overlap with fungal Harbinger transposons as annotated by CENSOR(34) .

### Supporting Information Table Legends (separate files)

**SI Table 1. Pipeline inputs.** This spreadsheet file contains two tabs as follows:

***RefSeq genomes*** contains accessions and associated metadata for the 319 RefSeq-annotated genomes used for the original inputs to the pipeline. BUSCO completeness data was obtained directly from the NCBI genomes (https://www.ncbi.nlm.nih.gov/datasets/genome/); blank values indicate that BUSCO scores were not provided on NCBI. Associated with SI Figure 4.

***GenBank genomes*** contains accessions and associated metadata for the additional 197 GenBank genomes used for phylogenetic dataset construction and phylostratigraphy.

**SI Table 2. HGT and metazoan interval annotations.** "HGT_interval" or "Metazoan_interval" columns (note that some HGT-chimeras have multiple HGT and/or metazoan intervals) for all proposed HGT-chimeras along with associated taxonomic information and assignment to similarity clusters. Each row corresponds to a single HGT-chimera protein GenBank or RefSeq accession, determined either through direct inference or through the secondary HGT-chimera search (see SI Methods). Note that some genes (gene accessions in the "gene" column) may have multiple protein isoforms. The "representative" column indicates whether the HGT-chimera was a representative sequence for one of the 104 HGT-chimera similarity clusters ("cluster" column). The total of 274 HGT-chimera genes reported in this study are tabulated by consideration of unique cluster ID-species pairs for all proteins derived from RefSeq genome annotations. This number rises to 348 when also considering GenBank-derived protein accessions, and to 662 when considering all protein isoforms across all genome annotation types. Associated with Figures 1D and SI Figure 4.

**SI Table 3. Number of HGT-chimera similarity clusters found per genome.** Each row represents a single species. "in_primary_search_set" indicates whether the species had a RefSeq genome included in the initial search of proteins from 319 genomes; FALSE indicates that the species only has a GenBank genome annotation considered in the search for secondary HGT-chimera sequences. "n_chimera_clusters" reports the number of unique HGT-chimera clusters found within the genome (raw data in SI Table 2); "n_chimera_genes" reports the number of unique chimeric genes (potentially including multiple genes per similarity cluster) (raw data in SI Table 2). SI Figure 4B was produced from the "n_chimera_clusters" column.

**SI Table 4. Distribution of each HGT-chimera cluster across species.** "cluster_id" indicates an arbitrary identifier assigned to each of 104 HGT-chimera clusters; "representative_chimera" indicates the RefSeq accession of the representative HGT-chimera protein per cluster; "n_species" is the number of unique species the HGT-chimera similarity cluster is found in (including both RefSeq and GenBank genomes); "span_name" is the name of the lowest taxonomic group in the NCBI taxonomy database that encompasses all species containing a representative of the HGT-chimera cluster; "span_taxid" is the NCBI taxonomic identifier for "span_name"; "span_rank" is the NCBI taxonomic rank for "span_name". Associated with SI Figure 4C and SI Figure 5.

**SI Table 5. Ages of HGT-chimeras.** The spreadsheet file contains two tabs as follows:

***Data*** contains minimum age estimates for HGT-chimeras derived using phylostratigraphy for 23 HGT-chimera similarity clusters found in more than one species. "span_name" is the name of the lowest taxonomic group in the NCBI taxonomy database that encompasses all species containing a representative of the HGT-chimera; “all_species” is the species used for phylostratigraphic dating; “Minimum age (MYA)” is the minimum age of the HGT-chimera cluster, defined as the estimated divergence time of all the species in the “all_species” column in millions of years; “Reference #” indicates literature citations in the “References” tab; values for rows lacking a numerical citation in this column were obtained from TimeTree 5 (105).

***References*** contains citations to supporting literature.

**SI Table 6. Gene copy number per genome for each HGT-chimera.** Tabulation in the “n_genes” column, with source data and gene accessions in SI Table 2.

**SI Table 7. Data supporting taxonomic origins of HGT-chimeras**. Each row contains supporting data for a HGT-chimera interval indicated in the column “interval_name”, with intervals named according to the format “genome_accession; protein_accession; HGT_(interval_start_coordinate,interval_stop_coordinate)”. The following columns contain data from tree-based inference: “tree_Note” contains any unusual features of the tree used for HGT inference; “tree_donor”: donor taxon inferred origin at the domain/kingdom level as described in SI Methods (associated with Figure 1C); “tree_sister_kingdom” and “tree_cousin_kingdom”: taxonomic origin of the most closely related non-arthropod sequence(s) and the second-most closely related non-arthropod sequence(s), respectively (blank if absent from the tree); “sister_species” and “cousin_species”: the species of the sister and cousin sequences; “symbiont_relatives”: any species of sister or cousin sequences from bacterial genera in the Symbiotic Genomes Database (92). The following columns contain data from DIAMOND BLASTp-based inference: “blast_seq”: sequence accession and description of the top non-metazoan hit; “blast_species”: the species of “blast_seq”; “blast_tax_label”: the taxonomic domain or kingdom of “blast_seq”; “blast_evalue”: the E-value of “blast_seq”; “blast_cov”: the percent length coverage of the query interval by the alignment with “blast_seq”; “blast_p_HGT300”: the percentage of non-metazoan NCBI taxonomic IDs among the top 300 DIAMOND BLASTp hits; “blast_AI”: the alienness index (see SI Methods). The following columns contain data from hmmsearch: “hmmer_seq”: the sequence accession of the top non-metazoan hmmsearch hit; “hmmer_species”: the species of “hmmer_seq”; “hmmer_kingdom”: the taxonomic kingdom/domain of “hmmer_species”; “hmmer_i-Evale”: the independent E-value for the hmmer alignment of the query profile for the HGT interval and “hmmer_seq.” Rows missing hmmsearch data correspond to intervals searched via DIAMOND BLASTp alone because they lacked non-self arthropod hits with bitscore greater than that of the top non-arthropod hits (see SI Methods).

**SI Table 8. Plausible ecological associations with HGT donors.** The spreadsheet file contains three tabs as follows:

***ecological_associations*** contains inferred associations aside from bacterial symbioses, with the following columns: “interval” is the HGT interval named according to “genome_accession;protein_accession; HGT_(interval_start_coordinate, interval_stop_coordinate)”; “species” is the species of the representative HGT-chimera; “donor” is the inferred donor organism; “evidence” indicates whether the donor is supported by maximum likelihood trees (“ML”), DIAMOND BLASTp (“blast”) or hmmsearch “hmmer”; “donor relationship” is the hypothesized relationship, with parenthetical citations to supporting literature in the “***references***” tab. Relationships without parenthetical citations were inferred directly from the “Source” description of the BLAST/hmmer/tree-inferred relative in SI Table 7 on the GenBank web browser, or from the associated NCBI BioProject or BioSample pages (for example, viral sequences isolated from arthropod tissue were hypothesized to be arthropod-infecting viruses).

***symbionts*** contains data on the presence of arthropod bacterial symbionts obtained from the Symbiotic Genomes Database (92) in the sister and cousin sequences of HGT-intervals, with the following columns: “name”: genus name; “node”: NCBI taxonomic IDs of “name”; “inters”: a list of HGT interval names with relatives in each genus; “n_inters”: the number of intervals with relatives.

***references*** contains citations to supporting literature referred to in the “***ecological_associations***” tab.

**SI Table 9. Results of searches for HGT-chimera relative sequences in genomes with HGT-chimeras.** Results of within-genome BLASTp and tree-based searches for relative sequences, with supporting data for origin via tandem duplication or retroduplication. “interval” indicates the HGT-chimera interval identifier as in SI Table 8, but with a “Meta” string in place of “HGT” for metazoan-derived intervals; “gene_loc” indicates the accession of the chimeric gene. The following columns contain data on the minimum non-chimeric E-value hit gene via BLASTp (see SI Methods): “min_eval” indicates the E-value of the hit; “min_eval_protein” indicates the accession of the protein hit; “min_eval_loc” indicates its gene accession. The following columns pertain to the closest non-chimeric BLASTp hit (via minimum number of intervening genes) on the same chromosome as the query HGT-chimera; cells are blank if no BLASTp hit was detected on the same chromosome: “min_gene_distance” is the number of intervening genes between the HGT-chimera and its non-chimeric hit; “min_gene_distance_loc” is the gene accession of the hit; “min_gene_distance_protein” is the RefSeq protein accession; “min_gene_distance_eval” is the E-value of the hit; “tandem” indicates whether the HGT-chimera was hypothesized to be a potential outcome of tandem duplication. The following columns pertain to analysis of exon counts for inference of retroduplication: “n_exons” indicates the number of exons spanning the interval in the chimeric gene; “relative_n_exons” indicates the number of exons spanning the aligned interval in the minimum E-value hit (“min_eval_protein”/”min_eval_gene”); “retro” indicates whether the HGT-chimera was hypothesized to be a potential outcome of retroduplication. The following columns contain data from tree-based parent inference: “tree_relative_protein” is the protein accession of the parent; “tree_relative_gene” is its gene accession. “FALSE” in both columns indicates that no hit was detected. Note that tree relatives were inferred in six cases where BLASTp failed to return any relatives because of the varying E-value cutoffs and search methodologies (BLASTp vs hmmsearch) in the two methods (see SI Methods). Hence, the final six rows of the table are blank for BLASTp-based inference of gene distance or retroduplication. Associated with SI Text 4 and SI Figure 9.

**SI Table 10. Results of searches for HGT-chimera relative sequences in genomes without HGT-chimeras.** Results of tree-based search for non-chimeric relatives of HGT-derived sequences in arthropod genomes that lack HGT-chimeras. “interval”: the HGT-chimera interval identifier as in SI Table 8; “non_chimeric_accessions”: protein accessions of non-chimeric relatives of HGT-chimeras, for cases where such relatives were found (cell is blank if the value of “n_non_chimera_species” is zero); “non_chimeric_species”: species of the non-chimeric hits, when found; “n_non_chimera_species”: number of species in “non_chimeric_species.” Associated with SI Text 4.

**SI Table 11. HGT-chimera GC content and codon use statistics.** Separated by metazoan and HGT intervals, by interval. “MILC”: Measure Independent of Length and Composition, a codon bias metric (109); “MILC_percen” percentile-normalized MILC values using a within-genome sampled distribution (see SI Methods); “GC”: GC content; “GC_percen”: percentile-normalized MILC values using a within-genome sampled distribution (see SI Methods). Associated with SI Figure 8A-B.

**SI Table 12. HGT-chimera transcription: RNA-Seq analysis.** RNA-Seq expression data for predicted HGT-chimeras from the NCBI genome browser. “cluster_id” refers to the similarity cluster identifier as in SI Table 4; “protein_name” is the RefSeq accession of the representative protein queried for detectable expression; “pct_ab_initio” is the percentage by length of the gene model not supported by RNA-Seq; “CDS_support” reports “full” if the complete coding region corresponding to “protein_name” is supported by RNA-Seq (if not, cell is blank); “HGT_support” and “Metazoan_support” provide RNA-Seq evidence (based on manual inspection of the RefSeq genome browser) for transcription of HGT and metazoan intervals (SI Table 2) when CDS_support is not full. The latter two columns are reported as “Yes” if 100% of the intervals have RNA-Seq coverage, “Partial” if coverage is >50%, or “No” otherwise.

**SI Table 13. HGT-chimera transcription: RT-PCR analysis.** RT-PCR data on HGT-chimera expression. The spreadsheet file contains three tabs as follows:

***sequencing_results*** contains the results of PCR and Sanger sequencing, with the following column names: “sequence name”: uniquely identifies each sequencing product with the format “species prefix_chimera id_seqnumber.” Each HGT-chimera has a unique species prefix_chimera id combination. Note that four HGT-chimeras (blue text in “sequencing results”: XM_045171894 from *Daphnia magna*, KAH9406650 from *Tyrophagus putrescentiae*, XM_035852275 from *Folsomia candida*, and XM_023473825 from *Eurytemora carolleeae*) have multiple sequencing products, in which case “prefix_chimera id” are identical but the sequence number varies. For example, *Typutr_chimera1_seq1* and *Typutr_chimera1_seq2* are different products of the same HGT-chimera obtained via cloning when Sanger sequencing suggested the presence of multiple allelic or splice form variants (see SI Methods); “genbank accession”: accession of coding and protein sequences submitted to GenBank. In this column, “None obtained” means that no interpretable Sanger sequencing product was obtained, so this was considered as a negative result and not submitted to GenBank; “reference protein accession”: NCBI protein accession of HGT-chimera that appeared from the screen; “reference mRNA sequence used for primer design”: nucleotide sequence used for primer design. Sequences not from RefSeq (i.e. not beginning with XM) were obtained by navigating to the protein page in GenBank and clicking on “CDS”; “species”: species name of HGT-chimera source; “chimera cluster #”: HGT-chimera similarity cluster ID number; “PCR bands obtained”: bands of the expected length obtained via RT-PCR; “Sequence obtained”: interpretable Sanger sequencing results obtained; “Chimera confirmed”: sequencing results support existence of HGT and metazoan intervals in a contiguous transcript without evidence of reading frame disruption or premature stop codons relative to the reference protein.

***primers*** contains primer data for RT-PCR, with the following column names: ”sequence_names”: unique sequencing product(s) obtained using the primer pair. As indicated in ***sequencing_results***, some primer pairs failed to produce PCR bands or interpretable sequencing products; “Forward”: Forward primer sequence; “Reverse”: Reverse primer sequence; “Annealing Temp [°C]”: annealing temperature for PCR reaction; “Extension Time [sec]”: Extension time for PCR reaction. We note that four HGT-chimera sequences were assembled from multiple overlapping PCR-products obtained with unique primer pairs (green text in “primers”: Aeaegy_chimera1_seq1 from *Aedes aegypti,* Aealbo_chimera1_seq1 from *Aedes albopictus*, Dapule_chimera1_seq1 from *Daphnia pulex*, Dapuli_chimera2_seq1 from

*Daphnia pulicaria*). Distinct sequence names from the same species with identical primer sequences correspond to cloning variants as in "***sequencing_results***". * indicates conditions for KOD One Polymerase.

***sample data*:** contains data for the sources of tissue samples for cDNA preparation, with the following column names: "Species name": Linnaean name, consistent with the NCBI taxonomy database; "Collection date": date when specimens were harvested; "Harvest method": method of tissue preservation prior to RNA extraction (LN2: flash-frozen in liquid nitrogen; Frozen: stored at -80 ºC); "geo_loc_name": location where specimens were obtained from; "Country: State (when available), city"; "Institution": University or company affiliation of the individuals responsible for specimen collection; "Collector": names of individuals who collected the specimens; "Dev-stage": Developmental stage of specimens; "Sex": sex of specimens if known; "Tissue type": tissue of specimens; "Isolation-source": information on colony or strain of the samples; "Shipping conditions": conditions that samples were shipped in.

**SI Table 14. Gene-wide dN/dS.** Values fitted with M0 in PAML (57, 58, 95). The "_f" suffix indicates that the parameter values were not fixed, i.e., estimated using maximum likelihood. "_1" suffix indicates dN/dS was fixed at 1. "lnL": log likelihood; "dnds": ratio of nonsynonymous to synonymous substitution rates; "dS": indicates total branch length in synonymous substitutions/site across the tree; "np" refers to the number of parameters in each model, "delta_p" is the change in number of parameters between the free parameter "_f" and null "_1" models; "p_val" obtained using a likelihood ratio test (chi-squared with "delta_p" degrees of freedom and statistic "2delta_L"); "p_val_corrected" with the Benjamini-Hochberg procedure. Includes data for all 23 HGT-chimera clusters found in more than one species, including clusters 3 and 18, which are excluded from Figure 2E.

**SI Table 15. dN/dS estimates partitioned by HGT vs. non-HGT ancestry with fixed-site models.** Suffix "2" indicates codons from HGT regions from model Mgene=4 in which dN/dS varies across sites; suffix "_f" indicates model fit with Mgene=2, in which dN/dS does not vary across sites; "nsites_total" is the total number of codons analyzed and "nsites_2" is the number of HGT codons in the interval (both after removing codons with gaps in the codon alignment). Remaining column descriptions are the same as in SI Table 14. Associated with Figure 2F.

**SI Table 16. Test of different dN/dS values across chimeric and non-chimeric branches.** "dnds_b0": background branch (non-chimeric); "dnds_b1": foreground branch (chimeric); "_f suffix": single dN/dS ratio for all branches; "_b" suffix for the two-branch test. Remaining column descriptions are the same as in SI Table 14. Associated with Figure 3E-F. We caution that dN/dS estimates may lose power to detect positive selection if dS values saturate at greater evolutionary distances (115), but note that our application of tree-based codon models (57, 58) and exclusive consideration of genus- or species-specific HGT-chimeras and their within-order relatives at least partially mitigates this concern. Associated with Figure 4F.

**SI Table 17. Functional predictions for HGT-chimeras.** Partitioned by interval. The spreadsheet file contains five tabs as follows:

The first four tabs (**GO_MF_HGT**, **GO_MF_Meta**, **GO_bp_HGT**, **GO_bp_Meta**) contain tabulations of gene ontology (GO) molecular function (MF) and biological process (bp) predictions for separated metazoan (Meta) and HGT intervals obtained via DeepGO-SE. "n_intervals" in each of these sheets tabulates the total number of intervals across in the given interval type with each GO prediction.

***combined_functions_per_interval*** compiles functional predictions across NCBI CDD search (“cdd”), InterProScan (“ipr”), DeepGO-SE (“go_mf” and “go_bp”), as well as any RepBase transposable element annotations obtained via CENSOR (“RepBase_TE”) overlapping with each interval.

**SI Table 18. Putative functional similarity between metazoan and HGT domains of HGT-chimeras.** HGT-chimeras with evidence of predicted functional similarity between metazoan and HGT intervals. Columns: “representative chimera” and “representative species” of the cluster representative analyzed for functional similarity; “HGT interval(s)” and “Metazoan interval(s)” contain predicted domains/functions for the HGT and metazoan-derived interval(s) of the HGT-chimera; “function evidence” columns indicate whether the functional inference presented in this table was supported by the NCBI Conserved Domain Database (“CDD”), InterProScan (“IPR”), DeepGO-SE, DIAMOND BLASTp, and/or CENSOR/RepBase, with source data in SI Table 17; “Predicted functional commonality” indicates the inferred functional similarity among the HGT and metazoan derived intervals. Rows highlighted in yellow have nucleic-acid associated functions in both interval types, while those highlighted in green have carbohydrate-associated functions in common. Functional similarity was inferred directly from domain descriptions on the InterPro or CDD websites, or from the references cited in parentheses and listed in the “**References**” tab.

**SI Table 19. Whole-protein HGT inference results.** Results of whole-protein HGT inference (without partitioning into intervals) on 104 HGT-chimera representatives, using the same criteria as round 2 BLASTp-based inference for demarcated intervals. “annotation”: ‘HGT’ if HGT-ancestry is inferred, ‘Meta’ if metazoan ancestry is inferred, ‘None’ if ancestry is ambiguous.

**Supporting Information Files Legends (separate files)**

**SI File 1.** Plots of round 1 and round 2 BLASTp results, for 104 representative HGT-chimera sequences. Alignments are shown as horizontal lines with coordinates on the x-axis indicating the start and end of the amino acid alignment on the query HGT-chimera sequence, and the y-axis representing the log-transformed E-value of each hit. Hits are colored according to their taxonomic origin, and thick orange or purple boxes at the bottom of each plot represent demarcated metazoan or HGT intervals, respectively. These plots are associated with Figure 1A and SI Figure 2.

**SI File 2.** Alignments of the 36 predicted HGT-chimera proteins for which we confirmed expression via RT-PCR, with the amino acid translations of their Sanger sequenced mRNA product(s). See SI Table 13.

### Supporting Information References

59. Schrödinger, LLC, *The PyMOL Molecular Graphics System, Version 3.0*.

96. Z. Yang, *Computational Molecular Evolution* (Oxford University Press, 2006).
