## Supplementary material for "Evolutionary innovation through fusion of sequences from across the tree of life": ST Tables 1 through 21: HGTc_SI table 20_v5.docx

**SI Table 20. Comparison of pipeline iterations 1 and 2.** An initial iteration (iteration 1) of the HGT-chimera detection pipeline was developed and implemented between 2022-2024, before the updates described in this table were implemented in 2025 prior to deposition of this study. The second pipeline iteration (iteration 2), as explained in detail in SI Methods, is independent of the first, except for the addition of 11 RT-PCR validated chimeras detected in iteration 1 to the clustered protein set searched here.

| **Feature** | **Pipeline Iteration 1 (previous)** | **Pipeline Iteration 2 (current)** |
| --- | --- | --- |
| Input RefSeq Genomes | 237 (accessed 1/2023) | 319 (Accessed 4/2025) |
| Input MMSeqs2-clustered proteins | N= 472,250 | N= 610,348 + 11 RT-PCR-validated chimeras recovered from Iteration 1 |
| BLASTp and hmmsearch database | NR (downloaded 12/2022) | NR (downloaded 10/2023) |
| E-value threshold for round 2 BLASTp-based HGT inference | 1e-1 | 1e-4 OR bit-score >50 (calibrated to previous study of Oskar (Blondel, Jones, and Extavour 2020) |
| Automated filter to remove chimeras in which the alignment of non-arthropod BLAST hits to metazoan intervals partially extend into HGT intervals or vice-a-versa | Present | Absent. Found to be redundant with round 2 BLASTp filtering and manual BLASTp plot inspection (last step of Phase III in SI Figure 1). |
| Ankyrin repeat filter | Absent | Present |
| Transposable element filter | Present, but by manual inspection of CENSOR/RepBase hits | Present, with automated thresholding from CENSOR/RepBase outputs |
| Expanded secondary chimera search database | NR database | Custom database of 319 RefSeq + 197 GenBank genomes |
| Phylogenetic tree construction from hmmsearch hits | Used for all intervals | Used only for intervals for which ≥ 2 BLASTp hits with bitscore ≥ max(non-arthropod bit-score), else used BLASTp hits directly |
| Bootstrap filter for clades of arthropod + non-metazoan sequences for HGT inference | Absent | >70% |
| Final number of HGT-chimera clusters | 50 | 104 |
