## Supplementary material for "Evolutionary innovation through fusion of sequences from across the tree of life": ST Tables 1 through 21: HGTc_SI table 21_v2.docx

**SI Table 21**: Source genomes for ten species with inferred representatives of HGT-chimera cluster 3 (SI Figure 7, SI Text 1). “Submitting Institution” was obtained from the Submitter field of the NCBI Genome page (<https://www.ncbi.nlm.nih.gov/datasets/genome>). We note that, with the exception of the genomes of two mites (*Oppiella nova* and *Oppia nitens*) that were both submitted by the University of Lausanne, all other genomes were submitted by labs at different institutions.

| **Species** | **Class** | **Accession** | **Submitting Institution** |
| --- | --- | --- | --- |
| *Oppia nitens* | Arachnida (order: Sarcoptiformes) | GCF_028296485.1 | University of Saskatchewan |
| *Tyrophagus putrescentiae* | Arachnida (order: Sarcoptiformes) | GCA_021730765.1 | Chinese University of Hong Kong |
| *Oppiella nova* | Arachnida (order: Sarcoptiformes) | GCA_905397405.1 | University of Lausanne |
| *Medioppia subpectinata* | Arachnida (order: Sarcoptiformes) | GCA_905368565.1 | University of Lausanne |
| *Blomia tropicalis* | Arachnida (order: Sarcoptiformes) | GCA_029204025.1 | Affiliated Wuxi People's Hospital of Nanjing Medical University |
| *Folsomia candida* | Collembola | GCF_002217175.1 | VU Amsterdam |
| *Allacma fusca* | Collembola | GCA_910591605.1 | University of Edinburgh |
| *Bradysia coprophila* | Insecta (family: Sciaridae) | GCF_014529535.1 | Brown University |
| *Bradysia odoriphaga* | Insecta (family: Sciaridae) | GCA_016920775.1 | Hebei Agricultural University |
| *Pseudolycoriella hygida* | Insecta (family: Sciaridae) | GCA_029228625.1 | University of Sao Paulo |
